# An adversarial collaboration to critically evaluate theories of consciousness

**DOI:** 10.1101/2023.06.23.546249

**Authors:** Cogitate Consortium, Oscar Ferrante, Urszula Gorska-Klimowska, Simon Henin, Rony Hirschhorn, Aya Khalaf, Alex Lepauvre, Ling Liu, David Richter, Yamil Vidal, Niccolò Bonacchi, Tanya Brown, Praveen Sripad, Marcelo Armendariz, Katarina Bendtz, Tara Ghafari, Dorottya Hetenyi, Jay Jeschke, Csaba Kozma, David R. Mazumder, Stephanie Montenegro, Alia Seedat, Abdelrahman Sharafeldin, Shujun Yang, Sylvain Baillet, David J. Chalmers, Radoslaw M. Cichy, Francis Fallon, Theofanis I. Panagiotaropoulos, Hal Blumenfeld, Floris P de Lange, Sasha Devore, Ole Jensen, Gabriel Kreiman, Huan Luo, Melanie Boly, Stanislas Dehaene, Christof Koch, Giulio Tononi, Michael Pitts, Liad Mudrik, Lucia Melloni

**Author notes:** Equally contributing co-first authors (Oscar Ferrante, Urszula Gorska-Klimowska, Simon Henin, Rony Hirschhorn, Aya Khalaf, Alex Lepauvre, Ling Liu, David Richter, Yamil Vidal). Equally contributing co-senior authors (Michael Pitts, Liad Mudrik, Lucia Melloni).

## Abstract

Different theories explain how subjective experience arises from brain activity^1,2^. These theories have independently accrued evidence, yet, confirmation bias and dependence on design choices hamper progress in the field^3^. Here, we present an open science adversarial collaboration which directly juxtaposes Integrated Information Theory (IIT)^4,5^ and Global Neuronal Workspace Theory (GNWT)^6–10^, employing a theory-neutral consortium approach^11,12^. We investigate neural correlates of the content and duration of visual experience. The theory proponents and the consortium developed and preregistered the experimental design, divergent predictions, expected outcomes, and their interpretation^12^. 256 human subjects viewed suprathreshold stimuli for variable durations while neural activity was measured with functional magnetic resonance imaging, magnetoencephalography, and electrocorticography. We find information about conscious content in visual, ventro-temporal and inferior frontal cortex, with sustained responses in occipital and lateral temporal cortex reflecting stimulus duration, and content-specific synchronization between frontal and early visual areas. These results confirm some predictions of IIT and GNWT, while substantially challenging both theories: for IIT, a lack of sustained synchronization within posterior cortex contradicts the claim that network connectivity specifies consciousness. GNWT is challenged by the general lack of ignition at stimulus offset and limited representation of certain conscious dimensions in prefrontal cortex. Beyond challenging the theories themselves, we present an alternative approach to advance cognitive neuroscience through a principled, theory-driven, collaborative effort. We highlight the challenges to change people’s mind ^13^ and the need for a quantitative framework integrating evidence for systematic theory testing and building.

## Main

Philosophers and scientists have sought to explain the subjective nature of consciousness (e.g., the feeling of pain or of seeing a colorful rainbow) and how it relates to physical processes in the brain^14,15^. This “explanatory gap” ^16^ or “hard problem” ^17^ has led to several competing theories of consciousness that have evolved in parallel^1–3^. Yet, those theories offer incompatible accounts of the neural basis of consciousness^1,2^. Moreover, empirical support for a given theory is often highly dependent upon methodological choices, pointing towards a confirmation bias when testing these theories^3^. Convergence upon a broadly accepted neuroscientific theory of consciousness will have profound medical, societal, and ethical implications.

With this goal in mind, we take an unusually concerted effort to testing theories of consciousness: a large-scale, open science, adversarial collaboration^11,12,18–20^ aimed at accelerating progress in consciousness research by building upon constructive disagreement. Our collaboration brings together proponents of Integrated Information Theory (IIT)^4,5^ and Global Neuronal Workspace Theory (GNWT)^6,21^, two of the most well-established theories in the field, in addition to theory neutral researchers. Together, we identified divergent predictions of the theories and jointly developed an experimental design to test them (Figure 1a-b). We preregistered foundational and novel predictions from the two theories, including pass/fail criteria for each prediction, as well as expected outcomes and their interpretation ex-ante^11,12^.

**Figure 1:**
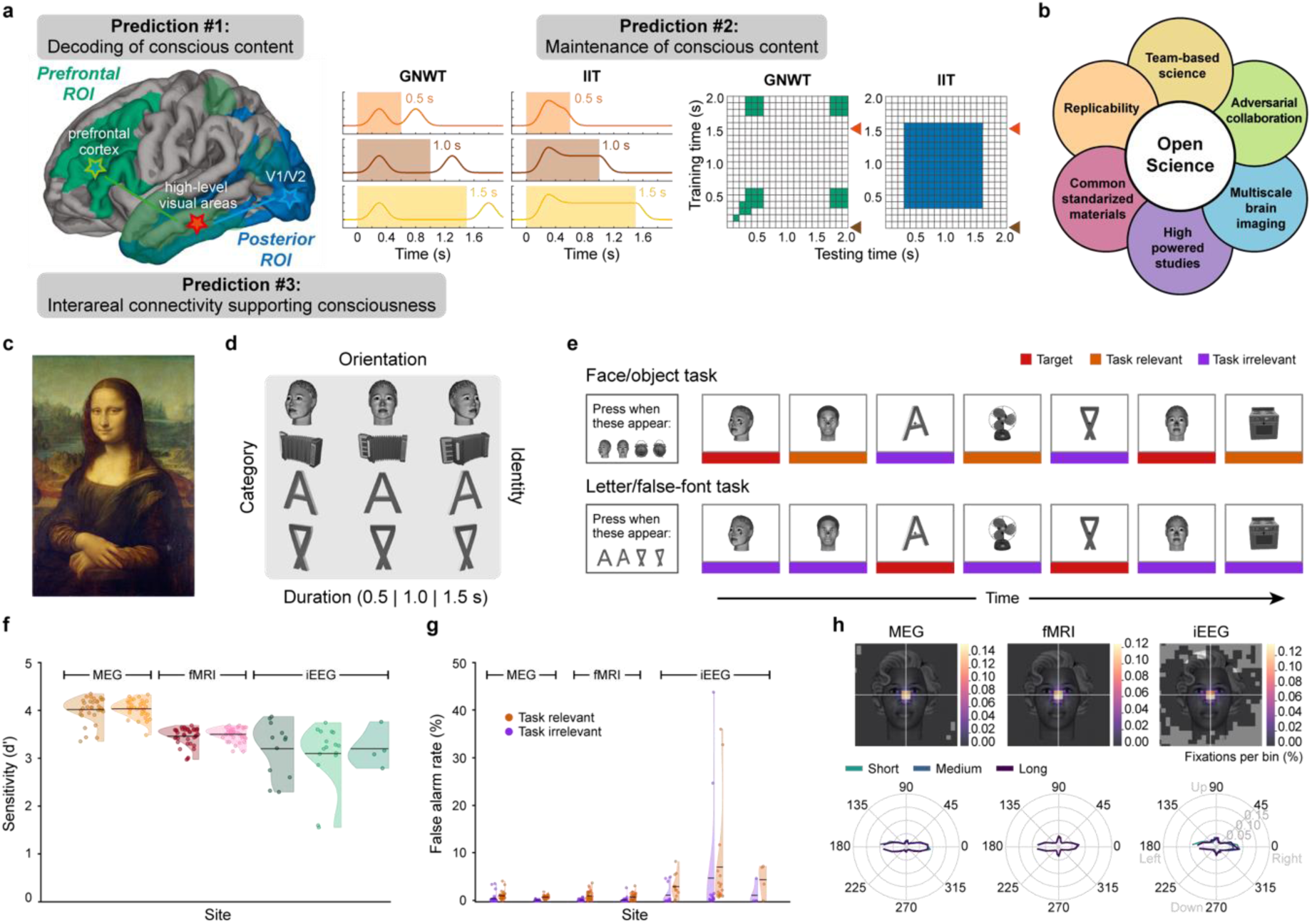
Theories predictions tested in an adversarial collaboration. **a.** Three key contrasting predictions of Integrated Information Theory (IIT) and Global Neuronal Workspace Theory (GNWT) tested in an adversarial collaboration framework. **Prediction #1:** Decoding of conscious content, evaluating which cortical areas hold information about different aspects of conscious content. IIT predicts that conscious content is maximal in posterior brain areas, while GNWT predicts a necessary role for PFC. **Prediction #2:** Maintenance of conscious content over time, evaluating the temporal dynamics by which the temporal extent of the conscious content is instantiated. IIT predicts that conscious content is actively maintained in posterior cortex throughout the extent of a conscious experience; while GNWT predicts brief content-specific ignition in PFC ∼0.3-0.5 s after stimulus onset and offset (when the workspace is updated), with content stored in a non-conscious silent state resembling activity-silent working memory in between. Waveforms and temporal generalization matrices depict the amplitude- and information-based temporal profiles predicted by the theories, respectively (left: colored rectangles indicate the three different stimulus durations, GNWT predicted waveforms pertain to PFC, IIT predictions to posterior cortex; right: brown arrows indicate stimulus onset, red arrows stimulus offset, green and blue colors reflect the predicted patterns of temporal generalization of conscious content according to each theory in PFC and posterior cortex for GNWT and IIT, respectively). **Prediction #3:** Interareal communication, evaluating the topological and temporal patterns of interareal connectivity subserving consciousness. The stars and arrows on the brain (left) depict the different predictions about the expected synchrony patterns (green: GNWT; blue: IIT). **b.** The scientific framework under which the Cogitate Consortium (Collaboration On GNW and IIT: Testing Alternative Theories of Experience), tested IIT and GNWT included: adversarial collaboration, (theory-neutral) team science, invasive and non-invasive multimodal data (iEEG, fMRI, MEG), large samples (>250 subjects), standardized protocols across multiple laboratories, built-in preregistered replication, and open methods, data and code. **c.** Conscious experience is multifaceted in content. Looking at the image of Mona Lisa by Leonardo da Vinci underscores the fact that conscious experiences are rich: The painting is experienced as occupying a location in space, pertaining to a given category (i.e., a face and not an object, or any other category), specifying an identify (i.e., Mona Lisa and not any other face), and a particular orientation (i.e., leftward oriented and not rightward or any other orientation). Moreover, the conscious experience is maintained over time for as long as one appreciates the painting, endowing it with a temporal extent (i.e., it feels extended in time). **d.** To experimentally capture the multifaceted aspect of phenomenological experience, we manipulated the content of consciousness by varying stimuli along four dimensions: category (faces, objects, letters and false fonts), identity (each category contained different exemplar), orientation (left, right, and front view), and duration (stimuli were presented for three durations i.e., 0.5 s, 1.0 s, and 1.5 s). Example stimuli used in the study are shown for reference. **e.** Overview of the experimental paradigm: At any one point in time, no more than one high-contrast, stimulus was present at fixation. In each trial, subjects were asked to detect target stimuli: either a face and an object or a letter and a false font in any of the three different orientations. Thus, each trial contained three stimuli types: targets (depicted in red), task relevant stimuli (belonging to the same categories as the targets, depicted in orange-red), and task irrelevant stimuli (belonging to the two other categories, depicted in purple). The pictorial stimuli (faces/objects) were task relevant in half of the trial blocks, while the symbolic stimuli (letters/false fonts) were relevant in the other half of the blocks. For illustration purposes only, a color line was added to depict the different trial types. Blank intervals between stimuli are not depicted here. **f.** Distribution of behavioral sensitivity scores (d’) separate per data modality and acquisition site. Crossing lines depict average d’ per site/modality. Dots depict individual participants d’s. Colors depict data modality: MEG N=65 (orange), fMRI N=73 (red), and iEEG N=32 (green), while the hue depicts each site within a modality. **g.** Distributions of false alarm (FA) rates per site and data modality, separated by task condition: Orange-red depicts task relevant stimuli. Purple depicts task irrelevant stimuli. Dots are individual participants FA rates. Other conventions as in f. **h.** Top row: Average fixations heatmaps computed over a 0.5 s window after stimulus onset. Heatmaps are displayed per data modality, zoomed into the stimulus area. Bottom row: Average saccadic direction maps per data modality. The three stimulus durations are shown separately.

IIT and GNWT explain consciousness differently: IIT proposes that consciousness is the intrinsic ability of a neuronal network to influence itself, as determined by the amount of maximally irreducible integrated information (phi) supported by a network. Theoretical and neuroanatomical considerations indicate that a complex of maximum phi likely resides primarily in the posterior cerebral cortex, in a temporo-parietal-occipital “hot zone”^4,5,22,23^. GNWT instead posits that consciousness arises from global broadcasting and late amplification (or “ignition”) of information across interconnected networks of higher-order sensory, parietal, and especially prefrontal cortex (PFC)^6,9,21^. Although GNWT holds the workspace to include prefrontal cortex and inferior parietal cortex^21^, in this adversarial collaboration we focused on PFC, as GNWT and IIT pose the most incompatible and hence maximally diagnostic predictions about this brain region.

We tested three core contrasting predictions of IIT and GNWT for how the brain enables conscious experience. **Prediction #1** pertains to brain areas in which conscious content should be found. IIT predicts that conscious content is instantiated primarily in posterior brain areas, while GNWT predicts a necessary role for PFC. **Prediction #2** pertains to how conscious percepts are maintained over time^24,25^: IIT predicts that conscious content is actively maintained by neural activity in the posterior ‘hot zone’ (PHZ) throughout the duration of a conscious experience. GNWT predicts, instead, that an ignition in PFC at stimulus onset, and at offset, updates the workspace, with activity-silent maintenance of information in between ^26^. **Prediction #3** pertains to interareal connectivity between cortical regions during conscious perception. IIT predicts short-range connectivity within posterior cortex, including lower-level sensory (V1/V2) and high-level category-selective areas (e.g., fusiform face area, lateral occipital cortex). In contrast, GNWT predicts long-range connectivity between high-level category-selective areas and PFC. This combination of predictions places a uniquely high bar for either theory to pass, especially considering the highly powered and multimodal studies we conducted. Finally, an additional goal of this experiment was to narrow down the cortical areas potentially participating in consciousness by excluding those reflecting confounding cognitive/task-related processes (***putative NCC analysis below***).

To test our predictions, we investigated the content and temporal extent of conscious visual experiences that are phenomenologically multifaceted and rich, even for a single stimulus. For example, when viewing the Mona Lisa (Figure 1c), one experiences it located in visual space, with a specific identity, a specific orientation, and the experience continues as long as one looks at the painting. To capture the multifaceted aspect of consciousness, we presented suprathreshold stimuli belonging to four different categories (faces, objects, letters, false fonts), with each category containing twenty individual identities presented in three orientations (front, left, right view) for three durations (0.5, 1.0, 1.5 s) (Figure 1d). Participants viewed the stimuli while searching for two infrequent targets, making some stimuli task relevant and others task irrelevant (Figure 1e). See supplementary video depicting the task. This paradigm offers several advantages: first, it provides robust conditions to test the theories’ predictions as it focuses on clearly experienced conscious content, studied through high signal-to-noise, suprathreshold, fully attended stimuli, making any failures of the theories’ predictions more significant. Second, it minimizes task and report confounds, better isolating neural activity specifically related to consciousness. Third, it diverges from the theories’ usual testing grounds to probe new predictions regarding how experience is maintained over time, making the results more informative.

All research was conducted by theory-neutral teams to guard against confirmatory bias. We evaluated the theories’ predictions in 256 subjects who performed the same behavioral task in three different neuroimaging modalities: functional magnetic resonance imaging (fMRI, *N*=120), magnetoencephalography (MEG, *N*=102), and intracranial electroencephalography (iEEG, *N*=34). The combination of several modalities maximized sensitivity, spatiotemporal resolution, and coverage, thereby providing stringent and comprehensive tests of the theories. Furthermore, each data type was collected by two (or three) independent laboratories to ensure generalization across populations, recording systems, and experimenters. Altogether, we aimed at fostering informativeness, reproducibility, and robustness of the results by (1) dissociating theory from data acquisition/analysis to prevent biases, (2) using a multimodal approach to test theories with an exquisite temporal and spatial precision, (3) acquiring data in a large sample of subjects to increase statistical power, (4) using standardized and preregistered protocols^12^ to evaluate theories under the same experimental framework and further minimize confirmatory bias^20^, and finally (5) combining an analysis optimization phase with a final testing phase using independent parts of our dataset to corroborate the robustness of the results (Figure 1b)^27^. Consequently, we present a large-scale international effort to evaluate two leading theories of consciousness under an integrated, rigorous and comprehensive adversarial collaboration framework, setting a precedent for theory testing.

We first established that our task manipulations were effective and comparable behaviorally across data modalities and sites (see supplementary for the full set of results). Subjects’ performance in the task was excellent, with high hit rates (M=96.84%, SD=4.19%), low false alarm rates (M=1.45%, SD=4.30%), and excellent fixation stability (mean accuracy <2°=89.62%, SD=10.61%; Figure 1f-h). Subjects’ performance across laboratories within each data modality was similar (all p=1.000 after multiple comparison correction, BF<0.12). Epilepsy patients showed slightly lower behavioral performance compared to neurotypical subjects, yet, behavior was still comparatively high (hit rate 93.90%, SD=12.29; false alarm rate M=4.25%, SD=20.17). We confirmed that subjects were conscious of the stimuli both in the task relevant and irrelevant trials in a separate experiment which included a surprise memory test (see supplementary).

As part of our testing framework, after excluding a limited number of subjects due to data quality checks, we conducted an initial optimization phase on 1/3 of the MEG (N=32) and fMRI (N=35) datasets to evaluate data quality across sites and to optimize analysis pipelines. Following the optimization phase, pipelines were preregistered (https://osf.io/92tbg/), and applied to the novel datasets containing twice as much data (MEG, N=65 and fMRI, N=73). In what follows we report results obtained on the novel, unseen datasets (see methods for the strategy used for iEEG and text for numbers of subjects that entered in each analysis). Results for all three tested predictions from the optimization phase were largely compatible, with some exceptions, with the replication phase (see supplementary).

### Prediction #1: Decoding of conscious content

According to IIT, information about the content of consciousness should be present primarily in posterior cortical areas, while for GNWT it should require the involvement of PFC. The main discrepancy between the theories is thus the necessity of PFC. IIT and GNWT further specify that brain areas evidencing conscious content should do so irrespective of other cognitive processes, e.g., report. This implies that conscious content should be present irrespective of tasks manipulations^28,29^. To test prediction #1, we evaluated decoding of stimulus category (pictorial: faces/objects and symbolic: letters/false fonts), and orientation (left/right/front facing). In each block, the subjects’ task was to identify two stimuli belonging to either the pictorial or the symbolic group of stimulus categories e.g., a specific face and a specific object (Figure 1e), making these two categories task relevant in that block. Hence, across the studies all categories were both task relevant and task irrelevant. Stimulus orientation was orthogonal to the task, and thus entirely task irrelevant.

Based on our preregistered predictions (https://osf.io/92tbg/), the theories would pass these tests if we observe decoding of one stimulus category pairing (e.g., faces/objects or letters/false fonts) *and* if orientation is decodable in at least one of the four categories, in the relevant brain regions and time windows. Testing for decoding of category and orientation constitutes a more stringent test of the theories as it requires two conditions to be satisfied, while also capturing a critical aspect of conscious content, i.e., its multidimensionality, or phenomenological richness (Figure1d).

For decoding of category, we also sought to demonstrate that information is evidenced irrespective of the task by training a classifier in one task and evaluating whether it generalizes to the other task condition, i.e., cross-task generalization. Here, we report the most robust results for decoding of category (faces/objects) and orientation (left/right/front views of faces). Qualitatively similar results were observed for decoding of letters/false fonts (Extended Data Figure 1a-d). Results for orientation, were consistent across stimulus categories and data modalities in posterior cortex, yet mostly absent in PFC (see supplementary).

In the iEEG data, we trained pattern classifiers on high gamma frequency band activity (70-150 Hz) at each time-point in the task irrelevant condition and tested across all time-points in the task relevant condition, for each stimulus duration, category, and across all electrodes within the theory-relevant ROIs (Figure 2a, Extended Data Table 2 and methods). In posterior cortex, face/object decoding showed significant cross-task generalization (>95% accuracy) for the approximate duration of the stimulus (Figure 2b, top row). In PFC, significant cross-task face/object decoding accuracy (∼70%) was also evident, but the temporal generalization of this decoding was restricted to ∼0.2-0.4 s (Figure 2b, bottom row). Training on task relevant and testing on task irrelevant trials showed similar results (Extended Data Figure 1e; within-task decoding provided in Extended Data Figure 3). The sustained (posterior) and phasic (PFC) patterns of cross-task temporal generalization of decoding thus matched both IIT’s and GNWT’s predictions, respectively.

**Figure 2:**
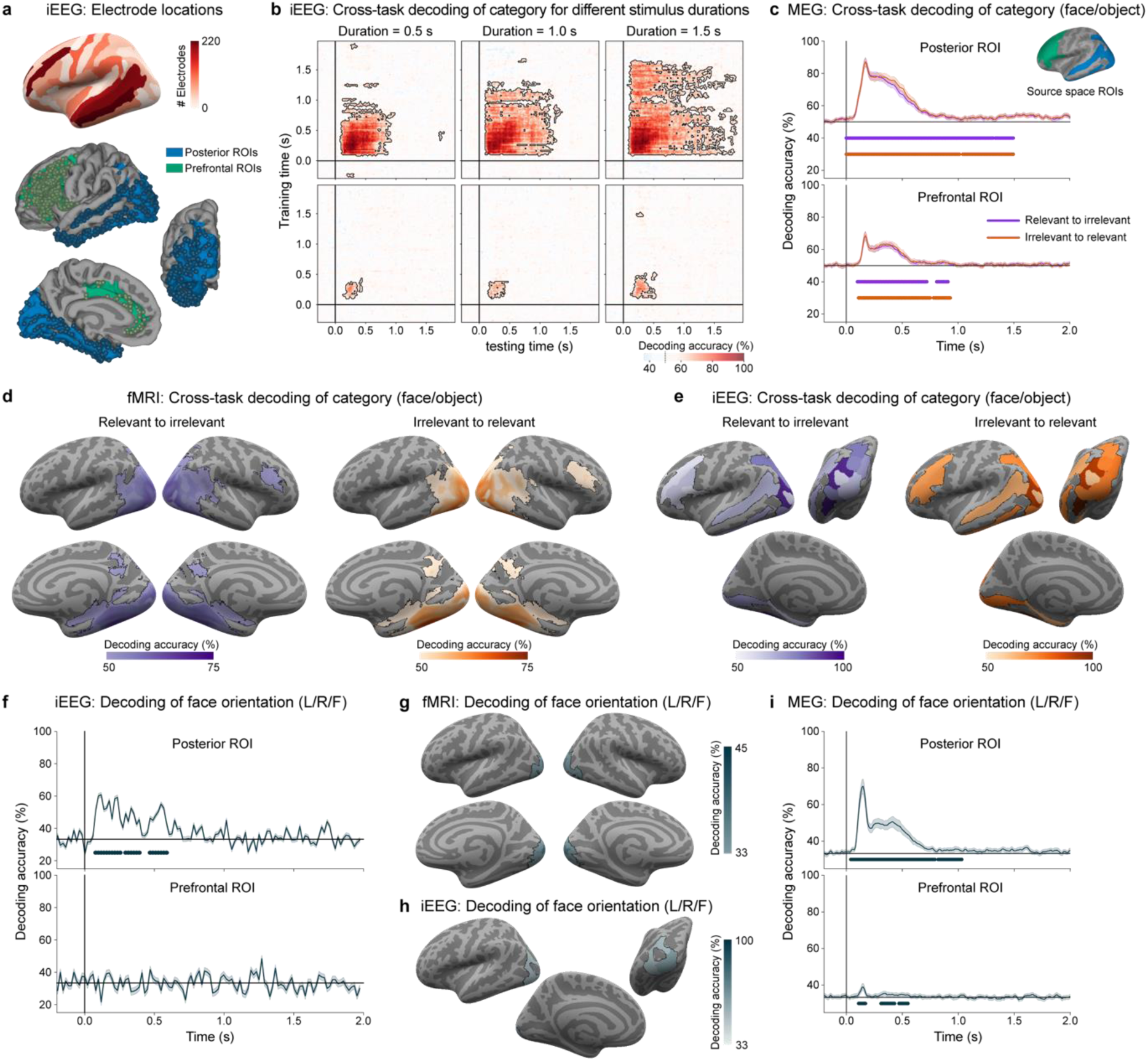
Prediction #1: Decoding of conscious content. **a.** Spatial coverage of intracranial electrodes across all patients included in the decoding analysis (N_subjects_=29), displayed on a standard inflated cortical surface map (top), and within the regions of interest (ROIs) for the two theories (bottom): posterior (blue, N_electrodes_=583), prefrontal (green, N_electrodes_=576). **b.** Cross-task temporal generalization of decoding of high gamma signal in iEEG in which pattern classifiers were trained to discriminate stimulus category (faces vs. objects) in the task irrelevant condition at each time-point and tested in the task relevant condition across all time-points. The three stimulus durations are plotted in columns (left: 0.5 s; center: 1.0 s; right: 1.5 s) and the two theory ROIs in rows (top: posterior ROIs; bottom: prefrontal ROIs). Significantly above-chance (50%) decoding is indicated by the outlined pink-red regions in the temporal generalization matrices. Contours indicate statistically significant decoding evaluated through a cluster-based permutation test. **c.** Cross-task decoding of stimulus category (faces vs. objects) in MEG cortical time series (N=65) when classifiers were trained on relevant stimuli and tested on irrelevant stimuli (purple); or trained on irrelevant stimuli and tested on relevant stimuli (red). Decoding was done separately within the whole posterior ROIs (top) and prefrontal ROIs (bottom). The inset shows inflated cortical surfaces depicting the two theory ROIs (posterior: blue; prefrontal: green) used for decoding. These decoding results combine data across the three stimulus durations, and used pseudotrial aggregation. The purple and red lines underneath the decoding functions indicate time-periods showing significantly above-chance (50%) decoding as assessed by cluster-based permutation test. 95% CI estimate across cross-validation folds. **d.** Cross-task decoding of stimulus category (faces vs. objects) in fMRI (N=73) using a searchlight approach, collapsed across the three stimulus durations. Left panel (purple): Pattern classifiers trained on relevant stimuli and tested on irrelevant stimuli. Right panel (orange-red): Pattern classifiers trained on irrelevant stimuli and tested on relevant stimuli. Regions showing significantly above-chance (50%) decoding, evaluated through a cluster-based permutation test, are indicated by the outlined colored regions on the inflated cortical surfaces (top: left/right lateral views; bottom: right/left medial views). **e.** Cross-task decoding of stimulus category (faces vs. objects) in iEEG within the theory-specific ROIs, collapsed across stimulus duration. Decoding accuracies are indicated in purple for classifiers trained on relevant stimuli and tested on irrelevant stimuli, and in orange-red when trained on irrelevant stimuli and tested on relevant stimuli, and are displayed on inflated surface maps from a left lateral view (top left), posterior view (top right) and left medial view (bottom). **f.** Decoding of stimulus orientation (left vs. right vs. front view faces) which was always task irrelevant, in single trial iEEG data, within posterior ROIs (top) and prefrontal ROIs (bottom), collapsed across the three stimulus durations. Lines under the decoding functions indicate time-points showing above chance (33%) decoding from a cluster-permutation test. Decoding using pseudotrial aggregation is shown in Extended Data Figure 5a. 95% CI estimate across cross-validation folds. **g.** Decoding of orientation (left vs. right vs. front view faces) in fMRI using the searchlight approach. Regions with significantly above-chance (33%) decoding accuracies are indicated in outlined blue on the inflated cortical surface maps (top: left/right lateral views; bottom: right/left medial views). **h.** Decoding of orientation (left vs. right vs. front view faces) in iEEG within the ROIs. Regions with electrodes showing above-chance (33%) accuracies are indicated in outlined blue on the inflated surfaces (top left: left lateral view; top right: posterior view; bottom: left medial view). **i.** Decoding of orientation (left vs. right vs. front view faces) in MEG cortical time series within the ROIs (top: posterior; bottom: prefrontal). Time-points showing significantly above-chance (33%) decoding are indicated by lines below the decoding functions. 95% CI estimate across cross-validation folds.

While electrode coverage across our sample of iEEG patients (N=29 for the decoding analyses) was exceptional in the relevant brain regions (Figure 2a, PFC ROIs N_electrodes_=576, Posterior ROIs N_electrodes_=583), we also evaluated these predictions in a larger population of healthy subjects (N=65) in MEG. Results from the cross-task decoding of stimulus categories using the MEG cortical time series (see methods) combining all parcels within the theory-relevant ROIs were consistent with the iEEG observations. Cross-task generalization of face/object decoding was significant in both posterior and prefrontal cortex (Figure 2c) within the theory-predicted time-windows. The extent of cross-temporal generalization of decoding in MEG was sustained in posterior cortex. In PFC, decoding was brief for all three stimulus durations (see supplementary).

A limitation of MEG is its spatial imprecision, which can impact source localization. We thus also tested the theories’ predictions in a large sample of healthy subjects (N=73) exploiting the high spatial resolution of fMRI. Using a searchlight approach (see methods), we found distributed and robust cross-task generalization (∼75%) in striate and extrastriate, ventral temporal, and intraparietal cortex (Figure 2d; see Extended Data Table 4 for anatomical details). Generalization in prefrontal cortex had lower accuracy (∼60%), and was spatially restricted to middle and inferior frontal cortex regions (Figure 2d). We obtained similar results with a decoding approach using theory-relevant ROIs defined in the Destrieux atlas (see supplementary). These results also closely matched a theory-relevant ROIs analysis in the iEEG data restricted to the time windows specified by the theories (Figure 2e). Hence, across recording modalities, we observed that face/object decoding was present both in the posterior and the prefrontal ROI, in line with IIT and GNWT predictions.

As the representation of conscious content is rich and multidimensional including features beyond category, we turned to decoding of stimulus orientation (which was always task irrelevant). Probing decoding of category *and* orientation places a higher bar for theory testing as it requires the satisfaction of two constrains, making it less probable to pass the test ^30^. Here, we found divergent results for the predictions of IIT and GNWT: decoding of face orientation (left/right/front views) was found in posterior cortex but not in prefrontal cortex, both in the iEEG theory-relevant ROIs decoding approach (Figure 2f, h; accuracy improved to ∼95% with pseudotrial aggregation as shown in Extended Data Figure 5a) and in the fMRI searchlight approach (Figure 2g, ∼45%). From the MEG cortical time series, decoding of face orientation was robust in posterior cortex (∼75% with pseudotrial aggregation), and reached above chance levels, albeit weakly (35%) in prefrontal ROIs (Figure 2i). Notably though, control analyses could not conclusively rule out that MEG decoding in the PFC stemmed from signal leakage from posterior regions (Extended Data Figure 5b). Decoding of orientation for the other stimulus categories (letters and false fonts but not for objects) was observed in posterior cortex but not in the prefrontal ROI across the three data modalities (see supplementary).

Finally, we tested IIT’s prediction that prefrontal regions do not contribute further information beyond that specified by posterior areas (or may even degrade performance as it could introduce noise into the classifiers). The results of this test would challenge IIT if the inclusion of PFC was found to increase decoding accuracy, while a lack of an increase would be consistent with both theories as GNWT holds that workspace neurons in PFC broadcast information from posterior processors rather than adding information. We compared decoding performance from classifiers exclusively trained on posterior regions with classifiers trained on posterior and prefrontal regions together (see methods). The results across all three data modalities (iEEG, MEG and fMRI) indicate that neither category nor orientation decoding improves, and in some cases even decreases, when adding prefrontal regions to posterior regions (Extended Data Figure 5c).

Considering the primary preregistered tests of both theories, for **prediction #1**, we found support for IIT: decoding of conscious content (both category and orientation) in posterior cortex was robust, independent of the task manipulation, and consistent across data modalities (iEEG, MEG and fMRI). Also, decoding of category and orientation was found to be the same, or to decrease, when including PFC to posterior regions. Supporting GNWT, we found decoding of category in PFC across all three imaging modalities. For decoding of orientation, results differed across modalities: only for MEG did cortical activity show decoding of orientation for faces but not for any other stimulus category in PFC. Yet, possible signal leakage from posterior sources could not be conclusively ruled-out. Considering the negative decoding results for orientation from fMRI and iEEG, which provide higher spatial resolution than MEG, this overall pattern of results challenges one of GNWT’s predictions.

### Prediction #2: Maintenance of conscious content over time

According to IIT, the state of the network that specifies the content of consciousness in posterior cortex is actively maintained for the duration of the conscious experience (manipulated here via different stimulus durations). In contrast, GNWT predicts brief content-specific ignition in PFC ∼0.3-0.5s after stimulus onset (when the workspace is updated). Then, activity decays to baseline, with information being maintained in an activity-silent state, until another ignition marks the offset of the current percept and the onset of a new percept (in our paradigm, the fixation screen following stimulus offset).

Based on our preregistered predictions, the theories pass if we observe the temporal dynamics for maintenance of conscious content that was predicted, i.e., sustained vs. phasic for IIT and GNWT (Figure 1a), respectively; for a minimum of one conscious feature (category, identity or orientation), in the relevant brain regions and time windows. Specifically, IIT would pass if sustained content-specific information and activation tracking of stimulus duration was found in posterior cortex for at least one of the above-mentioned features. GNWT would pass if prefrontal phasic activation (at onset and offset) associated with the maintenance of conscious content over time is found for at least one of those features. We evaluated those predictions studying both the strength of activation as a function of stimulus duration, and the informational content of that activation in each of the theory-relevant ROIs. We focused on the task irrelevant condition as it is most diagnostic for neural activity related to consciousness, minimizing the contribution of other, potentially confounding, cognitive processes (see supplementary for results on the task relevant condition). Due to the temporal nature of the predictions, they were tested on the two data modalities with millisecond temporal resolution, iEEG and MEG.

First, we tested the theories’ predictions investigating neural activation as a function of stimulus duration. In the iEEG data, we used linear mixed models (LMMs, see methods) to model the time course of neural activity in the high gamma (HG) frequency band (70-150 Hz), which correlates with spiking activity^31,32^, per electrode and theory-relevant ROI as a function of the theories’ predicted temporal models (Figure 1a. middle panel) and stimulus duration (LMMs, see methods). To increase sensitivity and to accommodate the (category) selective responses expected in higher-order sensory areas, we included an interaction term with category.

Electrode sampling in the posterior cortex and PFC was dense and comparable across the theory-relevant ROIs despite the serendipitous nature of the electrode implantation, enabling us to fairly and exhaustively test theories predictions directly in the human brain. Across the 31 epilepsy patients in this analysis, 194 of 657 (29.5%) posterior cortex electrodes and 123 of 655 (18.7%) PFC electrodes exhibited HG activity in response to the stimuli (see supplementary).

In posterior cortex, the results of the LMMs revealed 25 electrodes that exhibited sustained activity that tracked stimulus duration (Extended Data Table 6 for electrode localization and supplementary for results of the full model), in line with IIT’s prediction (Figure 3a). A subset of 12 electrodes showed sustained duration tracking irrespective of stimulus category predominantly in early visual areas (Figure 3b for an example electrode in occipital pole). The remaining 13 electrodes showed category-selective tracking (mostly to face stimuli) localized to the ventral temporal cortex (Figure 3b for an example electrode in lateral fusiform gyrus). Overall, the proportion of electrodes showing category-specificity and duration tracking was small, e.g., only 15% (8/53) of face selective electrodes showed sustained duration tracking as predicted by IIT, pointing to a sparse underlying neural substrate. These responses mostly localized to the lateral fusiform gyrus. The remaining face selective electrodes exhibited transient activations at stimulus onset, localized across striate, extrastriate and ventral areas (see supplementary).

**Figure 3:**
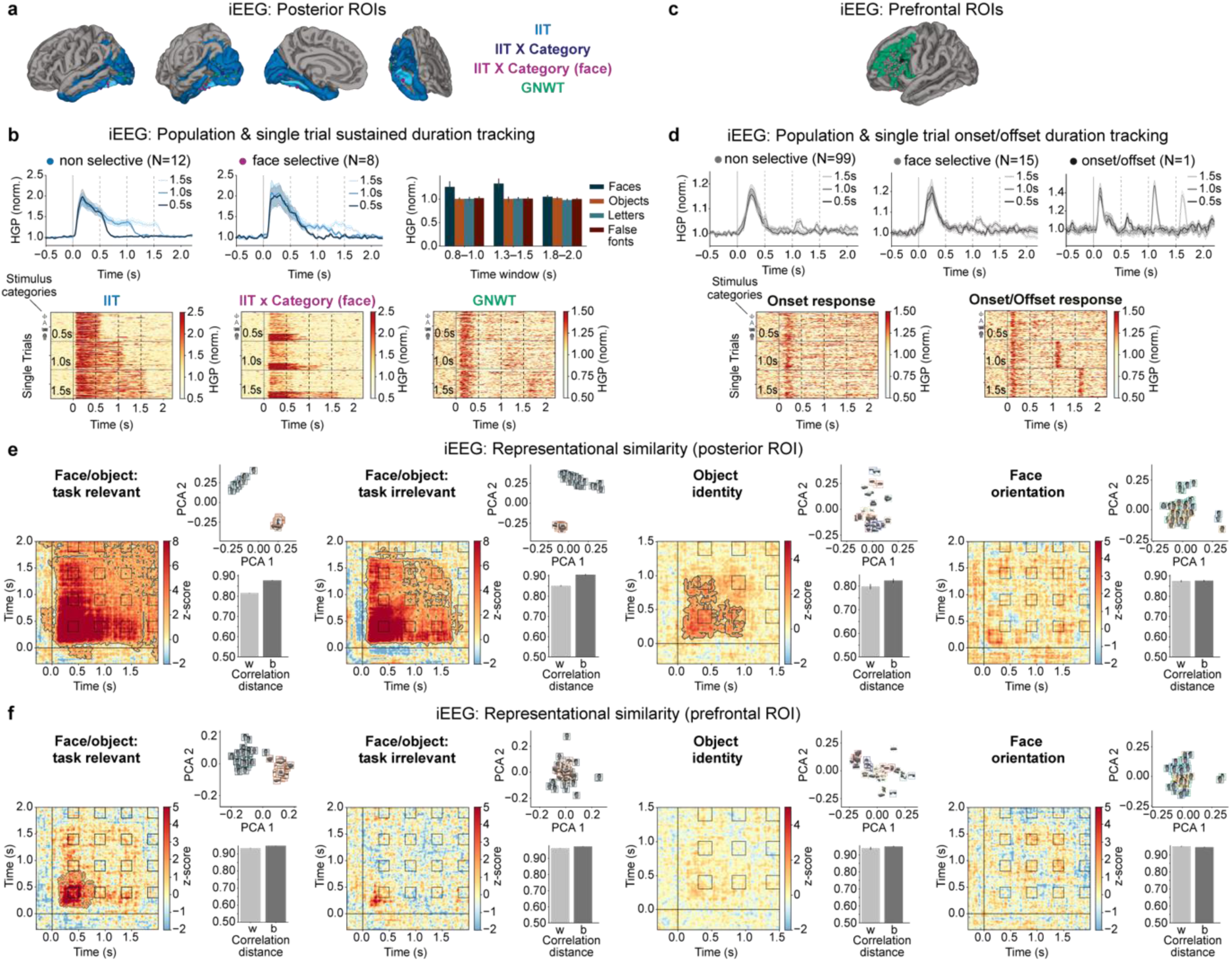
Prediction #2: Maintenance of conscious content over time. **a.** Electrodes in posterior cortex, delineated in blue, (N_subjects_=31, N_electrodes_=657) exhibiting sustained duration tracking compatible with IIT’s predictions, broken down by category-selective electrodes (N=13, dark blue), specifically for faces (N=8, purple), and non-category selective electrodes (N=12, light blue). Electrodes exhibiting the phasic duration tracking predicted by GNWT for PFC are depicted in green (N=11). **b.** Top panels. Averaged waveforms in posterior cortex for non-category selective (left) and face-selective (middle) sustained duration tracking electrodes, separately per stimulus duration, marked in shades of blue. (Right) Bar plot depicting mean high-gamma power averaged across all face-selective electrodes for each stimulus category separate per stimulus duration (faces: dark blue, objects: orange, letters: turquoise, false fonts: dark red). Bottom panels. Raster plots of example electrodes depicting non-category selective sustained duration tracking (left), face-selective sustained duration tracking (middle), and phasic onset and offset duration tracking responses predicted by GNWT for PFC (right). Rows depict single trials, sorted per stimulus duration (from top: 0.5, 1.0, 1.5 s), and then category (from top: false fonts, letters, objects, faces). **c.** Electrodes in PFC (N_subjects_=31, N_electrodes_=655) exhibiting phasic onset responses only (gray, N=114), 1 electrode (black) exhibiting a phasic onset and offset response but significantly earlier (0.15) than the time window predicted by GNWT (>0.3s). None of the 655 electrodes showed phasic onset and offset response (with activity silence in between) at the time windows predicted by GNWT. **d.** Top panels. Averaged waveforms in PFC for non-category selective (left) and face-selective (middle) onset only responsive electrodes, separately per stimulus duration, marked in shades of gray (as their pattern does not comply with any of the theory predictions). (Right) averaged waveforms for the electrode showing an onset & offset response that occur earlier than the predicted time-window. Bottom panels: Raster plots for one example electrode exhibiting an onset response only (left), and the early onset and offset response (right). Y Axis labels as in b. **e.** Cross-temporal representational dissimilarity matrices across all electrodes in posterior cortex (N_subjects_=28, N_electrodes_=583) for category (left), identity (middle) and orientation (right). Sustained representation of category was found irrespective of task (compare task relevant and task irrelevant RSA matrices). Principal component analysis revealed the stable separability across faces and objects, again irrespective of task. Bar plots show the within class dissimilarly (distances within the face and object category) and between class dissimilarity (faces vs. object distances). Larger between than within class separation was observed, consistent with the presence of category information. Sustained information about object identity was observed in posterior cortex, with larger between identity distances and within identity distances. Information about face orientation was weak and not sustained across the stimulus duration in posterior cortex. **f.** Cross-temporal representational dissimilarity matrices across all electrodes in PFC, as in Figure 3a. Transient representation of category was found irrespective of task (compare task relevant and task irrelevant RSA matrices). Principal component analysis revealed the stable separability across faces and objects, again irrespective of task. Bar plots as in Figure 3e. Larger between than within class separation was observed, consistent with the presence of category information. There was no identity nor orientation information in PFC in the relevant time windows predicted by GNWT, or at any other time point.

In PFC, 99 and 24 electrodes showed non-selective or category-selective onset responses, respectively (Figure 3d). Yet, none of the 655 electrodes tested matched the temporal profile predicted by GNWT (i.e., onset and offset). This null result was not due to the analysis approach, as the LMM was indeed sensitive to picking up the pattern predicted by GNWT in 10 electrodes outside the predicted ROI, i.e., in striate/extrastriate cortex (Figure 3b). An exploratory analysis to decode stimulus duration with unrestricted temporal profiles and time windows revealed a single electrode in the inferior frontal sulcus showing the GNWT-predicted pattern, yet earlier than expected (0.15 s) (Figure 3d). The very same electrode exhibited a biphasic event-related potential with a positive deflection early on (0.15 s) and a negative deflection at a later latency (see supplementary). Additional control analyses, including time-locking the analyses to stimulus offset, corroborated the temporal profile predicted by IIT in posterior areas, and the absence of the temporal profile predicted by GNWT in PFC (see supplementary).

For MEG, we used LMMs to investigate the temporal patterns of gamma frequency band power within posterior cortex (15 parcels) and PFC (11 parcels). Even though, gamma frequency band activity was strong in posterior areas none of the theory-based models provided a good fit to the data (see supplementary). Results on alpha frequency in iEEG and MEG were inconclusive and did not provide strong support for either of the theories. In iEEG, none of the prefrontal electrodes showed onset and offset response but instead this pattern was found in posterior sites. In MEG, temporal profiles consistent with GNWT were found in most areas in posterior cortex and in the anterior cingulate cortex but those results were highly dependent on parameter choices and contamination from posterior sites could not be ruled-out (see supplementary).

Together, the results from the temporal activation analysis support IIT’s predictions of sustained activation within posterior cortex. In contrast, we found no evidence in iEEG for the GNWT prediction concerning late phasic ignition of PFC at stimulus onset and offset, despite the presence of robust ignition at the onset of the stimuli. Evidence in the alpha band from MEG was inconclusive but not supported by iEEG despite the ample coverage of PFC. This pattern of results challenges GNWT’s predictions.

After analyzing the temporal profile of brain activity, we used cross-temporal Representational Similarity Analysis (RSA) both in the iEEG and MEG source data to test in which time windows the content of consciousness was represented (Figure 1a. middle panel). For IIT, conscious content should be maintained as long as the conscious experience lasts. GNWT instead predicts a phasic ignition of the workspace at stimulus onset with no active representation of the conscious content until another ignition marks the offset of the percept. Within each of the theory-relevant ROIs, we performed cross-temporal RSA for each stimulus dimension (category, identity, orientation) and correlated them with the temporal models predicted by the theories (Figure 1a, right panel). Here, we report the results for face and object stimuli. Qualitatively similar results were observed for letters/false fonts (Extended Data Figure 7).

In iEEG, we calculated the correlation distance between the patterns of HG activity across 583 electrodes in posterior (N_subjects_=28) and 576 electrodes in PFC cortex (N_subjects_=28), separately. Then, we applied principal component analysis (PCA) to visualize the similarity structure (see methods). We investigated the 1.5 s duration trials only as they enable a better contrast between the temporal profiles predicted by the theories.

In posterior cortex, the cross-temporal RSA revealed sustained face/object categorical representation, with larger correlation distances between categories (face/objects) than within category (face, object) (Figure 3e). The RSA matrix significantly correlated with the temporal model predicted by IIT, and outperformed the GNWT model (see supplementary for results of all contrast).

In PFC, the cross-temporal RSA revealed transient face/object categorical representation at stimulus onset, but not at stimulus offset. In line with this observation, we did not find any significant correlation with the GNWT onset & offset model (Figure 3f). This was the case also in the task relevant condition, where face/object information was stronger, more stable and longer lasting. We further confirmed the absence of GNWT predicted patterns in PFC through three control analyses using (a) feature selection, which improved RSA in PFC; (b) modified time-windows to investigate the possibility of an earlier ignition at stimulus offset; and (c) a decoding analysis time-locking trials to stimulus offset to maximize sensitivity (see supplementary). None of these control analyses changed the overall results.

It has been argued that because conscious experiences are specific, the representation of identity and orientation are more stringent tests of the neural substrate of conscious experience^35^. We thus also evaluated whether information about stimulus identity matched the theories predictions.

In posterior cortex, object identity information was sustained throughout the stimulus duration, with objects of the same identity showing smaller distances than different object identities (Figure 3e). The IIT model significantly correlated with the observed RSA matrix, and also better explained the data compared to the GNWT model. Comparable results were found for letter and false-font identity but not for face identity (Extended Data Figure 7). A different picture emerged in the PFC, where object identity information was absent, both at stimulus onset, offset, and generally throughout the time windows (Figure 3f). This pattern also held true for face, letter and false font identity. Furthermore, these results nicely align with two independent studies using comparable methods^33,34^ attesting to the robustness of the effects. Finally, we tested for the presence of orientation information. In posterior cortex, information about face orientation was weakly present at stimulus onset, yet was not sustained, decaying after 0.5 s (Figure 3e), contrary to IIT’s predictions. In PFC, no information about face orientation was found (Figure 3f). MEG time series were inconclusive, as none of the theories predictions were borne out when testing information about category, identity, or orientation (see supplementary).

Considering the primary preregistered tests of both theories, for **prediction #2**, we found support for IIT as activation and representation of conscious content was sustained in posterior cortex, including representation of category and identity across multiple stimuli. GNWT was however challenged as we found no convincing evidence in iEEG or MEG for a late phasic ignition of PFC at stimulus onset and offset, despite the presence of robust ignition at the onset of the stimuli. RSA analysis demonstrated category information in PFC, exclusively at stimulus onset and earlier than predicted; while information about stimulus identity and orientation was completely absent.

### Prediction #3: Interareal communication

IIT predicts neural connectivity within the posterior cortex, i.e., between high-level and low-level sensory areas (V1/V2), throughout any conscious visual experience. In contrast, GNWT postulates a brief and late metastable state (>0.25 s) with information sharing between PFC and category-specific areas manifested in long-range (gamma/beta) synchronization^36^.

Based on our preregistered predictions, the theories would pass this test if we observe interareal connectivity between the cortical nodes specified by the theories in the relevant time windows. For IIT, this implies *sustained* content-specific synchronization between face/object selective areas and V1/V2; while for GNWT connectivity should be *phasic* (0.3-0.5 s) between the category selective areas and PFC. Due to the temporal nature of the predictions, iEEG and MEG provide the most informative test. We computed pairwise phase consistency^37^ between each category-selective time series (face- and object-selective nodes) and either the V1/V2 or the PFC time series in the intermediate (1.0 s) and long-stimulus-duration (1.5 s), task irrelevant trials (see supplementary for task relevant trials).

For iEEG, we restricted analyses to electrodes showing face and object selectivity, using a different subset of electrodes to test connectivity with V1/V2 and PFC (see methods, Figure 4a for examples of face and object selective electrodes). Due to the sparse coverage, the requirement to focus on ‘activated’ electrodes (see methods) was relaxed. We found increased category selective, e.g., faces>objects synchrony between category-selective and V1/V2 electrodes (Figure 4b, top row). However, these effects were early and short-lived (e.g., <0.75 s), observed only at low frequencies, i.e., 2-25Hz, and mostly explained by the synchronous activity elicited by the stimulus evoked response (Extended Data Figure 8). Thus, the findings did not match IIT predictions, as the activity was not found in the gamma frequency predicted by IIT, and was not sustained. No content-selective PPC was found between face- and object-selective electrodes and PFC in the relevant time window, in contrast to GNWT’s prediction (Figure 4b, bottom row).

**Figure 4:**
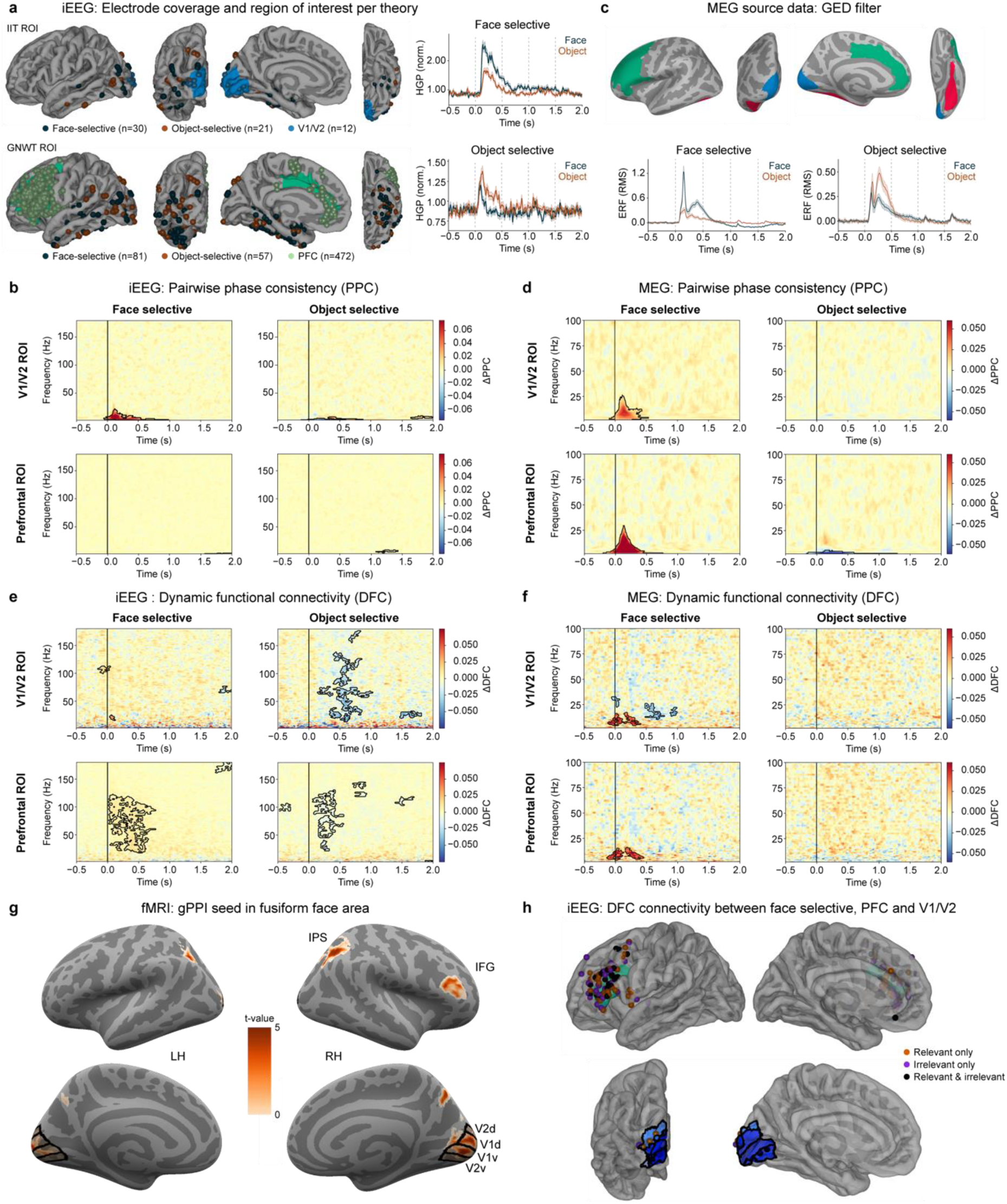
Prediction #3: Interareal communication. **a.** iEEG electrode coverage used to assess content-selective synchrony for IIT ROIs (top, N_subjects_=4) & GNWT ROIs (bottom, N_subjects_=21). Electrode coverage varied between ROIs as interareal connectivity was assessed between electrodes on a per-subject basis. In addition, two example category-selective electrodes are shown: one face-selective, and one object-selective. **b.** iEEG Pairwise phase consistency (PPC) analysis of task irrelevant trials reveals significant content-selective synchrony (e.g. faces > objects for face-selective electrodes; objects > faces for object-selective electrodes) in V1/V2 ROIs (top row), but not in PFC ROIs (bottom row). **c.** MEG cortical time series were extracted per participant from cortical parcels in V1/V2 (blue), PFC (green) and in a fusiform (red) ROIs. Category-selective signals were obtained by creating a category-selective GED filter (i.e., contrasting face/object trials against any other stimulus category trials) on the activity extracted from the fusiform ROI. Face-(bottom left) and object-selective (bottom right) responses averaged across participants are shown at the bottom. **d.** MEG PPC analysis of task irrelevant trials (N=65) reveals significant category-selective synchrony below 25 Hz for the face-selective GED filter (i.e., faces > objects for face-selective electrodes) in both V1/V2 (top row) and PFC ROIs (bottom row) and for the object-selective synchrony (objects > faces for object-selective electrodes) in PFC only. **e.** iEEG Dynamic functional connectivity (DFC) analysis of task irrelevant trials reveals significant content-selective synchrony only for object-selective electrodes in V1/V2 (e.g., top-right), but reveals significant content-selective synchrony for both categories in PFC (bottom row). **f.** MEG DFC analysis of task irrelevant trials (N=65) reveals significant content-selective synchrony below 25 Hz for the face-selective GED filter in both V1/V2 (top left) and PFC (bottom left), but not for the object-selective GED filter. **g.** fMRI gPPI (N=70) on task relevant and task irrelevant trials combined reveals significant content-selective connectivity when FFA is used as the analysis seed. A cluster-based permutation test was used to evaluate the statistical significance of the face > object contrast parameter estimates (p < 0.05). Various significant regions showing task related connectivity with the FFA seed were observed including V1/V2, right intraparietal sulcus (IPS), and right inferior frontal gyrus (IFG). **h.** Analysis of face-selective DFC synchrony across tasks is shown at the single electrode level in PFC (top) & V1/V2 (bottom) ROIs. Electrodes showing significant synchrony in relevant (orange-red), irrelevant (purple), or both relevant & irrelevant (black) task conditions combined are shown (averaged over 70-120 Hz and 0-0.5 s time window). DFC synchrony was observed in both tasks, but restricted to IFG for the GNWT analysis and V2 regions for IIT analysis, consistent with fMRI gPPI analysis shown in panel g.

For MEG, we used Generalized Eigenvalue Decomposition (GED) ^38^ to extract face- and object-selective components from ventral temporal areas (Figure 4c) and then computed PPC. We found selective synchronization between face-selective areas and both V1/V2 and PFC. However, these effects were early and restricted to low frequencies (2-25 Hz), which was inconsistent with both IIT and GNWT (Figure 4d) and mostly explained by stimulus evoked responses (Extended Data Figure 8).

The results of the preregistered PPC metric thus supported neither of the theories. PPC assesses oscillatory phase, and was chosen based on the theories’ mechanistic considerations. However, phase estimation is challenging in neural signals due to noise. We thus relaxed the constraints and tested the theories exploring a connectivity metric sensitive to co-modulations of signal amplitude - dynamic functional connectivity (DFC; see methods). We also removed the evoked responses given the observed impact in the PPC metric (Extended Data Figure 8 includes the evoked response).

In iEEG, we observed significant connectivity between object selective electrodes and V1/V2 (Figure 4e). Connectivity was evident in several frequency bands, most predominantly the gamma band. Yet, it was again brief, in contrast to IIT’s predictions. Connectivity between face selective electrodes and V1/V2 was scarce. Significant connectivity was observed between PFC and both the face and the object-selective areas, in the frequency (gamma) and time range predicted by GNWT. For MEG, brief DFC in the alpha-beta frequency bands was found only between face-selective nodes and both PFC and V1/V2 (Figure 4f).

Together, the results of the exploratory DFC metric in iEEG support GNWT’s predictions, while challenging IIT’s predictions, as connectivity with V1/V2 was not sustained. V1/V2 were however sparsely sampled with iEEG in our population, with only 12 electrodes localized to V1/V2 in contrast to 472 localized in PFC.

Finally, we then moved to fMRI, to evaluate connectivity across the entire cortex with homogeneous sampling. We computed generalized psychophysiological interaction (gPPI), defining Fusiform Face Area (FFA) and Lateral Occipital Complex (LOC) as seed regions per subject based on an anatomically constrained functional contrast (see methods) and combining task relevant and irrelevant trials. FFA showed content selective (face>object stimuli) connectivity with V1/V2, Inferior Frontal Gyrus (IFG) and Intraparietal Sulcus (IPS), consistent with the predictions of both IIT and GNWT (Figure 4g). No selective increases in interareal connectivity between object selective nodes and PFC or V1/V2 was found in fMRI, also when separating task relevant and irrelevant trials (Extended Data Figure 8). To determine whether connectivity to PFC and V1/V2 might be driven by the task in gPPI, we explored the iEEG data separating trials by the task. We found task independent, selective DFC connectivity (face>objects) for face selective electrodes with both IFG and V1/V2 (Figure 4h).

For **prediction #3**, no evidence for IIT or GNWT was found when considering our preregistered analysis. Neither the frequency band nor the temporal patterns of the PPC results were consistent with either theory. Exploring amplitude-based metrics of connectivity (DFC and gPPI), we found support for GNWT predictions, as both in the iEEG and fMRI we observed connectivity with PFC, further matching the timing (∼0.3 s) and spectral composition (gamma frequency) predicted by GNWT. For IIT, though connectivity with V1/V2 was present both in the iEEG and fMRI data, with the expected spectral signature (gamma frequency), it was not sustained throughout the duration of the stimulus, contrary to IIT’s prediction. Further investigations may be required given the sparse coverage of V1/V2 in iEEG.

### Putative Neural Correlates of Consciousness (pNCC)

Finally, we also aimed at narrowing down the cortical areas that are potentially involved in (visual) consciousness (i.e., ‘putative NCCs’), by detecting areas that consistently respond to visual stimuli while ruling out cortical areas responsive to other, accompanying (but confounding) cognitive processes, e.g., performing a task on the conscious content and motor responses ^39^. This test has implications for both theories, as they differ in their predictions about the NCC. IIT predicts that the cortical substrate of consciousness should include posterior areas while agreeing that certain PFC areas should be excluded due to task confounds. GNWT predicts an involvement of PFC even after ruling out task-based effects.

Based on our preregistered predictions, we used a contrast-conjunction approach (see Methods), both on univariate activation and decoding data. First, we identified voxels sensitive to the task itself, either to its goal, responding to the target, or to task relevance in general. These two contrasts revealed voxels in several prefrontal ROIs, including dorsolateral prefrontal, premotor, and motor cortex (Extended Data Figure 9). These voxels were excluded from further analyzes.

Then, among the remaining areas, we identified brain areas sensitive to changes in the content of consciousness, so that they consistently respond to at least one stimulus category (Stimulus>Baseline) in both the task relevant and task irrelevant conditions (Figure 5, see supplementary for results from all ROIs and contrast-conjunctions at the subject level). In posterior cortex, several regions in ventral occipito-temporal regions showed consistent task-independent activation for three or all four stimulus categories. In PFC, inferior and middle frontal gyrus and orbital cortex were activated for at least one of the stimulus categories. A number of areas showed deactivations both in posterior cortex (e.g., striate and some extrastriate areas) and PFC (e.g., inferior and middle frontal gyrus and orbital cortex). The complementary decoding approach revealed regions in extrastriate and early visual cortex and small, right lateralized clusters in PFC (Extended Data Figure 9 and supplementary for tables of effects within each ROI).

**Figure 5.**
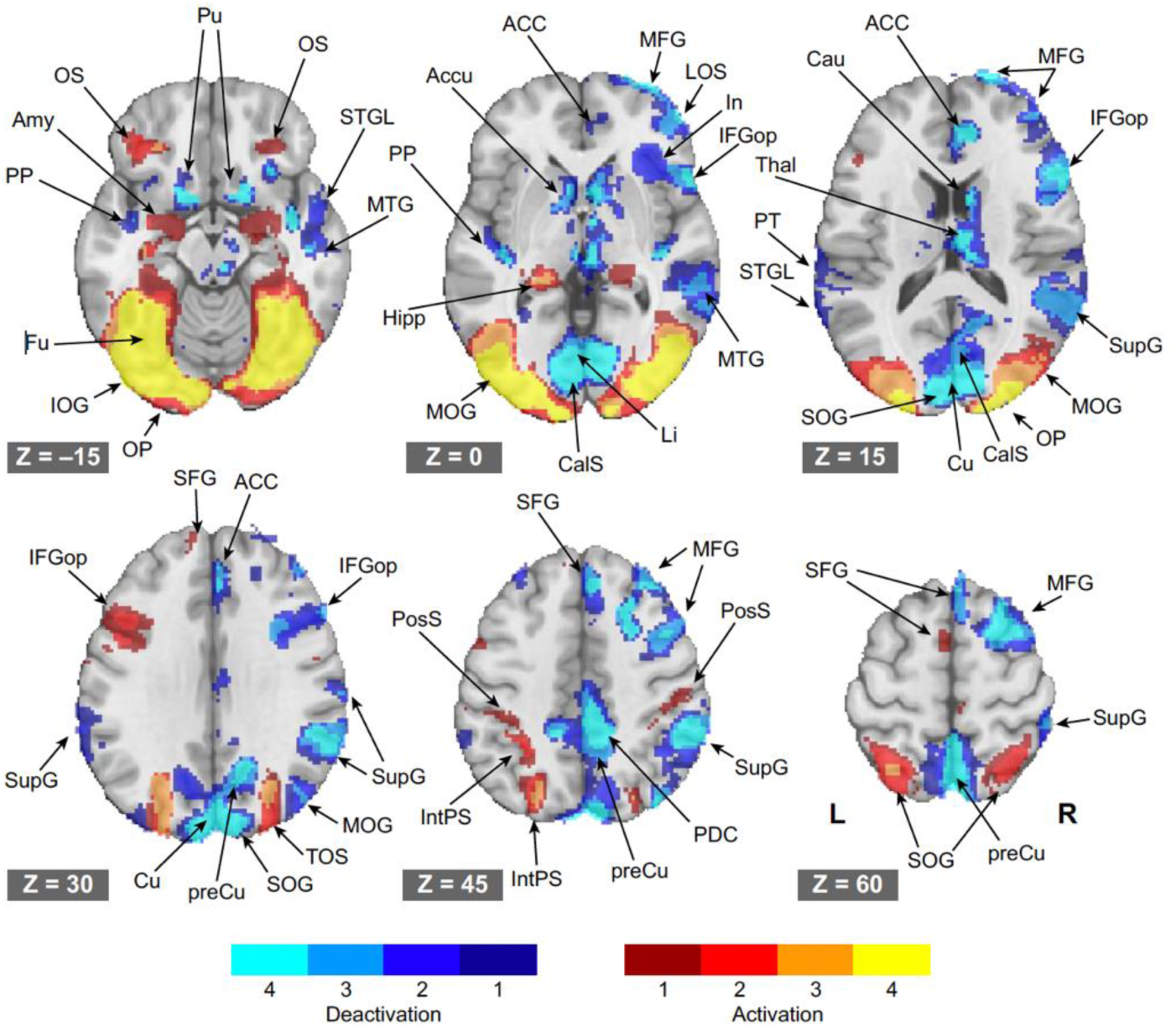
fMRI Univariate contrast conjunction analysis aimed at demarcating putative NCC. This conjunction analysis identifies visually-responsive cortical areas, after removing (confounding) task-responsive cortical regions. Axial brain slices show activations (reds-yellows) and deactivations (blues) (N=73), relative to a blank-screen baseline condition for each of the 4 stimulus categories. Color scales indicate the number of stimulus categories (1-4) passing the contrast-conjunction, as in [(task relevant stimulus > baseline) & (task irrelevant > baseline)] OR [(task relevant stimulus < baseline) & (task irrelevant < baseline)]. Cortical regions associated with task goals and task relevance, identified in two separate contrast-conjunction analyses (see Extended Data Figure 9), were removed from the activity maps shown here. Axial brain slices are displayed from inferior (top left) to superior (bottom right). Left and right hemisphere are displayed to the left and right, respectively. Neuroanatomical labels from the Destrieux atlas and additional subcortical regions : AC: Anterior Cingulate Gyrus; AG: Angular Gyrus; Accu: Nucleus Accumbens; Amy: Amygdala; CalS: Calcarine Sulcus; Cau: Caudate Nucleus; Cu: Cuneus; Fu: Fusiform gyrus; Hipp: Hippocampus; IFGop: Opercular part of the Inferior Frontal Gyrus; IFGtri: Triangular part of the Inferior Frontal Gyrus; In: Insula; IntPS: Intraparietal Sulcus; IOG: Inferior Occipital Gyrus; Li: Lingual Gyrus; LOS: Lateral Orbital Sulcus; MFG: Middle Frontal Gyrus; MOG: Middle Occipital Gyrus; MTG: Middle Temporal Gyrus; OP: Occipital Pole; OS: Orbital Sulci; PDC: Posterior Dorsal Cingulate; PP: Planum Polare of the Superior Temporal Gyrus; preCu: Precuneus; PosS: Postcentral Sulcus; PreSinf: Inferior part of the Precentral Sulcus; PT: Planum Temporale of the Superior Temporal Gyrus; Pu: Putamen; SFG: Superior Frontal Gyrus; SOG: Superior Occipital Gyrus; SPL: Superior Parietal Lobule; STGL: Lateral aspect of the Superior Temporal Gyrus; Sup: Supramarginal gyrus; Thal: Thalamus; TOS: Transverse Occipital Sulcus.

Together, the pNCC analysis revealed a pattern of candidate areas that was more spatially restricted than anticipated by the rather extensive preregistered theory ROIs. Specifically, the MFG, IFG and orbital cortex might participate in consciousness, as predicted by GNWT. Furthermore, the scant activation patterns found in PFC compared to the widespread deactivations was surprising, and suggests a reconsideration of the strong focus on activations (relative to deactivations) when assessing this region’s role in conscious perception. However, we consider this analysis an informative yet liberal test, given its potential to overestimate candidate cortical areas for consciousness by including non-conscious sensory precursors.

### General Discussion

This adversarial collaboration was aimed at overcoming researchers’ confirmation biases, breaking theoretical siloes^3^, identifying strengths and weaknesses of the theories^2,40^, rigorously testing them on common methodological grounds ^13,20^, and providing the means to change one’s mind given contradictory results^13^. In doing so, this approach enables progress in the field by catalyzing our ability to evaluate and arbitrate between theories of consciousness. Embracing this spirit, we opted for a discussion in three voices because even if we provide a stringent test and brought together incompatible theoretical views, different interpretations of the evidence may remain. In what follows, the theory-neutral consortium first presents the main challenges our study poses to the theories and then the adversaries offer their interpretation and future directions.

#### Cogitate consortium

Passed/failed predictions of the theories across data modalities are summarized in Extended Data Figure 10. The table highlights several challenges for both theories.

For IIT, the lack of sustained synchronization within posterior cortex represents the most direct challenge, based on our preregistration. Across several analyses, with various degrees of sensitivity, we only observed transient synchronization between category selective and early visual areas. This is incompatible with IIT’s claim that the state of the neural network, including its activity and connectivity, specifies the degree and content of consciousness ^5^. Although this null result could stem from methodological limitations (e.g., limited iEEG sampling of V1/V2 areas), our multimodal and highly powered study provided the best conditions so far for the predicted patterns to be found. We urge IIT proponents to direct future efforts to evaluate this prediction and to determine its significance and the extent of this failure.

More broadly, although IIT passed the preregistered duration prediction (#2), there was no evidence for a sustained representation of orientation, though orientation is usually a fundamental property of our conscious experience, and should have accordingly showed sustained representation^23^. This is an informative challenge for IIT, as orientation decoding was robust across all three data modalities, leaving open the question of how information about orientation is maintained over time.

Finally, our pNCC analysis suggested that portions of PFC might be important for consciousness. While IIT correctly predicted that the most consistent activation and decodability of content would be found in posterior cortex, it must explain the finding that the MFG and the IFG (for which we also found results in the decoding and synchrony analysis), were visually responsive and not ruled out as being task-related. This finding is particularly important to explain in the context of the current experiment where additional cognitive processing of the task irrelevant stimuli was minimized. ^41^

For GNWT, the most significant challenge based on our preregistered criteria pertains to its account for the maintenance of a conscious percept over time; and in particular, the lack of ignition at stimulus offset. In most of our main tests and control analyses across data modalities (for details, see supplementary), we failed to reveal an offset response in PFC (both in activation and in reinstatement of decoded content of any type). This result is less likely to stem from sensitivity limitations, since offset responses were robustly found elsewhere (e.g., visual areas); and in PFC, strong onset responses were found to the very same stimuli. The lack of ignition at stimulus offset is especially surprising given the change of conscious experience at the onset of the blank fixation screen. This clear update to the content of consciousness should have been represented somehow by the global workspace^12^. Thus, as our results do not support GNWT’s predictions regarding the maintenance of conscious experience, that aspect of consciousness remains unexplained within the GNWT framework.

Another key challenge for GNWT pertains to representing the contents of experience: though we found representation of category in PFC irrespective of the task, hereby demonstrating the sensitivity of our methods, no representation of identity was found, and representation of orientation was only evident in MEG (without being able to exclude source leakage effects), although these dimensions are a primal aspect of our conscious experience. This raises the question of whether PFC is involved in broadcasting *all* conscious content as predicted by GNWT ^21^ or only a subset (e.g., abstract concepts and categories, rather than low-level details), in which case the role of PFC in consciousness might need to be redefined.

Finally, the highly spatially restricted decoding of conscious content in PFC, alongside the restricted activations and deactivations in PFC observed in the pNCC analysis, point to a “localized spark” rather than the “wide-spread ignition” predicted by the theory, further challenging it. ^7^

Prior to the current study, the predictions from IIT and GNWT had mostly been tested with one data modality at a time ^21,22^, leaving interpretational freedom for negative results, which can easily be attributed to the limitations of a given modality ^42^. Here, the combination of techniques allowed us to cross-compensate for their respective limitations to thoroughly and systematically assess the theories’ predictions. This approach was mutually agreed upon by the theory leaders’ ex-ante as the most powerful and conclusive approach, making both positive and negative findings more meaningful.

Conceptually, our study focused on the mechanisms by which the content of the conscious experience of A differs from the experience of B (i.e., category, identity, orientation and duration), which addresses how the link between brain activity and subjective phenomenology changes between distinct conscious experiences. As such, we departed from the mainstream contrastive approach in which the presence of conscious experience is contrasted with its absence to study whether there was a conscious experience or not. Though widely used, the standard contrastive approach suffers from shortcomings which precludes it from directly revealing the processes related to consciousness, as it confounds consciousness with other cognitive processes such as decision-making, reporting, or the formation of episodic memory traces after a conscious experience ^39,43,44^. Studying the content of consciousness more directly links phenomenology to brain activity and overcomes several of the limitations of the contrastive method. Yet, some might argue that in doing so, we are tracking mere stimulus processing rather than consciousness per se. However, in the context of this adversarial collaboration, whose purpose is to falsify ^45^ divergent predictions of IIT and GNWT and not to provide confirmatory evidence, this perceived weakness is actually an asset: if the theories’ main positive predictions fail in the face of fully attended, consciously experienced stimuli, this provides evidence that the proposed neural mechanism is unlikely to be minimally necessary for conscious experience. Hence, our approach poses a principled test to both theories.

Beyond the direct challenges to the theories themselves, our study raises a number of important questions for theory testing and theory building, which apply broadly across most fields, e.g., how to weigh different theory predictions, and how to combine evidence across predictions, analyses and measures (in our case, fMRI, MEG and iEEG data). From the outset, we defined an independent set of predictions, setting criteria for failure to then weigh the results against these predictions. Yet, a formal framework that quantitively integrates evidence by weighing and quantitatively integrating over passes and failures, accounting for the centrality of the predictions for the theory, measurement error, and consistency across samples and measurements is direly needed to enable systematic theory building in the era of accumulation of results.

#### Integrated Information Theory: Melanie Boly, Christof Koch, Giulio Tononi

The results corroborate IIT’s overall claim that posterior cortical areas are sufficient for consciousness, and neither the involvement of PFC nor global broadcasting are necessary. They support preregistered prediction #1, that decoding conscious contents is maximal from posterior regions but often unsuccessful from PFC, and prediction #2, that these regions are sustainedly activated while seeing a stimulus that persists in time. They do not support prediction #3 concerning sustained synchrony, although there are potential explanations (see supplementary). Below we illustrate how these predictions were motivated by IIT.

Posterior regions are often considered mere ‘information processors’; their activation, it is claimed, may be necessary but not sufficient for experiencing specific contents. For example, they may show activations during deep sleep or anesthesia and for unreported stimuli under contrastive, near-threshold paradigms.^8^ This seems to warrant the need for additional ingredients, such as ‘global broadcasting’^8^ or ‘higher-order monitoring’ by PFC.^10^

For IIT, however, posterior regions are sufficient for consciousness as long as they satisfy the requirements for maximal integrated information. Why this prediction? Unlike other approaches, IIT infers the essential, physical requirements for the substrate of consciousness from the essential properties of experience.^4,5^ This leads to the claim that the quality and quantity of an experience are accounted for by the ‘cause–effect structure’ specified by a substrate with maximal integrated information, called the ‘main complex’.^4,5^ We conjectured that posterior cortical regions should provide an excellent substrate for the main complex owing to their dense local connections arranged topographically into a hierarchical, divergent– convergent 3D lattice,^5^ leading to prediction #1. Nevertheless, by IIT, posterior regions can only support consciousness if their physiology ensures high integrated information—which indeed breaks down^46^ due to bistability when consciousness is lost in deep sleep and anesthesia.^47–49^

Much of PFC, in contrast, seems to be organized not as a grid but as a patchwork of segregated columns,^50^ unfavorable for high integrated information. Even so, any PFC region organized in a grid-like way with dense interconnections with posterior regions may well be part of the main complex. As previously emphasized,^51^ *“…we bear no preconceived enmity to the prefrontal cortex. Indeed, searching for the NCC of specific aspects of experience…in certain anterior regions is an important task ahead*.” For example, parts of IFG might contribute to, say, an abstract/evaluative/actionable experiential aspect of faces, which could be consistent with some pNCC analysis results. However, IIT predicts that we would still experience faces (sans aspects contributed by PFC regions) if PFC were selectively inactivated. For IIT, all quality is structure: all properties of an experience are accounted for by properties of the cause–effect structure specified by the main complex. Every conscious content (face, object, letter, blank screen) is thus a (sub)structure of integrated information (irreducible cause-effects and their overlaps^4^); it is neither a message that is encoded and broadcasted globally,^8,52,53^ nor a distributed activity pattern, nor a neural process. Indeed, IIT’s research program aims to account for specific consciousness contents—why space feels extended, time feels flowing, and phenomenal objects feel like binding general concepts (invariants) with particular features—all exclusively in terms of their corresponding cause– effect structures.^4,23^ As highlighted in the Introduction, when we see Mona Lisa, we see that it is a face, with her particular features, at a particular location on the canvas, and we see her for as long as we look at her. This is why we predicted (prediction #2) that the NCC in posterior cortex would last for the duration of the percept, notwithstanding the widespread evidence for neural adaptation and onset/offset neural responses (probably due to transient excitation/inhibition imbalance), and (prediction #3) that synchrony would occur (reflecting causal binding) between units in higher and lower areas, supporting respectively invariant concepts and particular features.

To conclude, moving beyond the contrastive paradigm between seen and unseen stimuli and beginning to account for how experience feels is one key reason why the experiments reported in this adversarial collaboration mark an important development. Another is that they inaugurate a powerful new way of making progress on a problem often considered beyond the reach of science. The group that carried out this endeavor did so in a way that was explicit, open, and truly collaborative—in short, in a way that is paradigmatically scientific.

#### Global Neuronal Workspace Theory: Stanislas Dehaene

This unprecedented data collection effort brings several new insights relevant to our theory. Most importantly, the results confirm that PFC exhibits a metastable bout of activity (“ignition”) for about ∼200 ms, in a content-specific manner, even for task irrelevant stimuli, irrespective of stimulus duration (Figures 2b, 3f, Supplementary Figure 23), and with a concomitant transient increase in long-distance dynamic functional connectivity with face- and object-selective posterior areas (Figure 4e-h). Those findings, unpredicted by IIT but predicted by GNWT, support previous findings that PFC contains a detailed code for conscious visual contents ^28,54–58^. They also counter previous conclusions that were, in our opinion, too hastily drawn on the basis of insufficient evidence ^29^: with suitably sensitive experiments, content-specific PFC regions do show a transient ignition even for irrelevant stimuli. While agreeing with previous results ^58–62^, the convergence of iEEG, MEG and fMRI in the same task alleviates concerns associated with a possible mis-reconstruction of EEG/MEG sources. It also resolves a controversy related to the timing of conscious ignition, which was initially thought to be associated with the P300 ERP waveform ^8^, but can obviously arise earlier (∼200 ms post-onset) ^59,62^. GNWT would further predict that this latency should vary depending on the strength of both bottom-up accumulating evidence (e.g., contrast^63^) and top-down attention/distraction by other tasks^59,61,64^.

While some results do challenge GNWT, they do not seem unsurmountable given experimental limitations. First, note that there is a considerable asymmetry in the specificity of the theories’ predictions. None of the massive mathematical backbone of IIT, such as the φ measure of awareness, was tested in the present experiment. Instead, what are presented as unique predictions of IIT (posterior visual activation throughout stimulus duration) are just what any physiologist familiar with the bottom-up response properties of those regions would predict, since visual neurons still respond selectively during inattention or general anesthesia ^65–67^. Such posterior stimulus-specific, duration-dependent responses are equally predicted by GNWT, but attributed to non-conscious processing.

Unfortunately, here, it is impossible to decide which of the activations reflected conscious versus non-conscious processing, because the experimental design did not contrast conscious versus non-conscious conditions (fortunately, a second experiment by the Cogitate consortium will include such a contrast). The present experiment relied on the seemingly innocuous hypothesis that stimuli were “indubitably consciously experienced” for their entire duration. However, it is well known that perfectly *visible* stimuli, depending on attention orientation, may fail to be *seen* (attentional blink, inattentional blindness) ^68,69^ or may become conscious at a time decoupled from stimulus presentation (psychological refractory period, retro-cueing) ^64,70–72^. Here, it seems likely that subjects briefly gained awareness of all the images (since they remembered them later), but then reoriented their conscious thoughts to other topics, without waiting for image offset – and this interpretation perfectly fits the ignition profile that was found in PFC. It would be surprising if participants’ consciousness remained tied to each image for its full duration on every trial of this long experiment. It is also unclear whether participants were ever aware of stimulus orientation, which was always irrelevant. A new experiment, using quantified introspection ^64^, will be needed to assess for how long participants maintained the visual image in consciousness.

For the same reason, the absence of decodable activation at stimulus offset, while challenging, may simply indicate that participants never consciously attended to that event, which was always uninformative and irrelevant. Making stimulus offset more attractive, for instance by turning it into an occlusion event where an object hides behind a screen, could yield different results.

For GNWT, the prefrontal code for a conscious mental object is thought to involve a vector code distributed over millions of neurons which, unlike in posterior regions, are not clustered but spatially intermingled ^28,73^. Thus, we are not surprised that PFC responses are hard to decode from the macro- or mesoscopic signals measured by fMRI, MEG, or large intracranial electrodes that pool over tens of thousands of neurons. Therefore, the present positive results, indicating transient PFC ignition and decoding of faces and objects, seem to us more important than the null ones, especially as there is already much single-neuron evidence that PFC contains even more precise stimulus-specific neural codes ^28,54–56^.

Finally, while the theories concern the necessary regions for conscious experience, the present methods are purely correlational and do not evaluate causality. This limitation is not unique to the present work, but applies to any brain-imaging experiment. While applauding the present efforts, we therefore eagerly await the results of other adversarial collaborations using causal manipulations in animal models.

### Conclusion (Cogitate consortium)

At this point, the reader might expect the consortium to draw a final conclusion. Instead, we invite the reader to form their own conclusions considering the relative evidence we presented for each of the preregistered predictions, the scope of the evidence with > 250 subjects using the most sophisticated techniques available to human neuroscience, and the challenges in changing people’s minds. Science is a social enterprise, and the reader is as much a part of this enterprise as any of the authors from this consortium.

## Supporting information

Experimental Procedures Video

## Methods

### Preregistration and data availability

The full study protocol is available in <u>the preregistration</u> on the OSF webpage, including: (a) an exhaustive description of the experimental design, (b) the theories’ predictions and agreed upon interpretations of the results, (c) iEEG, MEG, and fMRI data acquisition details; (d) preprocessing pipelines; and (e) data analysis procedures. All data and code will be shared upon publication. Below, the main methods are concisely summarized.

### Ethics Statement

The experiment was approved by the institutional ethics committees of each of the data-collecting labs (see supplementary for details). All volunteers and patients provided oral and written informed consent before participating in the study. All study procedures were carried out in accordance with the Declaration of Helsinki. Epilepsy patients were also informed that clinical care was not affected by participation in the study.

### Participants

Healthy volunteers and patients with pharmaco-resistant focal epilepsy participated in this study. The datasets reported here consist of: (1) Behaviour, eye tracking and invasive electroencephalogram (iEEG) data collected at the Comprehensive Epilepsy Center at New York University (NYU) Langone Health, Brigham and Women’s Hospital, Boston Children’s Hospital (Harvard), and University of Wisconsin School of Medicine and Public Health (WU). (2) Behaviour, eye tracking, magnetoencephalographic (MEG) data collected at the Centre for Human Brain Health (CHBH) of the University of Birmingham (UB), and at the Center for MRI Research of Peking University (PKU). (3) Behaviour, eye tracking and functional magnetic resonance (fMRI) data collected at Yale Magnetic Resonance Research Center (MRRC) and at the Donders Centre for Cognitive Neuroimaging (DCCN), of Radboud University Nijmegen. For both the MEG and fMRI datasets, a 1/3 of the data that passed quality tests (henceforth, *Optimization dataset*; see <u>preregistration</u> for details about quality test criteria) were used to optimize the analysis methods, which were subsequently added to the preregistration as an additional amendment. These preregistered analyses were then run on the remaining 2/3 of the data (henceforth, *Replication dataset*) and constitute the data reported in the main study. For comparison, results from the optimization phase are reported in the supplementary material. This procedure was not used for the iEEG data due to the serendipitous nature of the recording and electrode placement, the rarity of this type of data and the increased difficulty of data collection due to the COVID-19 pandemic.

For the iEEG arm of the project, a total of 34 patients were recruited. Two patients were excluded due to incomplete data. Demographic, medical and neuropsychological scores for each patient, when available, are reported in Supplementary Table 25. Three iEEG patients whose behavior fell slightly short of the predefined behavioral criteria (i.e. hits < 70%, FA > 30%) were nonetheless included given the difficulty to obtain additional iEEG data. A total of 97 healthy subjects were included in the MEG sample (mean age 22.79 ± 3.59 years, 54 females, all right-handed), 32 of those datasets were included in the optimization phase (mean age 22.50 ± 3.43 years, 19 females, all right-handed), and 65 in the replication sample (mean age = 22.93 ± 3.66, 35 females, all right-handed). Five additional subjects were excluded from the MEG dataset: two due to failure to meet predefined behavioral criteria (i.e., hits < 80%, and/or FA > 20%), two due to excessive noise from sensors, and one due to incorrect sensor reconstruction. A total of 108 healthy participants were included in the fMRI sample (mean age 23.28 ± 3.46 years, 70 females, 105 right-handed), 35 of those datasets were included in the optimization sample (mean age 23.26 ± 3.64 years, 21 females, 34 right-handed), and 73 in the replication sample (mean age = 23.29 ± 3.37, 49 females, 71 right-handed). Twelve additional subjects were excluded from the fMRI dataset: eight due to motion artifacts, two due to insufficient coverage, and two due to incomplete data.

### Experimental procedure

#### Experimental design

To test critical predictions of the theories, five experimental manipulations were included in the experimental design: (1) stimulus category (faces, objects, letters and false fonts), (2) stimulus identity (20 different exemplars per stimulus category), (3) stimulus orientation (front, left and right view), (4) stimulus duration (0.5 s, 1.0 s, 1.5 s), and (5) task relevance (relevant targets, relevant non-targets, irrelevant).

Stimulus category, stimulus identity and stimulus orientation served to test predictions about the representation of the content of consciousness in different brain areas by the theories. In addition, stimulus duration served to test predictions about the temporal dynamics of sustained conscious percepts and interareal synchronization between areas. Task relevance served to rule out the effect of task demands, as opposed to conscious perception per se, on the observed effects ^1^.

#### Stimuli

Four stimulus categories were used: faces, objects, letters and false fonts. These stimuli naturally fell into two clearly distinct groups: pictures (faces and objects) and symbols (letters and false fonts). These natural couplings were aimed at creating a clear difference between task relevant and task irrelevant stimuli in each trial block (see Procedure). All stimuli covered a squared aperture at an average visual angle of 6° by 6°. Face stimuli were created with FaceGen Modeler 3.1; letter and false fonts stimuli were generated with MAXON CINEMA 4D Studio (RC - R20) 20.059; object stimuli were taken from the Object Databank^2^. Stimuli were gray-scaled and equated for luminance and size. To facilitate face individuation, faces had different hairstyles and belonged to different ethnicities and genders. The orientation of the stimuli was manipulated, such that half of the stimuli from each category had a side view (30° and −30° horizontal viewing angle, left and right orientation) and the other half had a front view (0°).

#### Procedure

Subjects performed a non-speeded target detection task (see supplementary video). The experiment was divided into runs, with four blocks in each run (see Trial counts below). On a given block, subjects viewed a sequence of single, supra-threshold, foveally presented stimuli belonging to four stimulus categories and presented for three stimulus durations. Within each block, half of the stimuli were task relevant and half task irrelevant. To manipulate task relevance, at the beginning of each block subjects were instructed to detect the rare occurrences of two target stimulus identities, one from each relevant category (pictures: face/object or symbols: letter/false-font), irrespective of their orientation. This was specified by presenting the instruction “detect face A and object B” or “detect letter C and false-font D”, accompanied by images for each target (See Figure 1e). Targets did not repeat across blocks. Each run contained two blocks of the Face/Object task and two blocks of the Letter/False-font task, with block order counterbalanced across runs.

Accordingly, each block contained three different trial types: i) *Targets*: the two stimuli being detected (e.g., the specific face and object identities); ii) *Task Relevant Stimuli*: all other stimuli from the task relevant categories (e.g., the non-target faces/objects); and iii) *Task Irrelevant Stimuli*: all stimuli from the two other categories (e.g., letters/false fonts). An advantage of this design is that the three trial types enabled a differentiation of neural responses related to task goal, task relevance, and simply consciously seeing a stimulus.

Stimuli were presented for one of three durations (0.5 s, 1.0 s or 1.5 s), followed by a blank period of a variable duration to complete an overall trial length fixed at 2.0 s. For the MEG and iEEG version, random jitter was added at the end of each trial (mean inter-trial interval of 0.4 s jittered 0.2-2.0 s, truncated exponential distribution) to avoid periodic presentation of the stimuli. The mean trial length was 2.4 s. For the fMRI protocol, timing was adjusted as follows: the random jitter between trials was increased (mean inter-trial interval of 3 s, jittered 2.5-10 s, with truncated exponential distribution), with each trial lasting approximately 5.5 s. This modification helped avoid non-linearities in BOLD signal which may impact fMRI decoding^3^. Second, to increase detection efficacy for amplitude-based analyses, three additional baseline periods (blank screen) of 12 s each were included per run (total = 24). The identity of the stimuli was randomized with the constraint that they appeared equally across durations and tasks conditions.

Subjects were further instructed to maintain central fixation on a black circle with a white cross and another black circle in the middle throughout each trial (see Figure 1e).

#### Trial counts

The MEG study consisted of 10 runs containing 4 blocks each with 34-38 trials per block, 32 non-targets (8 per category) and 2-6 targets, for a total of 1,440 trials. The same design was used for iEEG, but with half the runs (5 runs total), resulting in a total of 720 trials. For fMRI, there were 8 runs containing 4 blocks each with 17-19 trials per block, 16 non-targets (4 per category) and 1-3 targets, for a total of 576 trials. Rest breaks between runs and blocks were included.

### Data Acquisition

#### Behavioral data acquisition

The task was run on Matlab (PKU: R2018b; DCCN, UB and Yale: R2019b; Harvard: R2020b; NYU: R2020a, WU: 2021a) using Psychtoolbox v.3 ^4^. The iEEG version of the task was run on a Dell Precision 5540 laptop, with a 15.6“ Ultrasharp screen at NYU and Harvard and on a Dell D29M PC with an Acer 19.1” screen in WU. Participants responded using an 8-button response box (Millikey LH-8; response hand(s) varied based on the setting in the patient’s room). The MEG version was run on a custom PC at UB and a Dell XPS desktop PC on PKU. Stimuli were displayed on a screen placed in front of the subjects with a PROPixx DLP LED projector (VPixx Technologies Inc.). Subjects responded with both hands using two 5-button response boxes (NAtA or SINORAD). The fMRI version was run on an MSI laptop at Yale and a Dell Desktop PC at DCCN. In DCCN, stimuli were presented on an MRI compatible Cambridge Research Systems BOLD screen 32” IPS LCD monitor, and in Yale they were presented on a Psychology Software Tools Hyperion projection system to project stimuli on the mirror fixed to the head coil. Subjects responded with their right hand using a 2×2 Current Designs response box at Yale and a 1×4 Current Designs response box at DCCN.

#### Eye tracking data acquisition

For the iEEG setup, eye tracking and pupillometry data were collected using a EyeLink 1000 Plus on a remote mode, sampled monocularly at 500 Hz (from the left eye at WU, and depending on the setup at Harvard), or on a Tobii-4C eye-tracker, sampled binocularly at 90 Hz (NYU). The MEG and fMRI labs used the MEG and fMRI compatible EyeLink 1000 Plus Eye-tracker system (SR Research Ltd., Ottawa, Canada) to collect data at 1000 Hz. For MEG, eye tracking data were acquired binocularly. For fMRI, data were acquired monocularly from either the left or the right eye, in DCCN and Yale, respectively. For all recordings, a nine-point calibration was performed (besides Harvard, where thirteen-point calibration was used) at the beginning of the experiment, and recalibrated as needed at the beginning of each block/run.

#### iEEG data acquisition

Brain activity was recorded with a combination of intracranially subdural platinum-iridium electrodes embedded in SILASTIC sheets (2.3 mm diameter contacts, Ad-Tech Medical Instrument and PMT Corporation) and/or depth stereo-electroencephalographic platinum-iridium electrodes (PMT Corporation; 0.8-mm diameter, 2.0-mm length cylinders; separated from adjacent contacts by 1.5 to 2.43 mm), or Behnke-Fried depth stereo-electroencephalographic platinum-iridium electrodes (Ad-Tech Medical, BF08R-SP21X-0C2, 1.28 mm in diameter, 1.57 mm in length, 3 to 5.5 mm spacing). Electrodes were arranged as grid arrays (either 8 × 8 with 10 mm center-to-center spacing, 8 x 16 contacts with 3 mm spacing, or hybrid macro/micro 8 x 8 contacts with 10 mm spacing and 64 integrated microcontacts with 5 mm spacing), linear strips (1 × 8/12 contacts), depth electrodes (1 × 8/12 contacts), or a combination thereof. Recordings from grid, strip and depth electrode arrays were done using a Natus Quantum amplifier (Pleasonton, CA) or a Neuralynx Atlas amplifier (Bozeman, MT). A total of 4057 electrodes (892 grids, 346 strips, 2819 depths) were implanted across 32 patients with drug-resistant focal epilepsy undergoing clinically motivated invasive monitoring. 3512 electrodes (780 grids, 307 strips, 2425 depths) that were unaffected by epileptic activity, artifacts, or electrical noise were used in subsequent analyses. To determine the electrode localization for each patient, a post-operative computed tomography scan and a pre-operative T1 MRI were acquired and co-registered.

#### MEG data acquisition

MEG was acquired using a 306-sensor TRIUX MEGIN system, comprising 204 planar gradiometers and 102 magnetometers in a helmet-shaped array. The MEG gantry was positioned at 68 degrees for optimal coverage of frontal and posterior brain areas. Simultaneous EEG was recorded using an integrated EEG system and a 64-channel electrode cap (EEG data is not reported here, but is included in the shared dataset). During acquisition, MEG and EEG data were bandpass filtered (0.01 and 330 Hz) and sampled at 1000 Hz. The location of the head fiducials, the shape of the head, the positions of the 64 EEG electrodes and the head position indicator (HPI) coil locations relative to anatomical landmarks were collected with a 3-D digitizer system (Polhemus Isotrack). ECG was recorded with a set of bipolar electrodes placed on the subject’s chest. Two sets of bipolar electrodes were placed around the eyes (two at the outer canthi of the right/left eyes and two above/below the center of the right eye) to record eye movements and blinks (EOG). Ground and reference electrodes were placed on the back of the neck and on the right cheek, respectively. Subjects’ head position on the MEG system was measured at the beginning and end of each run, and also before and after each resting period, using four HPI coils placed on the EEG cap, next to the left and right mastoids and over left and right frontal areas.

#### Anatomical MRI data acquisition

For source localization of the MEG data with individual realistic head modeling, a high resolution T1-weighted (T1w) MRI volume (3T Siemens MRI Prisma scanner) was acquired per subject. Anatomical scans were acquired either with a 32-channel coil (TR/TE = 2000/2.03ms; TI = 880 ms; 8° flip angle; FOV = 256×256×208 mm; 208 slices; 1 mm isotropic voxels, UB) or a 64-channel coil (TR/TE = 2530/2.98ms; TI = 1100 ms; 7° flip angle; FOV = 256×256×208 mm; 198 slices;1 mm isotropic voxels, PKU). The FreeSurfer standard template was used (fsaverage) for participants lacking an anatomical scan (N=5).

#### fMRI data acquisition

MRI data were acquired using a 32-channel head coil on a 3T Prisma scanner. A session included high-resolution anatomical T1w MPRAGE images (GRAPPA acceleration factor = 2, TR/TE = 2300/3.03 ms, 8° flip angle, 192 slices, 1 mm isotropic voxels), and a whole-brain T2*-weighted multiband-4 sequence (TR/TE = 1500/39.6 ms, 75° flip angle, 68 slices, voxel size 2 mm isotropic, A/P phase encoding direction, FOV = 210 mm, BW = 2090 Hz/Px). A single band reference image was acquired before each run. To correct for susceptibility distortions, additional scans using the same T2*-weighted sequence, but with inverted phase encoding direction (inverted RO/PE polarity) were collected while the subject was resting at multiple points throughout the experiment.

### Preprocessing and analysis details

For readability, we first detail the preprocessing protocols for each of the modalities (iEEG, MEG, and fMRI) separately. Then, we describe the different analyses, combining information across the modalities, while noting any differences between them.

### iEEG preprocessing

Data were converted to BIDS^5^ and preprocessed using MNE-Python version 0.24^6^, and custom-written functions in Python and Matlab. Preprocessing steps included downsampling to 512 Hz, detrending, bad channel rejection, line noise and harmonic removal, and re-referencing. Electrodes were re-referenced to a Laplacian scheme^7^ while bipolar referencing was used for electrodes at the edge of a strip, grid or sEEG and the signal was localized at the midpoint (Euclidean distance) between the two electrodes. Electrodes with no direct neighbors were discarded. Seizure onset zone electrodes, those localized outside the brain, and/or containing no signal or high amplitude noise level were discarded. Line noise and harmonics were removed using a one pass, zero-phase non-causal band-stop FIR filter.

The high gamma power (HG, 70-150 Hz) was obtained by bandpass filtering the raw signal in 8 successive 10 Hz wide frequency bands, computing the envelope using a standard Hilbert transform, and normalizing it (dividing) by the mean power per frequency band across the entire recording. To produce a single HG envelope time-series, all frequency bands were averaged together^8^. Most analyses focused on the HG power as it closely correlated with neural spiking activity^9^ and with the BOLD signal ^10^. To obtain the Event Related Potentials (ERPs), the raw signal was low pass filtered at 30 Hz with a one pass, zero-phase non causal low pass FIR filter. Epochs were segmented between 1 s pre-stimulus until 2.5 s post-stimulus of interest.

#### Surface reconstruction and electrode localization

Electrode positions were determined based on a computed tomography scan coregistered with a pre-implant T1 weighted MRI. A three-dimensional reconstruction of each patient’s brain was computed using FreeSurfer (http://surfer.nmr.mgh.harvard.edu). For visualization, the individual subject’s electrode positions were converted to Montreal Neurological Institute (MNI)152 space. As each theory specified a set of anatomical regions of interest (ROIs), after electrode localization, electrodes were labeled according to the Freesurfer based Destrieux atlas segmentation^11,12^ and/or Wang atlas segmentation^13^.

#### Identification of task responsive channels

To identify task responsive electrodes, we computed the Area Under the Curve (AUC) for the baseline (−0.3-0 s) and the stimulus-evoked period (0.05-0.35s) separately for the task relevant and irrelevant conditions, and compared them per electrode using a Wilcoxon sign-rank test, corrected for False Discovery Rate (FDR^14^). A Bayesian t-test^15^ was used to quantify evidence for non-responsiveness.

#### Identification of category selective channels

To determine category selectivity for faces, objects, letters and false fonts on the HG, we followed the method of Kadipasaoglu and colleagues^16^. Per category, we computer a d’ (AUC, 0.05 −0.4 s) comparing the activation between the category-of-interest (u_j_) and each of the other categories (u_i_), normalized by the standard deviation of each category:

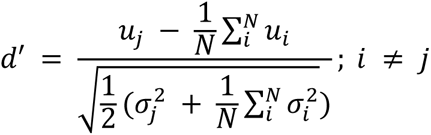

A permutation test (10,000 permutations) was used to evaluate significance. d’ was computed for the task relevant and irrelevant conditions, separately. An electrode was considered selective if it showed selectivity on both tasks.

#### Multivariate analysis electrodes combination

Due to the sparse and highly variable coverage of iEEG data, all collected electrodes were combined into a “super subject” multivariate analyses (RSA and decoding). To create a single trial matrix for the super subject, we equated the trial matrices of all our subjects by subsampling to the lowest number of trials in the relevant conditions. Subjects that did not complete the full experiment were discarded (N=3), resulting in a total of 29 subjects with 583 electrodes in posterior and 576 electrodes in prefrontal ROIs, respectively. In the case of analyses on stimuli identities, stimuli that were presented less than three times to any of the participants across intermediate and long trials in the task relevant and irrelevant trials were discarded. We then subsampled the trials for each identity to three trials per participant. The subsampling procedure was repeated 100 times to avoid random fluctuation induced by the subsampling. The analysis was computed for each repetition and average across repetitions.

### MEG preprocessing

The MEG data were converted to BIDS^17^ using MNE-BIDS^18^, and preprocessed following the FLUX Pipeline^19^ in MNE-Python v0.24.0^6^. Preprocessing steps included MEG sensor reconstruction using a semi-automatic detection algorithm and Signal-Space Separation (SSS)^20^ to reduce environmental artifacts. FastICA^21^ was used to detect and remove cardiac and ocular components from the data for each subject (M=2.90 components, SD=0.92). Prior to ICA, data were segmented, and segments containing muscle artifacts were removed. After preprocessing, data were epoched into a 3.5 s segment (1 s pre-stimulus to 2.5 s post-stimulus onset). Trials where gradiometers values exceeded 5000 fT/cm, magnetometers exceeded 5000 fT, and/or contained muscle artifacts were rejected from the MEG dataset.

#### Source modeling

MEG source modeling was performed using the dynamic statistical parametric mapping (dSPM) method^22^, based on depth-weighted minimum-norm estimates (MNE^23,24^), on epoched and baseline (−0.5 s to 0 s prior to stimulus onset) corrected data. To build a forward model, the MRI images were manually aligned to the digitized head shape. A single shell Boundary Elements Model (BEM) was constructed in MNE-Python based on the inner skull surface derived from FreeSurfer ^11,12^, to create a volumetric forward model (5 mm grid) covering the full brain volume. The lead field matrix was then calculated according to the head-position with respect to the MEG sensor array. A noise covariance matrix for the baseline and a covariance matrix for the active time window were calculated and the combined (i.e., sum) covariance matrix was used with the forward model to create a common spatial filter. Data were spatially pre-whitened using the covariance matrix from the baseline interval to combine gradiometer and magnetometer data ^25^.

### fMRI Preprocessing

Source DICOM data were converted to BIDS using BIDScoin v3.6.3^26^. This includes converting DICOM data to NIfTI using dcm2niix ^27^ and creating event files using custom Python codes. BIDS compliance of the resulting dataset was controlled using BIDS-Validator. Subsequently, MRI data quality control was performed using MRIQC ^28^ and custom scripts for data rejection. All (f)MRI data were preprocessed using fMRIPrep 20.2.3^29^, based on Nipype 1.6.1^30^. For further details on the fMRIprep pipeline, see preregistration.

#### Analysis-specific functional preprocessing

Additional, analysis-specific, fMRI data preprocessing was performed using FSL 6.0.2 (FMRIB Software Library; Oxford, UK^31^), Statistical Parametric Mapping (SPM 12) software^32^, and custom Python scripts after the above outlined general preprocessing. Functional data for univariate data analyses were spatially smoothed (Gaussian kernel with full-width at half-maximum of 5 mm), grand mean scaled, and temporal high-pass filtered (128 s). No spatial smoothing was applied for multivariate analyses.

#### Contrast of parameter estimates

We modeled BOLD signal responses to the experimental variables by fitting voxel-wise General Linear Model (GLM) to the data of each run using FSL FEAT. The following regressors were modeled in an event-related approach, with event duration corresponding to the stimulus duration (i.e., 0.5, 1.0, 1.5 s), and convolved with a double gamma hemodynamic response function: 12 regressors of interest (Targets, task relevant and task irrelevant stimuli per stimulus category i.e., faces, objects, letters, false fonts; and a regressors of no interest i.e., target screen display). We included the first-order temporal derivatives of the regressors of interest, and a set of nuisance regressors: 24 motion regressors (FSL’s standard + extended set of motion parameters) plus a CSF and a WM tissue regressor.

Each of the 12 regressors of interest was contrasted against an implicit baseline (used in the putative NCC analysis). Additionally, we obtained contrast of parameter estimates for ‘relevant faces vs. relevant objects’, ‘relevant letters vs. relevant false fonts’, ‘irrelevant faces vs. irrelevant objects’, ‘irrelevant letters vs. irrelevant false fonts’ (used for the definition of decoding ROIs), ‘relevant and irrelevant faces vs. relevant and irrelevant objects’ and ‘all stimuli vs. baseline’ (used for the definition of seeds for the generalized psychophysiological interaction analysis).

Data were averaged across runs per subject using FSL’s fixed effects analysis and subsequently averaged across participants using FSL’s FLAME1 mixed effect analysis. Gaussian random-field cluster thresholding was used to correct for multiple comparisons, using the default settings of FSL, with a cluster formation threshold of one sided p < 0.001 (z ≥ 3.1,) and a cluster significance threshold of p < 0.05.

### Anatomical Regions-of-interest (ROIs)

ROIs were defined a priori in consultation with the adversaries. They were determined per subject based on the Destrieux atlas^12^ including both hemispheres, and then resampled to standard MNI space (see Extended Data Table 2). For the connectivity analysis, areas V1/V2 (combining dorsal and ventral) were defined based on the Wang cortical parcellation^13^.

### Behavioral analyses

Log-linear corrected d’prime^33^, false alarms (FA) and reaction times (RT) were computed per category and stimulus duration, separately (FAs were also calculated per task relevance, without duration), and per modality (iEEG, MEG, fMRI). These measures were compared with Linear/Logistic mixed models, where appropriate. For the former, we report ANOVA omnibus F tests, and for the latter, omnibus χ² test from an analysis of deviance. We approximated degrees of freedom using the Satterthwaite method^34^. Pairwise t-tests following significant interactions were Bonferroni corrected. To estimate Bayesian Information Criterion (BIC) differences between the original and null logistic models, we used the p-values and sample size (^35^; p_to_bf package in R).

### Eye-tracking analyses

For Eyelink, gaze and pupil data were segmented, and missing data were excluded. Blinks were detected using the Hershman algorithm^36^, and removed with 200 ms padding^37^. The Eyelink standard parser algorithm was used for saccade and fixation detection. Saccades were further corroborated using the Engbert & Kliegl^38^ algorithm. Fixations were baseline corrected (−0.25 s to 0 s). Mean fixation distance, mean blink rate, mean saccade amplitude and mean pupil size were compared in a Linear Mixed Model (LMM) with category and task relevance as fixed effects and subject and item as random effects. Separate analyses were carried out on the first 0.5 s after stimulus onset including all trials; and on the 1.5 s trials including time window (0-0.5 s, 0.5-1.0 s, 1.0-1.5 s) as fixed effects. BIC was used to test the models against the null hypothesis models. For Tobii, gaze coordinate data was segmented, missing data were excluded, and coordinates were baseline corrected to depict heatmaps of patients’ gaze. Notably, the coordinate data was not added to the LMMs due to its poorer quality with respect to the EyeLink data.

### Decoding analysis

All decoding analyses were performed using a linear Support Vector Machine (SVM, scikit learn, https://scikit-learn.org/) classifier. Below we explain how this was done for each one of the predictions.

iEEG Decoding was done on the HG response, averaged over non-overlapping windows of 0.02 s separately for electrodes located in the GNWT and IIT ROIs. The top 200 electrodes (selectKbest^39^), as determined by F-test within a given set of electrodes from the theory ROIs, were used as features for the classifier. 200 features were selected to provide a balance between model optimization (e.g., feature selection) and subject representation (e.g., electrodes/features coming from multiple subjects). Statistical significance of decoding performance was assessed via permutation test, randomly permuting the sample labels and repeating the decoding analysis 1000 times, corrected for multiple comparisons using a cluster-based correction (cluster mass inference with cluster forming threshold at p < 0.05^40,41^). Also, to assess the decoding accuracy within unique ROIs (e.g., S_temporal_sup of the Destrieux atlas), separate classifiers were trained using all electrodes in a given parcel. Each classifier was fitted using all electrodes in a parcel and time window (GNWT: 0.3-0.5 s, IIT: 0.3-1.5 s) as features, resulting in a single accuracy value per parcel. SelectKbest (200 features iEEG) feature selection and 5-fold cross-validation with 3 repetitions was used. To assess the statistical significance of the decoding accuracy within unique ROIs (so only one accuracy score is obtained per ROI), p-values obtained via permutation tests were corrected for multiple comparisons across all ROIs using FDR correction (q ≤ 0.05^14^).

MEG Decoding was done on bandpass filtered (1-40 Hz) and downsampled (100 Hz) data. The reconstructed source-level MEG data within a subset of the predefined anatomical ROIs (GNWT: ‘G_and_S_cingul-Ant’,‘G_and_S_cingul-Mid-Ant’, ‘G_and_S_cingul-Mid-Post’, ‘G_front_middle’,‘S_front_inf’, ‘S_front_sup’, IIT: ‘G_cuneus’, ‘G_oc-temp_lat-fusifor’, ‘G_oc-temp_med-Lingual’,‘Pole_occipital’, ‘S_calcarine’,‘S_oc_sup_and_transversal’, as they show high response to the stimulus on the optimization dataset) were extracted for further analysis (500 vertices and 800 vertices per hemisphere for each of the anatomical ROI defined by the theories). We applied temporal smoothing (0.05 s window, 0.01 sliding window), computed pseudotrials^42^, normalized the data, and selected the top 30 features within a given ROI as features for the different classifiers. A group-level one-sample t-test per time point was performed on the decoding accuracy results, corrected for multiple comparisons using a cluster-based correction^41^.

The overall decoding strategy for fMRI was similar to that used on the iEEG and MEG data, yet with some differences. A Multi-Variate Pattern Analysis (MVPA) approach was used on the pattern of BOLD activity over voxels. A non-spatially-smoothed parameter estimate map was obtained by fitting a GLM per event with that event as the regressor of interest and all the other remaining events as one regressor of no interest^43^ as implemented in NiBetaSeries 0.6.0 package. The model also included the 24 nuisance regressors described in the fMRI preprocessing section.

Decoding was performed using a whole-brain approach and an ROI-based approach. The whole-brain analysis was performed using a searchlight approach with 4 mm radius. For ROI-based decoding, decoding ROIs were defined based on functional fMRI contrasts (see fMRI preprocessing section) and constrained with pre-defined anatomical ROIs (see Extended Data Table 2: Anatomical Regions-of-interest (ROIs)). One-sample permutation test was used to determine if decoding significantly exceeds chance level within each ROI. FDR was used to correct for multiple comparisons across ROIs. For whole-brain decoding, a cluster-based permutation test was used to evaluate the decoding statistical significance across subjects (p < 0.05). Additionally, stimulus vs. baseline searchlight decoding was performed using leave-one-run out cross validation and the resultant decoding accuracy maps were used as input for the multivariate putative NCC analysis (see below). To perform stimulus vs. baseline decoding, we subsampled the stimuli trials to a 2:1 ratio with respect to baseline. The SVM cost function was weighted by the number of trials from each class.

#### Decoding schemes for the different predictions

To test GNWT and IIT decoding predictions, stimulus category (faces vs. objects and letters vs. false fonts) was decoded separately for the task relevant and task irrelevant conditions (*within-task category decoding*) while orientation (front view vs. left view vs. right view) was decoded on the combined data from the two task conditions. In addition, *cross-task category decoding* from task relevant to task irrelevant condition and vice versa was performed to test generalization by training classifiers on one condition and testing on the other condition. Both within-task category and orientation decoding were performed in a leave-one-run-out cross validation scheme for fMRI and in an k-fold cross validation scheme for MEG and iEEG.

For *category decoding*, trials from each task condition (i.e., task relevant, irrelevant) were extracted for each category comparison of interest: 160 face/160 objects classification, 160 letters/160 false fonts classification within each task relevance condition for MEG, and half the trials for iEEG. For fMRI, there were 64 trials for each category in each task relevance condition. For *orientation decoding*, task relevant and task irrelevant trials were collapsed within category to increase Signal-to-Noise Ratio (SNR), resulting in 160 Front, 80 Left, and 80 Right trials per category for MEG, and half these numbers for iEEG. For fMRI, there were 64 Front, and 32 Left and Right trials per category. Decoding was evaluated using accuracy measures, tested against 50% chance level for category decoding (binary classification) and against 33% chance level for orientation decoding (3-class classification). For orientation decoding, balanced accuracy was used due to the unbalanced number of trials for the different orientations. The SVM cost function was weighted by the number of trials per class to reduce bias to the class with the highest number.

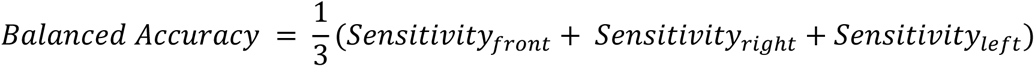

For *within-task decoding* (e.g., classification of categories across time), a classifier at each time-point was trained and tested separately using a 5-fold cross-validation (with 3 separate repeats of cross-validation). For cross-task decoding (task relevant −> irrelevant & task irrelevant −> relevant), each SVM model was trained on one task (e.g., faces/objects in the task relevant condition) and tested on the second task (e.g., faces/objects in the task irrelevant one). As cross-decoding in iEEG data is performed across all pooled electrodes, an additional cross-validation step was performed on this modality data to provide a confidence metric (e.g., confidence intervals) using a 5-fold cross-validation with 3 repetitions (e.g. train on 80% of task 1, and test on held-out 20% of task 2).

*Within-task temporal generalization* was performed by training a classifier at each time-point (using selectKbest feature selection) and testing its performance across all time-points using the same set of selected features and 3 repetitions of 5-fold cross-validation. To generalize from one task to another across all time-points, cross-temporal generalization was used: a classifier was trained at each time-point in task 1 (e.g., task relevant) using selectKbest feature selection, and tested across all time-points in task 2 (e.g., task irrelevant) using the same set of selected features. Cross-validation was performed in the same fashion as in cross-decoding.

Additional decoding analyses were performed on all trials aligned to the stimulus onset (e.g. −0.2-2 s relative to stimulus onset), and stimulus offset (−0.5-0.5 s around stimulus offset). For the latter analysis, all trials from different durations were aligned to the stimulus offset.

To assess the specific IIT prediction that including prefrontal regions along with posterior regions to the decoding of categories will not significantly affect decoding accuracy, we performed two additional decoding analyses in which the decoding performance of electrodes from the IIT region were compared with the decoding performance when electrodes from both the posterior + PFC ROIs are included. The PFC ROI included all PFC ROIs, except for inferior frontal sulcus, as it belongs to the IIT extended ROIs. Posterior ROI included all IIT ROIs shown in Extended Data Table 2. The first analysis compared the decoding accuracy for a model including all electrodes from posterior regions to a separate model in which electrodes (features) from posterior & PFC regions were combined (e.g., feature combination). In the second analysis, the decoding accuracy of the model including all electrodes from posterior regions was compared to a combined posterior + PFC model, in which two separate classifiers were trained and calibrated on posterior & PFC regions separately using isotonic calibration^44^, and posterior probabilities from each classifier were combined using a softmax normalization^45^. Training and testing of the individual models followed all previously described cross-validation procedures and model comparison was performed using a variance-corrected paired t-test^46^ and complemented with Bayesian analysis. Following Benavoli and colleagues^47^, the prior distribution of the mean difference in decoding scores between two classifier models was modeled as a Normal-gamma distribution conjugate to a normal likelihood, and the posterior distribution was obtained as a normal distribution. This posterior distribution was utilized to calculate the probability of one classification model being better than, worse than, or equivalent to the other model. As this estimation approach is applied using resampled datasets (e.g., using 5-fold cross-validation), the performance of the model becomes dependent on the folds, and thus a variance corrected t-distribution was used^46^.

We also tested this prediction on the fMRI data. To select features to be used for both analyses, the face vs. object contrast for each subject was masked by a predefined anatomical posterior ROIs as well as a PFC anatomical ROIs, defined the same way as described above. Within each of the two ROIs, the 150 voxels that are most selective to each of the to-be-decoded stimuli were defined as the decoding ROIs (300 voxels total) for each subject. The first analysis compared the decoding accuracies for a model that included 300 voxels from the posterior ROIs as features to another model that included 600 voxels (300 features from each ROI). In the second analysis, two separate models were constructed, calibrated, and combined as described above. For the two analyses, model comparison was performed using a group-level one-sample permutation test to determine if accuracies obtained by combining posterior and PFC ROIs are significantly higher than the accuracies obtained based on posterior ROIs only. FDR was used to correct for multiple comparisons.

### Duration analysis

Neural responses were extracted from three windows of interest (WoI) (0.8-1.0 s, 1.3-1.5 s, 1.8 −2.0 s) and compared using LMM. Four theory agnostic models were fitted: a null model, a duration model (3 durations), a WoI model, and a duration and WoI model. Two theory model were fitted: the GNWT model predicts activation (ignition) following stimulus offset (0.3-0.5 s) independent of duration, with virtually no response in between. The IIT model predicts sustained activation for the duration of the stimulus returning to baseline after stimulus offset. Both theoretical models were complemented with an interaction term between category (faces, objects, letters and false fonts) and the theories’ predictors, to account for regions showing selective responses to categories. Bayesian Integration Criterion (BIC) was used to define the winning model.

Models for iEEG were fitted per electrode on the predefined ROIs, using the HG (AUC), alpha (8-13 Hz, obtained through Morlet wavelets, f=8-13 Hz, in 1 Hz steps; f/2 cycles, AUC), and ERPs (peak to peak) as signal, separately for task relevant and irrelevant condition.

MEG models were fitted to source data on the predefined ROIs, using the gamma (60-90 Hz) and alpha band (8-13 Hz) as signal, separately for task relevant and irrelevant conditions. Time-frequency analyses were performed on source-data using Morlet wavelets (f=8-13 Hz, in 1 Hz steps; f/2 cycles; f=60-90 Hz, in 2 Hz steps, f/4 cycles), and were baseline corrected. Spectral activity was computed for each vertex, baseline corrected and then averaged across trials within each parcel included in the ROIs, yielding a unique time-course per ROI parcel. In addition, a single source time-course capturing the entire prefrontal ROI and the posterior ROI was computed by averaging the spectral activity within an ROI. Models were fitted on each parcel and ROI, as defined by the theories.

### Representational Similarity Analysis (RSA)

To examine how the neural representations evolved over time in response to the different stimulus properties (i.e., category, orientation and identity representation), we performed cross-temporal RSA on source level MEG data and iEEG HG power within each of the theory-defined ROIs. Specifically, at each set of data points, we computed a Representational Dissimilarity Matrix (RDM) by calculating the correlation distance (1-Pearson’s r, Fisher corrected) between all pairs of stimuli. Next, to quantify the representational space occupied by one class vs. another, we computed the average within-class distances vs. the average between-class distances. This analysis was performed in a cross-temporal manner, in which RDMs were computed between all stimuli at time point t1 and the corresponding set of stimuli at time points t1,2,…n.

Long trials (1.5 s) were used to investigate category and orientation representation. Since specific identities were repeated a limited number of times per duration, both intermediate (1.0) and long (1.5 secs) trials were combined and equated in duration by cropping the 1-1.5s time interval for long trials. This was done to allow for the analysis of at least three (3) presentations of the same identity.

To evaluate the theoretical predictions about when significant content representation should occur, we subsampled the observed cross-temporal representational matrices in four time windows (0.3-0.5, 0.8-1.0, 1.3-1.5, 1.8-2.0 s). The subsampled matrices were correlated to the model matrices predicted by GNWT and IIT (see Figure 1a, right panel) using Kendall’s Tau correlation. If the correlation was significant (see below) for at least one of the predicted matrices, we computed the difference between the transformed correlation ((*r* + 1) / 2) to each theory; and compared this difference against a random distribution to obtain a p-value. If the correlation with the theory predicted pattern in the theory ROI was significantly higher than the other model, we considered the theory prediction to be fulfilled.

To generate a null distribution of cross-temporal RSA surrogate matrices, we repeated the procedure outlined above 1024 times, randomly shuffling the labels. Next, the observed RSA matrix was z-scored using the null distribution as:

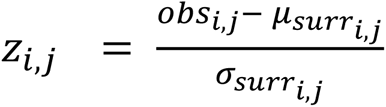

Where *obs_i,j_* is the observed within-vs.-between class difference at time points i and j, and μ_*surri,j*_, and σ_*surri,j*_ are the mean and standard deviation of the surrogate representational similarity matrix at time points i and j, respectively. Cluster based permutation tests ^48^, z-score threshold of z = 1.5 for clustering, were used to evaluate significance. RSA surrogates were also used to assess the significance of the correlation between the observed matrices and the theories’ predicted matrices. First, a null distribution of possible correlations was generated for each of the theories by correlating each of the surrogate matrices to each of the theory predicted matrices. Next, a p-value was obtained for each theory predicted matrix, by locating its observed correlation within the null correlation distribution. The same procedure was used to assess the significance of the difference in correlation to IIT and GNWT matrices (e.g., each of the surrogate matrices was correlated to each of the theory predicted matrices and the difference between the two was computed). P-values were FDR corrected (q ≤ 0.05)^14^.

For iEEG, the HG power per electrode within the predefined anatomical ROI was averaged in 0.02s non-overlapping windows. Electrodes were used as features for the RDM. The data were vectorized across all electrodes within a ROI (e.g., samples x significant electrodes) to compute the RDMs. 576 and 583 electrodes entered this analysis for the prefrontal and posterior ROI, respectively. The resultant RDM was subjected to a principal component analysis and the first two dimensions were plotted against each other to produce a 2-dimensional projection of dissimilarity scores across all pairs for each of the 100 subsampling repetitions. The PCA components were aligned across repetitions using Procrustes alignment and averaged together for visualization purposes^49,50^.

For MEG, the same analysis was run on the source reconstructed data within the predefined anatomical ROIs used for the Decoding analysis, bandpass filtered (1-40 Hz) and downsampled (100 Hz). For the category and orientation analysis, pseudo-trials and temporal moving-average methods were used to optimize the RSA analysis and improve the SNR. For identity, single trials were used. Vertices within the ROIs were used as features. The statistical testing differed from that conducted on the iEEG data, as it was performed at the subject level. Like the iEEG analysis, we first tested if the correlation between the data and the model predicted by each theory was greater than zero using the Kendall’s tau measure, and then compared between the theories using the Mann-Whitney U rank test on two independent samples.

### Functional Connectivity analysis

For both iEEG and MEG, pairwise phase consistency (PPC^51^) was computed between each category-selective time series (face- and object-selective) and either the V1/V2 or the PFC time series.

For iEEG, the PPC analysis included electrodes in V1/V2 visual areas, in PFC ROIs (see Extended Data Table 2), and face and object selective electrodes (see *Identification of task responsive channels*), as long as they were “active” during the task. As both theories predict different types of activation (e.g., ignition vs. sustained activation), channels were categorized as active if they showed an increase in HG power relative to baseline (−0.5 to −0.3 s, p<0.05, signed-rank test) evaluated across all trials (task relevant + irrelevant, intermediate + long trials, combined across both categories), for the 0.3-0.5 s window (GNWT), or in all time windows 0.3-0.5 s, 0.5-0.8 s, and 1.3-1.5 s (IIT).

For MEG, the category-selective single-trial time courses used to define the ROIs for PPC analysis were extracted using the Generalized Eigenvalue Decomposition (GED) method^52^. Two GED spatial filters were built by contrasting either faces or objects against all other categories during the first 0.5 s after stimulus onset. Single-trial covariance matrices were computed separately for signal and reference for all vertices within the fusiform ROI identified from the FreeSurfer parcellation using the Desikan atlas ^53^, and the Euclidean distance between them was z-scored. Trials exceeding 3 z-scores were excluded. The reference covariance matrix was regularized to reduce overfitting and increase numerical stability. The GED was then performed on the two covariance matrices, resulting in N (= rank of the data) pairs of eigenvectors and eigenvalues. The eigenvector associated with the highest eigenvalue was selected as a GED spatial filter, which in turn was applied to the data to compute the single-trial GED component time series. A GED spatial filter was extracted also for the PFC ROI, on parcels from the Destrieux atlas^12^, to identify the distributed pattern of sources that are responsive to visually-presented stimuli. Specifically, a spatial filter was built by contrasting source-level frontal slow-frequency activity (30-Hz low-pass filter) after stimulus onset (0 to 0.5 s) against baseline (−0.5 to 0 s). V1/V2 areas were identified using the Wang Atlas^13^ and a singular values-decomposition approach. For the GED, the 1.0 and 1.5 s duration trials were used to minimize overlap with the transient evoked at stimulus onset.

PPC was computed for each MEG time series/iEEG electrode pairing, for all face-trials and object-trials separately. Analyses were performed on 1.0 and 1.5 duration trials, separately on task relevant and irrelevant trials and also combined to maximize statistical power. To compute synchrony, time-frequency analysis of the broadband MEG and LFP signal was performed using Morlet wavelets (f=2-30 Hz, in 1 Hz steps; 4 cycles; f=30-180 Hz for iEEG or f=30-100 Hz for MEG, in 2 Hz steps, f/4 cycles), and PPC was then computed by taking the difference in phase angle between MEG time series/iEEG electrode at each time, t, and frequency f, for a specific trial and computing PPC across all trials in a category (e.g., faces) as:

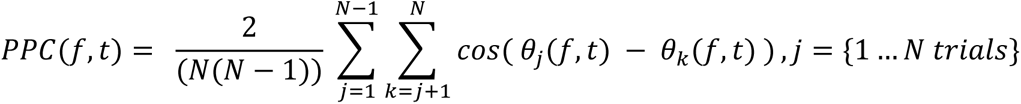

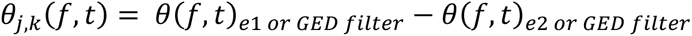, for all frequencies f, and at all times t.

For iEEG, PPC for each category-selective site was then averaged across all its pairings (e.g., all PFC electrodes pairings or all V1/V2 pairings within that patient). The variability in electrode coverage across patients precluded a within-subjects analysis. Therefore, to achieve sufficient statistical power, we pooled all derived PPC values from one electrode pairing (e.g., face-selective to PFC) across all patients into one ROI specific analysis. A similar approach was used on the MEG parcels.

To quantify content-specific synchrony enhancement, the difference in PPC was computed between within-category and across-category trials (e.g., for face-selective sites, the change in PPC was computed between faces vs. objects trials) using a cluster-based permutation test^41^. This was done for both modalities.

As an exploratory analysis, we also investigated dynamic functional connectivity using the Gaussian-Copula Mutual Information (GCMI^54^) approach to evaluate the dependencies between time series. This power-based measure of connectivity was implemented using the conn_dfc method from the Frites Python package ^55^. We used the same parameters as for the PPC analysis, with the following exceptions: For both MEG and iEEG, power was estimated through a multitaper-based method (using a frequency dependent dynamic sliding window: 2-30 Hz, T= 4 cycles; 30-100 Hz, T4/f using a 0.25-s sliding window. For iEEG the high frequency range was extended from 30-180 Hz, T=4/f cycles). DFC was performed per frequency band, 0.1 s sliding window, 0.02s steps.

For fMRI, connectivity was assessed through generalized Psycho-Physiological Interaction (gPPI) implemented in SPM^56^. The Fusiform Face Area (FFA) and Lateral occipital cortex (LOC) were defined as seed regions per subject based on an anatomically constrained functional contrast. Anatomically, FFA seeds were constrained to the “Inferior occipital gyrus (O3) and sulcus” and “Lateral occipito-temporal gyrus (fusiform gyrus, O4-T4)”. LOC seeds were constrained to the “Middle occipital gyrus (O2, lateral occipital gyrus)” and the “Middle occipital sulcus and lunatus sulcus” (Destrieux ROIs 2 and 21 for FFA and ROIs 19 and 57 for LOC, see Anatomical Regions-of-interest (ROIs)).

Candidate seed voxels within the above-mentioned anatomical ROIs were defined as those with a z value > 1 in the contrast of parameter estimates of all stimuli vs. baseline. Three subjects with less than 300 candidate seed voxels were excluded from the analysis. This was done to ensure that the seed voxels were visually driven. Next, using an unthresholded contrast of parameter estimates between ‘relevant and irrelevant faces’ and ‘relevant and irrelevant objects’, the 300 voxels most responsive to faces within the FFA anatomical ROIs were selected for the FFA seed, and the 300 voxels most responsive to objects within the LOC anatomical ROIs were selected for the LOC seed.

gPPI analysis was performed per subject and seed region separately, including an interaction term between the seed time series regressor (physiological term) and the task regressor (psychological term) at the subject-level GLM^56^, separately for task relevant and irrelevant conditions, and also combining across tasks to increase statistical power. For combined conditions, the model design matrix for each subject included regressors for task relevant and task irrelevant faces, objects, letters, and false fonts collapsed across conditions (four regressors) as well as a regressor for targets (irrespective of their category), yielding five regressors in total. As for separated conditions, the model design matrix included regressors for task relevant and task irrelevant faces, objects, letters, and falsefonts (eight regressors) as well as a regressor for targets (irrespective of their category), yielding nine regressors in total. For each seed, group level analysis was performed using a cluster-based permutation test to evaluate the statistical significance of face > object contrast parameter estimates across subjects (p < 0.05).

### Putative NCC analyses

A series of conjunction analyses were performed on the fMRI data to identify a) areas responsive to task goal, b) areas responsive to task relevance, and c) areas putatively involved in the neural correlate of consciousness. We note that the contrasts proposed below might overestimate the neural correlates of consciousness and that the fast event-related design adopted here might be suboptimal to detect activity changes in the salience network^57^, i.e., potentially underestimating some regions that might be involved in conscious processing. We therefore have adopted a conservative approach that distinguishes between areas that might participate in consciousness vs. those that definitely do not.

The conjunction defining *areas responsive to task* goals was defined as [TaskRelTar > bsl] & [(TaskRelNonTar = bsl) & (TaskIrrel = bsl)]. This contrast captures areas that show an increase of BOLD signal for targets but not for other stimuli. The following conjunction identified *areas responsive to task relevance*: [(TaskRelTar > bsl) & (TaskRelNonTar ≠ bsl)] & [TaskIrrel = bsl]. This contrast identifies areas displaying differential activity for all task relevant stimuli, but are insensitive to non-task relevant ones. Finally, the following conjunction was used to identify the *putative NCC areas*: [(TaskRelNonTar (stim id) > bsl) & (TaskIrrel (stim id) > bsl)] OR [(TaskRelNonTar (stim id) < bsl) & (TaskIrrel (stim id) < bsl)], critically detecting areas that responsive to any stimulus category irrespective of task, with consistent activation or deactivation.

To compute conjunctions, we first ran a GLM (see above) corrected for multiple comparisons (Gaussian random-field cluster-based inference). Equivalence to baseline was established using a JZS Bayes Factor test, with a Cauchy prior (r scale value of 0.707). Evidence maps were thresholded at BF01 > 3. The thresholded z maps and the Bayesian evidence maps on the group level were used for the conjunction analysis. For conjunctions including an ‘unequal to’, a ‘logical and’ operation was used between the directional z maps, after thresholded maps were binarized. For the putative NCC contrast, conjunctions were performed separately for activations and deactivations, using a ‘logical and’ operator for the task relevant and irrelevant z maps. The resulting maps were combined using a ‘logical or’ operation to discard areas showing effects of opposite direction for task relevant and task irrelevant stimuli. This analysis was also done at the subject level, masked using the anatomical ROIs, to account for inter-subject variability. For each ROI, the proportion of subjects with voxels included in the conjunction is reported. The multivariate version of the putative NCC analysis was done using the thresholded statistical maps obtained from the whole-brain searchlight decoding based on a subject-level stimulus vs. baseline decoding accuracy maps (for details regarding the decoding approach used, see *Decoding Analysis*).

## Data availability

Raw behavioral data, Raw iEEG, M-EEG, Imaging and Eye tracking data, unthresholded group-level statistical brain maps from neuroimaging analyses and source data to reproduce all figures will be made publicly available upon publication.

## Code availability

Task and analysis code will be publicly available upon publication here: https://github.com/Cogitate-consortium/cogitate-msp1

## Acknowledgements

Special thanks to Dawid Potgieter for spearheading the ARC program; to Daniel Kahneman for guidance to navigate adversarial collaborations; to Heather Berlin, William Jaworski, Hakwan Lau and Cyriel Pennartz for insightful discussions during the two-day meeting organized by the Templeton World Charity Foundation at the Allen Institute, Seattle in March 2018; to Hakwan Lau for helping conceptualize the proposed experiments; to Orrin Devinsky, Werner Doyle, Patricia Dugan and Daniel Friedman for supporting the recruitment and patient care at NYU; to Essa Yacoub, Michael Kahana, Peter Zeidman, Karl Friston, Jean-Remi King, Michael Cohen, Fosca Al Roumi, Shlomit Yuval-Greenberg and Dejan Draschkow for guidance on diverse data analysis; to Caspar Schwiedrzik for insightful discussions and feedback throughout the different phases of this study (conceptualization, data analysis and initial draft); to Sarah Brendecke and Felix Bernoully for help with figures and stimulus materials; to Monique Smulders and Sarah Kusch for help with fMRI data acquisition; to the patients and their families for generously supporting this study.

## Authors contributions

Conceptualization: O.F., A.K., A.L., L.L., D.R., N.B., T.B., P.S., S.B., D.J.C., R.M.C., F.F., F.I.P., H.B., O.J., F.P.L., H.L., M.B., S.D., C.K., G.T., L.M., M.P., L.M. Data curation: O.F., U.G.K., S.H., R.H., A.K., A.L., L.L., D.R., N.B., T.B., P.S., M.A., T.G., D.H., C.K., D.R.M., S.M., A.S., A.S., S.Y., H.B., S.D., O.J., H.L., L.M., L.M. Data Quality: O.F., U.G.K., S.H., R.H., A.K., A.L., L.L., D.R., Y.V., M.A., K.B., T.G., D.H., C.K., S.M., A.S., H.B., S.D., O.J., H.L., M.B., L.M., M.P., L.M. Formal analysis: O.F., S.H., R.H., A.K., A.L., L.L., D.R., Y.V., N.B., K.B., C.K., S.B., R.M.C., H.B., S.D., O.J., H.L., L.M., M.P., L.M. Funding acquisition: C.K., L.M., M.P., L.M. Investigation: O.F., U.G.K., A.K., A.L., L.L., D.R., M.A., K.B., T.G., D.H., J.J., C.K., D.R.M., S.M., A.S., S.Y., H.B., H.L., L.M. Methodology: O.F., U.G.K., S.H., R.H., A.K., A.L., L.L., D.R., Y.V., T.B., K.B., C.K., S.B., R.M.C., F.I.P., H.B. S.D., O.J., F.P.L., H.L. M.B., C.K., L.M., M.P., L.M. Project administration: O.F., U.G.K., S.H., A.K., A.L., L.L., D.R., N.B., T.B., T.G., D.H., S.M., A.S., S.Y., H.B., O.J., G.K., H.L., L.M., M.P., L.M. Resources: O.F., R.H., A.K., A.L., L.L., T.B., S.M., A.S., A.S., S.Y., H.B., S.D., O.J., H.L., L.M., M.P., L.M. Software: O.F., U.G.K., S.H., R.H., A.K., A.L., L.L., D.R., Y.V., N.B., P.S., K.B., J.J., S.M., A.S., A.S., H.B., O.J. Supervision: O.F., S.H., A.K., L.L., D.R., N.B., T.B., K.B., A.S., H.B., S.D., O.J., G.K., F.P.L., H.L., M.B., L.M., M.P., L.M. Validation: O.F., U.G.K., S.H., R.H., A.K., A.L., L.L., Y.V., C.K., H.B., O.J., L.M., M.P., L.M. Visualization: O.F., S.H., R.H., A.K., A.L., L.L., D.R., Y.V., T.B., M.A., H.B., S.D., O.J., L.M., M.P., L.M. Writing – original draft: O.F., U.G.K., S.H., R.H., A.K., A.L., L.L., D.R., Y.V., T.B., F.I.P., H.L., M.B., S.D., G.T., L.M., M.P., L.M. Writing – review & editing: O.F., U.G.K., S.H., R.H., A.K., A.L., L.L., D.R., Y.V., N.B., T.B., M.A., S.B., D.J.C., R.M.C., F.F., F.I.P., H.B., S.D. O.J., F.P.L., H.L., M.B., S.D., C.K., G.T., L.M., M.P., L.M.

## Additional information

Correspondence and requests for materials should be addressed to Lucia Melloni.

## Extended Materials

**Extended Data Figure 1:**
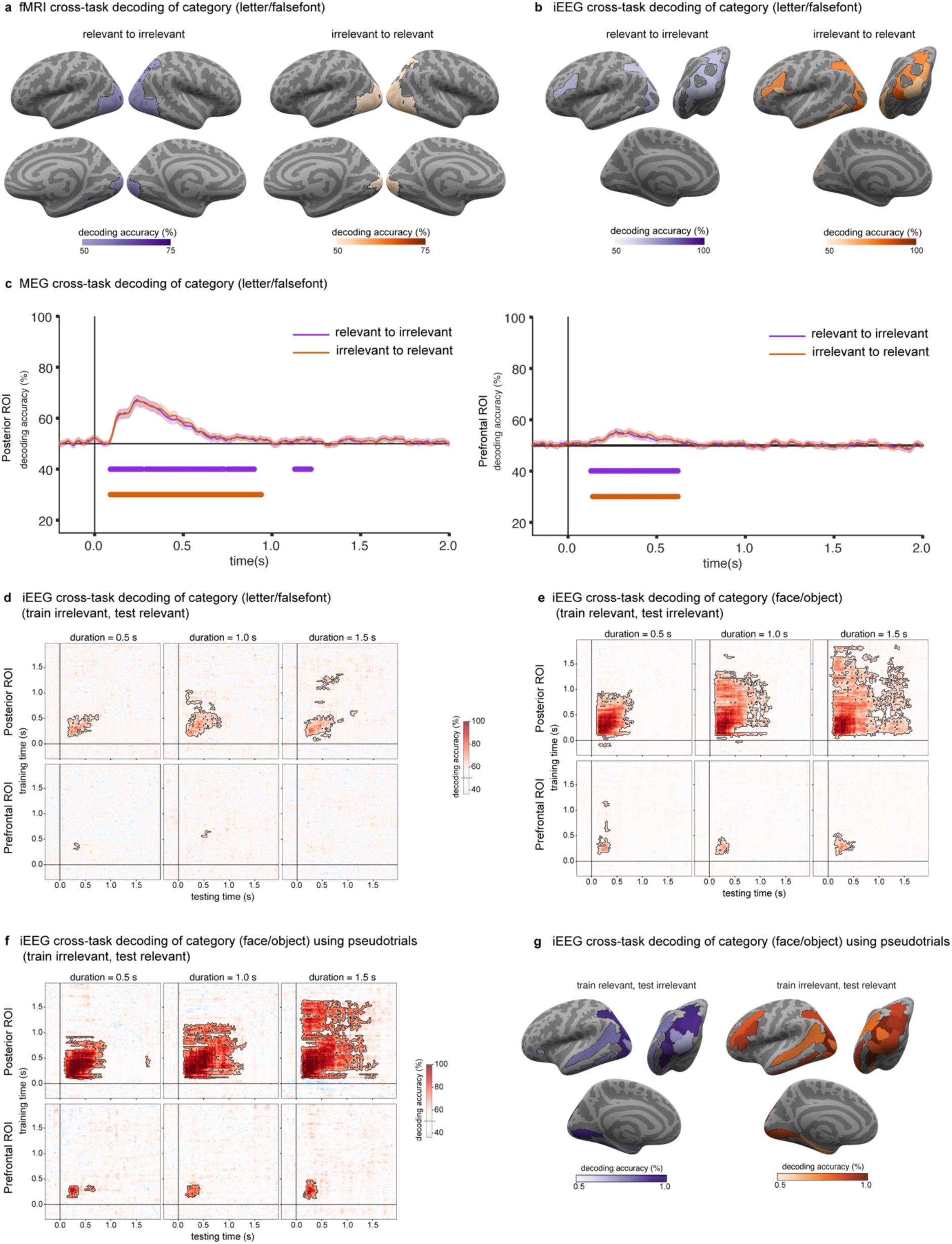
Prediction#1 Decoding of conscious content for letters, false fonts, faces and objects. **a.** fMRI decoding accuracies (letters vs. false fonts) using a searchlight approach, collapsed across the three stimulus durations. Left: decoding for classifiers trained on task relevant and tested on task irrelevant stimuli (purple). Right: decoding for classifiers trained on task irrelevant and tested on task relevant stimuli (orange-red). Regions showing significantly above-chance (50%) decoding accuracies are indicated by the outlined colored regions on the inflated cortical surfaces (top: left/right lateral views; bottom: right/left medial views). **b.** iEEG decoding accuracies (letters vs. false fonts) within the theory-relevant ROIs collapsed across stimulus duration. Left: decoding for classifiers trained on task relevant and tested on task irrelevant stimuli (purple). Right: decoding for classifiers trained on task irrelevant and tested on task relevant stimuli (orange-red). ROIs showing significantly above-chance (50%) decoding are displayed on inflated surface maps from a left lateral view (top left), posterior view (top right) and left medial view (bottom). **c.** MEG cross-task decoding of category for letter vs false font. (orange-red: train on test irrelevant, test on task relevant; purple: train on task relevant, test on task irrelevant). Left: results in posterior ROIs. Right: results in prefrontal ROIs. **d.** iEEG cross-task temporal generalization of category decoding (letters vs. false fonts) classifiers trained on task relevant stimuli and tested on task irrelevant stimuli. The three stimulus durations are plotted in columns (left: 0.5 s; center: 1.0 s; right: 1.5 s) and the two theory ROIs in rows (top: posterior ROIs; bottom: prefrontal ROIs). Significantly above-chance (50%) decoding is indicated by the outlined pink-red regions in the temporal generalization matrices. **e.** iEEG cross-task temporal generalization of category decoding (faces vs. objects) in the opposite direction as in Figure 2b (classifiers trained on task relevant stimuli and tested on task irrelevant stimuli). Conventions as in c. **f.** iEEG cross-task temporal generalization of category decoding (faces vs. objects), Classifiers are trained on task relevant and tested on task irrelevant stimuli. Pseudotrials are used to boost decoding accuracy. Conventions as in c. **g.** iEEG decoding accuracies within the theory-relevant ROIs using pseudotrial aggregation to boost decoding accuracies, collapsed across stimulus duration. Conventions as in b.

**Extended Data Figure 3:**
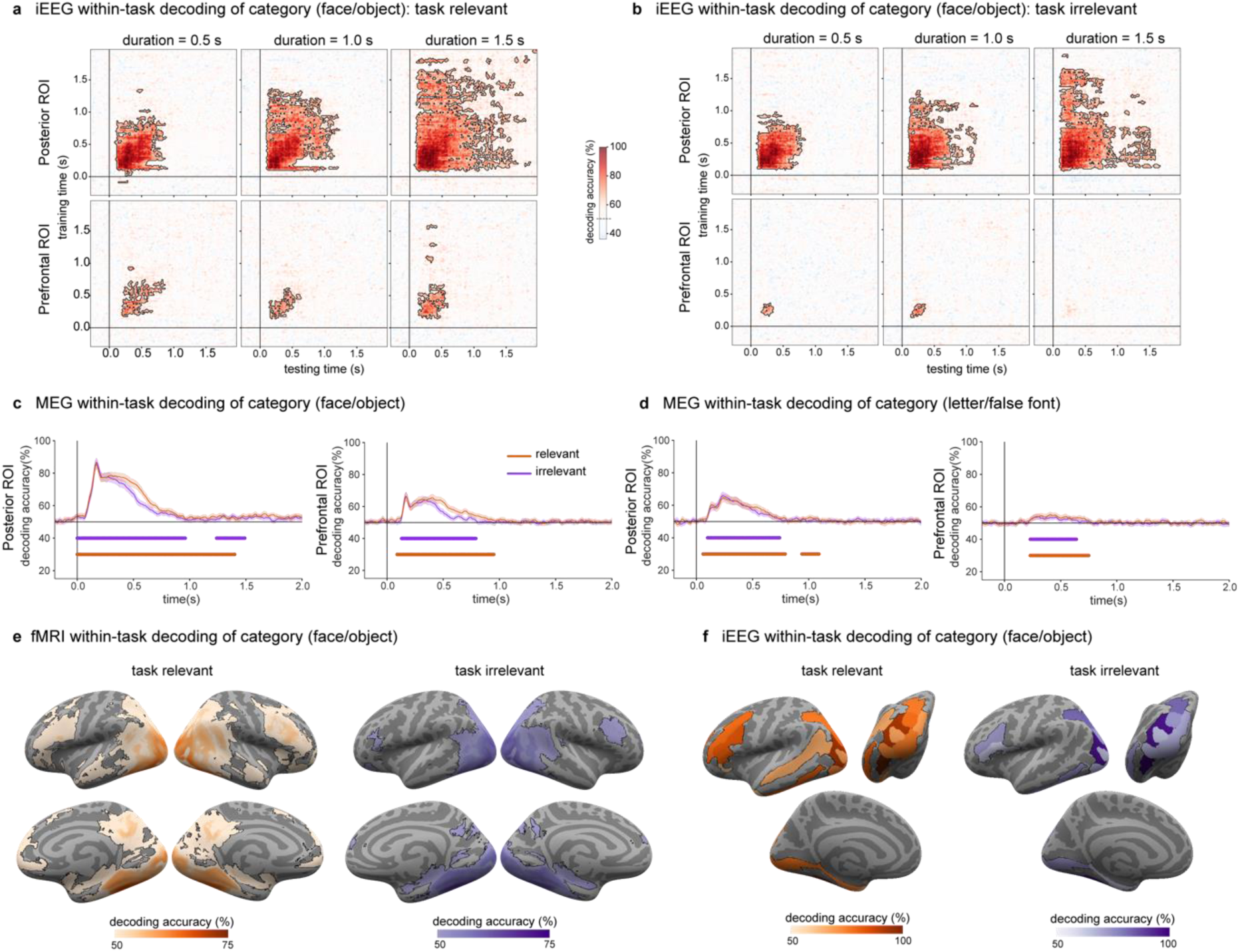
Within-task temporal generalization of decoding of stimulus category (faces vs. objects). **a.** iEEG decoding accuracies for pattern classifiers trained and tested on task relevant stimuli. As in Figure 2b, the three stimulus durations are plotted in columns (left: 0.5 s; center: 1.0 s; right: 1.5 s) and the two theory ROIs in rows (top: posterior ROIs; bottom: prefrontal ROIs). Significantly above-chance (50%) decoding is indicated by the outlined pink-red regions in the temporal generalization matrices. **b.** iEEG decoding accuracies for pattern classifiers trained and tested on task irrelevant stimuli. Same plotting conventions as in panel a. **c.** MEG within task decoding of category for faces vs objects (red-task relevant; purple-task irrelevant). Left: results in posterior ROIs. Right: results in prefrontal ROIs. **d.** MEG within task decoding of category for letters vs false fonts (red-task relevant; purple-task irrelevant). Left: results in posterior ROIs. Right: results in prefrontal ROIs. **e.** fMRI decoding using a searchlight approach, collapsed across the three stimulus durations. Left: decoding accuracies for pattern classifiers trained and tested on task relevant stimuli (orange-red). Right: decoding accuracies for pattern classifiers trained and tested on task irrelevant stimuli (purple). Regions showing significantly above-chance (50%) decoding accuracies are indicated by the outlined colored regions on the inflated cortical surfaces (top: left/right lateral views; bottom: right/left medial views). **f.** iEEG decoding accuracies within the theory-relevant ROIs, collapsed across stimulus duration. Left: decoding for classifiers trained and tested on task relevant stimuli (orange-red). Right: decoding for classifiers trained and tested on task irrelevant stimuli (purple). ROIs showing significant above-chance (50%) decoding are displayed on inflated surface maps from a left lateral view (top left), posterior view (top right) and left medial view (bottom).

**Extended Data Figure 5:**
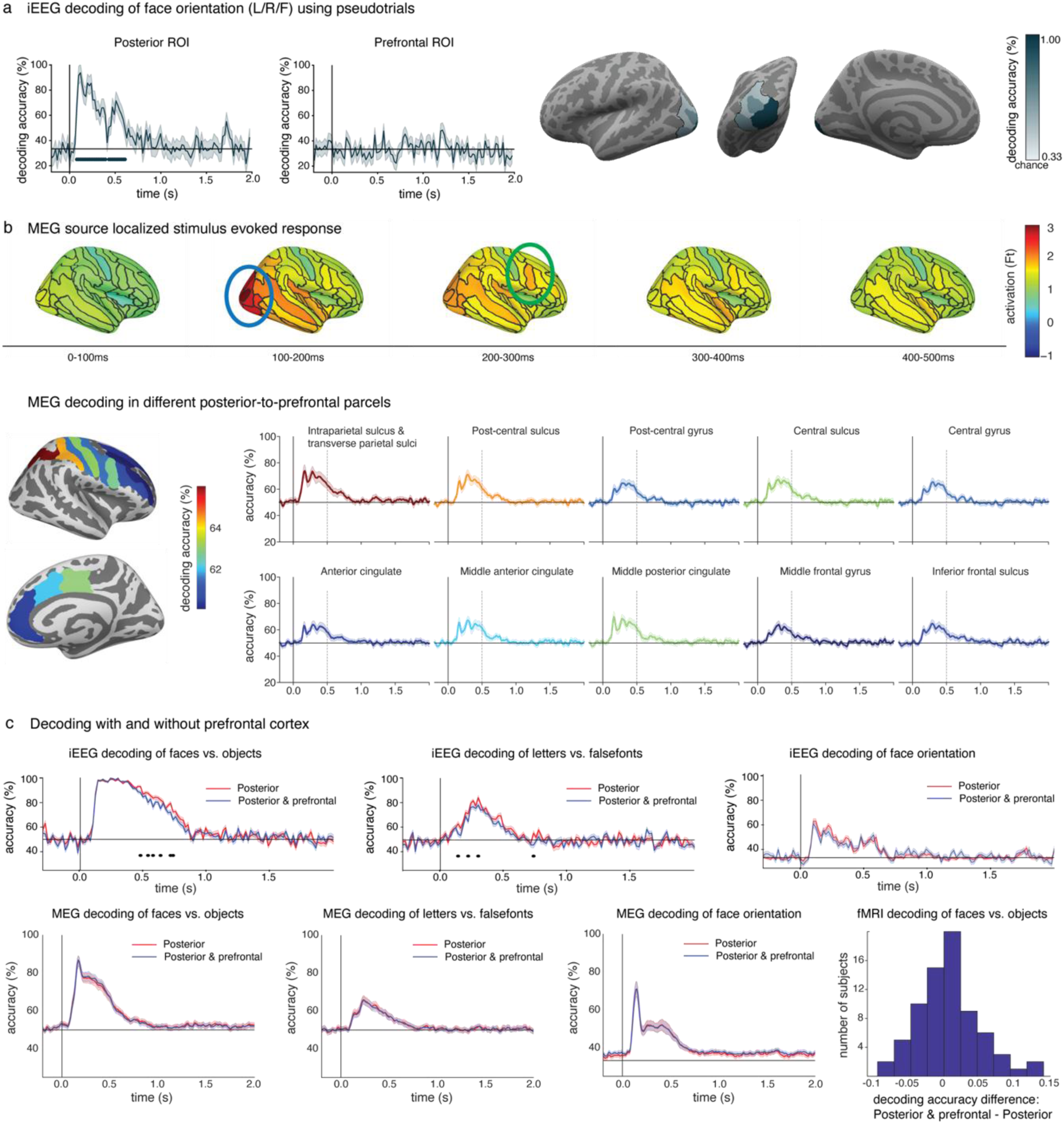
Control analyses for the decoding prediction. a. Left panel: iEEG decoding results of orientation (left vs. right vs. front view faces) within the theory ROIs over time as in Figure 2, using pseudotrials akin to the MEG analysis. Right panel: Regions with electrodes showing above-chance (33%) accuracies are indicated in outlined blue on the inflated surfaces (left: left lateral view; middle: posterior view; right: left medial view). **b.** Two analyses were performed to evaluate potential leakage in the MEG decoding results. These analyses were conducted on independent data from the optimization phase (N=32). Top panel: Stimulus-evoked response in face task relevant trials combined across three stimulus durations were investigated at different latencies and projected on the inflated surfaces. Blue and green ellipses denote posterior and prefrontal areas, respectively. Activity in posterior areas showed the highest peak ∼0.1-0.2 s while prefrontal areas showed the highest peak in a later time window ∼0.2-0.3 s. These differential peak timings serve as evidence against the leakage interpretation. Bottom panels: Face vs. object decoding performance in task relevant trials combining trials across the three durations was investigated separately within parcels in parietal and PFC to evaluate the possibility of a posterior to anterior decoding gradient. Left panel: average face vs. object decoding accuracy in an early time window (0.25-0.5 s) projected on two differently inflated surfaces to better depict gyri and sulci in parietal and prefrontal areas. Right panel, time-resolved decoding performance in parietal and frontal parcels. Decoding performance is highest in posterior areas and lowest in anterior areas, with fairly similar time courses, consistent with the possibility of leakage in decoding from posterior to anterior areas. This effect is better appreciated when considering the high decoding of faces vs. objects in motor related areas, with a gradient from postcentral to precentral sulcus. **c.** Decoding analysis including or excluding prefrontal areas alongside posterior areas to evaluate changes in decoding performance. IIT predicts that including PFC to posterior areas should have either no effect or decreased decoding performance (Posterior + Prefrontal: blue; posterior only: red). Top: iEEG decoding of faces vs. objects (left), letters vs. false fonts (middle) and face orientation (right). Lines underneath the decoding functions indicate time-periods showing significantly worse decoding accuracies when including PFC. Bottom: MEG decoding results, same order as iEEG. Right, fMRI decoding of faces vs. objects. Histogram shows the differences in classification including and excluding frontal areas. iEEG and MEG results consistently show similar (or worse) decoding performance when including prefrontal areas. fMRI accuracies of PFC + Posterior show slight increase of 1.2% on average compared to posterior accuracies, observed in 56% of the subjects. However, it is important to note that these increases are not considered robust due to several factors, including the small magnitude of the accuracy difference and the fact that this slight increase was observed only in the combined features analysis and not the combined models’ analysis (see Methods). The negative outcomes observed in iEEG and MEG data support our interpretation of the fMRI results.

**Extended Data Figure 7:**
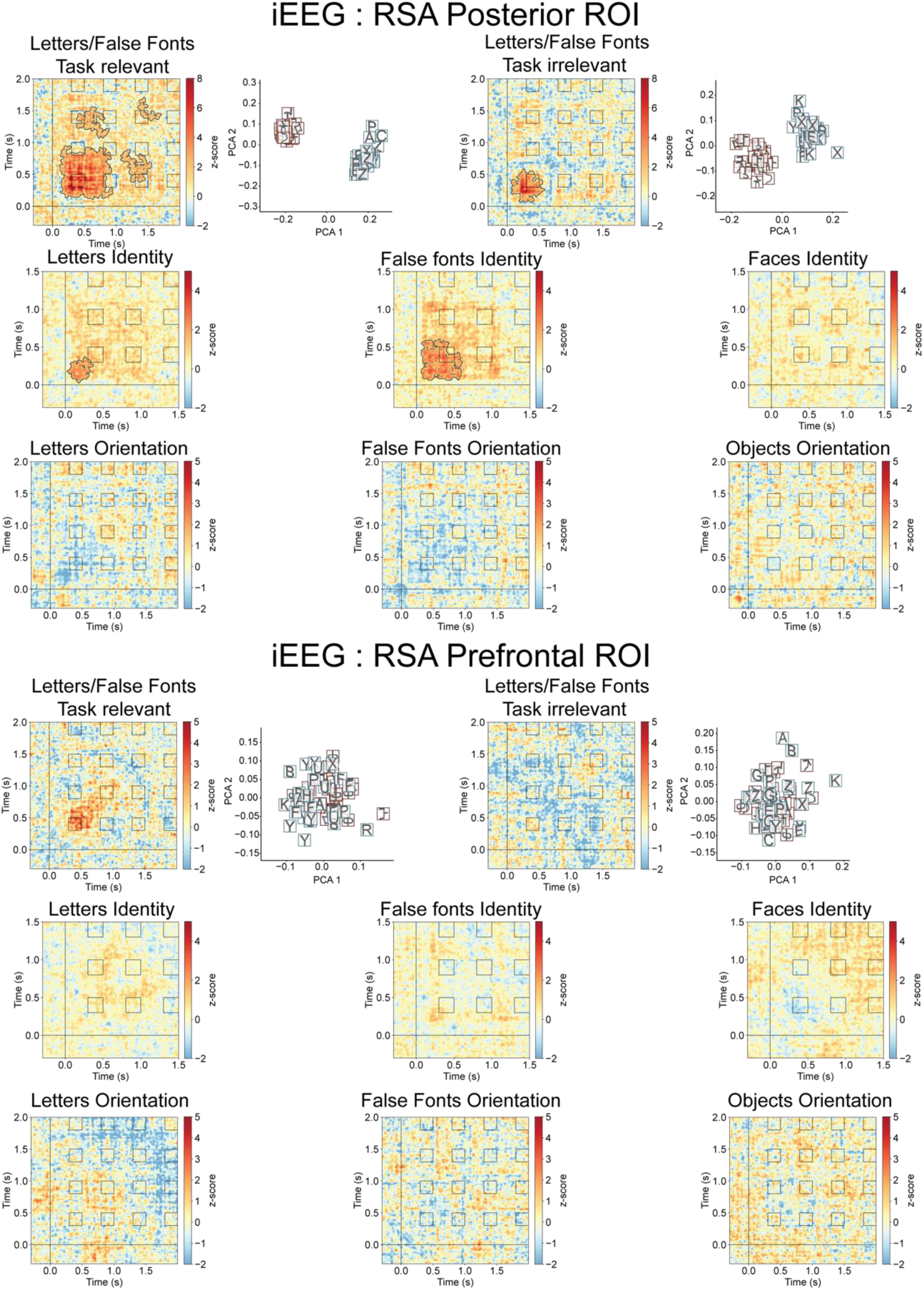
Maintenance of conscious content over time for stimulus categories, identity and orientation. Cross temporal representational similarity matrices across all electrodes in posterior cortex for letters vs. false fonts (upper row), identity (middle) and orientation (bottom) for posterior (upper half) and PFC (lower half) ROI, respectively. Contours in the matrices represent statistical significance, established using cluster-based permutation tests (upper tail test at alpha=0.05). Clear separability between letters and false fonts in posterior cortex is illustrated using Principal Component Analysis at 0.3 s irrespective of the task (left – task relevant, right - task irrelevant). Separability was mostly sustained in the task relevant condition, but not from ∼0.95 to 1.4 s. In the task irrelevant condition, however, separability was statistically significant for a brief period in the beginning. Identity information was statistically significant for letters and false fonts, but not faces. Identity information was not sustained for the entire stimulus duration (however, z-scores were elevated until 1 s, hinting at a limitation in statistical power). No statistically significant orientation information was evident for any of the categories. None of the contrasts yielded statistically significant results in the PFC ROI.

**Extended Data Figure 8:**
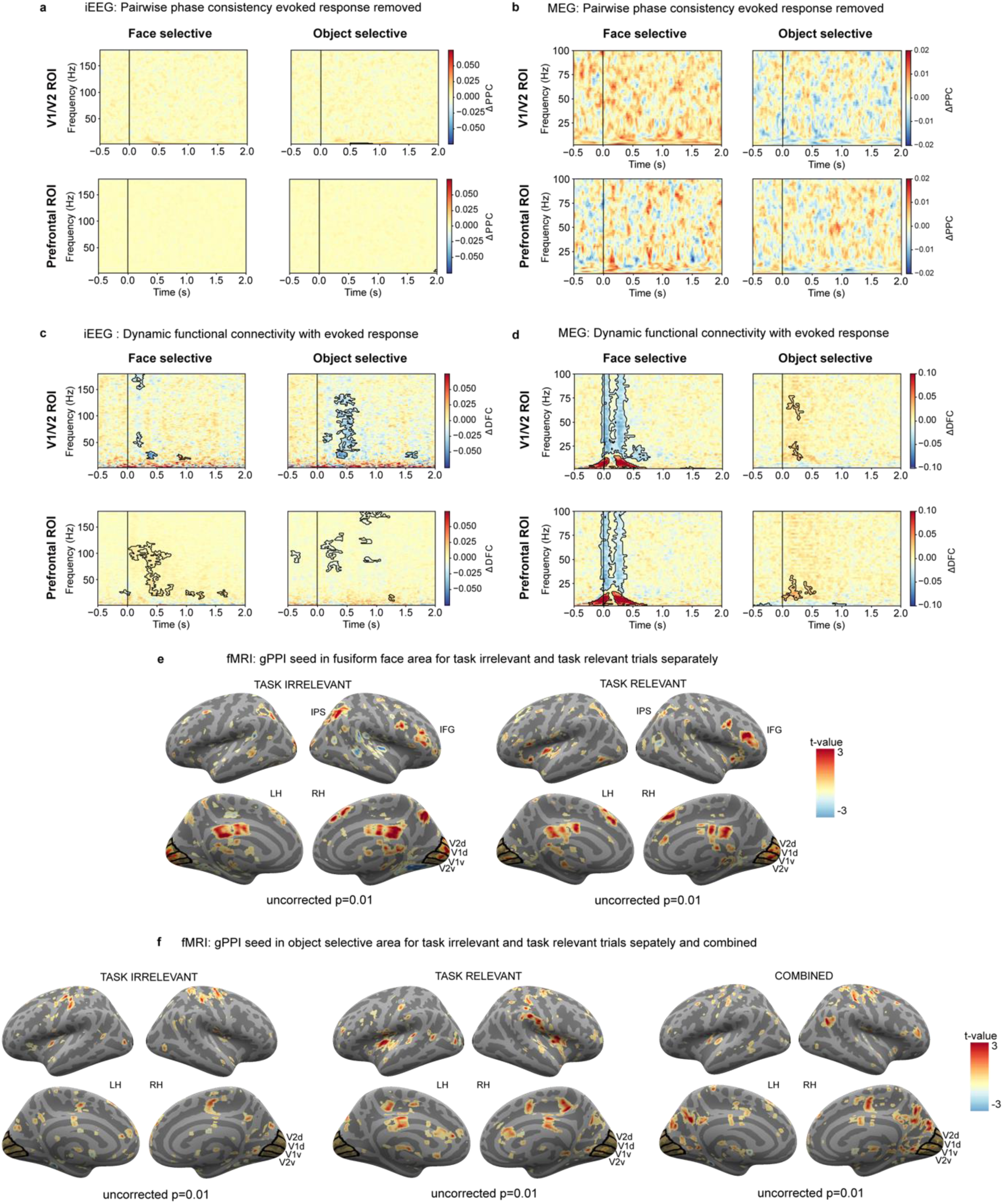
Control analysis for the interareal communication prediction. **a.** iEEG Pairwise phase consistency (PPC) analysis of task irrelevant trials did not reveal any significant category-selective synchrony cluster neither in the posterior ROI nor in the PFC ROI after removing the evoked response. Colorbars represent the change in PPC (face-object trials) for each node (face-selective, object-selective). Positive values reflect stronger connectivity for faces. Negative values reflect stronger connectivity for objects. **b.** MEG PPC analysis of task irrelevant trials did not reveal any significant category-selective synchrony cluster neither in the posterior ROI nor in the PFC ROI after removing the evoked response. The same conventions of Figure 8a are used here. **c.** iEEG Dynamic functional connectivity (DFC) analysis of task irrelevant trials without removing the evoked response reveals significant content-selective connectivity between object-selective electrodes and V1/V2 electrodes (top-right), reflected as broadband (25-125 Hz) decrease in the change in DFC (e.g., faces < objects). Similar broadband content-selective changes in DFC (faces > objects) were observed for face-selective electrodes in PFC (bottom-left). Smaller, yet significant effects, were detected for connectivity between face-selective electrodes and V1/V2 electrodes (top-left) and for object-selective electrodes and PFC electrodes (bottom-right). Conventions as in Figure 8a. **d.** MEG DFC analysis of task irrelevant trials without removing the evoked response reveal significant content-selective synchrony between the face-selective GED filter node and both V1/V2 (top-left) and PFC (bottom-left). This is reflected in an increase in low-frequency connectivity (<25 Hz) combined with a decrease in high-frequency connectivity (25-100 Hz). Smaller yet significant effects were detected for the object-selective GED filter (right). Conventions as in Figure 8a. **e.** Generalized psychophysiological interactions (gPPI) task-related connectivity analysis of task irrelevant (left) and task relevant (right) conditions revealed weak clusters of content-selective connectivity when FFA is used as the analysis seed (p < 0.01, uncorrected). Common significant regions showing task related connectivity in task irrelevant, task relevant, and combined conditions (Figure 4) include V1/V2, right intraparietal sulcus (IPS), and right inferior frontal gyrus (IFG). **f.** gPPI task-related connectivity analysis of task irrelevant (left), task relevant (middle), and combined conditions revealed weak clusters of content-selective connectivity when lateral occipital complex (LOC) is used as the analysis seed (p < 0.01, uncorrected). The results of the gPPI showed that there are no common significant regions showing task related connectivity in task irrelevant, task relevant, and combined conditions.

**Extended Data Figure 9:**
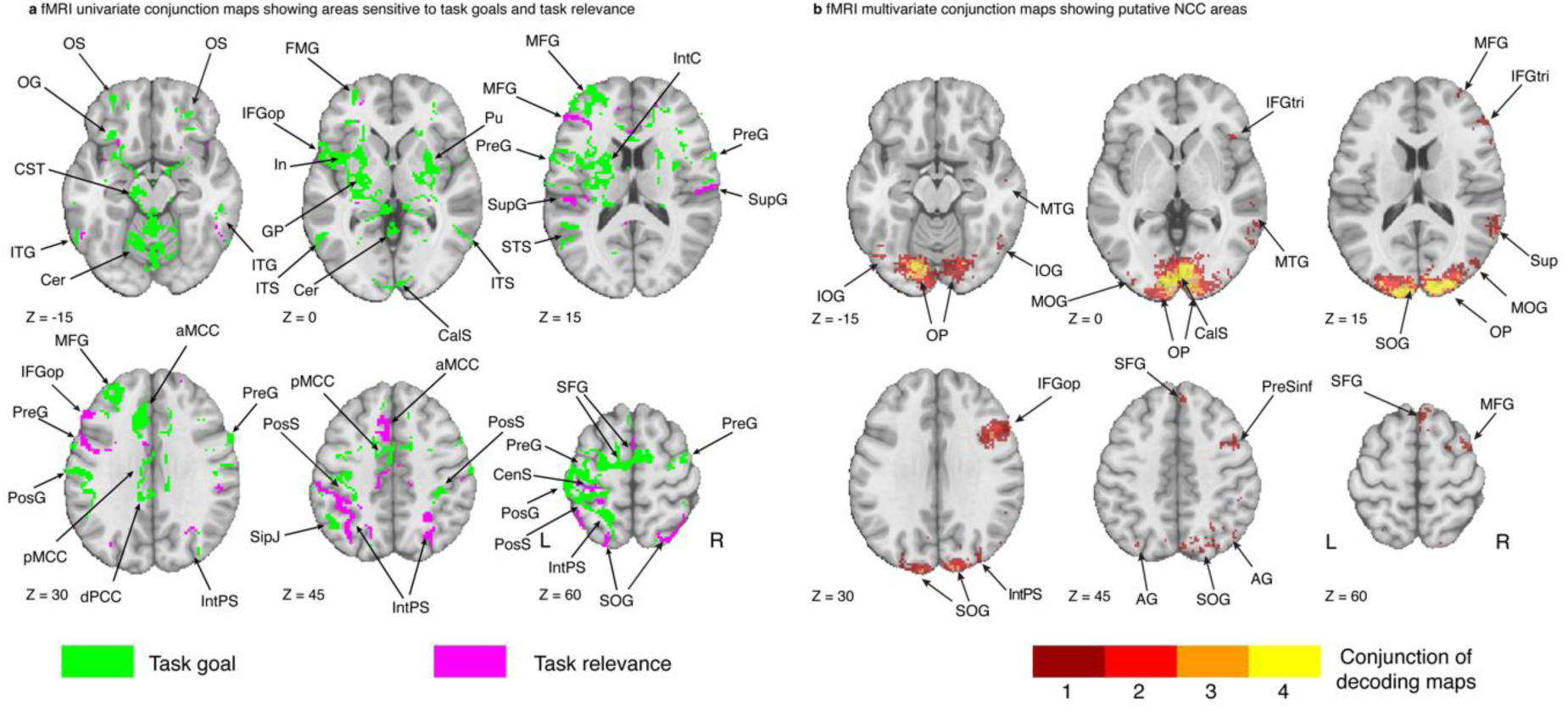
fMRI maps of areas involved in Task goals, Task relevance and presumptive NCC. **a.** Results from the univariate fMRI (N=73) contrast-conjunction analysis identifying task goals (green) and task relevance (magenta) areas. Task goals areas were identified as follows: targets > bsl & task relevant = bsl & task irrelevant = bsl. Task relevance were identified as follows: targets > bsl & task relevant ≠ bsl & task irrelevant = bsl. Axial brain slices are displayed from inferior (top left) to superior (bottom right). Left and right hemisphere are displayed to the left and right, respectively. Neuroanatomical labels from the Destrieux atlas and additional subcortical regions as in Figure 5. **b.** fMRI multivariate contrast-conjunction analysis (N=73) identifying areas showing consistent whole-brain searchlight decoding of stimulus vs. baseline using thresholded statistical maps obtained at the subject level after removing areas identified in a. Conjunction was defined as above chance decoding both for task relevant & task irrelevant stimuli for each stimulus category separately. Colorbar shows the number of stimulus categories passing the conjunction.

**Extended Data Figure 10:**
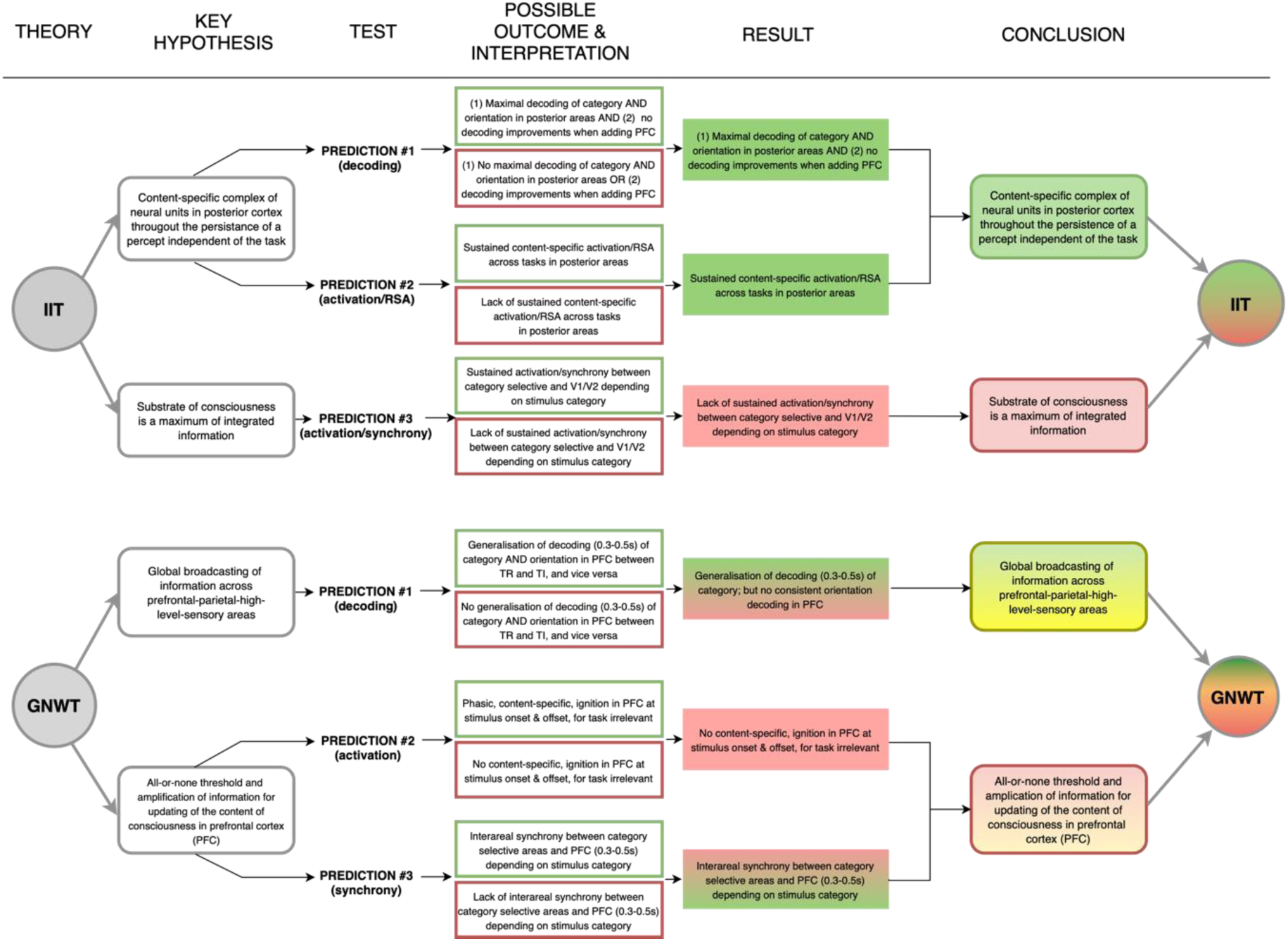
An overview of theoretical predictions, experimental outcomes and interpretations. On the left, the original predictions made by the IIT (top) and GNWT (bottom), preregistered here (see also ^12^; Figure 1). The table describes the key hypotheses (second column, ‘Key hypotheses’) made by the theories (see also Figure 1a), and probed in three different test analyses (third column, ‘Test’; decoding (prediction #1; Figure 2), activation & RSA (prediction #2; Figure 3) and synchrony (prediction #3; Figure 4). Next, we describe the possible outcomes of each of these analyses, and how they would inform the theoretical predictions (fourth column, ‘Possible outcome and interpretation’). Outcomes that conform with the prediction are presented in a green frame (i.e., ‘pass’), while outcomes that contradict the prediction are presented in a red frame (i.e., ‘fail’). Thus, the left side of the table presents the a-priori predictions and expected outcomes, prior to conducting the experiment. The right side of the figure presents instead the actual findings of this experiment, integrating over the three modalities and multiple tests. We first describe the key findings with respect to each prediction (fifth column; ‘Result’). Green marks results that confirm the theories, red marks results that challenge them, the mixture of green/red marks cases in which the combination of results yielded a mixture of a pass and a fail. Finally, we integrate over these results to generate the final conclusion based on the key hypotheses: Here, green/red is used for pass/fail. Yellow marks cases in which we considered that the results did not allow a confident interpretation. Namely, for GNWT’s prediction #1, we found cross-task generalization of decoding of faces vs. objects, in line with the prediction. However, the only evidence for orientation decoding was found in the MEG data, where we could not conclusively rule out leakage from posterior areas. Thus, as passing this prediction requires both decoding of category and orientation to be found, we cannot determine with high confidence if this prediction should be counted as a pass or a fail. For GNWT’s prediction #3, we do find evidence supporting it, yet with an exploratory analysis (DFC), after the preregistered analysis failed to show support for the prediction. Thus, it cannot be regarded as a preregistered ‘pass’. Altogether: for IIT, a mixture of a passed prediction (of content-specific complex of neural units in posterior cortex, throughout the persistence of a percept, independent of the task) and a challenge prediction (of maximum integrated information), and for GNWT, two mixtures of partly challenged predictions (of global broadcasting of information in the PFC and an all-or-none threshold and amplification of information updating the content of consciousness in PFC).

**Extended Data Table 2:** Theory defined anatomical ROIs (Destrieux atlas)

| Parcellation | Relevant for |
| --- | --- |
| Fronto-marginal gyrus (of Wernicke) and sulcus | IIT excluded |
| Inferior occipital gyrus (O3) and sulcus | IIT, FFA |
| Transverse frontopolar gyri and sulci | IIT excluded |
| Anterior part of the cingulate gyrus and sulcus (ACC) | GNWT, IIT excluded |
| Middle-anterior part of the cingulate gyrus and sulcus (aMCC) | GNWT, IIT excluded |
| Middle-posterior part of the cingulate gyrus and sulcus (pMCC) | GNWT |
| Cuneus (O6) | IIT |
| Opercular part of the inferior frontal gyrus | GNWT, IIT excluded |
| Orbital part of the inferior frontal gyrus | GNWT, IIT excluded |
| Triangular part of the inferior frontal gyrus | GNWT, IIT excluded |
| Middle frontal gyrus (F2) | GNWT, IIT excluded |
| Superior frontal gyrus (F1) | IIT excluded |
| Middle occipital gyrus (O2, lateral occipital gyrus) | IIT, LOC |
| Superior occipital gyrus (O1) | IIT |
| Lateral occipito-temporal gyrus (fusiform gyrus, O4-T4) | IIT, FFA |
| Lingual gyrus, lingual part of the medial occipito-temporal gyrus, (O5) | IIT |
| Parahippocampal gyrus, parahippocampal part of the medial occipito-temporal gyrus, (T5) | IIT |
| Orbital gyri | IIT extended |
| Angular gyrus | IIT extended |
| Supramarginal gyrus | IIT extended |
| Precentral gyrus | IIT extended |
| Straight gyrus, Gyrus rectus | IIT excluded |
| Subcallosal area, subcallosal gyrus | IIT excluded |
| Lateral aspect of the superior temporal gyrus | IIT extended |
| Planum temporale or temporal plane of the superior temporal gyrus | IIT extended |
| Inferior temporal gyrus (T3) | IIT |
| Middle temporal gyrus (T2) | IIT extended |
| Horizontal ramus of the anterior segment of the lateral sulcus (or fissure) | GNWT |
| Vertical ramus of the anterior segment of the lateral sulcus (or fissure) | GNWT |
| Occipital pole | IIT |
| Temporal pole | IIT |
| Calcarine sulcus | IIT |
| Inferior frontal sulcus | IIT extended, GNW |
| Middle frontal sulcus | GNW, IIT excluded |
| Superior frontal sulcus | GNW, IIT excluded |
| Sulcus intermedius primus (of Jensen) | IIT extended |
| Intraparietal sulcus (interparietal sulcus) and transverse parietal sulci | IIT |
| Middle occipital sulcus and lunatus sulcus | IIT, LOC |
| Superior occipital sulcus and transverse occipital sulcus | IIT |
| Anterior occipital sulcus and preoccipital notch (temporo-occipital incisure) | IIT extended |
| Lateral occipito-temporal sulcus | IIT extended |
| Lateral orbital sulcus | IIT excluded |
| Medial orbital sulcus (olfactory sulcus) | IIT excluded |
| Orbital sulci (H-shaped sulci) | IIT excluded |
| Inferior part of the precentral sulcus | IIT extended |
| Suborbital sulcus (sulcus rostrales, supraorbital sulcus) | IIT excluded |
| Inferior temporal sulcus | IIT extended |
| Superior temporal sulcus (parallel sulcus) | IIT |

Anatomical regions-of-interest (ROIs) labelled in Destrieux et al. (2010) atlas used for testing theories predictions (unless otherwise specified). IIT and GNWT ROIs are the ROIs in posterior cortex and prefrontal cortex (PFC) relevant for IIT and GNWT predictions, respectively. IIT extended ROIs specify areas within PFC in which IIT considers that effects might be found, rendering them non-diagnostic for the evaluation of the theory. IIT excluded ROIs are regions within PFC where IIT does not predict any effect (i.e., no increase in decoding accuracy when these regions are added to posterior regions in the analysis). Thus, an effect found on those areas would pose a challenge for IIT. Additionally, ROIs used to define the Face Fusiform Area (FFA), and Lateral Occipital Complex (LOC), relevant for connectivity analyses are included.

**Extended Data Table 4:**
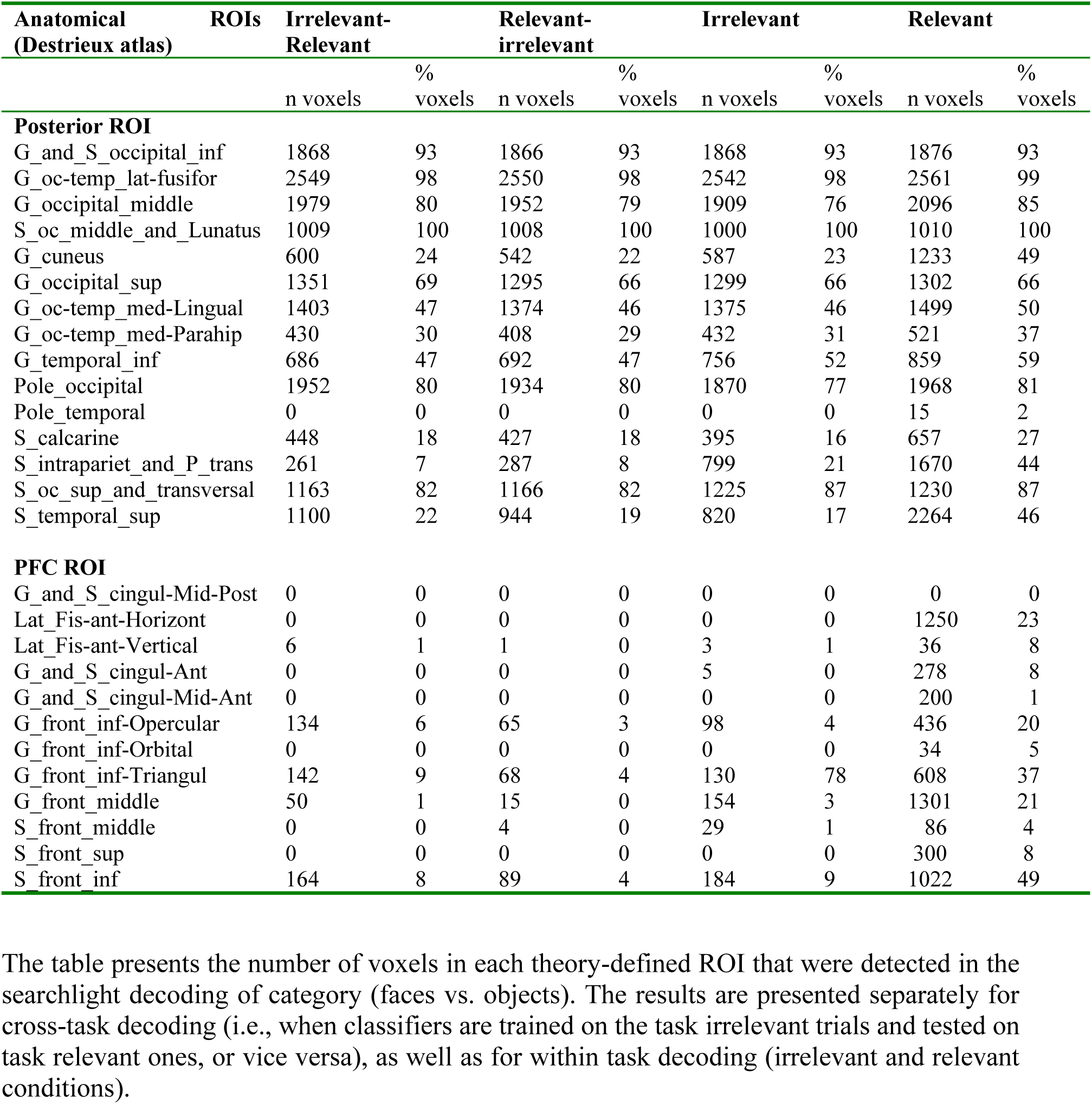
Decoding of faces vs. category in the theory-defined ROIs.

**Extended Data Table 6:** Electrode locations found to be significant in the LMM analysis.

| Channel | x | y | z | Destrieux ROI | Wang ROI | Desikan ROI | Model |
| --- | --- | --- | --- | --- | --- | --- | --- |
| SE107-O2PH16 | -0.03618 | -0.08678 | 0.000733 | S_oc_middle_and_Lunatus | TO1 | ctx-lh-lateraloccipital | IIT x Cate |
| SE120-T3bOT10 | -0.05876 | -0.06964 | -0.02078 | G_oc-temp_lat-fusifor | Unknown | ctx-lh-fusiform | IIT x Cate |
| SE120-T3bOT9 | -0.05712 | -0.0689 | -0.02016 | G_oc-temp_lat-fusifor | Unknown | ctx-lh-fusiform | IIT x Cate |
| SF102-LO1 | -0.01976 | -0.10359 | 0.001174 | Pole_occipital | V2d | ctx-lh-lateraloccipital | IIT x Cate |
| SF102-LO2 | -0.02301 | -0.09792 | 0.005426 | Pole_occipital | V2d | ctx-lh-lateraloccipital | IIT x Cate |
| SF103-PIT1 | -0.04072 | -0.06213 | -0.02039 | G_oc-temp_lat-fusifor | Unknown | ctx-lh-fusiform | IIT x Cate |
| SF103-PIT2 | -0.04156 | -0.04393 | -0.02499 | G_oc-temp_lat-fusifor | Unknown | ctx-lh-fusiform | IIT x Cate |
| SF104-LO1 | -0.01396 | -0.10275 | 0.008659 | Pole_occipital | V2d | ctx-lh-lateraloccipital | IIT x Cate |
| SF104-LO2 | -0.01663 | -0.10338 | 0.005258 | Pole_occipital | V2d | ctx-lh-lateraloccipital | IIT x Cate |
| SF109-IO3 | 0.006178 | -0.07586 | -0.00279 | G_oc-temp_med-Lingual | V2v | ctx-rh-lingual | IIT x Cate |
| SF109-IO4 | 0.005093 | -0.07816 | -0.0047 | G_oc-temp_med-Lingual | V2v | ctx-rh-lingual | IIT x Cate |
| SF113-RIT1 | 0.038119 | -0.04974 | -0.02225 | G_oc-temp_lat-fusifor | Unknown | Cerebellum-Cortex | IIT x Cate |
| SF113-RIT2 | 0.040545 | -0.04845 | -0.02346 | G_oc-temp_lat-fusifor | Unknown | Cerebellum-Cortex | IIT x Cate |
| SE107-O1b3 | -0.01196 | -0.06305 | -0.00094 | G_oc-temp_med-Lingual | Unknown | ctx-lh-lingual | GNW |
| SE107-O2PH14 | -0.03383 | -0.08203 | 6.93E-05 | S_oc_middle_and_Lunatus | LO2 | ctx-lh-lateraloccipital | GNW |
| SE107-O2PH15 | -0.0354 | -0.08519 | 0.000512 | S_oc_middle_and_Lunatus | LO2 | ctx-lh-lateraloccipital | GNW |
| SE108-O2b14 | -0.0294 | -0.09064 | -0.00472 | S_oc_middle_and_Lunatus | LO1 | ctx-lh-lateraloccipital | GNW |
| SE120-O2*5 | -0.04225 | -0.09646 | -0.00451 | G_and_S_occipital_inf | Unknown | ctx-lh-lateraloccipital | GNW |
| SE120-O2*6 | -0.04354 | -0.09769 | -0.00357 | G_and_S_occipital_inf | Unknown | ctx-lh-lateraloccipital | GNW |
| SE120-T3c6 | -0.05264 | -0.08681 | 0.025426 | S_temporal_sup | Unknown | ctx-lh-inferiorparietal | GNW |
| SF104-LO3 | -0.02255 | -0.10253 | 0.000551 | Pole_occipital | V2d | ctx-lh-lateraloccipital | GNW |
| SF109-DL4 | 0.022039 | -0.07051 | 0.008421 | S_calcarine | Unknown | ctx-rh-pericalcarine | GNW |
| SF109-DL5 | 0.02433 | -0.07204 | 0.008081 | S_calcarine | Unknown | ctx-rh-pericalcarine | GNW |
| SF109-G45 | 0.04645 | -0.08224 | -0.00242 | G_occipital_middle | Unknown | ctx-rh-lateraloccipital | GNW |
| SE108-O2b13 | -0.02856 | -0.08853 | -0.00505 | G_and_S_occipital_inf | Unknown | ctx-lh-lateraloccipital | IIT |
| SE110-O2*10 | 0.036288 | -0.1042 | -0.00079 | G_and_S_occipital_inf | Unknown | ctx-rh-lateraloccipital | IIT |
| SE110-O2*7 | 0.031792 | -0.09698 | -0.00721 | S_oc-temp_lat | Unknown | ctx-rh-lateraloccipital | IIT |
| SE110-O2*8 | 0.03359 | -0.09987 | -0.00464 | G_and_S_occipital_inf | Unknown | ctx-rh-lateraloccipital | IIT |
| SE110-O2*9 | 0.035389 | -0.10276 | -0.00207 | G_and_S_occipital_inf | Unknown | ctx-rh-lateraloccipital | IIT |
| SE120-O1b10 | -0.02828 | -0.11893 | 0.004408 | Pole_occipital | V2d | ctx-lh-lateraloccipital | IIT |
| SF102-LO3 | -0.0356 | -0.08904 | -0.00424 | G_occipital_middle | LO2 | ctx-lh-lateraloccipital | IIT |
| SF107-O1 | 0.024693 | -0.10108 | -0.00812 | Pole_occipital | Unknown | ctx-rh-lateraloccipital | IIT |
| SF107-O2 | 0.027381 | -0.09982 | -0.00773 | Pole_occipital | Unknown | ctx-rh-lateraloccipital | IIT |
| SF107-O3 | 0.042207 | -0.08618 | -0.00419 | G_occipital_middle | Unknown | ctx-rh-lateraloccipital | IIT |
| SF113-RO1 | 0.034984 | -0.08617 | 0.010333 | G_occipital_middle | V3B | ctx-rh-lateraloccipital | IIT |
| SF113-RO2 | 0.040244 | -0.08034 | 0.011692 | G_occipital_middle | LO2 | ctx-rh-inferiorparietal | IIT |

Electrodes location in MNI coordinates, as well as in the corresponding parcellations of the Destrieux Atlas, Wang Atlas and Desikan Atlas.

