## Supplementary Information for "An adversarial collaboration to critically evaluate theories of consciousness"

#### Table of Contents

*Behavioral analysis* ... 4

**Pre-registered analyses** ... 4
d' analysis ... 4
Hit rate analysis ... 6
False Alarm analysis ... 6
Reaction times analysis ... 9

*Eye movements analyses* ... 11

**Pre-registered analyses** ... 11
First 0.5 s: fixation distance from center ... 11
First 0.5 s: number of blinks ... 12
First 0.5 s: saccade amplitude ... 13
First 0.5 s: pupil size ... 14
Long trials analysis: fixation distance from center ... 15
Long trials analysis: number of blinks ... 16
Long trials analysis: saccade amplitude ... 18
Long trials analysis: pupil size ... 19

*Control experiment: surprise memory test* ... 22

**Methods** ... 22
Participants ... 22
Apparatus ... 22
Stimuli and procedure ... 22
Analysis ... 23

**Results** ... 24
Exposure phase ... 24
Memory phase ... 24

**Conclusions** ... 26

*Replicability of findings: optimization vs. replication results* ... 27

**Behavioral analysis** ... 27

**Eye movements analysis** ... 27

**Decoding analysis** ... 29
MEG ... 29
fMRI ... 30

**Levels of activation analysis** ... 31
MEG ... 31

**Synchrony analysis** ... 33
MEG ... 33

**Putative NCC analysis** ... 35

### Behavioral analysis

#### Pre-registered analyses

##### d' analysis

A linear mixed model was used to test if d' is modulated by stimulus category (Faces, Objects, Letters, False Fonts), stimulus duration (0.5, 1, 1.5 s) or modality (iEEG, fMRI, MEG). These factors were defined as fixed effects, and subject was defined as a random effect[1]. The dependent variable was an adjusted d' score (using a log-linear correction). A main effect of modality was found ($F(2,167.04)=102.39$, $p<0.001$, Bayes Factor (BF)=$7.80\times10^{26}$), with MEG subjects showing the highest adjusted d' (M=4.02, SD=0.43), followed by the fMRI sample (M=3.48, SD=0.34) which in turn was higher than that of iEEG patients (M=3.15, SD=0.72). The difference in d' between the MEG and fMRI subjects likely stems from the different numbers of trials which affects the d' correction, while the lowest d' in iEEG patients was expected given the clinical setting (Supplementary Figure 1; all post-hoc contrasts $p<0.001$). A main effect of category was also found ($F(3, 238.33)=32.17$, $p<0.001$, BF=$4.34\times10^{14}$), with faces showing a slightly lower d' (M=3.49, SD=0.57) than all other categories (Objects: M=3.68, SD=0.53; Letters: M=3.68, SD=0.58; False Fonts: M=3.64, SD=0.60, all p values <0.001). In addition, d' was lower for false fonts compared to objects (p=0.017). No significant differences were found between the other categories: false fonts vs. letters: p=0.633, letters vs. objects, p=1.000). The slightly lower d' found for faces could potentially reflect the fact that target faces were harder to individualize and remember compared to the stimuli within the other categories. Notably though, this effect seemed to differ by modality, as revealed by an interaction between modality and category ($F(6, 232.01)=7.47$, $p<0.001$, BF=$2.94\times10^{4}$). Follow up analyses showed that the general difference between faces and all other categories stemmed from the MEG sample (all three p values <0.001), while for the iEEG sample it was observed for faces vs. objects ($p<0.001$) and faces vs. letters (p=0.012), but not false fonts (p=0.334), and an additional difference was found in the iEEG sample between false fonts and objects ($p<0.001$). No category differences were found for the fMRI population (p values range between 0.124 and 1.000). Finally, a main effect was also found for stimulus duration ($F(2, 767.32)=11.74$, $p<0.001$, BF=776.74), with the longest duration stimuli evoking a slightly higher d' (M=3.67, SD=0.55) than the shortest (M=3.58, SD=0.59, $p<0.001$) but not the intermediate duration stimuli (M=3.63, SD=0.58, p=0.053). The shortest and intermediate duration stimuli also differed from one another (p=0.019). No additional interactions were found (p values range between 0.898 (BF=0.06) and 1.000 (BF=0.02)).

---

[1] Notably, in our preregistration we mistakenly defined Item (i.e., exemplar stimuli within each category) as another random factor, yet this cannot be performed as d' is calculated across items.

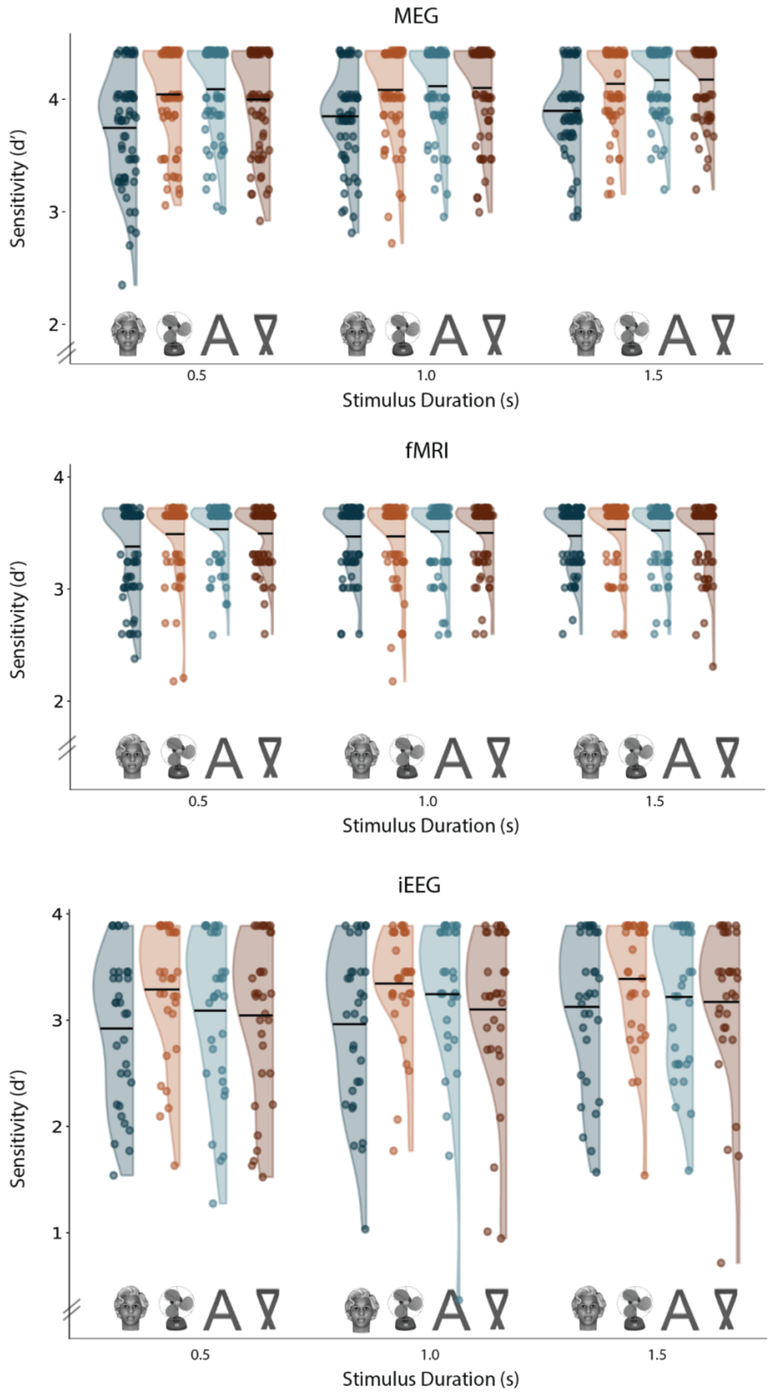

**Supplementary Figure 1**. d' for MEG (upper panel), fMRI (middle panel) and iEEG (lower panel), for each one of the categories (faces, objects, letters, false fonts, marked with exemplary stimuli, and drawn in blue, orange, turquoise and brown, respectively) and each duration (horizontal axis; 0.5, 1.0 and 1.5 s). Each dot is an individual subject, plotted together with the overall distribution. Black horizontal lines depict the mean for each condition.

Importantly, there were no differences in performance, measured using d' between labs within each modality (iEEG: $F(2, 15.15)=0.11$, $p=1.000$ (all p values were Bonferroni corrected), BF=0.05; fMRI: $F(1, 70.87)=0.96$, $p=1.000$, BF=0.05; MEG: $F(1, 62.22)=0.08$, $p=1.000$, BF=0.04).

#### Hit rate analysis

To test whether the modality effect in sensitivity was affected by the different number of trials in each modality, we conducted an exploratory analysis where we ran the same model but with hit rate as a dependent variable, instead of d'. A small difference in hit rates (~3.5%) between different modalities was found ($p<0.001$, BF=77.71), such that hit rates in the iEEG modality (M=93.90, SD=12.29) were found to be lower than both fMRI (M=97.47, SD=7.26, $p<0.001$) and MEG (M=97.54, SD=5.20, $p<0.001$). No difference was found between the fMRI and MEG modalities ($p=1.000$), which further strengthens the conclusion that the higher d' values reported above for MEG stemmed from the higher number of trials (and the log-linear correction); see also False Alarms analysis below, where no difference was found between the MEG and fMRI samples). In addition, hit rates did not differ between different categories ($p=1.000$), and no interactions were found (p values range between 0.330 and 1.000). No difference in hit rates was found between labs within each modality (fMRI: $F(1, 72.81)=0.23$, $p=0.633$, BF=0.04; MEG: $F(1, 54.69)=3.99$, $p=0.356$, BF=0.23; iEEG: $F(2, 8.91)=0.71$, $p=1.000$, BF=0.06). Stimulus duration did modulate hit rates ($p<0.001$, BF=66.98), with the short duration stimuli (M=96.05, SD=8.92) showing slightly lower hit rates than both the intermediate (M=96.91, SD=7.78; $p=0.041$) and the long duration stimuli (M=97.51, SD=6.98; $p<0.001$); the intermediate and long duration stimuli did not differ from one another in terms of hit rates ($p=0.102$).

#### False Alarm analysis

A logistic mixed model was used to test if false alarms were modulated by task relevance (Relevant, Irrelevant), stimulus category (Faces, Objects, Letters, False Fonts), or modality (iEEG, fMRI, MEG), defined as fixed effects. Subject was again defined as a random effect.[2] As expected, task relevance affected false alarms ($\chi^2(1)=241.05$, $p<0.001$, BF=$3.89\times10^{51}$), such that more false alarms were found in the task relevant condition (M=1.73%, SD=3.92) compared with the task irrelevant one (M=0.73%, SD=3.90; $p<0.001$, Supplementary Figure 2). This finding reinforces the effectiveness of our task manipulation.

A main effect of category was also found ($\chi^2(3)=182.23$, $p<0.001$, BF=$3.11\times10^{36}$), such that faces (M=1.78%, SD=4.01) led to higher false alarm rates compared with letters (M=0.97%, SD=3.71, $p<0.001$) and objects (M=0.98%, SD=3.59, $p<0.001$), but not compared to false fonts (M=1.43%, SD=4.63, $p=0.130$), which in turn also evoked more false alarms than letters ($p=0.002$). Modality was also found to affect false alarms ($\chi^2(2)=64.05$, $p<0.001$, BF=$7.33\times10^{11}$), with iEEG (M=4.22%, SD=8.22) patients having higher false alarm rates compared to both MEG (M=0.63%, SD=0.62, $p<0.001$) and fMRI (M=0.59%, SD=0.54, $p<0.001$) subjects, which, in turn, did not differ from each other ($p=0.480$). In addition, the interaction between category and modality was

[2] Here, we first ran the model with Item as a random effect, as preregistered, yet the model failed to converge, probably due to the very low number of false alarms.

significant ($\chi^2(6)=35.61$, $p<0.001$, BF=2.75x$10^3$) such that in the fMRI sample there were no differences between the stimulus categories (p values range between 0.070 and 1.000); In the iEEG sample, false fonts (M=4.95%, SD=9.94) let to more false alarms than both letters (M=3.48%, SD=8.08; p=0.005) and objects (M=3.14%, SD=7.91; p<0.001). In addition, faces led to higher false alarm rates compared to both letters (p=0.018) and objects (p<0.001), but did not differ from false fonts (p=1.000). Objects and letters did not differ (p=1.000). This result strengthens the interpretation that the MEG modality was driving the category face effect, with faces (M=1.18%, SD=0.87) showing a higher false alarm rate compared to letters (M=0.31%, SD=0.53; p<0.001), false fonts (M=0.59%, SD=0.77; p=0.013) and objects (M=0.45%, SD=0.70; p<0.001). Notably, the difference between false fonts and letters in the MEG modality was found to be significant as well (p=0.002; false fonts vs. objects p=1.000, letters vs. objects p=0.088).

The interaction between modality and task relevance was also significant ($\chi^2(2)=29.48$, $p<0.001$, BF=2.28x$10^4$), such that while in all modalities participants made more false alarms to task relevant stimuli, the magnitude of such differences changed between modalities (iEEG relevant M=5.21%, SD=8.12, irrelevant M=2.98%, SD=8.67; MEG relevant M=0.92%, SD=0.65, irrelevant M=0.27%, SD=0.68; fMRI relevant M=0.93%, SD=0.81, irrelevant M=0.16%, SD=0.37; p<0.001 in all modalities). The interaction between stimulus category and relevance was also significant ($\chi^2(3)=56.15$, $p<0.001$, BF=2.32x$10^9$) with no difference in false alarm rate between different categories when task irrelevant (all p-values range between 0.173 and 1.000). When task relevant, faces evoked higher false alarm rates (M=2.65%, SD=5.00) compared to all other stimuli (p<0.001 for all contrasts; false fonts: M=1.94%, SD=5.07, letters: M=1.17%, SD=3.59, objects: M=1.17%, SD=3.51). In addition, false fonts had higher false alarm rates compared to both letters and objects (p<0.001 for both). Letters and objects did not differ from each other (p=1.000). The triple interaction was not significant (p=0.696, BF=0.09).

Akin to the d' analysis, no difference in false alarm rates was found between labs within each modality (fMRI: $\chi^2(1)=1.26$, $p=1.000$, BF=0.08; MEG: $\chi^2(1)=3.20$, $p=0.517$, BF=0.20; iEEG: $\chi^2(2)=1.34$, $p=1.000$, BF=0.07).

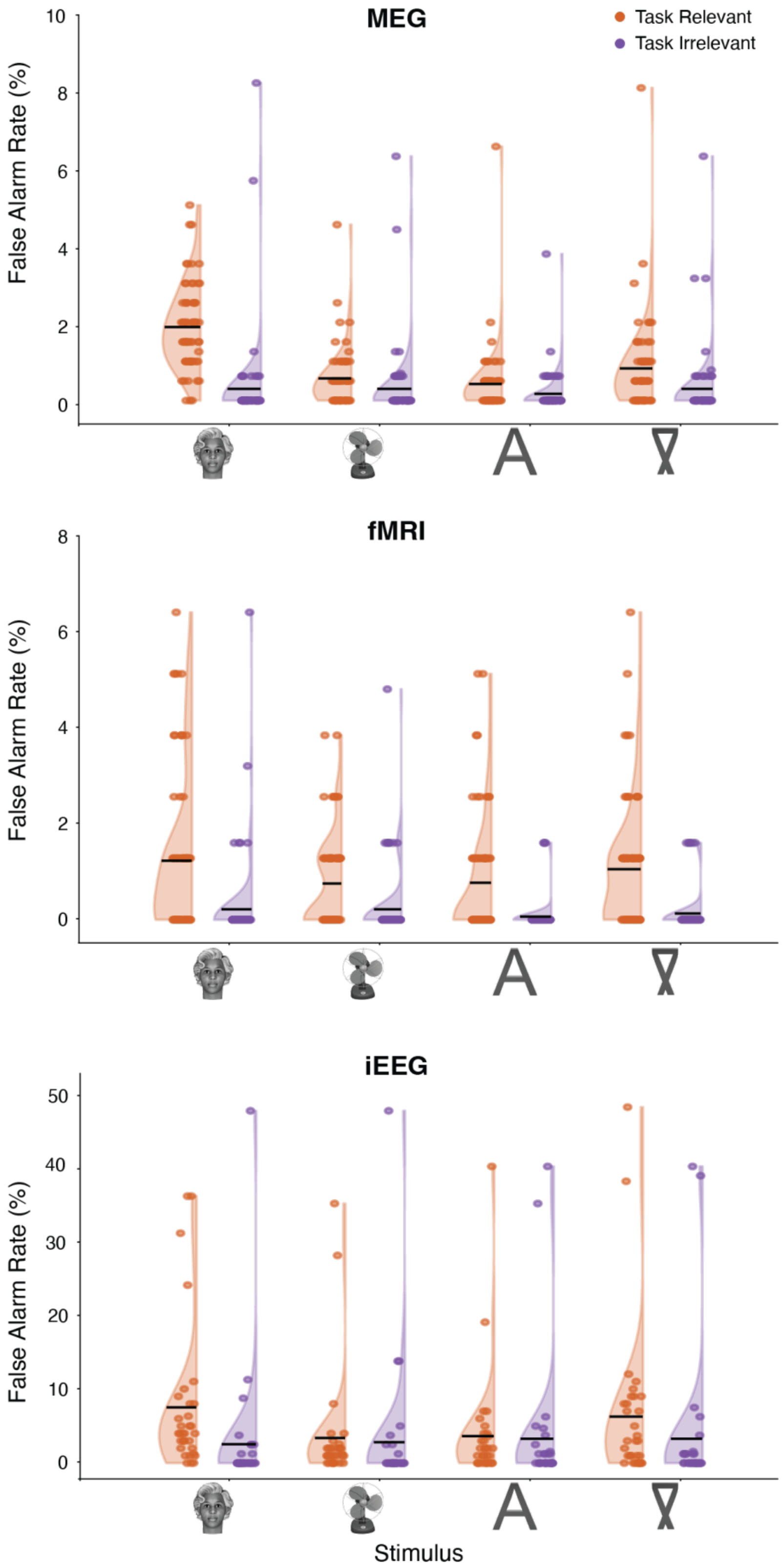
MEG
Task Relevant
Task Irrelevant
False Alarm Rate (%)
10
8
6
4
2
0
fMRI
False Alarm Rate (%)
8
6
4
2
0
iEEG
False Alarm Rate (%)
50
40
30
20
10
0
Stimulus

**Supplementary Figure 2**. FAs for MEG (upper panel), fMRI (middle panel) and iEEG (lower panel), for each one of the categories (faces, objects, letters, false fonts) in the task relevant (orange) and task irrelevant (purple) conditions. Each dot is an individual subject, plotted together with the overall distribution. Note the different scales for the different modalities, with some iEEG patients having a substantially larger number of false alarms compared to the other modalities. Black horizontal lines depict the mean for each condition.

#### Reaction times analysis

A linear mixed model was used to test if reaction times for hits were modulated by stimulus category (Faces, Objects, Letters, False Fonts), stimulus duration (0.5 s, 1.0 s, 1.5 s) or modality (iEEG, fMRI, MEG). These factors were defined as fixed effects, and subject and item were defined as random effects. A main effect of category was found ($F(3, 221.3)=40.23$, $p<0.001$, $BF=2.95 \times 10^{17}$), with letters - which are arguably the easiest and most automated to identify - evoking the fastest responses (M=0.58s, SD=0.20), compared with faces (M=0.63, SD=0.21, $p<0.001$), objects (M=0.62, M=0.19, $p<0.001$) and false fonts (M=0.64, SD=0.21, $p<0.001$; Supplementary Figure 3). In addition, reaction times for objects were faster compared to both faces and false fonts ($p<0.001$ for both). Faces and false fonts did not differ from each other ($p=1.000$).

A main effect of modality was also found ($F(2, 164.9)=14.99$, $p<0.001$, $BF=2.44 \times 10^{3}$), with the MEG sample showing shorter reaction times (M=0.59s, SD=0.06) than the fMRI (M=0.67, SD=0.11; $p<0.001$) and iEEG samples (M=0.65, SD=0.10; $p=0.006$). Reaction times between the iEEG and fMRI samples were not found to be different ($p=0.693$). An interaction between modality and category was also found ($F(6, 14480.8)=3.45$, $p=0.014$, though BF=1.23), with MEG showing a letter advantage (for all contrasts, $p<0.001$; faces: M=0.60, SD=0.08; false fonts: M=0.60, SD=0.06; objects: M=0.59, SD=0.07; letters: M=0.56, SD=0.07), the rest of the contrasts were not significant in the MEG sample (objects vs. false fonts $p=0.961$, faces vs. false fonts=1.000, faces vs. objects $p=1.000$). In the iEEG sample, the letter advantage (M=0.62, SD=0.11) was found when compared to faces (M=0.67, SD=0.12; $p<0.001$) and false fonts (M=0.66, SD=0.12; $p<0.001$), but not to objects (M=0.65, SD=0.10; $p=0.094$). Faces did not differ from false fonts ($p=1.000$) and objects ($p=0.393$), and objects did not differ from false fonts ($p=0.760$). In the fMRI data, faces (M=0.69, SD=0.13) were different from both letters (M=0.64, SD=0.11) and objects (M=0.65, SD=0.11; $p<0.001$ in both cases), and the same was found for false fonts (M=0.70, SD=0.13; again, $p<0.001$). No other effects survived the Bonferroni correction (p values range between 0.302 (BF=0.06) and 1.000 (BF=0.01)).

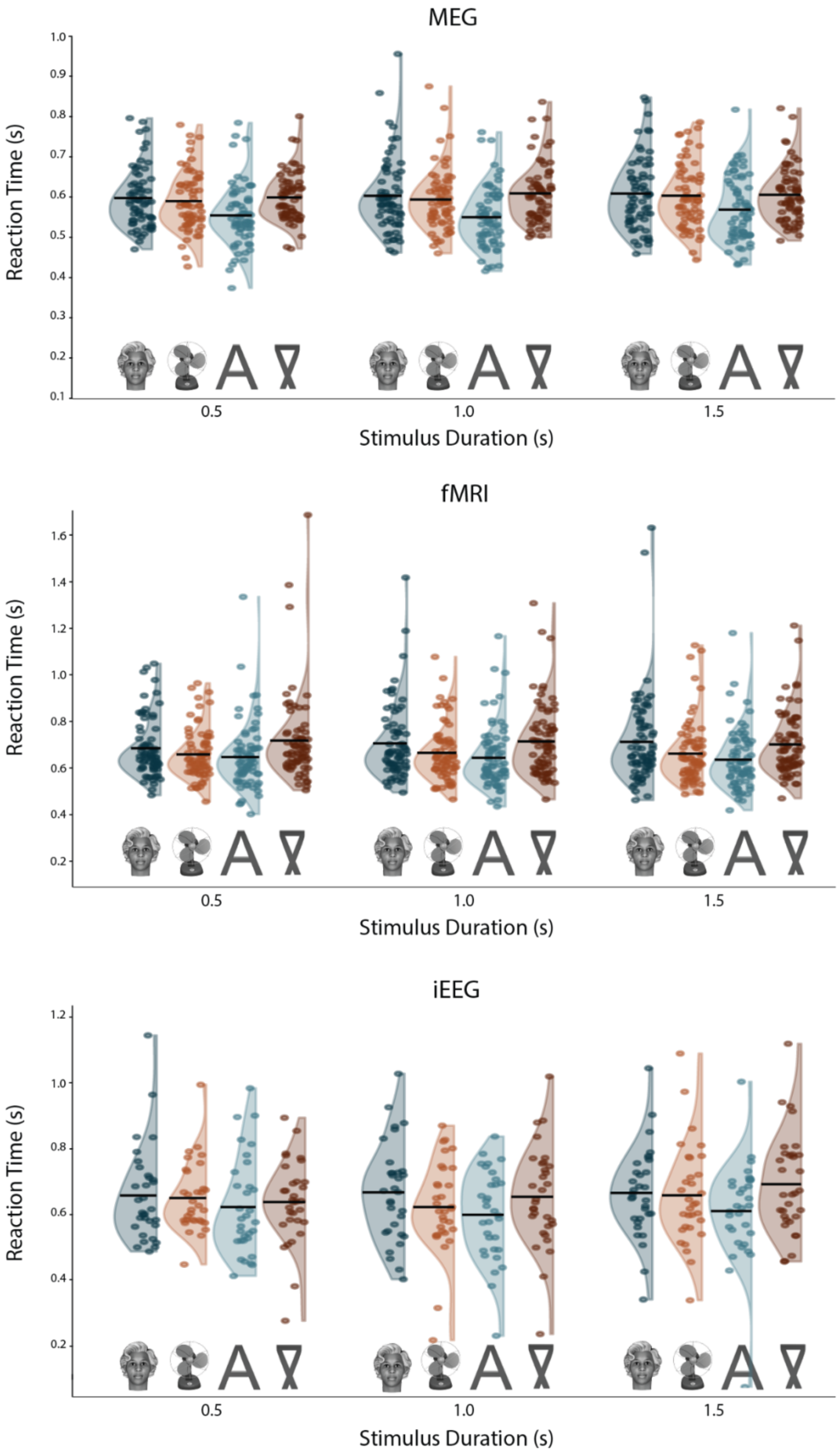

**Supplementary Figure 3**. Reaction times for the different modalities, conditions and durations. The same conventions as in Extended data Figure 1 are used.

### Eye movements analyses

#### Pre-registered analyses

Eye movement patterns were analyzed with respect to four dependent variables: fixation distance from screen center, saccade amplitude, number of blinks and pupil size. For each such variable, we asked how it was modulated by task relevance (Relevant, Irrelevant) and category (Faces, Objects, Letters, False fonts). These were defined as fixed effects, while subject and item were defined as random effects in a linear mixed model. We first focused on the first 0.5 s of the stimulus, that were shared for all three durations. Then, to explore how these variables changed over time, we only analyzed the long duration stimuli (1.5 s), and added time window (0-0.5 s / 0.5-1.0 s / 1.0-1.5 s) as a fixed effect. Thus, 8 models were run overall (4 dependent variables X 2 analyses). This analysis was conducted for all sites using EyeLink; NYU data, which was recorded using Tobii, is only included in the heatmaps (Supplementary Figure 14), but not in any of the other analyses, due to the low quality of the data. Notably, eleven participants were excluded from this analysis: three participants due to not having eye tracking data to begin with (2 iEEG patients, 1 MEG subject) and eight for having data of insufficient quality (5 iEEG patients, 1 MEG subject, and 2 fMRI subjects).

##### First 0.5 s: fixation distance from center

As we report in the main text, overall, subjects were very good at maintaining fixation. Importantly, their ability to maintain fixation within the first 0.5 s, as assessed by the median distance of fixation from the screen center (Supplementary Figure 4), was not affected by task relevance ($F(1, 59012)=0.25$, $p=1.000$, $BF=0.003$). Stimulus category did affect fixations ($F(3, 59021)=4.59$, $p=0.010$, though $BF=0.29$), with lower distance from the center for faces ($M=1.61$, $SD=1.26$) compared to false fonts ($M=1.64$, $SD=1.31$; $p=0.003$; the rest of the comparisons were not significant and ranged between 0.145 and 1.000). The interaction between task relevance and category was also not significant ($F(3, 59010)=0.31$, $p=1.000$, $BF=0.003$).

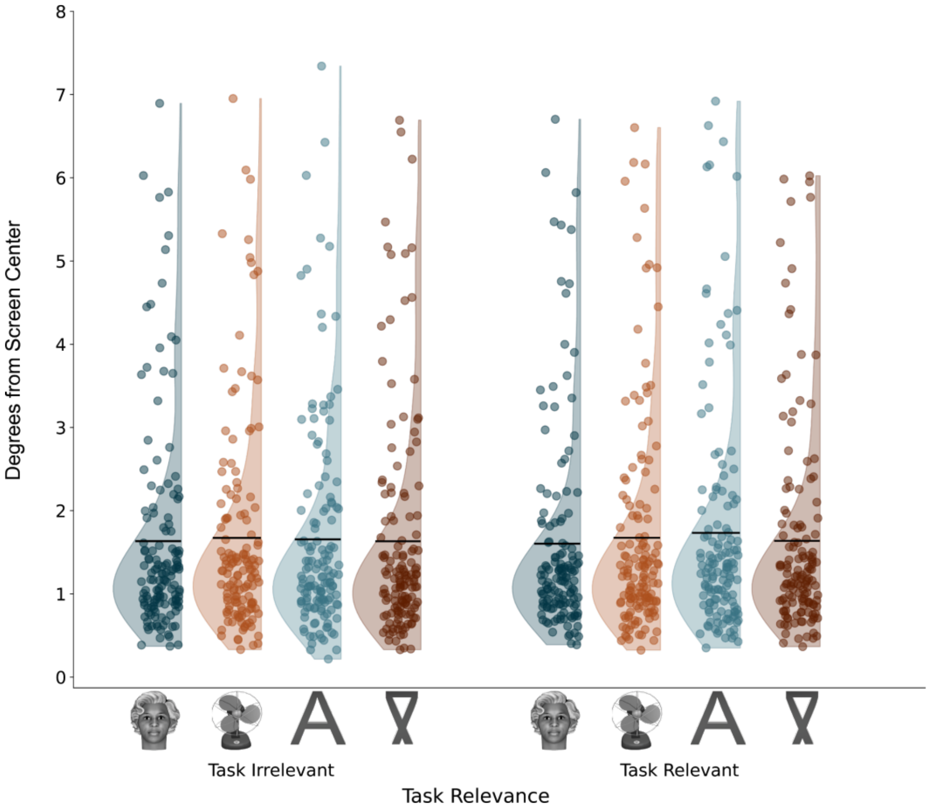

**Supplementary Figure 4**. Mean distance of fixations from the center, across all modalities and durations, per stimulus category and task relevance. The same conventions as in Supplementary Figure 1 are used.

#### First 0.5 s: number of blinks

Overall, the number of blinks was very low, with about one blink per trial on average (M=1.10, SD=1.08; Supplementary Figure 5). Small differences were nevertheless found between the conditions. Namely, a main effect was found for category ($F(3, 188)=31.11$, $p<0.001$, $BF=4.28x10^{12}$), such that faces evoked fewer blinks on average (M=0.14, SD=0.15) than objects (M=0.15, SD=0.17, $p<0.001$), letters (M=0.16, SD=0.16, $p<0.001$) and false fonts M=0.16, SD=0.17, $p<0.001$). The rest of the categories did not differ from one another (p values range between 0.099 and 1.000). A main effect was found for task relevance ($F(1, 120346)=90.75$, $p<0.001$, $BF=5.73x10^{17}$), with less blinks in the task relevant (M=0.14, SD=0.16) than the task irrelevant (M=0.16, SD=0.17, $p<0.001$) condition. An interaction between the two factors was also found ($F(3, 120342)=5.28$, $p=0.004$, though $BF=0.77$). Post-hoc comparisons showed that while for the task relevant condition, faces differed from all conditions ($p<0.001$ for the three comparisons, others were not significant and ranged between 0.477 and 1.000), in the irrelevant condition, faces differed only from letters and false fonts ($p<0.001$ for both), but not from objects ($p=0.155$). In addition, objects also differed from both false fonts ($p<0.001$) and letters ($p=0.023$). False fonts and letters did not differ from each other ($p=1.000$).

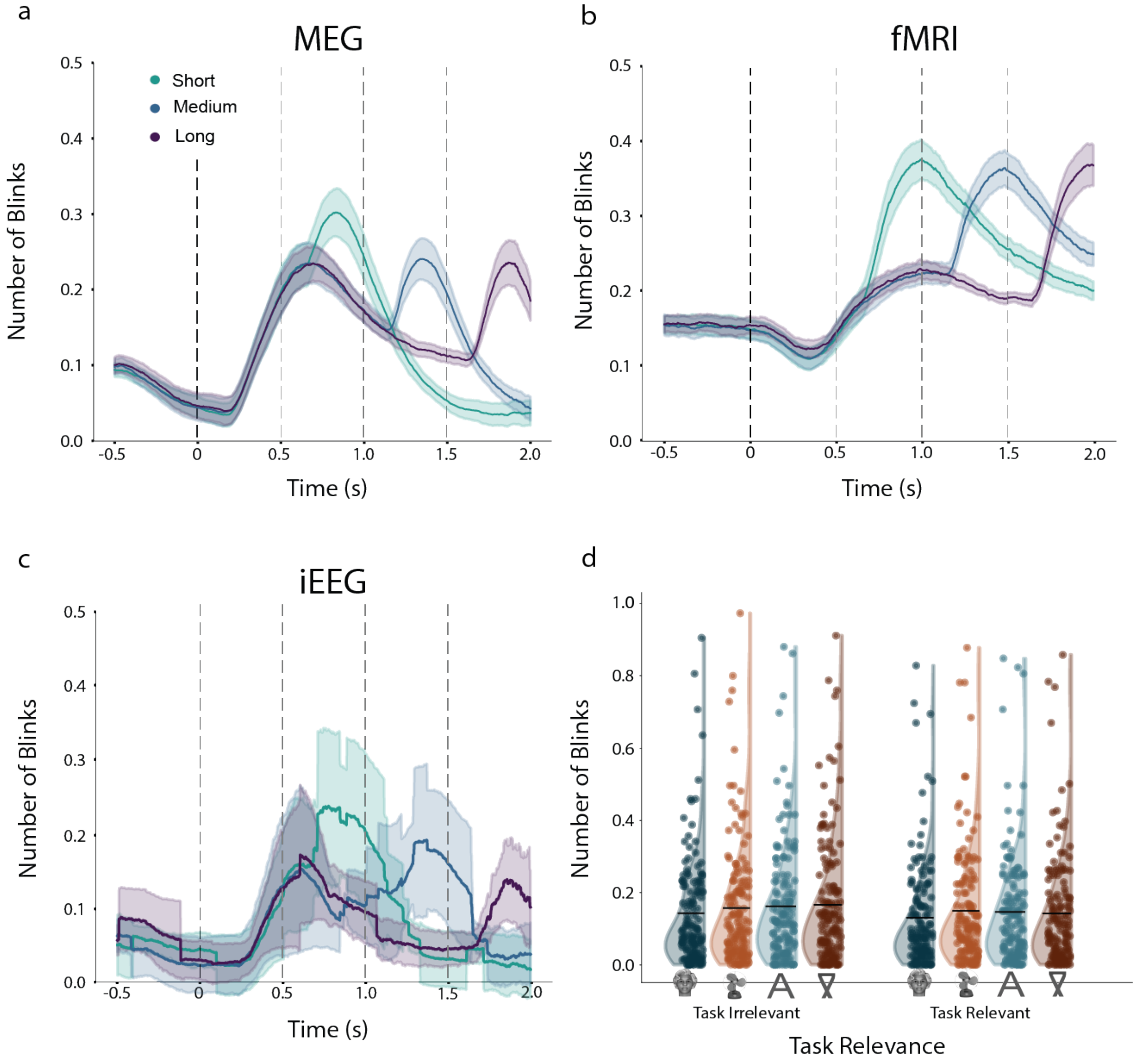

**Supplementary Figure 5**. **a-c**: Averaged number of blinks over time across subjects for the three durations (short, medium and long, in green, blue and purple, respectively), for the three modalities (a: MEG; b: fMRI; c: iEEG). The black dashed line marks the onset of the stimulus, while gray dashed lines mark the three offsets. **d**. Distributions of the averaged number of blinks per subject in the first 0.5 s of all trials (represented by the dots), across all modalities and durations, broken down by stimulus category (faces, objects, letters, false-fonts; horizontal axis) and task conditions (irrelevant on the left, and relevant on the right).

#### First 0.5 s: saccade amplitude

We further investigated the amplitudes of the saccades within the first 0.5 s of stimulus presentation. A main effect of stimulus category was found (F(3, 211)=5.87, p=0.002, though BF=1.31; Supplementary Figure 6d). This stemmed from the letter stimuli evoking greater amplitudes (M=1.40, SD=1.72) compared to faces (M=1.25, SD=1.33, p<0.001), false fonts (M=1.27, SD=1.32, p=0.002) and objects (M=1.24, SD=1.13, p=0.006). Other differences between categories were not found (p=1.000 for all the other comparisons). In addition, no effects were found for either task relevance (F(1, 35980)=0.06, p=1.000, BF=0.003) nor the interaction between relevance and category (F(3, 36025)=1.28, p=0.840, BF=0.005). In addition to saccade amplitudes, we also report the number of saccades over time in the different modalities (Supplementary Figure 6).

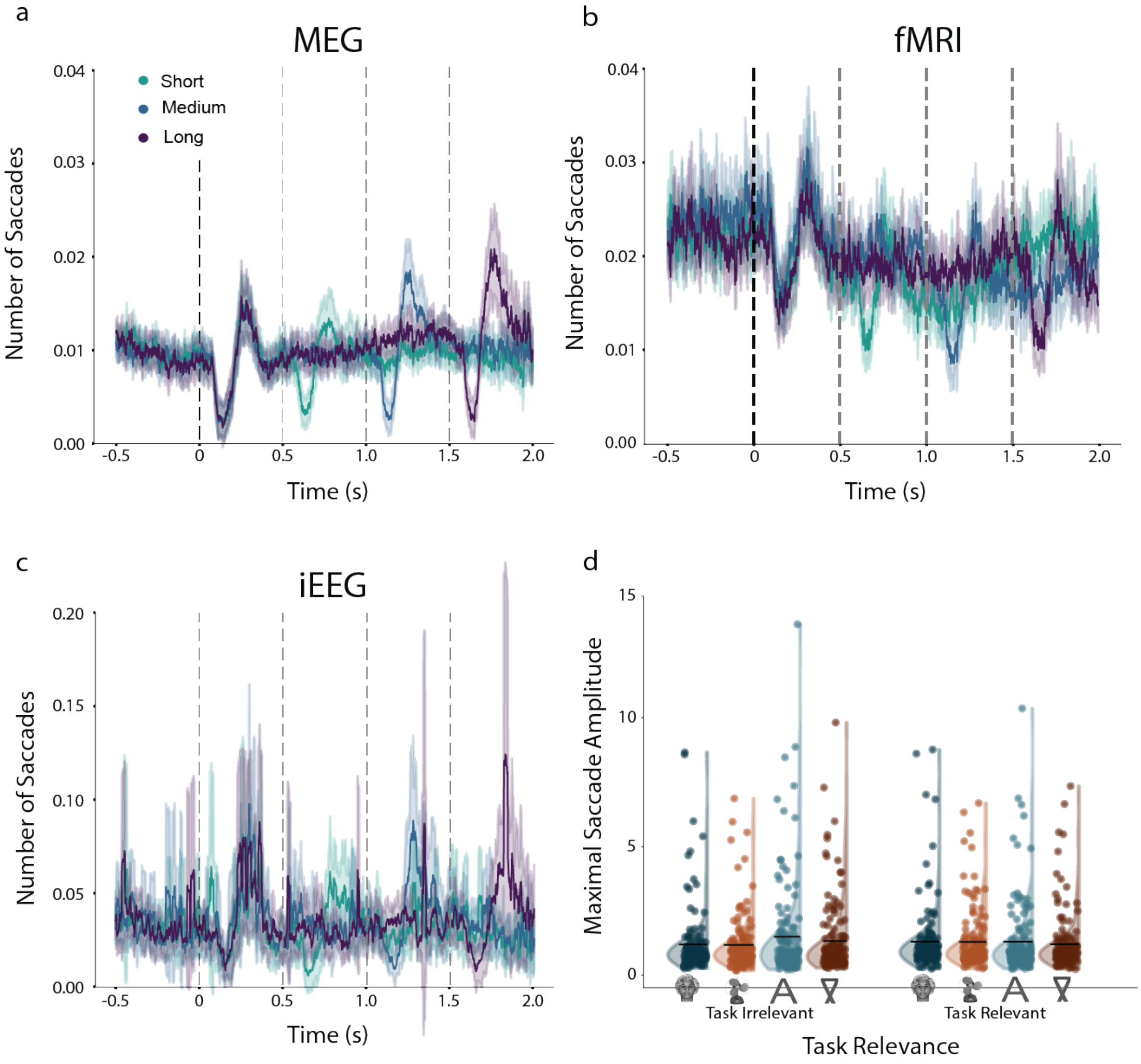

**Supplementary Figure 6**. **a-c.** Number of saccades over the trial, separately for the different modalities. **d.** Distributions of the maximal saccade amplitude per subject in the first 500 ms of all trials. The same conventions as in Supplementary Figure 5 are used.

#### First 0.5 s: pupil size

Pupil size was modulated only by the interaction between stimulus category and task relevance (Supplementary Figure 7; F(3, 112912)=3.83, p=0.028, though BF=0.10); However, post hoc comparisons yielded no differences between the categories both when task relevant and irrelevant (p values range between 0.133 and 1.000 for the task irrelevant stimuli, and between 0.230 and 1.000 for the task relevant stimuli). The other main effects were not significant (p=1.000 and BF=0.003 for both).

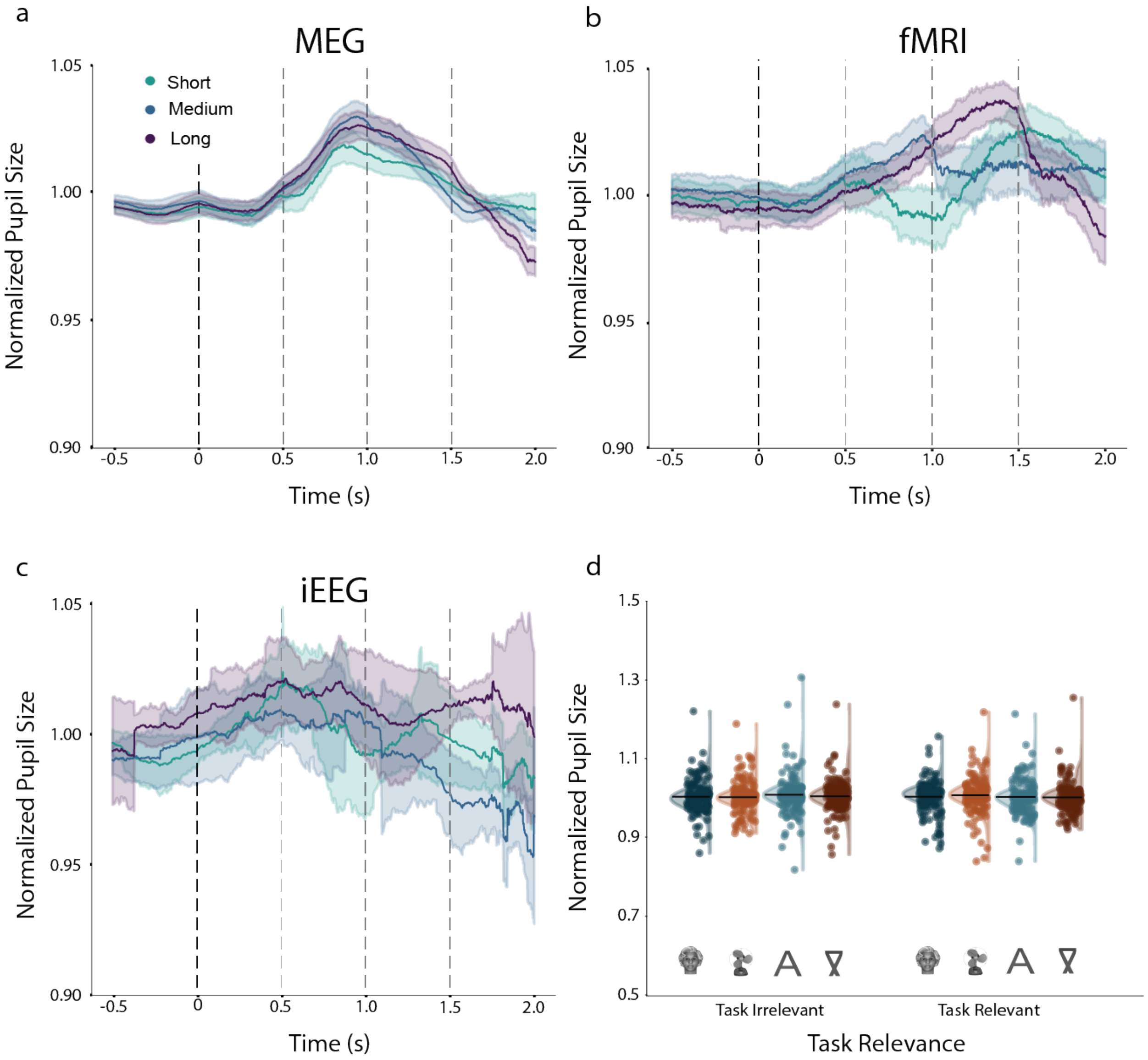

**Supplementary Figure 7**. **a-c.** Averaged standardized pupil size over the trial, separately for the different modalities. **d.** Distributions of the standardized pupil size per subject in the first 500 ms of all trials. The same conventions as in Supplementary Figure 5 are used.

#### Long trials analysis: fixation distance from center

When assessing fixation distance from the center of the screen for the 1.5 s stimuli, a main effect of time window was found ($F(2, 57130)=578.05$, $p<0.001$, $BF=3.27 \times 10^{245}$; Supplementary Figure 8), such that fixation distance differed between all three time windows, growing more distant later in time: first time window: M=1.64 dva, SD=1.36); second time window: M=1.65, SD=1.34; third time window: M=1.71, SD=1.32; p<0.001 for all comparisons). No other effect survived Bonferroni correction (p values range between 0.217 (BF=0.03) and 1.000 (BF=0.006)).

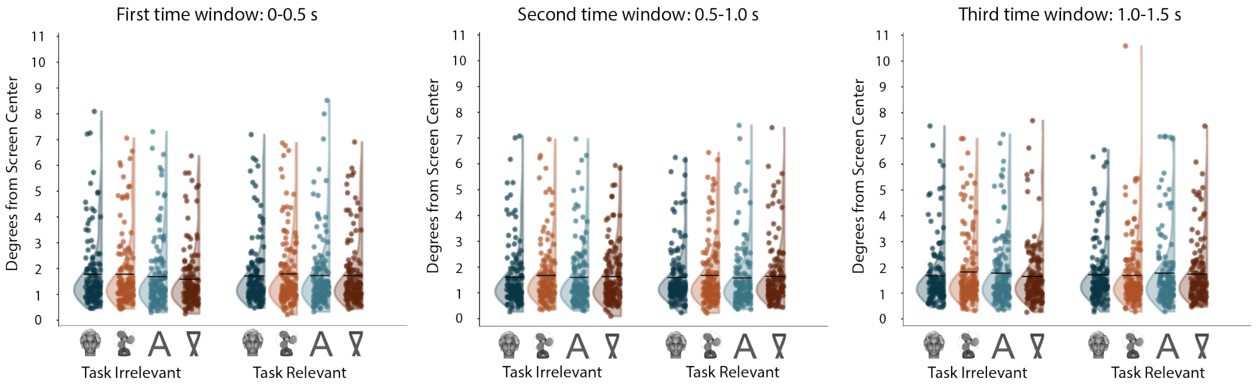

**Supplementary Figure 8**. The averaged distance of fixations from the screen center (in degrees of visual angle) for the first (left), second (middle) and third (right) time windows, in each category (horizontal axis) for task irrelevant (left) and relevant (right) stimuli. Each dot is an individual subject, plotted together with the overall distribution. The black horizontal lines mark the averaged value per condition. For illustration purposes, we excluded one outlier participant whose averaged gaze was highly affected by 2 trials where they diverted their gaze away from the center (M=16.80). This participant was not excluded from the analysis.

#### Long trials analysis: number of blinks

Here, despite the overall very low number of blinks during long stimulus duration trials (M=1.15, SD=1.10), many differences were found between the conditions (Supplementary Figure 9). For convenience, all effects are summarized in Supplementary Table 1.

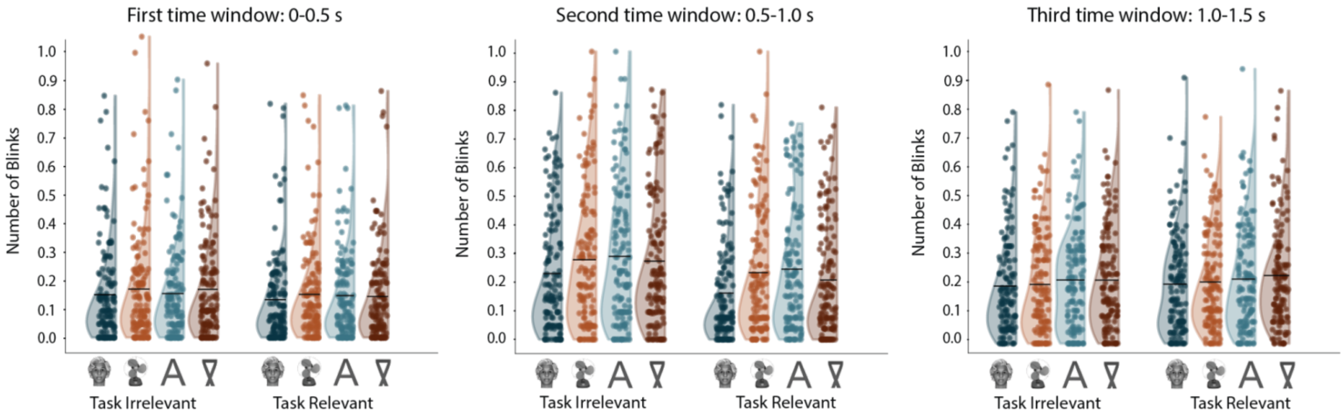

**Supplementary Figure 9**. Averaged number of blinks for the first (left), second (middle) and third (right) time window, in each category. The same conventions are used as in Supplementary Figure 8.

| | df | F | Adjusted p value |
|---|---|---|---|
| **Category** | **(3, 123183)** | **49.62** | **<0.0001 (BF=1.95x10$^{28}$)** |
| Category: face vs. false font | Inf | | <0.0001 |
| Category: face vs. letter | Inf | | <0.0001 |
| Category: face vs. object | Inf | | <0.0001 |
| Category: false font vs. letter | Inf | | 1.000 |
| Category: false font vs. object | Inf | | 0.710 |
| Category: letter vs. object | Inf | | 0.024 |
| **Task Relevance** | **(1, 123185)** | **101.90** | **<0.0001 (BF=1.59x10$^{20}$)** |
| Task Relevance: irrelevant vs. relevant | Inf | | <0.0001 |
| **Time Window** | **(2, 123180)** | **907.97** | **<0.0001 (BF=inf)** |
| Time Window: first vs. second | Inf | | <0.0001 |
| Time Window: second vs. third | Inf | | <0.0001 |
| Time Window: first vs. third | Inf | | <0.0001 |
| **Category x Task Relevance** | **(3, 123187)** | **1.27** | **1.000 (BF=0.005)** |
| **Time Window x Category** | **(6, 123180)** | **13.28** | **<0.001 (BF=2.28x10$^{11}$)** |
| First: face vs. false font | Inf | | 0.002 |
| First: face vs. letter | Inf | | 0.015 |
| First: face vs. object | Inf | | 0.020 |
| First: false font vs. letter | Inf | | 1.000 |
| First: false font vs. object | Inf | | 1.000 |
| First: letter vs. object | Inf | | 1.000 |
| Second: face vs. false font | Inf | | <0.0001 |
| Second: face vs. letter | Inf | | <0.0001 |
| Second: face vs. object | Inf | | <0.0001 |
| Second: false font vs. letter | Inf | | 0.012 |
| Second: false font vs. object | Inf | | 1.000 |
| Second: letter vs. object | Inf | | 0.215 |
| Third: face vs. false font | Inf | | 0.001 |

| | | | |
|---|---|---|---|
| Third: face vs. letter | Inf | | 0.003 |
| Third: face vs. object | Inf | | 1.000 |
| Third: false font vs. letter | Inf | | 1.000 |
| Third: false font vs. object | Inf | | 0.016 |
| Third: letter vs. object | Inf | | 0.031 |
| **Time Window x Task Relevance** | **(2, 123180)** | **69.02** | **<0.001 (BF=8.64x10$^{26}$)** |
| First: irrelevant vs. relevant | Inf | | <0.001 |
| Second: irrelevant vs. relevant | Inf | | <0.001 |
| Third: irrelevant vs. relevant | Inf | | 0.022 |
| **Time Window x Category x Task Relevance** | **(6, 123180)** | **2.19** | **0.288 (BF=0.02)** |

**Supplementary Table 1**. Statistics for all comparisons made in the analysis of number of blinks for long duration stimuli, with the factors: category, task relevance, and time window. Main effects and interactions are highlighted in bold, and significant ones are followed by post-hoc tests.

#### Long trials analysis: saccade amplitude

A main effect of time window was found on saccade amplitudes (F(2, 35796)=26.63, p<0.001, BF=3.43x10$^{8}$; Supplementary Figure 10), such that the saccade amplitude in the first time window (M=1.23, SD=1.60) was lower than in both the second (M=1.39, SD=1.86, p<0.001) and the third time windows (M=1.35, SD=1.61; p<0.001), while these latter two windows did not differ from each other (p=1.000). The interaction between category and task relevance was also significant (F(3, 35575) = 4.31, p=0.034, though BF=0.198), such that when task irrelevant, letters (M=1.47, SD=2.23) differed from both faces (M=1.25, SD=1.57; p=0.002) and objects (M=1.23, SD=1.38, p=0.044). The rest of the contrasts were not significant (p value ranges between 0.106 and 1.000). When task relevant, saccade amplitudes were lower for faces (M=1.23, SD=1.12) than objects (M=1.56, SD=2.20, p=0.034). The rest of the contrasts did not differ (p values range between 0.277 and 1.000).

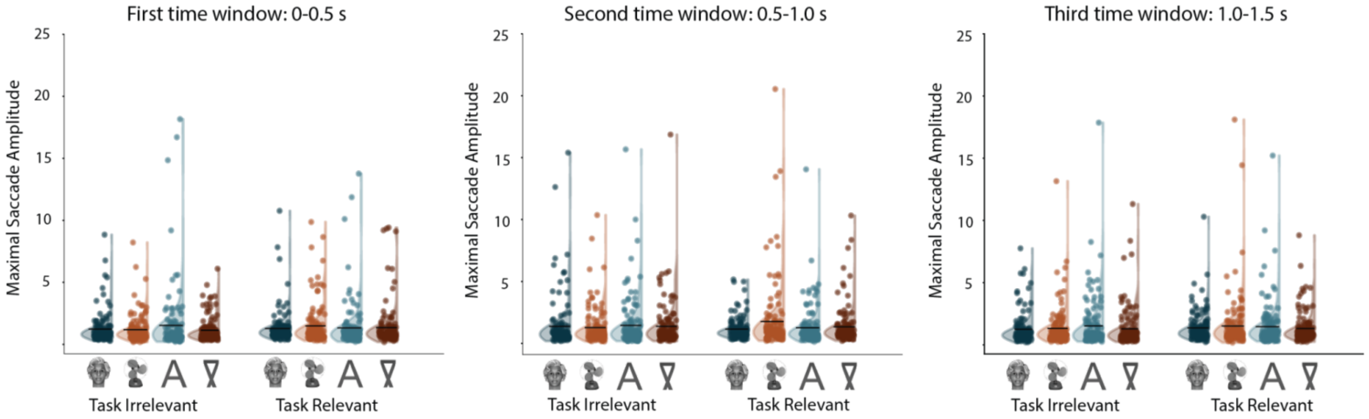

**Supplementary Figure 10**. The averaged maximal saccade amplitude (in arbitrary units) for the first, second and third time window. The same conventions are used as in Supplementary Figure 8.

#### Long trials analysis: pupil size

Though only an interaction effect was found for standardized pupil size when examining the first 0.5 s post-stimulus (with no significant post-hoc differences; see analysis above), several differences emerged when only the long stimulus duration trials were analyzed (Supplementary Figure 11). A main effect of task relevance was found ($F(1, 105556)=81.26$, $p<0.001$, $BF=4.73\times10^{15}$), with smaller pupil sizes for task irrelevant (M=1.01, SD=0.07) compared to task relevant stimuli (M=1.02, SD=0.07; $p<0.001$). Additionally, a main effect of time window was found ($F(2, 105719)=136.34$, $p<0.001$, $BF=1.30\times10^{56}$), with pupil size in the first time window (M=1.00, SD=0.06) being smaller than both the second (M=1.02, SD=0.07; $p<0.001$) and the third windows (M=1.02, SD=0.07; $p<0.001$), with no difference between the second and third windows ($p=0.443$). The interaction between task relevance and time window was also significant ($F(2, 105642)=58.13$, $p<0.001$, $BF=1.61\times10^{22}$). Post hoc analysis revealed that while in the first time window there was no difference in pupil size between task relevant and irrelevant stimuli ($p=0.132$), pupil size was smaller for task irrelevant stimuli both in the second (irrelevant: M=1.01, SD=0.07, relevant: M=1.02, SD=0.07; $p<0.001$) and third (irrelevant: M=1.01, SD=0.07, relevant: M=1.04, SD=0.07; $p<0.001$) time windows. The interaction between time window and stimulus category was also significant ($F(6, 105638)=3.51$, $p=0.013$, though $BF=0.53$), with no differences between stimuli in the first time window ($p=1.000$ for all comparisons). In the second time window, faces (M=1.01, SD=0.06) led to smaller pupil sizes than both false fonts (M=1.02, SD=0.07; $p<0.001$) and letters (M=1.02, SD=0.06; $p=0.013$; other p values range between 0.099 and 1.000). In the third time window, the only difference in pupil size was between faces (M=1.02, SD=0.06) and false fonts (M=1.03, SD=0.07, $p=0.037$). All other factors were not significant. Supplementary Table 2 summarizes the means, SDs, and statistics for the different conditions.

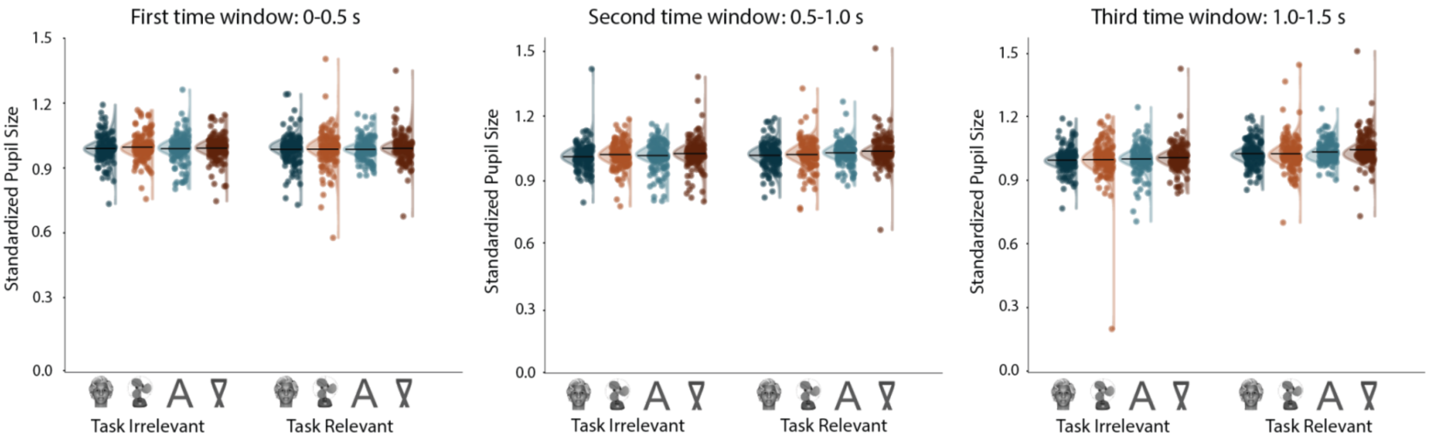

**Supplementary Figure 11**. Standardized pupil size for the different time windows. The same conventions as in Supplementary Figure 8 are used.

| | df | F | Adjusted p value |
|---|---|---|---|
| Category | (3, 228) | 3.65 | 0.093<br>(BF=0.07) |
| Task Relevance | (1, 105556) | 81.26 | <0.0001 (BF=4.73x1015) |
| Task Relevance: irrelevant vs. relevant | Inf | | <0.0001 |
| Time Window | (2, 105719) | 136.34 | <0.0001 (BF=1.30x1056) |
| Time Window: first vs. second | Inf | | <0.0001 |
| Time Window: first vs. third | Inf | | <0.0001 |
| Time Window: second vs. third | Inf | | 0.443 |
| Category x Task Relevance | (3, 105565) | 2.86 | 0.247<br>(BF=0.03) |
| Category x Time Window | (6, 105638) | 3.51 | 0.013<br>(BF=0.53) |
| Time Window x Task Relevance | (2, 105642) | 58.13 | <0.0001 (BF=1.61x1022) |
| First: irrelevant vs. relevant | Inf | | 0.132 |
| Second: irrelevant vs. relevant | Inf | | <0.0001 |
| Third: irrelevant vs. relevant | Inf | | <0.0001 |
| Time Window x Category x Task Relevance | (6, 105637) | 0.43 | 1.000<br>(BF=0.003) |

**Supplementary Table 2**. Statistics for all comparisons made in the analysis of pupil size for the long duration stimuli, with the factors: category, task relevance, and time window. The same conventions as in Supplementary Table 1 are used.

### Control experiment: surprise memory test

In our main experiment, the stimuli were presented at high contrast, in isolation, at the center of fixation, for long durations. However, given the low prevalence of the targets and the paucity of the display (a stimulus was presented approximately every 2.2 seconds), it could be argued that subjects may have missed some stimuli, not being consciously aware of some of them. This could be especially true for the task irrelevant stimuli, on which subjects had no need, and no incentive, to focus.

To investigate whether task irrelevant stimuli might not be consciously perceived during the main task, we ran a control experiment (which was already described in the published Study Protocol of this experiment[2]). We used the same stimuli and task structure as the main experiment, but performed a surprise memory task at the end of this control experiment. We used memory encoding as a proxy of visibility and focused on whether (old-new) recognition differed between the task relevant and task irrelevant conditions. We reasoned that memory performance would not be high overall, as subjects did not expect to be tested on the stimuli, and had no reason and/or incentive to memorize them. Yet we still asked if those stimuli that were task irrelevant, and potentially might not have been perceived during the memory task, give rise to differential recognition rates compared to those that were task relevant and for whom it is reasonable to assume that they were consciously seen.

#### Methods

##### Participants

Thirty-nine participants (26 females, aged between 18 and 59, mean=32.6, SD=12.82, all right-handed) took part in the study, which was run at the Max Planck Institute (MPI) for Empirical Aesthetics in Frankfurt/Germany. All participants had normal or corrected-to-normal vision. They were recruited from the participant pool of the MPI and received monetary compensation for their participation. They all provided written informed consent prior to participation. All experimental procedures were approved by the Ethics Council of the Max Planck Society.

##### Apparatus

Stimuli (6° × 6°) were presented foveally on an LCD monitor (ASUS VG24QE, 24in., refresh rate 100 Hz, resolution 1920 × 1080 pixel) in a darkened, sound-attenuating booth. Stimulus delivery and response collection were controlled using Psychtoolbox 3 [3] on Matlab 2017a, on a PC running Windows 10.

##### Stimuli and procedure

Stimuli and procedure were identical to the ones used in the main study, with the following differences: First, 20 filler stimuli (five per category) were created for the memory test stage following the same procedure as those used in that experiment. Second, only two blocks of 40 trials each were presented. Accordingly, 10 stimuli of each category (Faces, Objects, Letters,

False-fonts) were presented in a block. Third, and most importantly, an additional session - a surprise memory test – was administered after the study (henceforth, we refer to the main study as the "exposure phase"). In the surprise memory test 40 stimuli (10 from each category) from the exposure phase (old) were presented alongside the 20 filler (novel) stimuli. Subjects were asked to determine whether they had seen the stimuli in the previous exposure phase. From the 10 old stimuli belonging to a given category, half were taken from the non-targets in the task relevant condition, while the other half were taken from the task irrelevant condition. This enabled us to compare the effect of task and stimulus category on incidental memory.

The stimuli presented during the exposure phase were randomized and counterbalanced across participants such that all stimuli appeared equally often in the memory test, while also controlling for their appearance in the task condition (relevant and irrelevant), target, category, orientation and duration in the exposure phase. In each trial, a single stimulus (old/novel) was presented at fixation subtending approximately 6˚ by 6° of visual angle. The stimulus was shown until subjects responded by pressing the left or right arrow keys on a keyboard to indicate old/new. No time-out period was implemented. Key-response attribution was randomized across participants. Participants then gave a confidence rating from 1 to 5 as to how confident they were of having seen the stimuli during the exposure phase. Subjects used the 1-5 keys on a keyboard to give their response: 1 corresponds to not sure at all and 5 to being absolutely sure that they had seen the stimulus during the exposure phase. Response attribution was kept constant across participants to avoid confusion in the response mapping.

#### Analysis

Separate analyses were performed on the data of the exposure and the memory phase. Analysis on the exposure phase data was aimed at assessing subjects' overall performance in the target detection tasks. For each participant, four targets were presented, one for each category. Due to the low number of target trials, we ran a Generalized Linear Mixed (GLM) model. The dependent variable had a binomial outcome, with 1 for correct detections, 0 for misses. There were therefore four data points per participant (one per category), each with a value of one or zero depending on whether participants detected a given target. The GLM model was computed with a binomial distribution and a logit link function, with stimulus category as fixed effect and participants as random effect.

To investigate visibility across task relevant and task irrelevant stimuli, we evaluated subjects' performance in the memory test, defined as d', as well as the degree of confidence that subjects exhibited in having seen the stimuli during the exposure phase. We computed d' defining a Hit when subjects declared seeing a stimulus that was indeed presented in the exposure phase (old), and a FA when subjects declared seeing a stimulus that was not presented in the exposure phase (filler). d' was computed separately per task and stimulus category and analyzed in a two-way repeated measure ANOVA with task (relevant/irrelevant) and category (faces/objects/letters/false-fonts) as within-subject factors. The median confidence per participant was computed for the different stimuli categories and task relevance separately for correct and incorrect responses. If confidence ratings are reliable, median confidence ratings should be higher for correct decisions than for incorrect ones.

All analyses were performed in MATLAB and Statistics Toolbox (Release 2019, The MathWorks, Inc., Natick, Massachusetts, United States). The repeated measures ANOVA were computed using the fitrm function. The post-hoc multiple comparisons were performed using the multcompare command with Bonferroni correction for multiple comparison correction. Linear mixed models (LMMs) were computed using fitlme matlab function.

### Results

#### Exposure phase

Hit counts were consistently high across participants, in line with the findings of the main experiments (mean hit rate across categories = 0.92) and similar across categories ($F(3,152) = 0.24$; $p = 0.871$; Faces: 38/39; Objects:37/39; Letters: 37/39, False fonts: 32/39) indicating that all subjects complied with task instructions.

#### Memory phase

A repeated measures ANOVA on d' revealed an effect of category ($F(3,114) = 11.61$, $p<0.001$), whereby overall d' for Objects was higher than for faces (diff = 0.61, $p = .003$), letters (diff = 0.59, $p < .001$) and false-fonts (diff = 0.83, $p <0.001$) indicating that Objects were overall better remembered in this task. The effect of category was modulated by task relevance ($F(3,114) = 3.93$, $p=0.01$). As Supplementary Figure 12 shows, objects and letters were remembered similarly across the task relevance conditions ($p>0.5$); whereas memory for faces and false-fonts was higher for the task relevant than for the task irrelevant condition (face diff = 0.35, $p = .011$; false-fonts diff = 0.33, $p = .002$). Critically, no differences between the task relevant and task irrelevant conditions were observed for stimulus categories that were overall better remembered (e.g., objects); and such differences were mostly present for those stimulus categories that were less well encoded in memory (e.g., false-fonts).

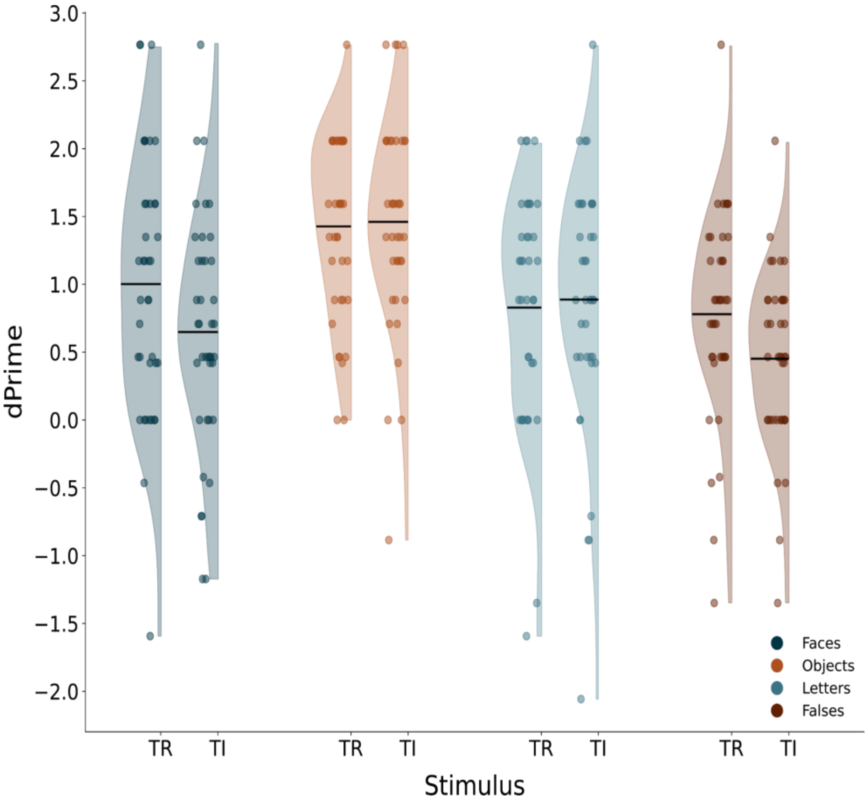

**Supplementary Figure 12**. d' in the surprise memory test for the different stimulus categories (faces: blue; objects: orange: letters: turquoise; false-fonts: brown) and task relevance conditions (TR: task relevant; TI: task irrelevant).

Next, we investigated confidence ratings. We compared the median confidence rating in a linear mixed model with factors: category, task relevance and accuracy (i.e., whether the response was correct or not). As expected, confidence ratings were higher for correct than for incorrect responses ($F(1,533) = 28.23$; $p < 0.001$), validating the subjects' responses in the task. Importantly, as shown in Supplementary Figure 13, confidence ratings were similar across tasks and categories. Accordingly, no main effect of task or interaction between task, and category or accuracy was found (all $p > 0.05$).

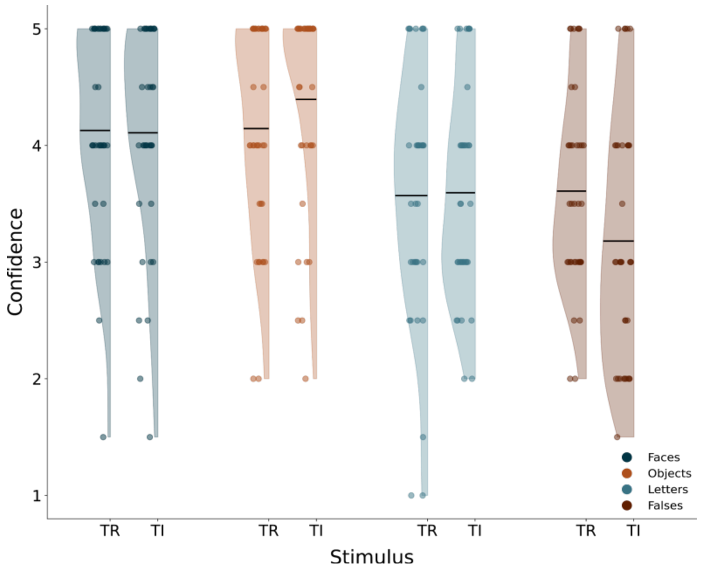

**Supplementary Figure 13**: Mean of the confidence rating medians to correct answers of the participants in the surprise memory phase for task relevant (TR) and task irrelevant (TI) stimuli. Values ranged from 1 to 5 with 1 representing not being confident at all and 5 being absolutely sure of having seen the stimulus. Other plotting conventions are the same as Supplementary Figure 12.

Confidence ratings did differ across categories ($F(1,533) = 15.26$; $p < 0.001$), and were modulated by accuracy ($F(1,533) = 4.36$; $p = 0.005$). To further investigate the interaction between accuracy and category, two separate linear mixed models were run with category as a fixed factor separately for the correct and incorrect responses. Confidence ratings across all 4 categories were comparable for the incorrect responses ($F(3,247)=2.32$, $p=.076$), whereas they differed across categories for the correct responses ($F(3,294)=21.09$, $p<.001$). For correct responses, confidence ratings for faces and objects were similar ($t(73)=-0.84$, $p=.0403$) but overall higher than both the confidence ratings for letters and false-fonts (all $p<0.001$), which were comparable among themselves ($t(69)=1.49$, $p=.140$).

### Conclusions

The results of this control experiment mitigate the concern that subjects might not have been aware of the task irrelevant stimuli (or were aware less often than for the task relevant stimuli). When presented with a surprise memory test on the previously shown stimuli, subjects' performance for task relevant and task irrelevant objects and letters did not differ. Similarly, there were no differences in confidence ratings on the memory test across task relevant vs. irrelevant stimuli. As there is no reason to think that subjects might not be aware of task relevant stimuli, the overall similar performance and confidence for task irrelevant ones renders the claim that the latter might have not been consciously perceived highly unlikely.

### Replicability of findings: optimization vs. replication results

As explained in the main text, we divided the MEG and fMRI data into two sets, with one-third of the data (i.e., the optimization dataset; 67 subjects total, with 35 fMRI and 32 MEG subjects) used for development of analysis details. This allowed us to then test the replicability of the results on the remaining two-thirds of the data (i.e., the replication dataset). Below we report the results of the optimization dataset for all analysis reported in the main paper, alongside the results of the replication dataset for comparison. The results of both the optimization and replication phase were analyzed with the exact same procedures.

#### Behavioral analysis

Overall, the optimization phase subjects showed very high hit rates (M=97.50%, SD=2.93% across all included subjects), very few false alarms (M=0.59%, SD=0.38%), and reasonably fast reaction times (M=0.64s, SD=0.10). Importantly, akin to the replication data, there were no differences in performance in the optimization data, measured using d', between labs within each modality (fMRI: $F(1, 33.02)=0.33$, $p=1.000$, BF=0.056; MEG: $F(1, 28.49)=0.13$, $p=1.000$, BF=0.055). Similarly, there were no differences in reaction times between labs within the same modality (fMRI: $F(1, 32.79)=0.004$, $p=1.000$, BF=0.02; MEG: $F(1, 30.0)=2.27$, $p=1.000$, BF=0.04). With respect to false alarms, as there were too few of them, the data could not be adequately modeled.

#### Eye movements analysis

Overall, much like the replication data, the subjects in the optimization dataset were very good at maintaining fixation (Supplementary Figure 14; for a description of saccade direction, see Supplementary Figure 15), and their ability to do so within the first 0.5 s was not modulated by task relevance (Linear Mixed Model: $F(1, 23555.5)=0.14$, $p=1.000$, BF=0.004). However, a main effect for stimulus category was not found in the optimization dataset ($F(3,232.4)=0.37$, $p=1.000$, BF=0.004), but an interaction between category and task relevance was observed ($F(3, 23573.4)=4.70$, $p=0.008$, BF=0.491): When task irrelevant, fixations were slightly closer to the center for letters (M=1.52, SD=1.24) than faces (M=1.63, SD=1.29, $p=0.04$). When task relevant, there was no significant difference between stimulus categories (p ranging between 0.194 and 1.000), like in the replication dataset.

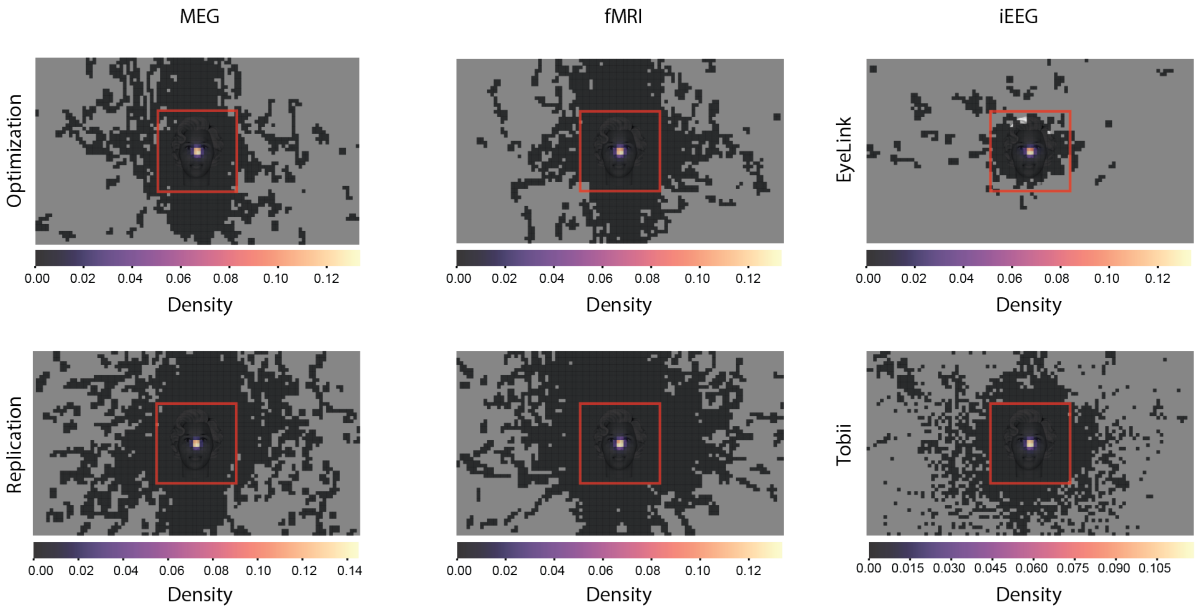

**Supplementary Figure 14**. Averaged heat maps of fixations throughout the experiment for the three modalities, for the optimization phase subjects (upper row) and the replication phase subjects (lower row), for MEG (left) and fMRI (center). The iEEG column presents the fixation patterns for the EyeLink sample (top) and the Tobii one (bottom), to complement Figure 1 in the main text, where only the stimulus area was presented.

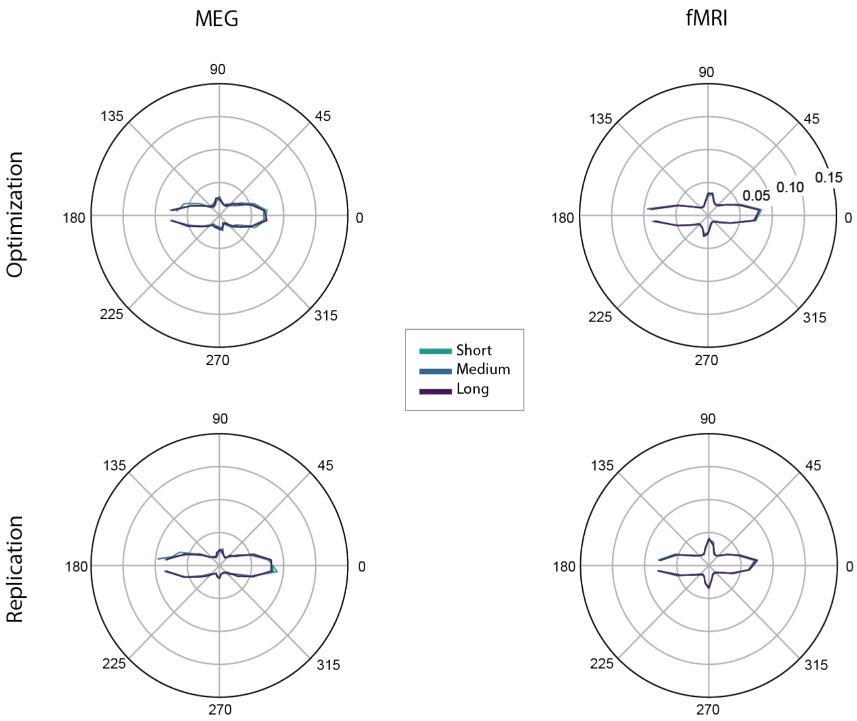

**Supplementary Figure 15**. Saccade direction for the optimization phase subjects (upper row) and the replication phase subjects (lower row), for MEG (left) and fMRI (right). Plotting conventions are the same as in Figure 1 of the main paper.

### Decoding analysis

#### MEG

Overall, the results of the optimization and replication datasets were highly consistent, with face vs. object decoding in both the posterior and prefrontal regions, which generalized from the task irrelevant to the task relevant condition and vice versa (Supplementary Figure 16).

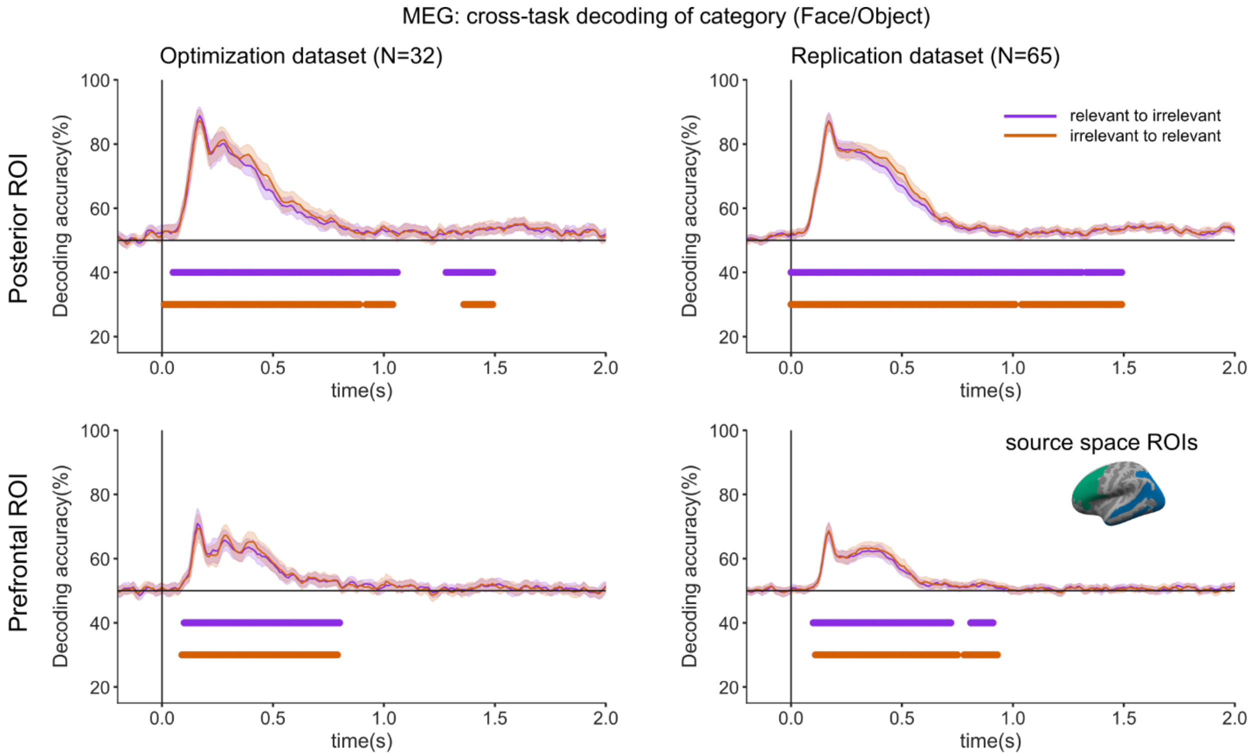

**Supplementary Figure 16**. Category decoding accuracy for the optimization (left) and replication (right) datasets, when training classifiers on the task relevant condition and testing on the task irrelevant condition (purple), or training on the task irrelevant condition and testing on the task relevant condition (orange), within MEG source space for posterior ROIs (top) and prefrontal ROIs (bottom).

Similarly, decoding of face orientation again yielded similar results for the two datasets, with strong decoding of orientation in posterior areas and very weak decoding of orientation in prefrontal ones (Supplementary Figure 17).

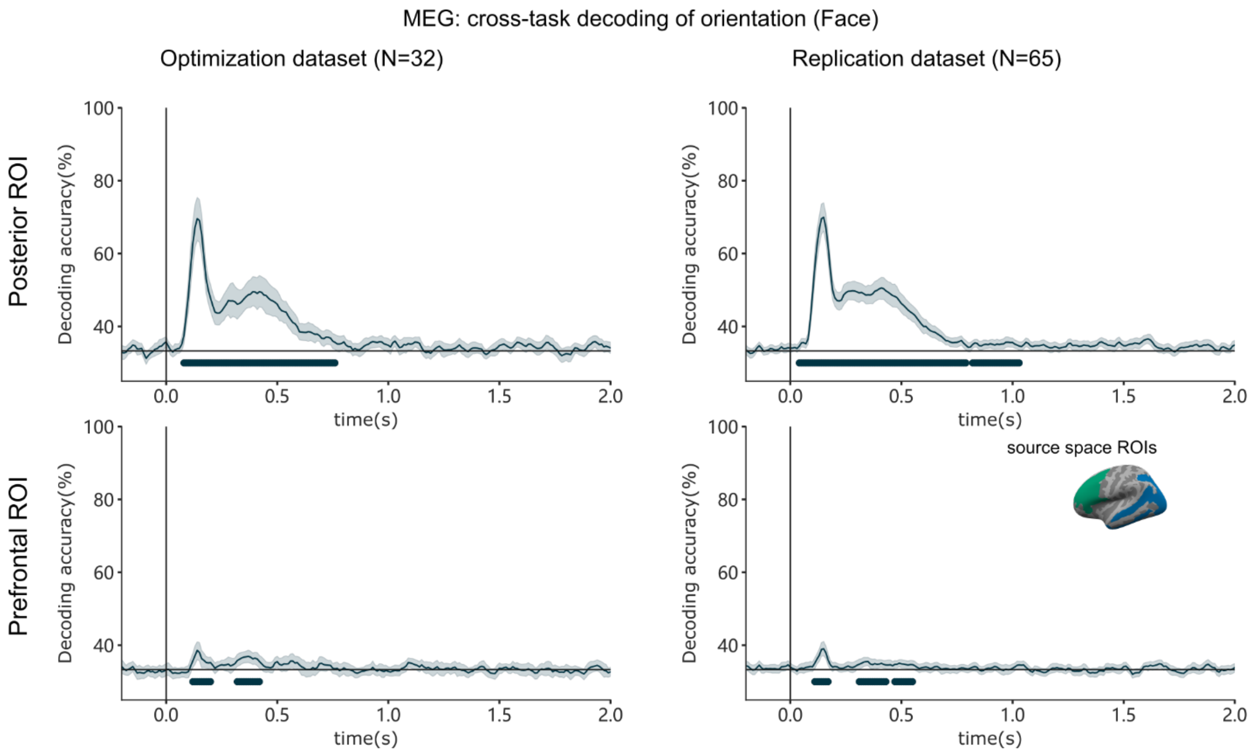

**Supplementary Figure 17**. Orientation decoding accuracy for faces (left vs. right vs. front view) compared across the optimization (left) and replication (right) datasets, when training and testing classifiers on the task irrelevant feature of face orientation (blue) in MEG source space for posterior ROIs (top) or prefrontal ROIs (bottom).

#### fMRI

Comparable results were found for the searchlight decoding between the optimization and replication datasets (Supplementary Figure 18), with significant decoding evident in similar posterior and prefrontal regions, and generalization from the task irrelevant to the task relevant conditions, and vice versa. Notably though, in the optimization results, the regions showing significant decoding were smaller in spatial extent, most likely due to the smaller sample size of the optimization dataset.

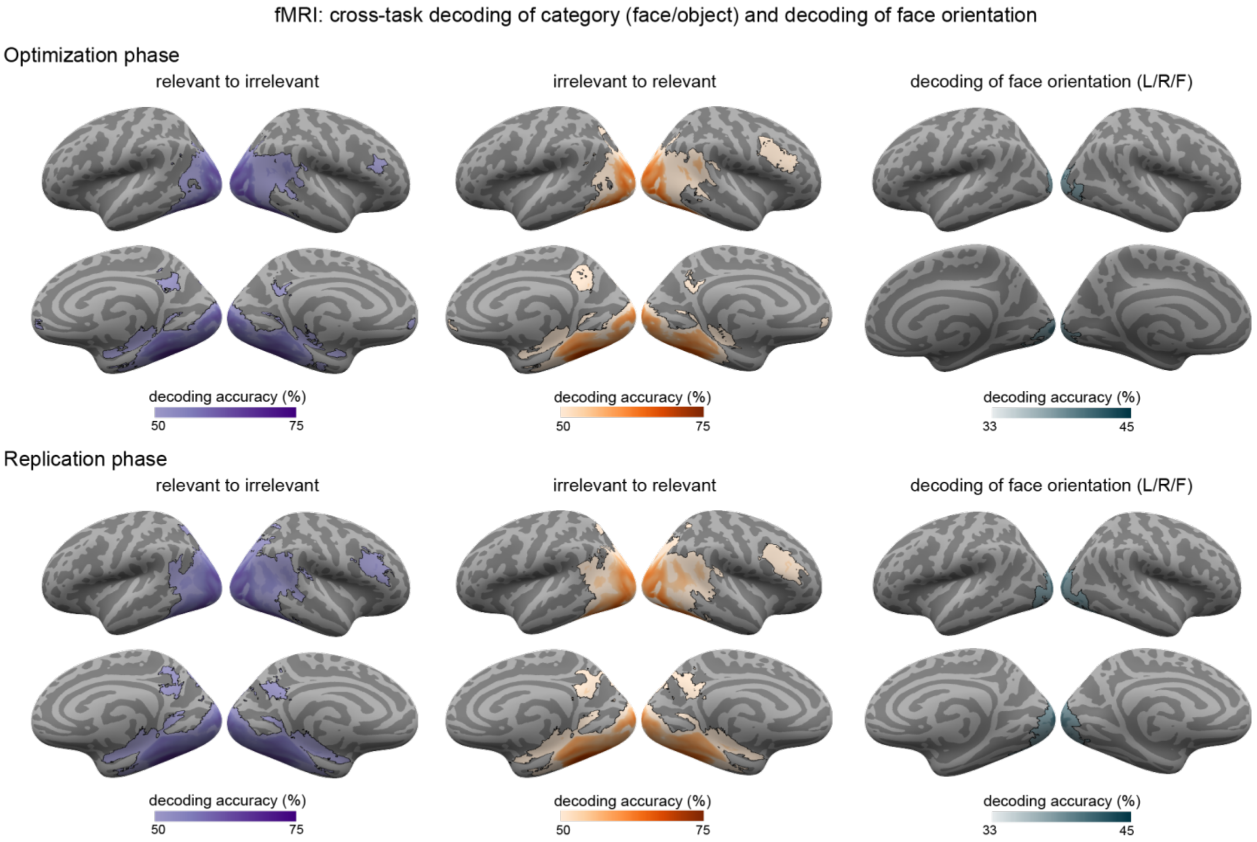

**Supplementary Figure 18**. Comparison between fMRI decoding results in the optimization (top) and replication (bottom) datasets. Cross-task decoding of stimulus category (faces vs. objects) when training classifiers on task relevant stimuli and testing on task irrelevant stimuli (left) or vice versa (center) and for face orientation (left vs. right vs. front view) which was always irrelevant, using a searchlight approach, collapsed across the three stimulus durations. Regions showing significantly above-chance (50%) decoding accuracies are indicated by the outlined colored regions on the inflated cortical surfaces (within each panel, top: left/right lateral views; bottom: right/left medial views).

### Levels of activation analysis

#### MEG

The LMM analysis on gamma and alpha band signals in the prefrontal and posterior ROIs for the optimization dataset yielded similar results as the replication dataset (Supplementary Figure 19 and Supplementary Table 3). In both datasets, none of the models derived from the theories showed a better fit for the gamma band activity compared to the non-theoretical models in any of the prefrontal or posterior parcels. However, the non-category-specific theories' models best fitted the alpha band activity in eleven parcels in the posterior ROI.

Altogether, the results of the analysis on the optimization data were very much in line with our main findings. In the gamma band, no support was found for either theory. In the alpha band, the late time bins provided compelling support for the IIT's model in the Occipital Pole, though not being content-specific. However, the phasic alpha response to stimulus onset and offset in the anterior and middle-posterior cingulate cortex found in the replication dataset was not found in the optimization dataset. This difference might either reflect lack of power in the optimization dataset given the lower number of participants, or suggest that this finding might be a false positive in the replication dataset.

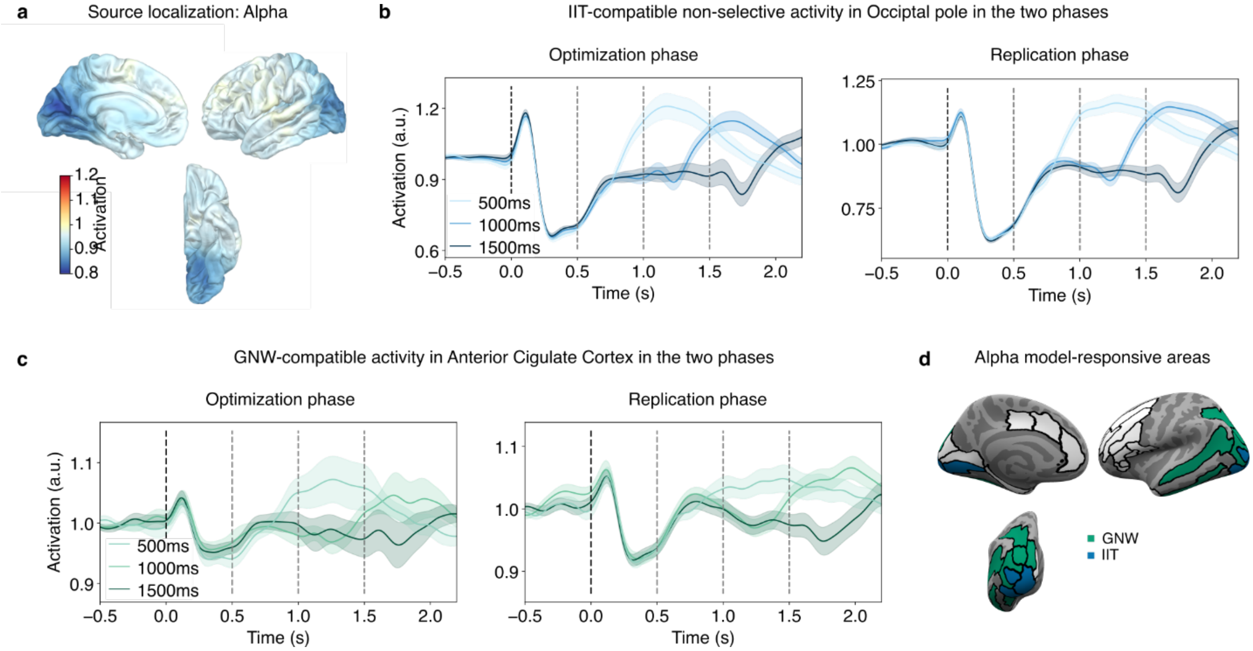

**Supplementary Figure 19**. Results of the MEG levels of activation analysis in the optimization and replication datasets. **a.** Source localization of the alpha band activity. **b.** Time course of the averaged alpha band activity in the occipital pole on optimization and replication datasets separately. **c.** Time course of the averaged alpha band activity in the anterior cingulate cortex on optimization and replication data separately. **d**. Representation of the LMM results on alpha band data for each parcel included in the prefrontal and posterior ROIs.

| ROI | # parcels | Task relevance | Signal | IIT model | IIT x Category | GNWT model | GNWT x Category |
|---|---|---|---|---|---|---|---|
| Prefrontal | 12 | Irrelevant | Alpha | 0 | 0 | 0 | 0 |
| | | | "Late" alpha | 1<br>$F_{IIT}$ = 32.78<br>$p < 0.0001$ | 0 | 0 | 0 |
| | | | Gamma | 0 | 0 | 0 | 0 |
| Posterior | 15 | Irrelevant | Alpha | 4<br>$F_{IIT}$ = 10.31<br>$p < 0.0014$ | 0 | 7<br>$F_{GNW}$ = 8.94<br>$p < 0.0029$ | 0 |

| | | | | |
|---|---|---|---|---|
| "Late" alpha | 15<br>$F_{IIT} = 8.27$<br>$p < 0.0041$ | 0 | 0 | 0 |
| Gamma | 0 | 0 | 0 | 0 |

**Supplementary Table 3**. Counts of parcels for each of the fitted models of interest per ROI on the MEG optimization dataset for alpha, "late" alpha, and gamma signal separately.

### Synchrony analysis

#### MEG

The results of the synchronization analysis conducted on the optimization dataset shared some, but not all of the results found in the replication dataset (Supplementary Figure 20). Namely, while a significant difference in phase-synchronization between the category-selective nodes and PFC was found within the 0-0.5 s time window in both datasets, no such difference was found between the face-selective node and V1/V2 in the optimization dataset. The DFC connectivity results showed the expected results between PFC and the face-selective node in both datasets during the ignition time window, as predicted by GNWT, while no sustained connectivity was found between category-selective areas and V1/V2 in the optimization dataset.

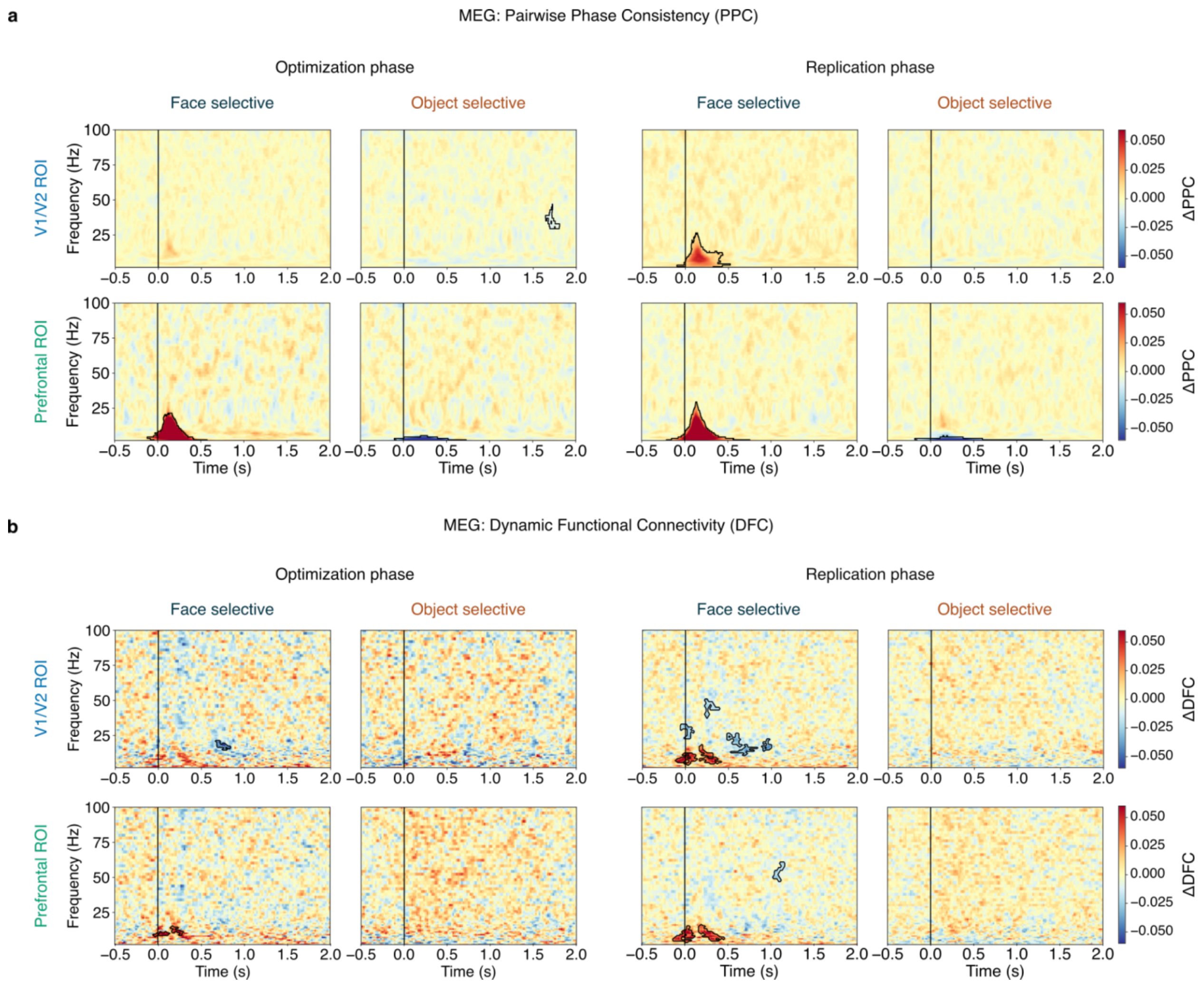

**Supplementary Figure 20**. Comparison of the MEG synchronization results between the optimization dataset (left) and the replication dataset (right). a) PPC results were consistent between replication and optimization datasets in PFC, but not in V1/V2. b) DFC results were also consistent between replication and optimization datasets in PFC, but not in V1/V2.

#### fMRI

Contrary to the other results which were highly similar between the two datasets, the gPPI findings from the optimization dataset did not match the results from the replication dataset (Supplementary Figure 21). As reported in the main text, the replication dataset results showed common clusters across irrelevant, relevant, and combined conditions in V1/V2, IPS, and IFG. On the other hand, the results from the optimization dataset showed scattered clusters that were not shared across the task irrelevant, relevant, and task combined conditions. Moreover, none of the clusters from the optimization dataset survived the correction for multiple comparisons, while only clusters from the combined analysis of the replication dataset survived the correction. We suggest that this discrepancy reflects the fact that the gPPI analysis requires a very large sample size [4], and accordingly yielded results only in the replication dataset, where we had twice the number of subjects.

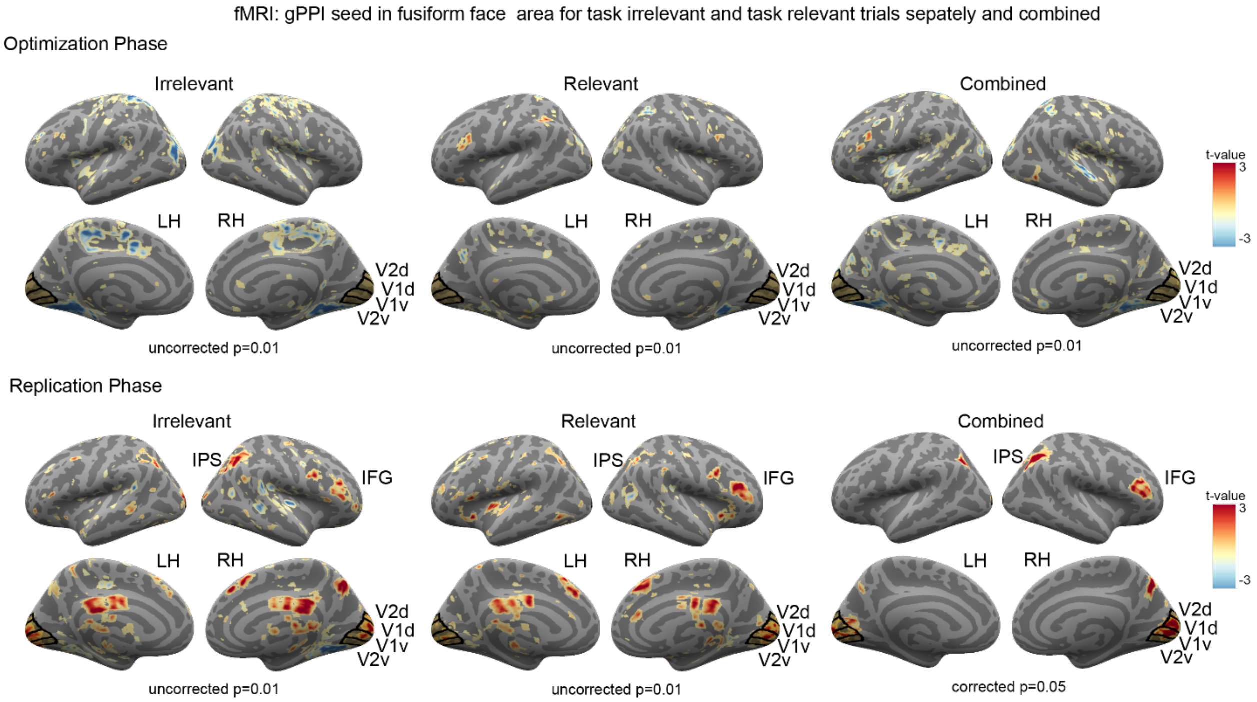

**Supplementary Figure 21**. Comparison between the main fMRI gPPI results with FFA as a seed in the optimization dataset (top) and the replication dataset (bottom). Results from the optimization and replication datasets were not consistent with each other, most likely due to the relatively smaller sample size (N=35) in the optimization dataset compared to the replication dataset (N=73).

### Putative NCC analysis

#### fMRI

Almost all of the areas identified as putative NCCs in the optimization dataset fell within the areas which were also identified in the replication dataset (Supplementary Figure 22). Notably, in the replication data we found more extended pNCCs (e.g., Fusiform Gyrus for letters and false fonts, Inferior Frontal Gyrus pars Opercularis for all stimulus categories), presumably because of the larger sample size compared with the optimization dataset.

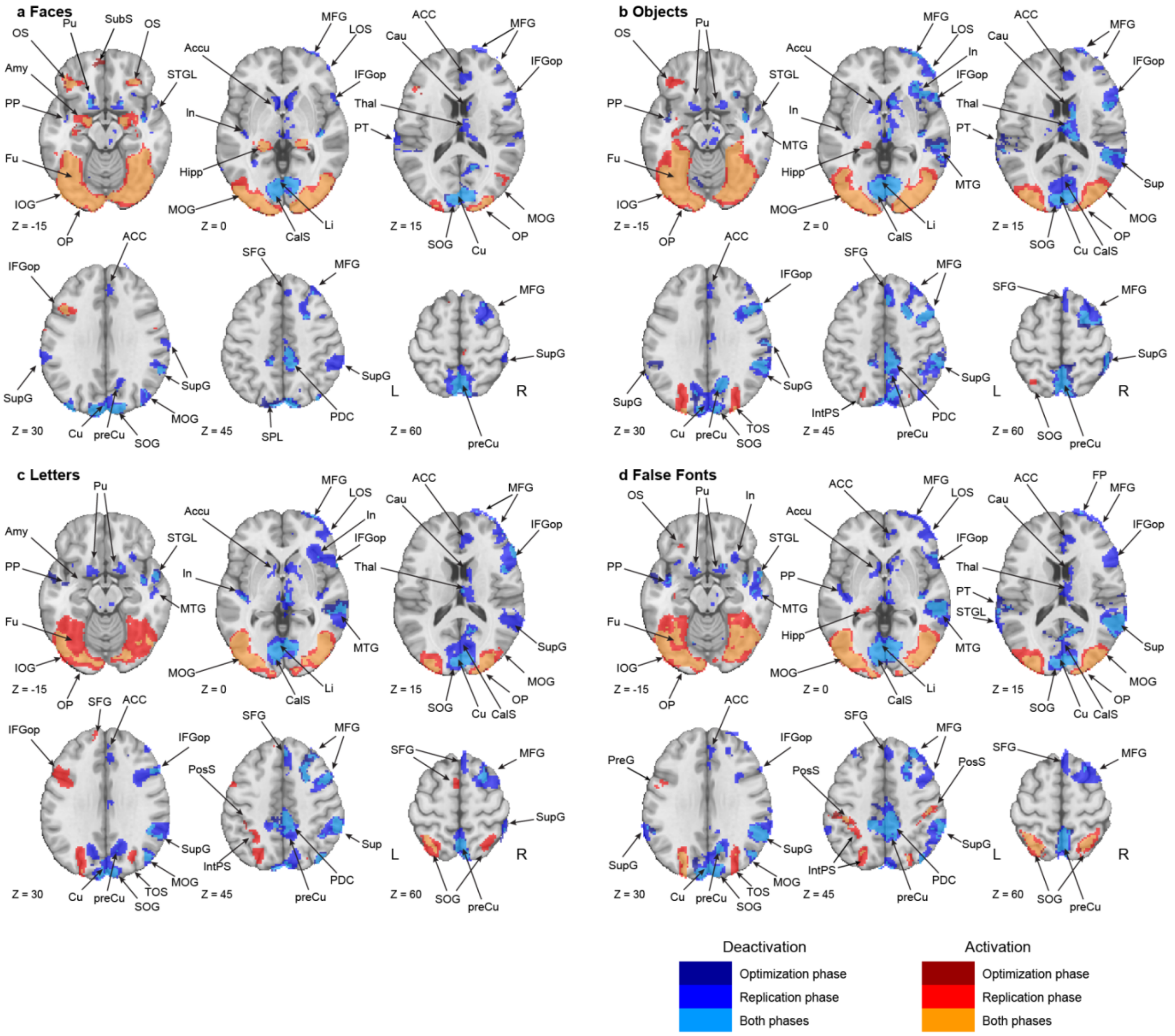

**Supplementary Figure 22**. Putative NCC analysis results for each of the stimulus categories (**a.** faces; **b.** objects; **c.** letters; **d.** false fonts), for the optimization and replication datasets. Color codes indicate in which of the datasets (optimization, replication, or both) results were found: Results that were found only in the optimization dataset are marked in dark red/blue for activations and deactivations, respectively, results found only in the replication dataset are marked in intermediate red/blue, and results found in both datasets (i.e., the overlap) are marked in light blue/orange. Abbreviations: ACC: Anterior Cingulate Gyrus; Accu: Nucleus Accumbens; Amy: Amygdala; CalS: Calcarine Sulcus; Cau: Caudate Nucleus; Cu: Cuneus; Fu: Fusiform gyrus; Hipp: Hippocampus; IFGop: Opercular part of the Inferior Frontal Gyrus; In: Insula; IOG: Inferior Occipital Gyrus; Li: Lingual Gyrus; LOS: Lateral Orbital Sulcus; MFG: Middle Frontal Gyrus; MOG: Middle Occipital Gyrus; MTG: Middle Temporal Gyrus; OP: Occipital Pole; OS: Orbital Sulci; PDC: Posterior Dorsal Cingulate; PP: Planum Polare of the superior temporal gyrus; preCu: Precuneus; PreG: Precentral Gyrus; PT: Planum Temporale of the Superior Temporal Gyrus; Pu: Putamen; SFG: Superior Frontal Gyrus; SOG: Superior Occipital Gyrus; STGL: Lateral aspect of the Superior Temporal Gyrus; SupG: Supramarginal Gyrus; Thal: Thalamus; TOS: Transverse Occipital Sulcus; SubS: Suborbital Sulcus; SPL: Superior Parietal Lobule.

### Prediction #1: Decoding of conscious content

#### Pre-registered analyses

##### fMRI: Category Decoding

In the main paper, we report fMRI decoding results for face vs. objects (Figure 2 and Extended Data Table 4) and letters vs. false fonts (Extended Data Figure 1) using a searchlight decoding approach. For completeness, we report here a table with results for the letters vs. false fonts searchlight decoding (Supplementary Table 4).

| **Anatomical (Destrieux atlas) ROIs** | **Irrelevant–Relevant** | | **Relevant Irrelevant** | | **Irrelevant** | | **Relevant** | |
|---|---|---|---|---|---|---|---|---|
| | n voxels | % voxels | n voxels | % voxels | n voxels | % voxels | n voxels | % voxels |
| **Posterior ROI** | | | | | | | | |
| G_and_S_occipital_inf | 1135 | 57 | 1053 | 52 | 748 | 37 | 1181 | 59 |
| G_oc-temp_lat-fusifor | 280 | 11 | 280 | 11 | 85 | 3 | 535 | 21 |
| G_occipital_middle | 902 | 37 | 889 | 36 | 379 | 15 | 1389 | 56 |
| S_oc_middle_and_Lunatus | 587 | 58 | 580 | 57 | 150 | 15 | 713 | 71 |
| G_cuneus | 293 | 12 | 277 | 11 | 139 | 6 | 285 | 11 |
| G_occipital_sup | 309 | 16 | 303 | 15 | 109 | 6 | 508 | 26 |
| G_oc-temp_med-Lingual | 347 | 12 | 340 | 11 | 223 | 7 | 228 | 8 |
| G_oc-temp_med-Parahip | 0 | 0 | 0 | 0 | 0 | 0 | 1 | 0 |
| G_temporal_inf | 362 | 25 | 329 | 22 | 133 | 9 | 638 | 44 |
| Pole_occipital | 984 | 41 | 959 | 40 | 580 | 24 | 1102 | 45 |
| Pole_temporal | 0 | 0 | 0 | 0 | 0 | 0 | 0 | 0 |
| S_calcarine | 124 | 5 | 131 | 5 | 55 | 2 | 121 | 5 |
| S_intrapariet_and_P_trans | 250 | 7 | 276 | 7 | 27 | 1 | 1368 | 36 |
| S_oc_sup_and_transversal | 464 | 33 | 488 | 34 | 63 | 4 | 876 | 62 |
| S_temporal_sup | 0 | 0 | 0 | 0 | 0 | 0 | 48 | 1 |
| **PFC ROI** | | | | | | | | |
| G_and_S_cingul-Mid-Post | 0 | 0 | 0 | 0 | 0 | 0 | 0 | 0 |
| Lat_Fis-ant-Horizont | 0 | 0 | 0 | 0 | 0 | 0 | 0 | 0 |
| Lat_Fis-ant-Vertical | 0 | 0 | 0 | 0 | 0 | 0 | 0 | 0 |
| G_and_S_cingul-Ant | 0 | 0 | 0 | 0 | 0 | 0 | 0 | 0 |
| G_and_S_cingul-Mid-Ant | 0 | 0 | 0 | 0 | 0 | 0 | 0 | 0 |

| | | | | | | | | |
|---|---|---|---|---|---|---|---|---|
| G_front_inf-Opercular | 0 | 0 | 0 | 0 | 0 | 0 | 23 | 1 |
| G_front_inf-Orbital | 0 | 0 | 0 | 0 | 0 | 0 | 0 | 0 |
| G_front_inf-Triangul | 0 | 0 | 0 | 0 | 0 | 0 | 0 | 0 |
| G_front_middle | 0 | 0 | 0 | 0 | 0 | 0 | 67 | 1 |
| S_front_middle | 0 | 0 | 0 | 0 | 0 | 0 | 0 | 0 |
| S_front_sup | 0 | 0 | 0 | 0 | 0 | 0 | 52 | 1 |
| S_front_inf | 0 | 0 | 0 | 0 | 0 | 0 | 1 | 0 |

**Supplementary Table 4**: Number of voxels in each ROI detected in the searchlight decoding of category (letters vs. false fonts) broken down by theory-defined ROI and for cross-task and within-task decoding.

#### MEG: temporal generalization of category decoding

In the main paper (Figure 2c), we presented results for the cross-task decoding of stimulus category (faces vs. objects). Below we show results of the cross-temporal generalization of decoding in posterior and prefrontal areas, both for faces vs. objects and letters vs. false fonts.

Supplementary Figure 23 shows face vs. object decoding: posterior regions exhibited decoding of category which spread across time, with regions showing decoding at both early and late time windows. Thus, decoding patterns appear to be present, disappear and are then reinstated at later times regardless of the stimulus duration. In contrast, prefrontal regions displayed only an early and transient decoding profile, predominantly along the diagonal of the cross-time decoding generalization matrix, albeit with more limited temporal generalization within this restricted time-window. While the temporal generalization patterns in PFC were consistent with those observed in the iEEG data (Figure 2b); those within the posterior cortex were different than the iEEG data, which showed a clear pattern of duration-tracking. The source of this difference between the MEG and iEEG results in posterior regions is unclear, but may stem from the less precise spatial localization of MEG compared to iEEG, or alternatively from the different brain signal used for the two decoding analyses (High Gamma (HG) for iEEG, due to its tight relation with spiking activity, and low frequencies in the LFP for MEG). Further studies are required to better understand the sources of these discrepancies between MEG and iEEG decoding in posterior cortex.

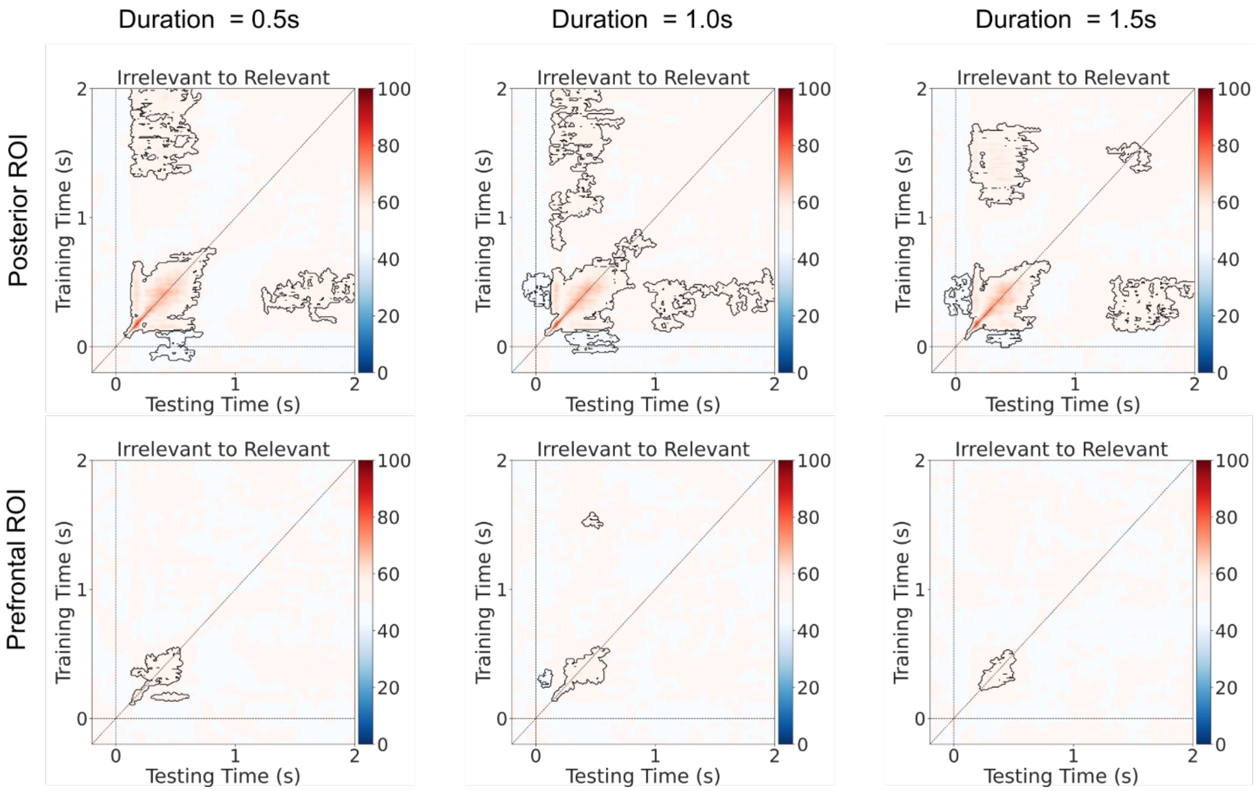

**Supplementary Figure 23.** Cross-time and cross-task generalization analysis for face vs. object decoding, shown separately per stimulus duration (left: 0.5 s, middle: 1.0 s, right: 1.5 s). Task irrelevant trials were used for training and task relevant trials for testing the pattern classifiers. Upper row: posterior ROI. lower row: prefrontal ROI.

The same analysis was performed for decoding of letters vs. false fonts. As shown in Supplementary Figure 24, posterior regions showed a transient profile along the diagonal of the cross-time decoding generalization matrix, while prefrontal regions did not contribute to above-chance decoding of these stimulus categories when broken-down by stimulus duration (significant letter vs. false font decoding was only evident when trials across all stimulus durations were combined, as reported in Extended Data Figure 1).

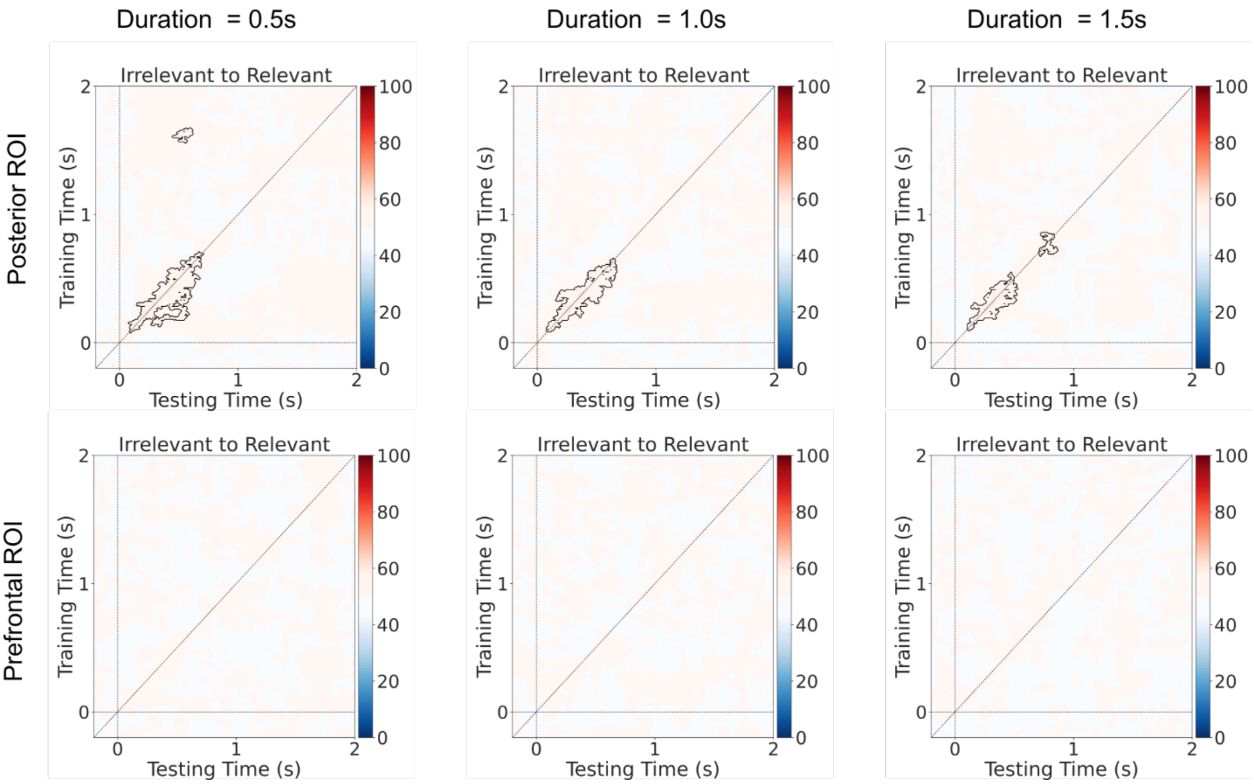

**Supplementary Figure 24.** Cross-time and cross-task generalization analysis for letters vs. false font decoding, shown separately per stimulus duration (left: 0.5 s, middle: 1.0 s, right: 1.5 s). Task irrelevant trials were used for training and task relevant trials for testing the pattern classifiers. Upper row: posterior ROI. lower row: prefrontal ROI.

#### fMRI: ROI-based category decoding

In the main paper we reported fMRI decoding results with a searchlight approach. The main advantage of this approach is that results reflect an unrestricted search across the entire brain, yet they may lack sensitivity. As GNWT and IIT proponents pre-defined anatomical ROIs reflecting their predictions (see Extended Data Table 2: Anatomical Regions-of-interest (ROIs)), we implemented a decoding approach that decodes stimulus category within each of the theory-defined ROIs separately, to maximize sensitivity. This ROI-based approach further enabled us to better compare the results across modalities (e.g., with the iEEG, which were carried out on the theory-defined ROIs only).

Similar to the searchlight decoding approach, stimulus category was decoded in the task relevant and task irrelevant conditions separately. In addition, generalization of decoding from task relevant to task irrelevant conditions, and vice versa, was tested. For the task relevant/irrelevant conditions, faces vs. objects and letters vs. false fonts classification were conducted in a leave-one-run-out cross validation scheme. To test for generalization of category decoding across conditions, faces vs. objects and letters vs. false fonts classification were done by training the classification model on one condition and testing on the other condition. Parameter estimate maps[5] of the categories of interest and a SVM classifier were employed to identify stimulus category similar to the searchlight approach.

A permutation test was used to evaluate the statistical significance of decoding within each ROI and test whether the ROIs show accuracies that significantly exceeded the chance level (> 0.5). Correction for multiple comparisons across ROIs was performed using the false discovery rate (FDR) method ($p < 0.05$). Significant ROIs were identified and the corresponding average accuracy across subjects at each of these ROIs were calculated and displayed on a brain surface (Supplementary Figure 25).

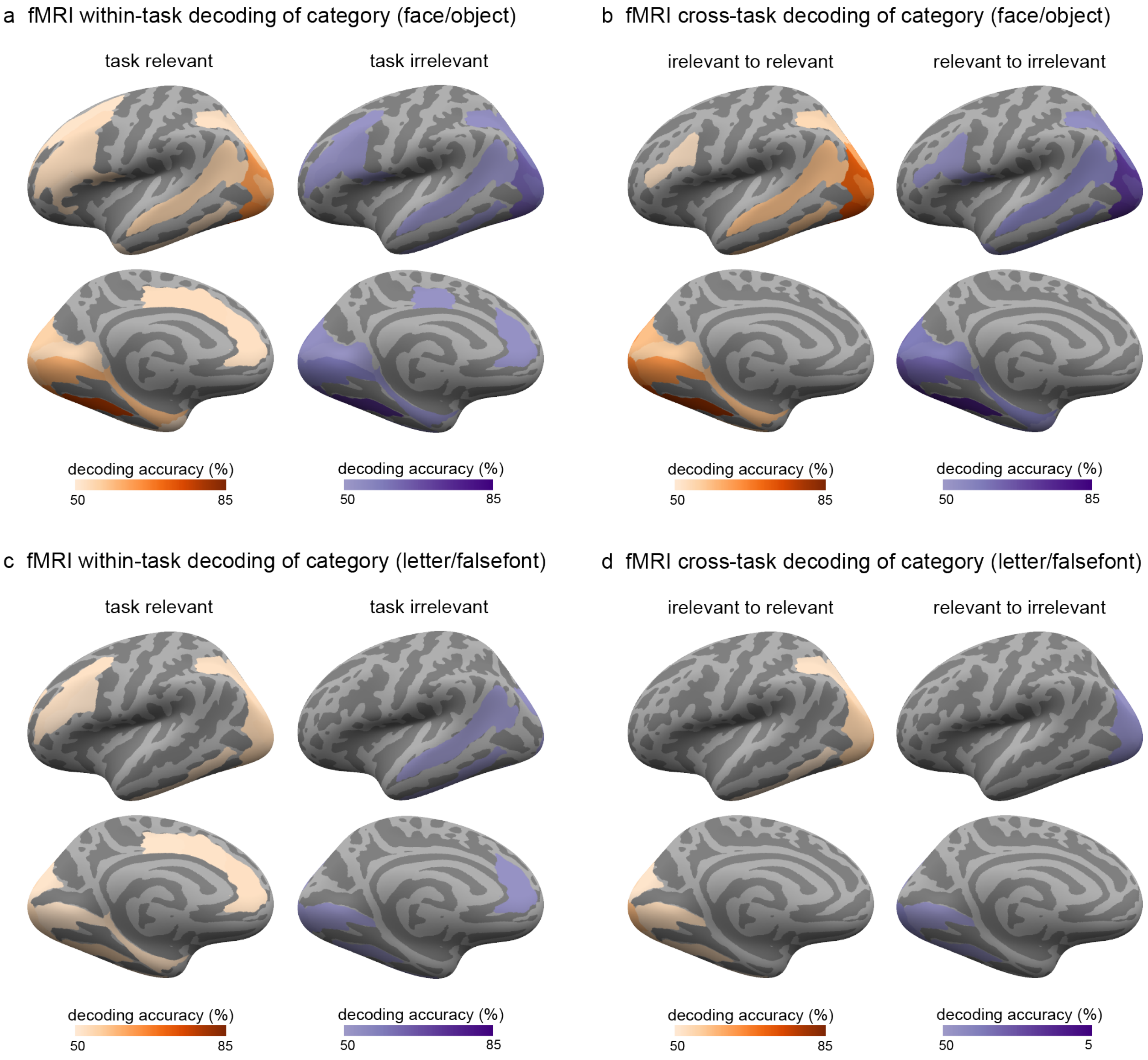

**Supplementary Figure 25**. Within-task and cross-task decoding of stimulus category (**a & b**: faces vs. objects; **c & d**: letters vs. false fonts) in fMRI using an ROI approach, collapsed across the three stimulus durations. Regions showing significantly above-chance (50%) decoding are indicated by the outlined colored regions on the inflated cortical surfaces (top: left lateral views; bottom: left medial views).

In the posterior cortex, faces vs. objects decoding showed significant cross-task generalization in occipital, posterior temporal, and posterior parietal cortex. In the prefrontal cortex, significant cross-task decoding of category was also observed in the inferior frontal sulcus (both relevant to irrelevant and irrelevant to relevant generalization) and the opercular part of the inferior frontal gyrus (relevant to irrelevant generalization). Significant within-task decoding was also found in both prefrontal and posterior cortices. Letters vs. false fonts decoding showed cross-task generalization and within-irrelevant decoding only in the posterior cortex while within-task decoding of task relevant stimuli showed significance in both posterior and prefrontal regions.

#### Orientation decoding within each stimulus category

In the main paper, we reported results for orientation decoding for faces. Below we provide the full set of results for all stimulus categories (i.e., faces, objects, letters and false fonts) across all three data modalities (i.e., iEEG, MEG and fMRI). Across all techniques, we found evidence for orientation decoding in posterior areas. In PFC however, only MEG showed above chance decoding, and only for face stimuli. However, as reported in Extended Data Figure 5b, the possibility of leakage from posterior areas in MEG source space could not be ruled out.

##### iEEG: Orientation decoding within each stimulus category

Orientation decoding (left vs. right vs. front view faces) was performed on each stimulus category separately, both with and without pseudotrial aggregation (see Methods) for each theory ROI separately (N=29, GNWT ROIs $N_{electrodes}$=576, IIT ROIs $N_{electrodes}$=583). All task conditions were collapsed as orientation was always task irrelevant, and all stimulus durations were collapsed to increase trial numbers entered into the analysis. In posterior ROIs, stimulus orientation was decodable for all stimulus categories besides objects in an early time window (e.g., < 0.5 s) when using pseudotrial aggregation (Supplementary Figure 26). In prefrontal ROIs, stimulus orientation was not decodable for any category, with or without pseudotrial aggregation.

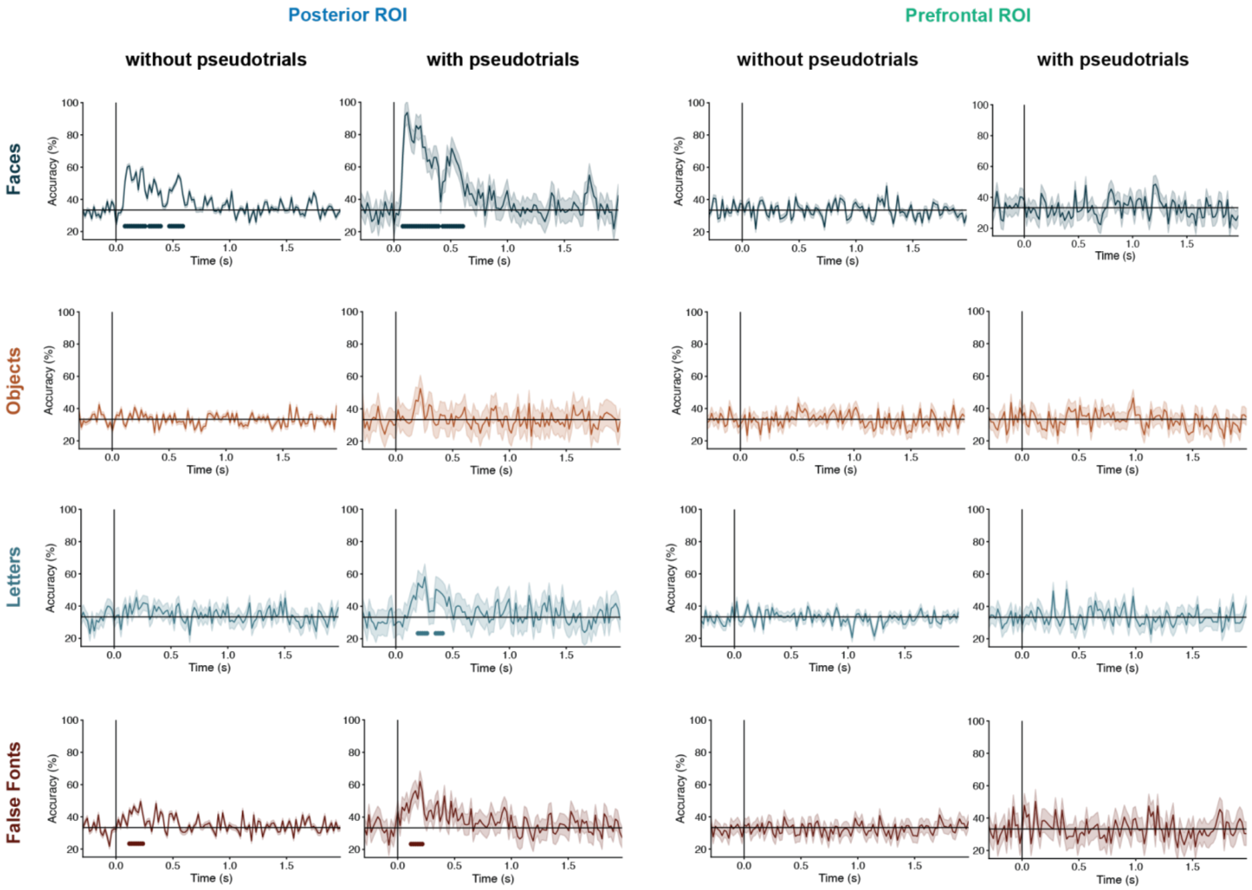

**Supplementary Figure 26.** Decoding of stimulus orientation (left vs. right vs. front views), which was always task irrelevant, shown for all stimulus categories within posterior ROIs (left 2-columns) and prefrontal ROIs (right 2-columns), collapsed across the three stimulus durations. Separate classifiers were trained without (left) and with (right) pseudotrial aggregation. Lines under the decoding functions indicate time-points showing above chance (33%) decoding accuracies.

#### MEG: Orientation decoding within each stimulus category

As shown in Supplementary Figure 27, in the posterior ROI we observed significant decoding of orientation (left vs. right vs. front views) for all four categories, while in the prefrontal ROI we only observed above-chance decoding of orientation for the face category. Like in iEEG, all stimulus durations and tasks-conditions were combined for this analysis to increase sensitivity.

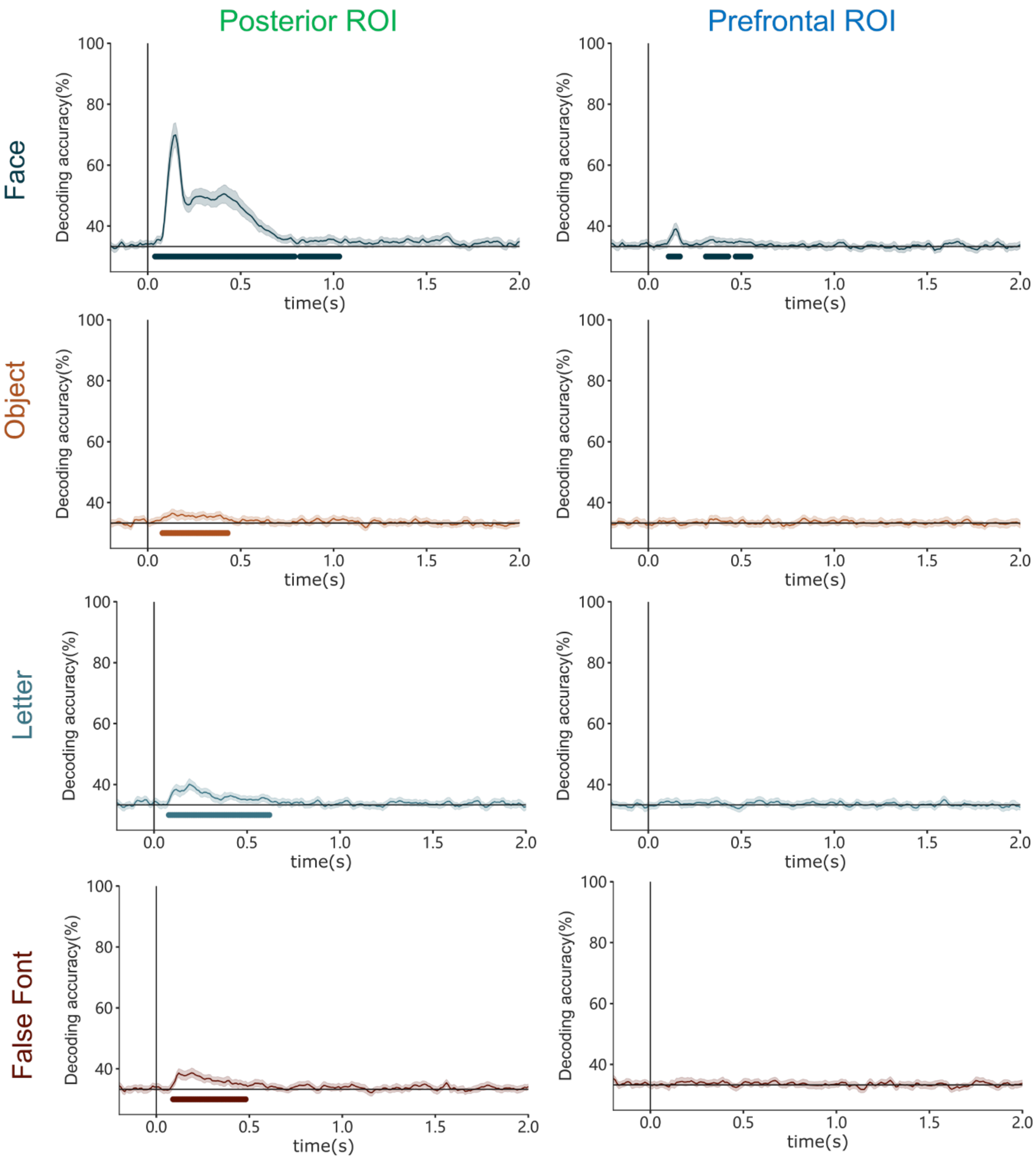

**Supplementary Figure 27**. Orientation decoding in MEG source space for each stimulus category in posterior (left) and prefrontal ROIs (right). Lines under the decoding functions indicate time-points showing above chance (33%) decoding accuracies.

#### fMRI: Searchlight decoding of orientation within each stimulus category

We tested orientation decoding (left vs. right vs. front views) for faces, objects, letters, and false fonts using the leave-one-run-out approach. The analyses were performed using the same searchlight decoding pipeline described in the main text.

Significant orientation decoding was observed in different regions of the posterior cortex for faces, letters, and false fonts while there were no significant regions anywhere in the brain showing object orientation decoding. Specifically, the occipital pole and superior occipital gyrus showed decoding accuracies significantly above-chance for orientation of faces, letters, and false fonts (Supplementary Figure 28). No prefrontal regions showed above-chance orientation decoding for any of the stimulus categories.

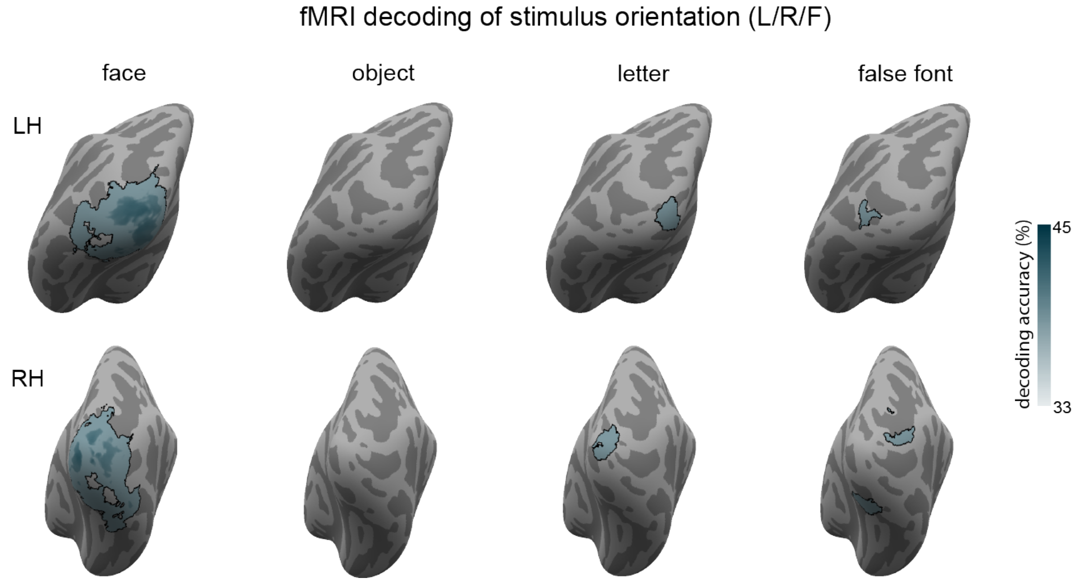

**Supplementary Figure 28**. Decoding of orientation (left vs. right vs. front view faces) in fMRI using the searchlight approach for faces, objects, letters, and false fonts. Regions with significantly above-chance (33%) decoding accuracies are indicated in outlined blue on the inflated cortical surface maps (top: left posterior views; bottom: right posterior views). No regions in prefrontal cortex showed above-chance decoding of orientation for any of the stimulus categories.

### Prediction #2: Maintenance of conscious content over time

#### Pre-registered analyses: tracking of duration

In the main paper, we reported the results for the theories' predictions on modulations in the gamma band power (iEEG & MEG). However, the preregistered predictions specified that they could be met in either the gamma, the alpha bands, or in Event-Related Potentials/Fields (ERPs/ERFs). For completeness, we report all of these results here.

##### iEEG: duration tracking in the different signals

In Supplementary Table 5, we describe the results for all conditions and all preregistered signals. The number of electrodes aligned with each theoretical model within each task condition, and for each type of signal (alpha, HG power and event-related potentials, ERPs) is provided. This table is complemented by Supplementary Figure 29, where these same electrodes are displayed on the brain surface, showing how they are localized in the brain.

| ROI | # electrodes | Task relevance | Signal | IIT model | IIT x Category | GNWT model | GNWT x Category |
|---|---|---|---|---|---|---|---|
| PFC | 655 | Irrelevant | Alpha | 0 | 0 | 0 | 0 |
| | | | HGP | 0 | 0 | 0 | 0 |
| | | | ERP | 2<br>$F_{IIT} > 24.67$<br>$P < 0.0001$ | 0 | 1<br>$F_{GNWT} = 34.83$<br>$P < 0.0001$ | 0 |
| | | Relevant | Alpha | 0 | 0 | 0 | 0 |
| | | | HGP | 0 | 0 | 1<br>$F_{GNWT} = 28.80$<br>$P < 0.0001$ | 0 |
| | | | ERP | 1<br>$F_{IIT} = 86.49$<br>$P < 0.0001$ | 2<br>$F_{IIT \times cate} > 5.76$<br>$P < 0.001$ | 3<br>$F_{GNWT} > 21.01$<br>$P < 0.0001$ | 0 |
| Posterior | 657 | Irrelevant | Alpha | 1<br>$F_{IIT} = 10.51$<br>$P = 0.001$ | 0 | 5<br>$F_{GNWT} > 16.59$<br>$P < 0.0001$ | 0 |
| | | | HGP | 12<br>$F_{IIT} > 27{,}26$<br>$P < 0.0001$ | 13<br>$F_{IIT \times cate} > 7.54$<br>$P < 0.0001$ | 11<br>$F_{GNWT} > 20.55$<br>$P < 0.0001$ | 0 |
| | | | ERP | 11<br>$F_{IIT} = 23.45$<br>$P < 0.0001$ | 1<br>$F_{IIT \times cate} = 8.88$<br>$P < 0.0001$ | 29<br>$F_{GNWT} > 20.24$<br>$P < 0.0001$ | 2<br>$F_{GNWT\ x\ cate} >$ 4.99<br>$P < 0.001$ |

| | | | | | |
|---|---|---|---|---|---|
| Relevant | Alpha | 1 $F_{IIT} = 36.90$ P < 0.0001 | 0 | 6 $F_{GNWT} >15.29$ P < 0.0001 | 0 |
| | HGP | 13 $F_{IIT} > 14.70$ P < 0.001 | 22 $F_{IIT \times cate} > 3.1$ P < 0.02 | 13 $F_{GNWT} > 19.35$ P < 0.0001 | 1 $F_{GNWT \times cate} =$ 2.10 P = 0.09 |
| | ERP | 16 $F_{IIT} = 27.23$ P < 0.0001 | 1 $F_{IIT \times cate} = 4.16$ P = 0.006 | 31 $F_{GNWT} > 11.25$ P < 0.0001 | 0 |

**Supplementary Table 5.** Number of prefrontal and posterior electrodes with a significant fit to the theories' models in the LMM analysis, separately for the alpha, HGP and ERP analysis (prefrontal ROI N subjects=31, posterior ROI N subjects=31).

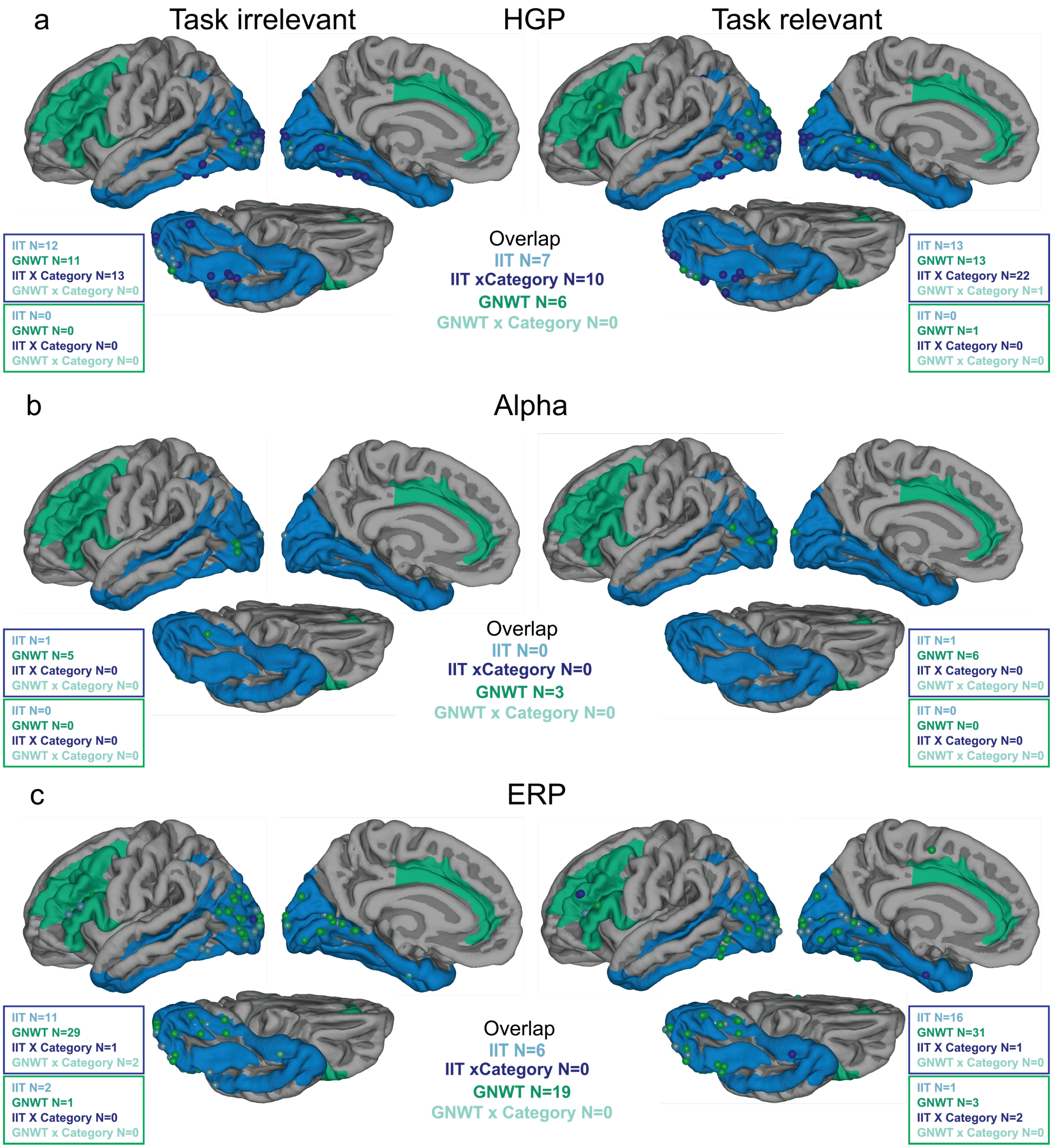

**Supplementary Figure 29***:* Location of electrodes found to be consistent with the theories predicted activation patterns in the task irrelevant (left) and relevant (right) conditions separately in the HGP (a), alpha (b) and ERP (c) signals. In each case, the count of electrodes is reported for each theory ROI (blue box for posterior ROI, N subjects=31, green box for prefrontal ROI, N subjects=31) per theory-derived model separately for each task relevance condition, as well as the overlap (i.e., counts of electrodes for which the same model was of best fit for both task relevant and irrelevant condition, labeled as overlap, and presented in the middle of the figure).

As both the figure and the table show, in the task irrelevant condition the results of the alpha band analysis did not provide strong support for either of the theories: none of the prefrontal electrodes showed the GNWT expected activity pattern, while in posterior sites, only one electrode showed the expected IIT pattern, and none showed the expected interaction between the IIT model and stimulus category. The ERP results, however, did provide some support for GNWT and IIT, with one prefrontal electrode whose activity was consistent with the GNWT prediction, 11 posterior electrode whose activity fitted the IIT model, and 1 electrodes showing the interaction between the IIT model and category (Supplementary Figure 30).

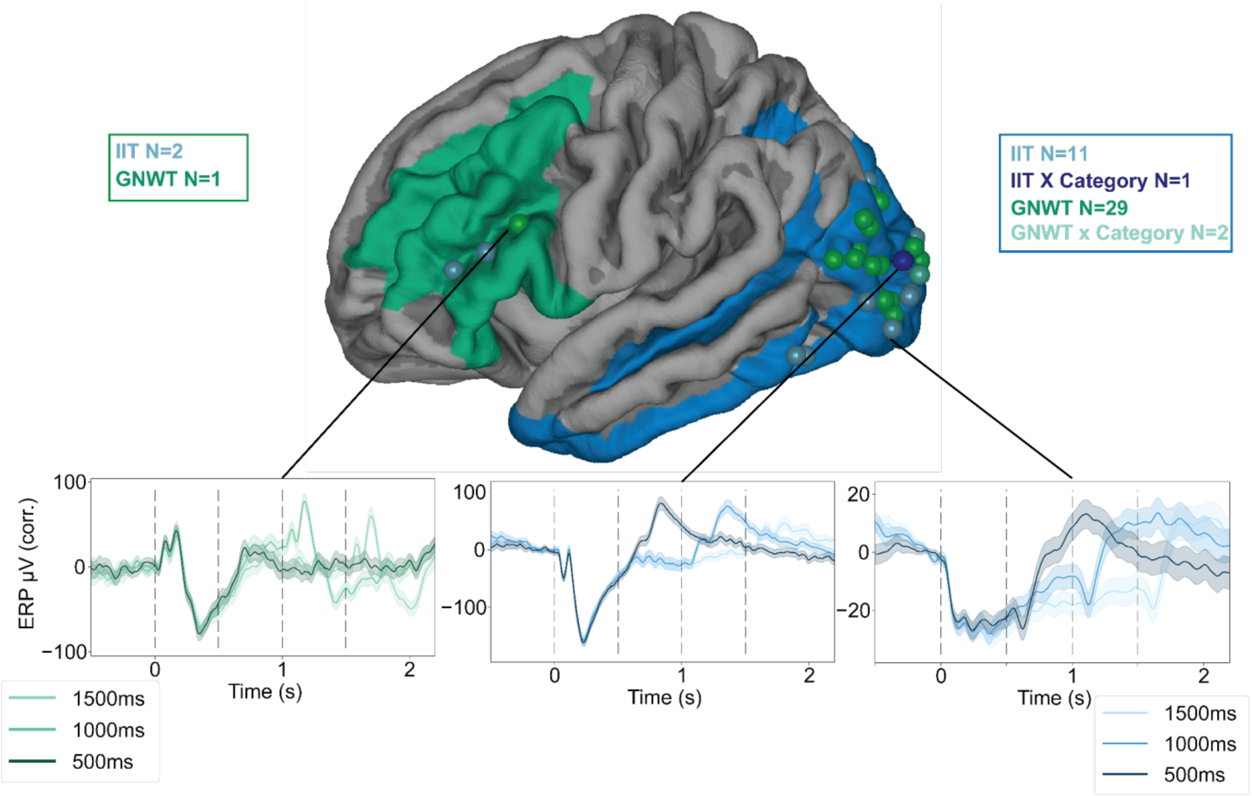

**Supplementary Figure 30***:* Results of the LMM on the ERP signal in the task irrelevant condition. The location of the electrodes found to be consistent with the theories' models are shown on the brain surface, with the prefrontal and posterior ROIs depicted in green and blue, respectively. The time series below show the average activation across trials, separately for each duration (shaded areas represent s.e.m. across trials). On the left, an electrode found to be consistent with the GNWT prediction in the prefrontal region, in the middle is an electrode consistent with IIT prediction with category interaction, on the right is an electrode consistent with IIT without a category interaction.

In the task relevant condition, more prefrontal electrodes (N=3) were found to be consistent with the GNWT model for the ERP signal, and one electrode fitted the GNWT model in the HGP. Notably however, these results might be driven by the task rather than by consciousness per se (accordingly, all theory predictions were critically tested on the task irrelevant conditions).

As for consistency across task relevance conditions, as can be seen in Supplementary Figure 31 for the HGP signal, electrodes captured by the models in the task irrelevant condition largely overlapped with those captured in the task relevant condition. The only notable exception were 12 out of the 22 electrodes captured in the task relevant condition as showing a category interaction with the IIT model; these electrodes showed no interaction in the task irrelevant condition. This result can partly be explained by the somewhat weaker signal found in the task irrelevant condition, compared with the task relevant one (see onset responsiveness, category selectivity and Representational Similarity Analysis (RSA) sections), as might be indeed expected given the task manipulation. Because testing for interaction involves sub-grouping of trials, and accordingly require stronger effect sizes to be detected, it is probable that these electrodes were not captured by the model in the task irrelevant condition due to the weaker signal.

#### iEEG: Onset responsiveness and category selectivity

Electrode implantation varies considerably across epilepsy patients, as it is dictated based on medical considerations. Thus, we first aimed at characterizing the neural responses observed across the populations of electrodes in response to the stimuli presented. This analysis was done independently from the theories' predictions.

Overall, 15.9% (558 out of 3512) of the electrodes showed responses to our stimuli: out of those, 357 showed amplitude increases with respect to baseline, while 201 showed amplitude decreases. Supplementary Figure 31 shows the electrodes found to be responsive in the task relevant and irrelevant trials. Next, we characterized the latency of neural responses across brain areas. Response latencies were the shortest and least variable in occipital cortex, while latencies in other regions were found to be more variable (see Supplementary Figure 31c). We then examined the electrodes that were located within the ROIs defined by the theories. Within the posterior ROI we observed a higher number of electrodes showing amplitude increases compared to baseline than electrodes showing amplitude decreases (141 vs. 29). In the PFC ROI, the proportion of activated to deactivated electrodes was comparable (55 vs. 59).

We then established the robustness of this result using a Bayes factor t-test (Cauchy scale factor of 0.707). We observed comparable results: 440 electrodes were found to be responsive using this method, with 95.2% of those electrodes consistent with the electrodes identified using the methods reported above.

We further characterized the selectivity of the electrodes in our data set (see methods). A total of 223 electrodes were found to be selective to a given category. Faces was the category with the most selective electrodes (108), followed by objects (78), false-fonts (31) and letters (6). On average, the selectivity strength (quantified as a d') was similar across faces (M=0.88, SD=0.66), objects (M=0.70, SD=0.36) and false-fonts (M=0.71, SD=0.28), although the strongest d' values were observed for the face-selective electrodes. 89 of the 223 category selective electrodes (40.0%) were located within the posterior ROI defined by IIT (51 face selective, 24 object selective, 13 false-fonts selective and only 1 letter selective electrodes). While fewer category selective electrodes were observed in the GNWT ROI (24/223, 10.8%), selectivity for all stimulus categories was observed (15 face selective, 6 object selective, 2 letter selective and 1 false-font selective electrodes). Most of the category selective electrodes clustered within a few ROIs (inferior frontal sulcus, 8, middle frontal gyrus, 7, triangular part of the inferior frontal gyrus, 4, opercular part of the inferior frontal gyrus, 4 on the Destrieux).

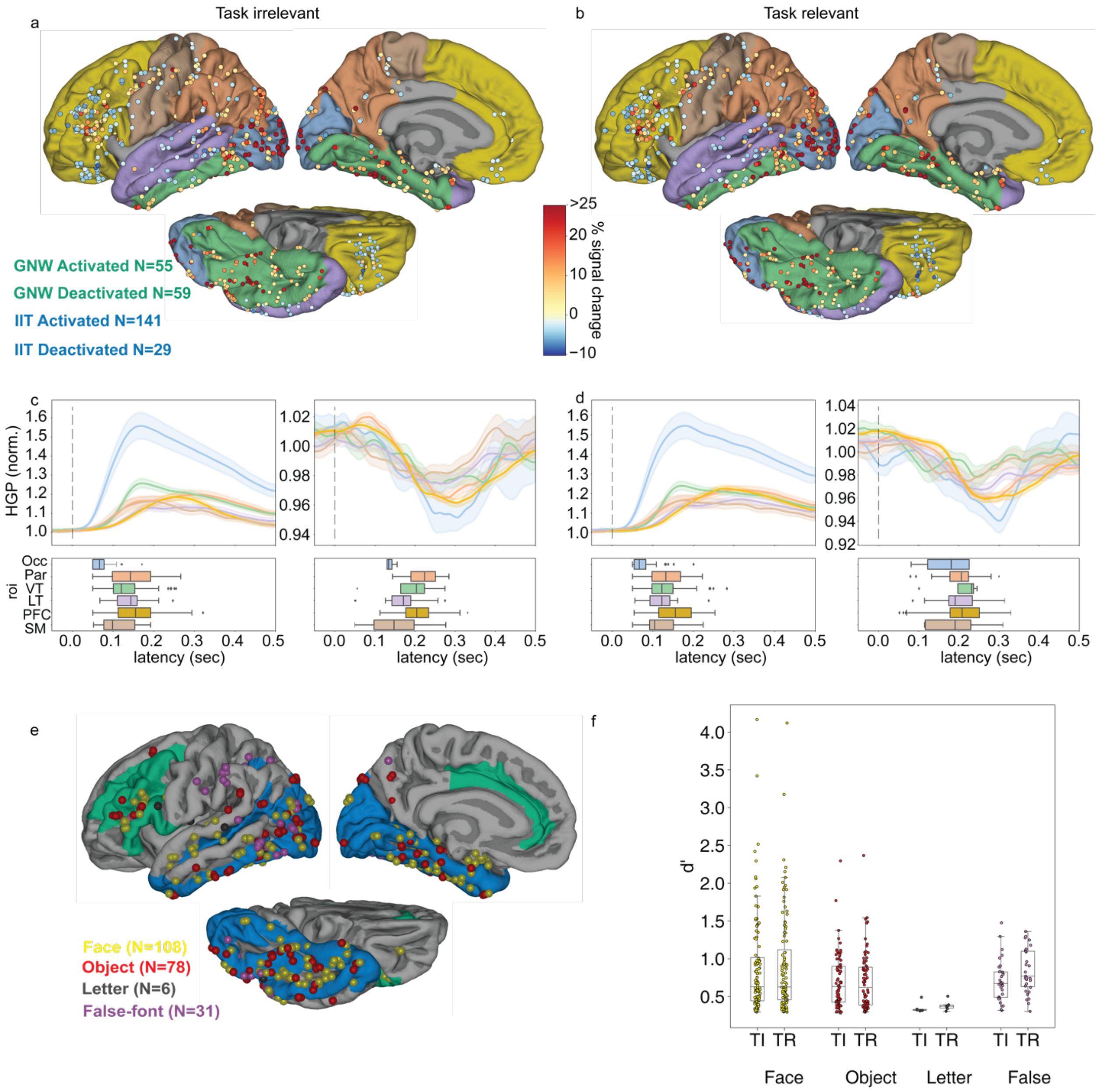

**Supplementary Figure 31.** Results of the onset responsive and category selectivity analysis (N subjects=32). **a-b.** The location of onset responsive electrodes, color coded by the percentage of signal change between baseline (-0.3-0 s) and onset window (0.05-0.35 s) in the task irrelevant (**a**) and relevant (**b**) conditions. The color on the surfaces represents anatomical ROIs (blue: occipital (occ), orange: parietal (Par), green: ventral temporal (VT), purple: lateral temporal (LT), yellow:

PFC, brown: sensorimotor (SM)). **c.** Average HGP across electrodes in the task irrelevant condition showing activation (left) and deactivation (right, shaded area represent the s.e.m. across electrodes) separately for each ROI (colors matching the surfaces) as well as the response latencies across electrodes within ROIs in the form of boxplots. **d.** Same as panel c but for the task relevant condition. **e.** Location of category selective electrodes color coded per category. The color on the surfaces represent the theory ROIs. **f.** Box plots depicting the distribution of selectivity strengths (d') for each category and task relevance condition separately (each point represents the d' of an electrode selective to the corresponding category).

#### MEG: duration tracking in the different signals

For MEG, we used LMMs to investigate the temporal patterns of gamma band (60-90 Hz) and band alpha (8-13 Hz) power, as well as event-related fields (ERFs). Akin to the iEEG section above, here too we provide results for all conditions and all preregistered signals, within the regions of interest (ROIs) defined by the theory proponents (posterior ($N_{parcels}$=15), parietal ($N_{parcels}$=1) and prefrontal ($N_{parcels}$=11) parcels). Supplementary Table 6 details the number of parcels aligned with each theoretical model according to regions of interest, task condition, and type of signal (gamma and alpha bands as well as event-related fields, ERFs). We additionally report the results for the "Late" alpha analysis, in which we tested later time windows (200 ms later than specified in the preregistration), given the expected delay of the peak alpha signal decrease relative to the peak gamma signal increase (see below for details).

| ROI | # parcels | Task relevance | Signal | IIT model | IIT x Category | GNWT model | GNWT x Category |
|---|---|---|---|---|---|---|---|
| Posterior | 15 | Irrelevant | Alpha | 1<br>FIIT = 15.48<br>p = 0.0001 | 0 | 13<br>FGNW > 14.65<br>P < 0.0002 | 1<br>FGNW x cate = 0.18<br>p = 0.91 |
| | | | "Late" alpha | 12<br>FIIT > 30.21<br>p < 0.0001 | 3<br>FIIT x cate > 2.91<br>p < 0.03 | 0 | 0 |
| | | | Gamma | 0 | 0 | 0 | 0 |
| | | | ERF | 0 | 0 | 1<br>FGNW = 43.60<br>p < 0.0001 | 0 |
| | | Relevant | Alpha | 2<br>FIIT > 16.60<br>p < 0.0001 | 0 | 12<br>FGNW > 10.19<br>p < 0.0014 | 0 |
| | | | "Late" alpha | 15<br>FIIT > 38.55<br>p < 0.0001 | 0 | 0 | 0 |
| | | | Gamma | 0 | 0 | 0 | 0 |
| | | | ERF | 0 | 0 | 3<br>FGNW > 19.11 | 0 |

| | | | | | | | |
|---|---|---|---|---|---|---|---|
| | | | | | | $p < 0.0001$ | |
| Parietal | 1 | Irrelevant | Alpha | 0 | 0 | 1<br>FGNW = 28.19<br>$p < 0.0001$ | 0 |
| | | | “Late” alpha | 1<br>FIIT > 23.80<br>$p = 0.0001$ | 0 | 0 | 0 |
| | | | Gamma | 0 | 0 | 0 | 0 |
| | | | ERF | 0 | 0 | 0 | 0 |
| | | Relevant | Alpha | 0 | 0 | 1<br>FGNW = 20.44<br>$p < 0.0001$ | 0 |
| | | | “Late” alpha | 1<br>FIIT = 12.66<br>$p = 0.0004$ | 0 | 0 | 0 |
| | | | Gamma | 0 | 0 | 0 | 0 |
| | | | ERF | 0 | 0 | 0 | 0 |
| Prefrontal | 11 | Irrelevant | Alpha | 0 | 0 | 2<br>FGNW = 19.62<br>$p < 0.0001$ | 0 |
| | | | “Late” alpha | 2<br>FIIT > 22.52<br>$p < 0.0001$ | 0 | 0 | 0 |
| | | | Gamma | 0 | 0 | 0 | 0 |
| | | | ERF | 0 | 0 | 0 | 0 |
| | | Relevant | Alpha | 0 | 0 | 0 | 0 |
| | | | “Late” alpha | 2<br>FIIT > 24.51<br>$p < 0.0001$ | 0 | 0 | 0 |
| | | | Gamma | 0 | 0 | 0 | 0 |
| | | | ERF | 0 | 0 | 0 | 0 |

**Supplementary Table 6**. Number of posterior, parietal and prefrontal parcels with a significant fit to the theories’ models in the LMM analysis, separately for the alpha, “late” alpha, gamma and ERF analyses.

In the task irrelevant condition, the results provided support for GNWT, with two prefrontal parcels (Anterior and Middle-Anterior Cingulate Cortex) showing the GNWT-expected pattern of alpha band activity. However, this finding was not supported by the alpha band analysis in the late time bins. More convincing evidence was found in support of IIT, specifically in the two alpha band analyses. The observed effects were category-specific. The observed effects were specific to a particular category in three posterior parcels (Cuneus, Superior Occipital Gyrus, Superior and Transverse Occipital Sulcus), specificity predicted by IIT. The results from the gamma band and ERF analyses did not provide any support for either theory.

In the task relevant condition, neither of the theories' predictions were fully supported in the alpha band. Specifically, none of the prefrontal parcels showed the pattern of activity predicted by GNWT. At the same time, we did not observe the category-selective effect predicted by IIT in any of the posterior parcels. As in the irrelevant condition, the results from the gamma band and ERF analyses did not provide any support for either theory. More detailed information regarding the specific analyses on the alpha and ERF signals are provided below.

#### Gamma power

The LMM analysis on the task-irrelevant condition showed that none of the theory-based models provided a good fit to the data (Supplementary Table 6a). We did not detect any sustained gamma band activity in posterior cortex, as predicted by IIT, neither any phasic onset and offset response in PFC, as predicted by GNW (see Supplementary Table 6b for an example of a posterior and PFC parcel). Gamma band power was strong in posterior areas (see Supplementary Figure 32), but it is possible that the null results were due to small signal amplitudes.

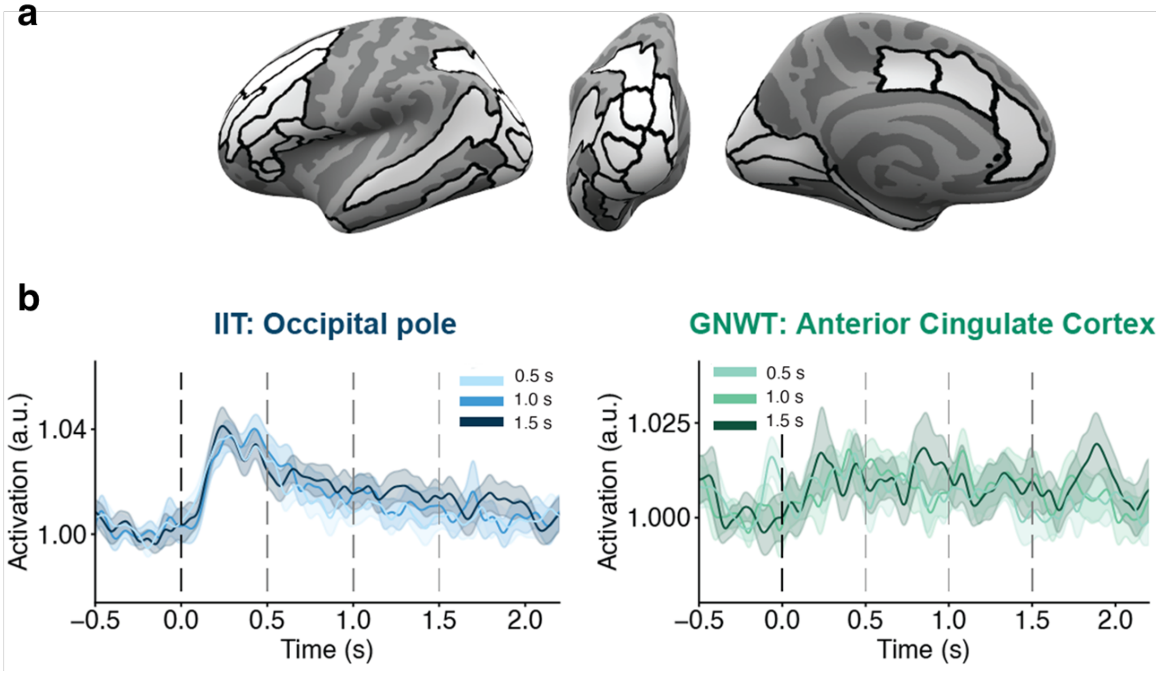

**Supplementary Figure 32**. A. Parcels denoting ROIs showing gamma activity consistent with none of the temporal patterns predicted by the theories (white). B. Left: averaged gamma activity in the occipital pole for the three stimulus durations (shades of blue), showing a pattern of activity not in line with IIT's prediction. Right: averaged gamma activity in the anterior cingulate cortex for the three stimulus durations (shades of green), showing a pattern of activity not in line with GNWT's prediction.

##### Alpha power

Alpha band activity can be reliably detected in MEG and it is held to be anti-correlated with the gamma signal. To make sure this is indeed the case also in our data, we explored the time course of the two rhythms in the task irrelevant condition, averaged across durations. Such anti-correlation was indeed found in our data (Supplementary Figure 33), with the alpha and gamma band activities both localized in posterior areas and exhibiting an inverse correlation (i.e., while gamma activity increases, alpha activity decreases).

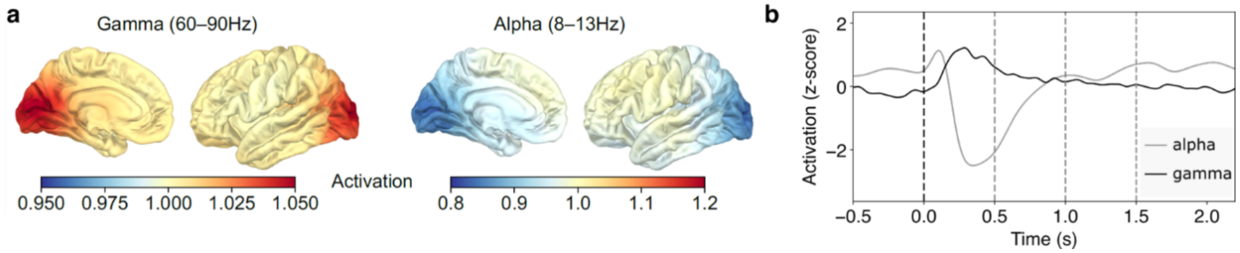

**Supplementary Figure 33**. a. Source localization maps for the gamma band (60-90 Hz, left) and alpha band (8-12 Hz, right) activity. b. Time course of the gamma and alpha band activity in occipital cortex for the task irrelevant condition. The data are averaged across all stimulus durations. The vertical axis represents the z-scored power in the two frequency bands. Gamma and alpha band activities exhibit an anti-correlated relationship.

The LMM analysis on the task-irrelevant condition showed two prefrontal parcels (Anterior and Middle-Anterior Cingulate Cortex) and a posterior parcel (Occipital Pole) matching GNWT's and IIT's predictions, respectively (Supplementary Figure 34a). In posterior cortex, alpha power decreased in a sustained manner scaling with stimulus duration (with a further phasic decrease at stimulus offset) in the occipital pole, in line with IIT predictions. In PFC, the anterior and middle-anterior cingulate cortex showed the GNWT temporal profile, with a phasic decrease in the alpha power in the 300-500 ms interval after stimulus offset (Supplementary Figure 34b). Supplementary Table 7 provides the Bayesian Information Criteria (BICs) for all tested models for each of these parcels.

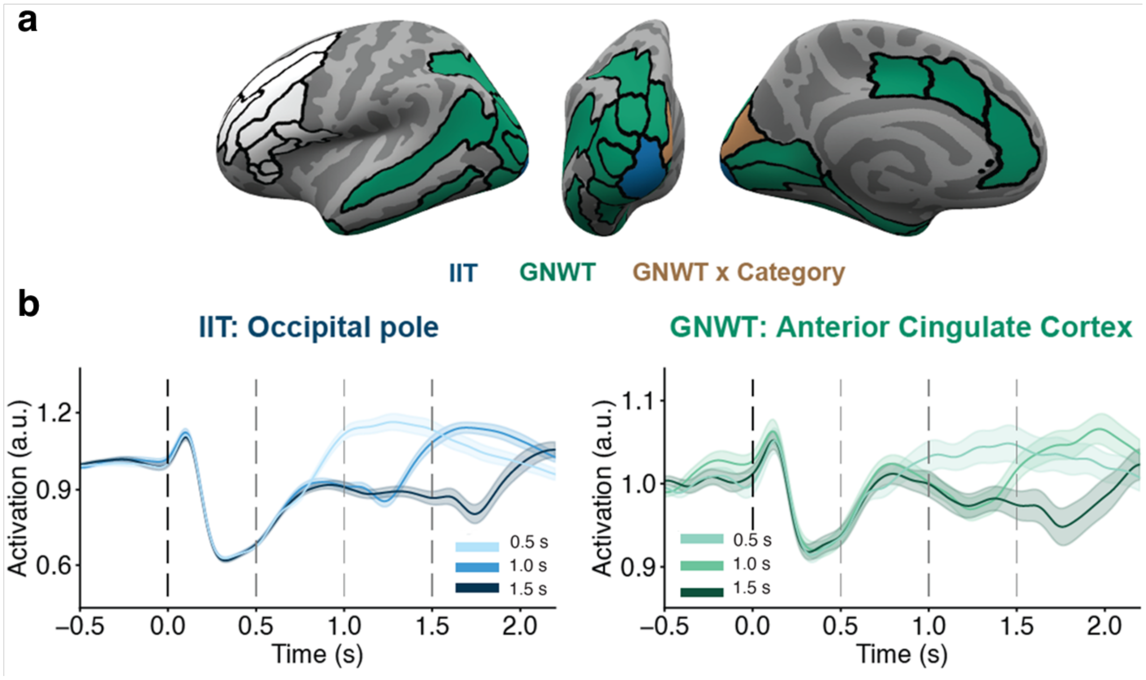

**Supplementary Figure 34**. a. Parcels denoting ROIs showing alpha activity consistent with the sustained duration tracking predicted by IIT (blue), the phasic onset & offset duration tracking predicted by GNWT (green), and areas showing none of the temporal patterns predicted by the theories (white). b. Left: averaged alpha activity in the occipital pole for the three stimulus durations (shades of blue), best fitted by the IIT model. Right: averaged gamma activity in the anterior cingulate cortex for the three stimulus durations (shades of green), best fitted by the GNWT model.

| Model | Anterior Cingulate Cortex | Middle-Anterior Cingulate Cortex | Occipital Pole |
|---|---|---|---|
| Null | -2428.03* | -2307.79* | -593.34* |
| Time window | -2415.60* | -2297.43* | -625.27* |
| Duration | -2461.89* | -2336.43* | -840.26* |
| Time window + Duration | -2449.52* | -2326.17* | -878.06* |
| Time window + Duration + IIT | -2446.19* | -2324.87* | -885.72* |

| | | | |
|---|---|---|---|
| Time window + Duration + GNWT | -2465.26* | -2337.96* | -876.21* |
| Time window + Duration + IIT x Category | -2419.60* | -2302.07* | -874.86* |
| Time window + Duration + GNWT x Category | -2437.61* | -2308.88* | -863.20* |

**Supplementary Table 7**. BIC values from the linear mixed model on the alpha activity for each model and parcel showing theory-predicted activity. Stars indicate that the fit of the model converged significantly (p-value < .05).

Given that alpha is a slow rhythm, it might be that the predicted patterns should actually be found later in time (compared to the gamma patterns for which the time windows were optimized). We accordingly conducted a control analysis using later time bins. Specifically, we analyzed the activity in the time bins 1.0 -1.2 s, 1.5-1.7 s and 2.0-2.2 s. In the posterior ROI, all the parcels showed an alpha activity in line with IIT.s prediction, with two parcels (Cuneus and Superior Occipital Gyrus) also showing category-selective responses. In contrast, none of the parcels in PFC showed the pattern of activity predicted by GNWT. While these findings confirm the sustained posterior activity previously associated with alpha, providing further support for IIT's prediction, they raise doubts about the phasic alpha band response in the PFC.

##### ERF signals

ERFs were calculated for each parcel in both the prefrontal and posterior ROIs (in line with the main analyses reported in the main text). The results provided no evidence for either of the theories. In the prefrontal ROIs, none of the theories' models outperformed the non-theoretical models. In the posterior ROIs, the GNWT's model best fitted the ERFs in the Inferior Occipital Cortex parcel, and the IIT's model did not outperform the other models in any of the remaining parcels (See again Supplementary Table 6 above).

#### Exploratory analyses: duration predictions

Several control analyses were conducted to exclude possible confounds, to better understand the obtained results we found, or to provide the theories with the best chance possible of confirming their predictions. These control analyses are described below.

#### iEEG: Exploratory decoding analysis, with unrestricted temporal profiles and time windows

The models presented above test the theories predictions very strictly, as they only investigate activation in pre-specified time-windows. To explore if the data includes any evidence for duration tracking that is not restricted to these time windows, we complemented our planned analysis with an additional analysis that searched for these patterns using decoding along the entire signal. We accordingly trained an SVM classifier to decode stimulus duration for each electrode from 0.5 to 2 s of stimulus onset (excluding the first 0.5 s as indiscriminate of stimuli duration), using time points as features (after 0.200 s non-overlapping moving average of single trials activation). The SVM was run separately for each stimulus category (faces, objects, letters, false fonts) and task manipulation (task relevant and irrelevant). No FDR correction was applied. Electrodes were considered to track duration if the decoding accuracy was significantly above chance (label shuffle permutation test) in both task relevance conditions for at least one of the stimulus categories (see Supplementary Table 8). This approach also allowed us to investigate the robustness of the LMM in identifying the predicted patterns as well as capture any other activation patterns associated with duration. The results are depicted in Supplementary Figure 35 (alongside the results of the original LMM analysis, and the following sustained activity tracking analysis, described below).

| ROI | # electrodes | Across categories | Faces | Objects | Letters | False fonts |
|---|---|---|---|---|---|---|
| Posterior | 657 | 47 | 34 | 31 | 26 | 28 |
| Prefrontal | 655 | 4 | 2 | 3 | 1 | 1 |

**Supplementary Table 8**: Count of electrodes for which duration decoding was found, separately for prefrontal and posterior ROIs (N subjects=31). Electrodes showing any duration tracking, for at least one of the categories, are reported in the column ‘Across categories’. The other columns describe the number of electrodes showing duration tracking separately for each of the categories. Notably, there was substantial overlap (i.e., most electrodes responded to more than one category).

In this exploratory analysis, we asked two questions. First, how many of the electrodes captured by the LMM analysis were also detected in this exploratory decoding analysis. Second, how many of the electrodes detected in the exploratory analysis show a pattern that matches either the IIT or the GNWT predictions (based on visual inspection of the results). As Supplementary Figure 35 shows (panels a, b and d), the first question yielded a strong result, validating the two analyses, as substantial overlap was found between them, with all electrodes detected by the LMM approach also found in the decoding approach, except for one. As for the second question, with this more liberal approach we were only able to detect one electrode that showed an onset and offset response consistent with GNWT’s predictions, yet at earlier time windows than expected (reported in the main text; see Figure 3d). The three other electrodes picked up by this analysis did not show the expected onset & offset response.

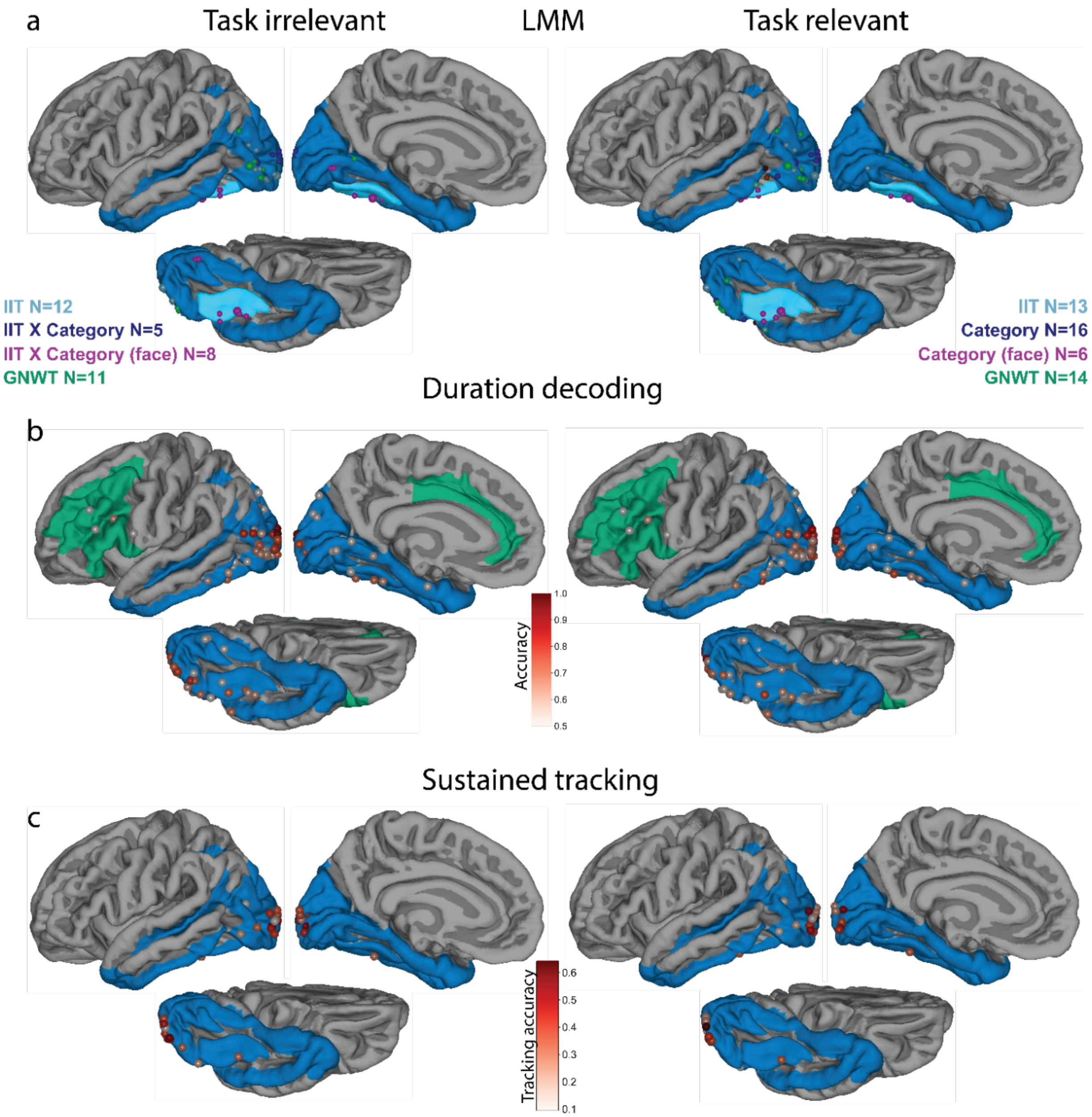

Methods overlap

d

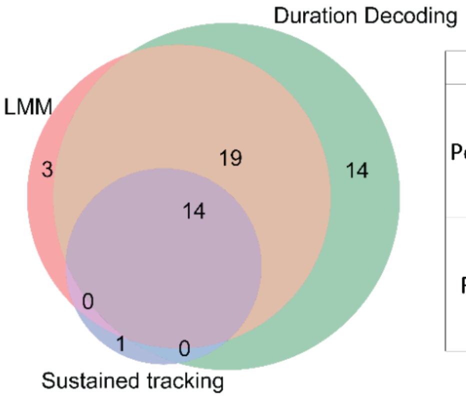

| | LMM | Duration decoding | Sustained tracking |
|---|---|---|---|
| Posterior | IIT: 12 | 12 (12) | 8 (11) |
| | IIT x Category: 13 | 13 (13) | 5 (5) |
| | GNW: 11 | 8 (10) | 1 (3) |
| | None | 14 | 1 |
| Frontal | IIT: 0 | 0 | |
| | IIT x Category: 0 | 0 | |
| | GNW: 0 | 0 | |
| | None | 4 | |

**Supplementary Figure 35**. Comparison of the results of the different analyses used to identify electrodes showing activation patterns associated with stimuli duration, separately for the task irrelevant (left) and relevant (right) conditions (prefrontal ROI, N subjects=31; posterior ROI, N subjects=31). **a.** Location of electrodes identified by the LMM models (reported in the main text) as showing activation patterns consistent with IIT (light blue), GNWT (green), IIT with category interaction (dark blue), on which face selective electrodes showing sustained activation in a category dependent manner are marked in purple. The number next to the legend corresponds to the total number of electrodes found for each analysis. **b.** Location of electrodes with above chance duration decoding in at least one of the 4 stimuli categories, color coded by the highest significant accuracy observed across the 4 stimuli categories. **c.** Location of electrodes with significant sustained activation (as determined by the sustained tracking method) for at least one of the stimulus categories, color coded by the tracking accuracy (i.e. proportion of trials for which activation was sustained for as long as the stimulus was presented on the screen, +/- 0.15 s). **d.** Overlaps across the three methods for the task irrelevant trials only. The Venn diagram (left) represents the total number captured by each method and the overlap across those. The table (right) shows the breakdown of the electrodes that were identified in the different theoretical models in the LMM analysis (left column), according to their overlap with the other methods (duration tracking, middle column; sustained activity, right column). In the latter columns, the numbers correspond to the count of LMM-detected electrodes that were also found to be significant by each method in both task conditions (alpha=0.05) while the number between parentheses corresponds to the counts of electrodes found to be significant in the task irrelevant condition (no correction). The row "none" describes the number of electrodes detected by the two analyses, which were not picked up by the LMM. Note that the sustained tracking method was not applied on the prefrontal ROIs, hence these cells are empty.

#### iEEG: Onset/offset analysis

The second control analysis was again driven by the lack of prefrontal electrodes showing the GNWT predicted pattern of both onset and offset ignitions. This failure could stem from three possible findings: lack of electrodes showing an onset response, lack of electrodes showing an offset response, and/or a lack of electrodes showing both an onset and an offset response. In addition, in the LMM approach, activation was averaged over broad time windows (0.2 s). If the offset responses are very short-lived in the prefrontal ROI, they might have been missed by our analysis. To disentangle between these options, we performed a time-resolved analysis. Specifically, we ran a sliding t-test across all stimuli durations from 0.3-0.5 s (against baseline from -0.2 to 0 s) following stimulus onset and offset separately. Activation was considered to be significant when a minimum of 0.02 s consecutive samples were below $p<0.05$. Only electrodes within the GNWT-defined region of interest were investigated. The results are summarized in Supplementary Table 9.

| # electrodes | Significant window | Across categories | Faces | Objects | Letters | False-fonts |
|---|---|---|---|---|---|---|
| 655 | Stimulus onset | 9 | 2 | 3 | 2 | 7 |
| | Stimulus offset | 0 | 0 | 0 | 0 | 0 |
| | Both | 0 | 0 | 0 | 0 | 0 |

**Supplementary Table 9**: Counts of electrodes showing increased activation following stimulus onset, offset or both in the prefrontal ROI (N subjects=31). This analysis was performed for each task and category separately. The reported counts in the across categories column refer to the electrodes with activation significantly above baseline in both task conditions within at least one of the stimulus categories. The reported counts in the faces, objects, letters and false-fonts column refer to the counts separately for each category. With this analysis, we identified nine electrodes that showed an onset response between 0.3 and 0.5 s, while no electrode showed either an offset response alone (at the respective time following stimulus offset), or both an onset and offset response.

This result demonstrates first that our analyses are sensitive enough to capture the relevant responses when they occur, and second that onset responses are indeed found in PFC ROIs within the predicted time window (though in a relatively small subset of the electrodes). This pattern of results suggests that the reason the main preregistered analyses failed to support the GNWT model was the lack of offset responses.

#### iEEG: Sustained duration tracking

Finally, the third control analysis was aimed at testing for sustained activity in the electrodes identified by the modeling approach as matching the patterns predicted by IIT. Notably, the preregistered modeling approach was designed to maximize discriminability between both theories by investigating only specific time windows following stimulus onset. Thus, it is possible that the identified electrodes do not show sustained activation, but rather increased activation only in the investigated time windows.

To control for this possibility, we applied the method developed by Gerber and colleagues[6] .This method is aimed at determining how long a stimulus was presented in single trials by isolating the time point at which activation drops below a threshold defined as the median between the average baseline (-0.5 to -0.2 s) activation and post-stimulus activation (1.2 to 1.5 s) of the longest stimuli (1.5 s) only. Activation in a given trial was considered to accurately track duration if the predicted duration was within +/- 0.15 s of stimulus offset. For a given electrode, the proportion of accurately predicted trial duration was computed and compared to a null distribution of duration tracking accuracy obtained by durations label shuffles. This analysis was performed separately for each task condition and stimulus category on the 1.0 and 1.5 s trials. Electrodes were considered to track duration if duration was correctly identified in both task conditions, for at least one of the stimuli categories. This method was applied to electrodes located in the IIT defined region of interest using the HG signal, and the results are summarized in Supplementary Table 10. Using this method, we found 15 electrodes showing sustained activity, out of the 25 electrodes found using the modeling approach.

| # electrodes | Across categories | Faces | Objects | Letters | False fonts |
|---|---|---|---|---|---|
| 657 | 15 | 10 | 10 | 9 | 5 |

**Supplementary Table 10**. Counts of electrodes showing duration tracking in the posterior ROI (N subjects=31). We performed the analyses for each task and category separately. The reported counts in the across categories column refer to the electrodes in which duration was identified significantly above chance in both task conditions within at least one of the categories. The reported counts in the faces, objects, letters and false-fonts column refer to the counts separately for each category.

#### MEG: gamma power in unified ROIs

As described previously, for the MEG data, we did not find any meaningful results with respect to the theories' predictions in the gamma range when considering the individual parcels within the prefrontal and posterior ROIs. We further tested if the predictions are borne out by the data when inspecting the entire posterior/prefrontal ROIs without dividing them into parcels. Here, we did find an early increase in gamma power with the onset of the visual stimuli in the posterior ROI, but not in the prefrontal one (Supplementary Figure 36). However, this posterior gamma activity was not sustained with respect to the duration of the stimuli, contrary to the prediction by IIT. These observations were confirmed by the results of the LMM analysis, which did not find support for the IIT models in the posterior ROI, nor for the GNWT models in the prefrontal ROI (Supplementary Table 11).

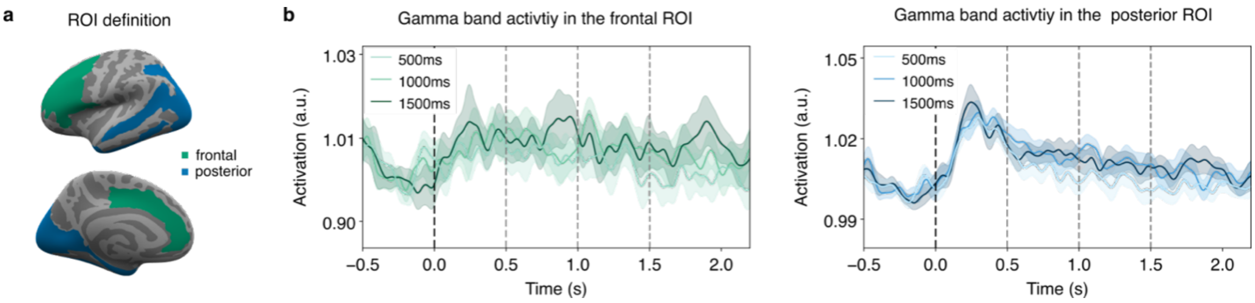

**Supplementary Figure 36**. Modulation of gamma band (60-90 Hz) power in the entire prefrontal (left) and posterior (right) ROIs. **a.** Brain surface plot showing the definition of the entire prefrontal and posterior ROIs in lateral (top) and medial (bottom) views. **b.** Time course of the gamma band activity in the prefrontal (left, green) and posterior (right, blue) ROIs for each stimulus duration, averaged across participants. Vertical dotted lines represent stimulus onset (t=0 s) and offset (t=0.5, 1.0 and 1.5 s). Shaded areas denote the confidence intervals. In the posterior ROI, we found an early increase in gamma power with the onset of the visual stimuli, which was however not sustained with respect to the duration of the stimuli.

| Model | Prefrontal ROI | Posterior ROI |
|---|---|---|
| Null | -8634.19* | -9131.94* |
| Time window | -8620.56* | -9125.84* |
| Duration | -8637.89* | -9138.72* |
| Time window + Duration | -8624.27* | -9132.71* |
| Time window + Duration + IIT | -8620.26* | -9125.63* |
| Time window + Duration + GNW | -8618.65* | -9126.16* |
| Time window + Duration + IIT x Category | -8577.34* | -9086.74* |
| Time window + Duration + GNWT x Category | -8575.98* | -9088.87* |

**Supplementary Table 11**. BIC values from the linear mixed model on the gamma activity for each model and ROI. Stars indicate that the fit of the model converged significantly (p-value < .05).

#### MEG: Alpha power in unified ROIs

Similarly, we tested the predictions on the entire posterior/prefrontal ROIs using alpha power, and observed an initial reduction in activity followed by a sustained but smaller decrease throughout the duration of the stimuli (Supplementary Figure 37). These observations were subsequently analyzed using a LMM. In the posterior ROI, the response best fitted the GNWT model, different from IIT's predictions. For the prefrontal ROI, the models including the GNWT predictors outperformed all other models (Supplementary Table 12). Taken together, these results support the hypotheses proposed by GNWT and those proposed by IIT.

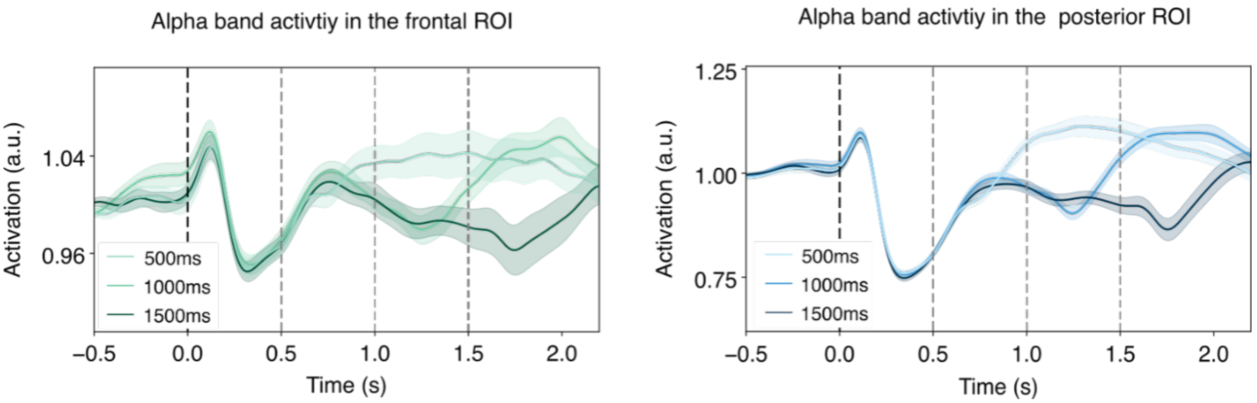

**Supplementary Figure 37**. Alpha band (8-13 Hz) power in the combined prefrontal (left, green) and the combined posterior (right, blue) ROIs, averaged across participants for each stimulus duration. Alpha power in prefrontal and posterior cortices was reduced following an initial increase after stimulus onset, and then increased back to (or above) baseline levels later in time depending on stimulus duration. Vertical dotted lines represent stimulus onset and offset. Shaded areas denote the confidence intervals.

| Model | Prefrontal ROI | Posterior ROI |
|---|---|---|
| Null | -3260.23* | -1814.58* |
| Time window | -3250.18* | -1817.67* |
| Duration | -3301.51* | -2017.08* |
| Time window + Duration | -3291.59* | -2022.05* |
| Time window + Duration + IIT | -3287.74* | -2018.97* |
| Time window + Duration + GNW | -3303.48* | -2074.43* |
| Time window + Duration + IIT x Category | -3262.99* | -2003.74* |
| Time window + Duration + GNWT x Category | -3276.18* | -2055.83* |

**Supplementary Table 12**. BIC values from the linear mixed model on the alpha activity, separate per models and ROIs. Stars indicate that the fit of the model converged significantly (p-value < .05).

#### MEG: ERFs in unified ROIs

When inspecting the combined ROIs, both the prefrontal and posterior ROIs showed an initial evoked response locked to the stimulus onset (Supplementary Figure 38). We also observed an offset response, yet it was mainly visible in the posterior ROI, where neither of the theories predicted such a pattern. A LMM analysis on the ERFs was then performed on each ROI. In both cases, none of the theory models outperformed the time window model (see Supplementary Table 13).

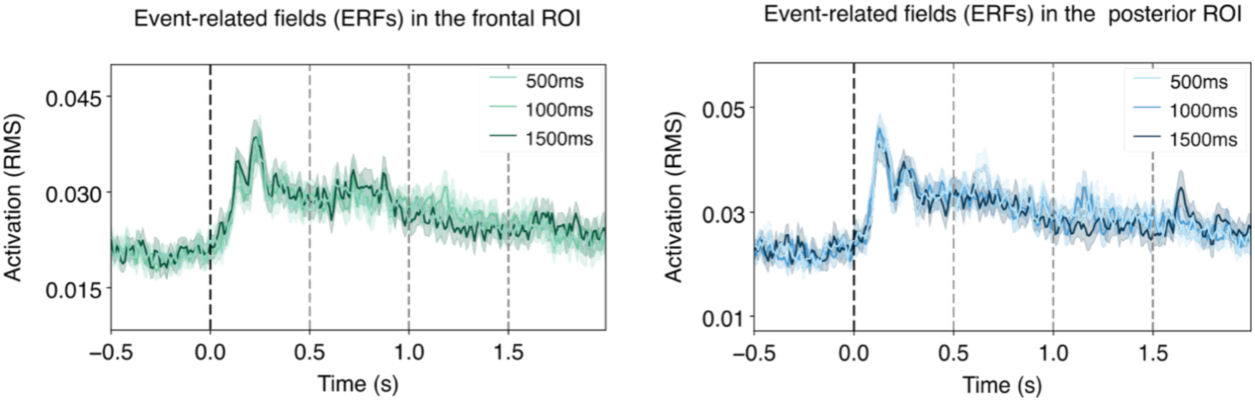

**Supplementary Figure 38**. Modulation of the ERF response in the combined prefrontal (left) and combined posterior (right) ROIs, averaged across participants for each stimulus duration (Root Mean Square; RMS). We found an early evoked response at the onset of the visual stimuli in both the prefrontal and posterior ROIs and a smaller offset response limited to the posterior ROI. Vertical dotted lines represent stimulus onset (black) and offset (gray, for the three different offsets). Shaded areas denote the confidence intervals.

| Model | Prefrontal ROI | Posterior ROI |
|---|---|---|
| Null | -14128.76* | -13689.17* |
| Time window | -14223.10* | -13726.36* |
| Duration | -14115.14* | -13673.99* |

| | | |
|---|---|---|
| Time window + Duration | -14209.57* | -13711.19* |
| Time window + Duration + IIT | -14205.25* | -13706.36* |
| Time window + Duration + GNW | -14203.24* | -13705.87* |
| Time window + Duration + IIT x Category | -14170.60* | -13669.54* |
| Time window + Duration + GNWT x Category | -14168.86* | -13668.17* |

**Supplementary Table 13.** BIC values from the linear mixed model on the ERFs for each model and ROI. Stars indicate that the fit of the model converged significantly (p-value < .05).

#### MEG: Alpha power using delayed time-windows on the unified ROIs

Given that alpha is a slow rhythm, it might be that the predicted patterns should actually be found later in time (compared to the gamma patterns). We accordingly conducted a control analysis using later time bins. Specifically, we analysed the activity in the time bins 1.0 -1.2 s, 1.5-1.7 s and 2.0-2.2 s. The IIT's model outperformed all other models in both posterior and in the prefrontal ROI. Notably however, this model was not category selective (as opposed to the content-selective IIT x Category model), contrary to IIT's predictions (Supplementary Table 14).

| Model | Prefrontal ROI | Posterior ROI |
|---|---|---|
| Null | -3482.61* | -1976.74* |
| Time window | -3473.54* | -1987.24* |
| Duration | -3524.23* | -2173.01* |
| Time window + Duration | -3515.32* | -2186.06* |
| Time window + Duration + IIT | -3525.33* | -2354.12* |
| Time window + Duration + GNWT | -3511.91* | -2266.14* |

| | | |
|---|---|---|
| Time window + Duration + IIT x Category | -3493.69* | -2341.51* |
| Time window + Duration + GNWT x Category | -3478.20* | -2248.88* |

**Supplementary Table 14**. BIC values from the linear mixed model on the alpha activity for later time bins, separate per models and ROIs. Stars indicate that the fit of the model converged significantly (p-value < .05).

#### MEG: onset/offset analysis

To further investigate alpha band activity in the different parcels in posterior cortex and PFC, we conducted an additional control analysis by comparing the power at either 0.4 or 0.6 s following stimulus onset and offset separately against a baseline at -0.25 s relative to stimulus onset and offset, respectively. These two specific time points were chosen to test both the preregistered and late alpha time bins while attempting to minimize the temporal smoothing of the wavelet analysis which used a 0.5 s gaussian window (i.e., 0.25 s before and after the time point). The baseline before stimulus onset was not used for the offset analysis in an attempt to minimize the contribution of the sustained decreased in alpha activity due to sustained tracking. To increase SNR, the power at each specific time was computed by averaging the activity from -1 to +1 ms, providing a total of three data points. Furthermore, as the onset and offset activity are confounded in the 0.5-s duration, that condition was excluded from the analysis. For this analysis 1.0 and 1.5 s trials from the task irrelevant condition were combined. We considered alpha deactivation to be significant if the p-value in the t-test remained significant after FDR correction over each contrast i.e., onset and offset separately.

In the task irrelevant condition, the onset/offset analysis yielded a significant decrease in alpha band power at stimulus onset in all PFC parcels in the early time.  In the late time, the alpha decrease was detected only in the anterior cingulate cortex (Supplementary Figure 39) and orbital inferior frontal gyrus in the late time. All posterior parcels showed an onset response in both early and late time (Supplementary Figure 39). The offset response was significant in 8 out of the 11 PFC parcels in the early time. In the late time window, none of the parcels showed a significant offset response. Similarly, no offset response was detected in the late time in any of the posterior parcels (Supplementary Figure 39), while it was detected in the intraparietal sulcus, inferior temporal gyrus and temporal pole in the early time analysis.

In the task relevant condition, we observed a significant onset response in all posterior and PFC parcels, in both the early and late time periods. However, none of the parcels showed a significant offset response at any time point.

To address the variability in activity during the stimulus presentation, which was used as a baseline in the offset analysis, we conducted an additional control analysis on the task irrelevant condition where an offset response in PFC was detected. We calculated the difference between the alpha power value at each time point and the value estimated at the preceding time point (N-1). This method eliminated any sustained modulation during stimulus presentation, thus making our baseline activity more reliable. All other aspects of the analysis remained the same as in the previous onset-offset analysis. The results demonstrated that once the variability in the sustained response was eliminated, none of the posterior or PFC areas exhibited a significant onset or offset response at 0.4 and 0.6 s, which a phasic offset response occurring in an earlier time period (< 0.25 s) in both posterior cortex and PFC (Supplementary Figure 39).

In summary, the onset/offset analysis shows a phasic offset response in PFC, as well as in posterior cortex, at the preregistered time. These results align with the results of the LMM analysis providing evidence in favour of GNW's predictions. However, the results of the late time analysis - which is considered to take into account the slow dynamic of the alpha band activity - do not fully support this conclusion, as most of the PFC parcels do not show a significant offset response during this period. Moreover, we did not observe an offset response in PFC when analysing the task relevant condition, nor when controlling for sustained effects in the baseline time period. Finally, given the similarity with the results in posterior cortex, the fact that the offset response in posterior cortex was significantly larger in magnitude (see Supplementary Figure 39 for a comparison), and the known difficulties to record activity from deep and medial (frontal) sources, we cannot rule out the possibility that the effect in PFC stems from leakage from posterior sources (see Extended Data Figure 5b).

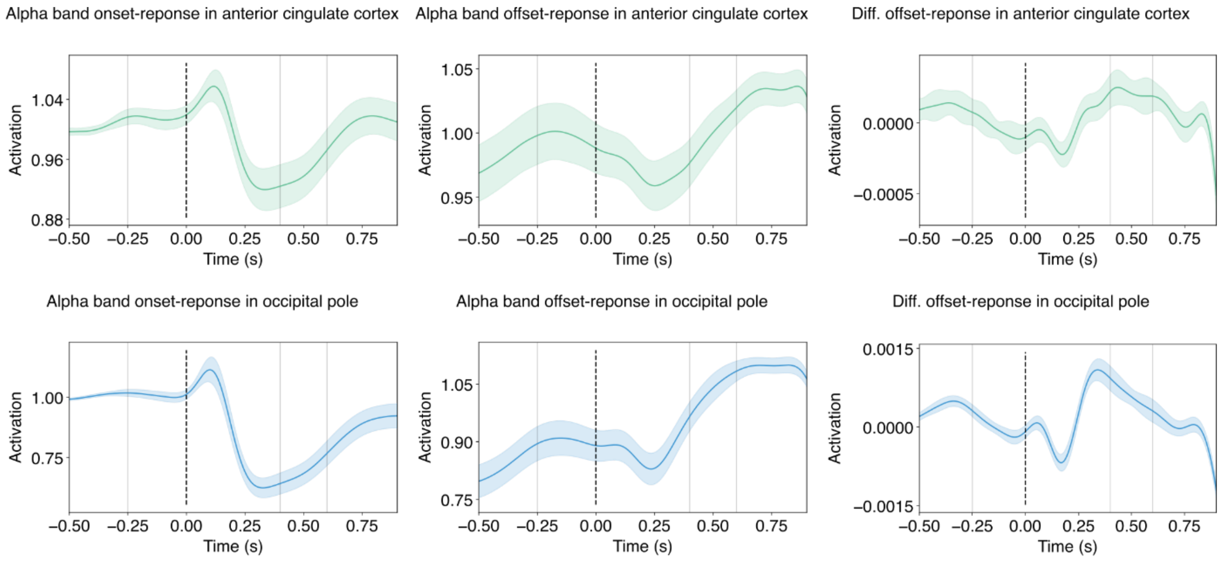

**Supplementary Figure 39**. Results of the onset/offset analysis performed on the alpha band activity in the task irrelevant condition. The top row of the figure presents the alpha band activity in the anterior cingulate cortex locked to stimulus onset (left) and stimulus offset (middle), as well as the offset response after removing the sustained activity (right). The bottom row presents the same plots for the alpha band activity in the occipital pole. The solid vertical lines represent the baseline (-0.25 s), early (0.4 s) and late activity (0.6 s) time points used in the analysis.

#### MEG: Task relevant (non-target) analysis

To examine if the pattern of results might be different when the stimuli are task relevant, we repeated the LMM analysis on the gamma band, alpha band and ERF signals, for each prefrontal and posterior parcel. None of the parcels showed the gamma activity pattern predicted by either of the two theories. As for the alpha band, the GNWT model best fitted the activity in the same parietal parcel where it was found best for the task irrelevant condition (Middle-Posterior Cingulate Cortex), as well as in twelve of the posterior parcels (see Supplementary Figure 40). The non-content selective IIT model best fitted the alpha signal in two posterior parcels (Occipital Pole, Lingual part of the Medial Occipito-Temporal gyrus). When examining the alpha band activity in late time bins, the non-content selective IIT model provided the best fit in the parietal Middle-Posterior Cingulate Cortex, and the frontal Anterior Cingulate Cortex, in the Inferior Orbital Gyrus and in all of the posterior parcels. Finally, while none of the theories' models best fitted the ERF signal in the prefrontal ROIs, the GNWT model outperformed all the other models in three posterior parcels (Cuneus, Lateral Occipito-Temporal Gyrus, Middle Occipital and Lunatus Sulcus).

The analysis on task relevant trials was also performed on the combined ROIs. In the gamma band analysis, none of the theory models outperformed the null model in the prefrontal ROIs and the duration model in the posterior ROIs, in contrast with the theories' predictions. When considering the alpha band activity, none of the theory models fit the response better than the duration model in the prefrontal ROIs, different from GNWT's predictions; in the posterior ROIs, the GNWT model outperformed the other models, against IIT's predictions. The ERF analysis results were consistent with the other results, with none of the theories' models outperforming the time window model in neither of the ROIs. Altogether, these results showed little evidence in favor of the theories' predictions.

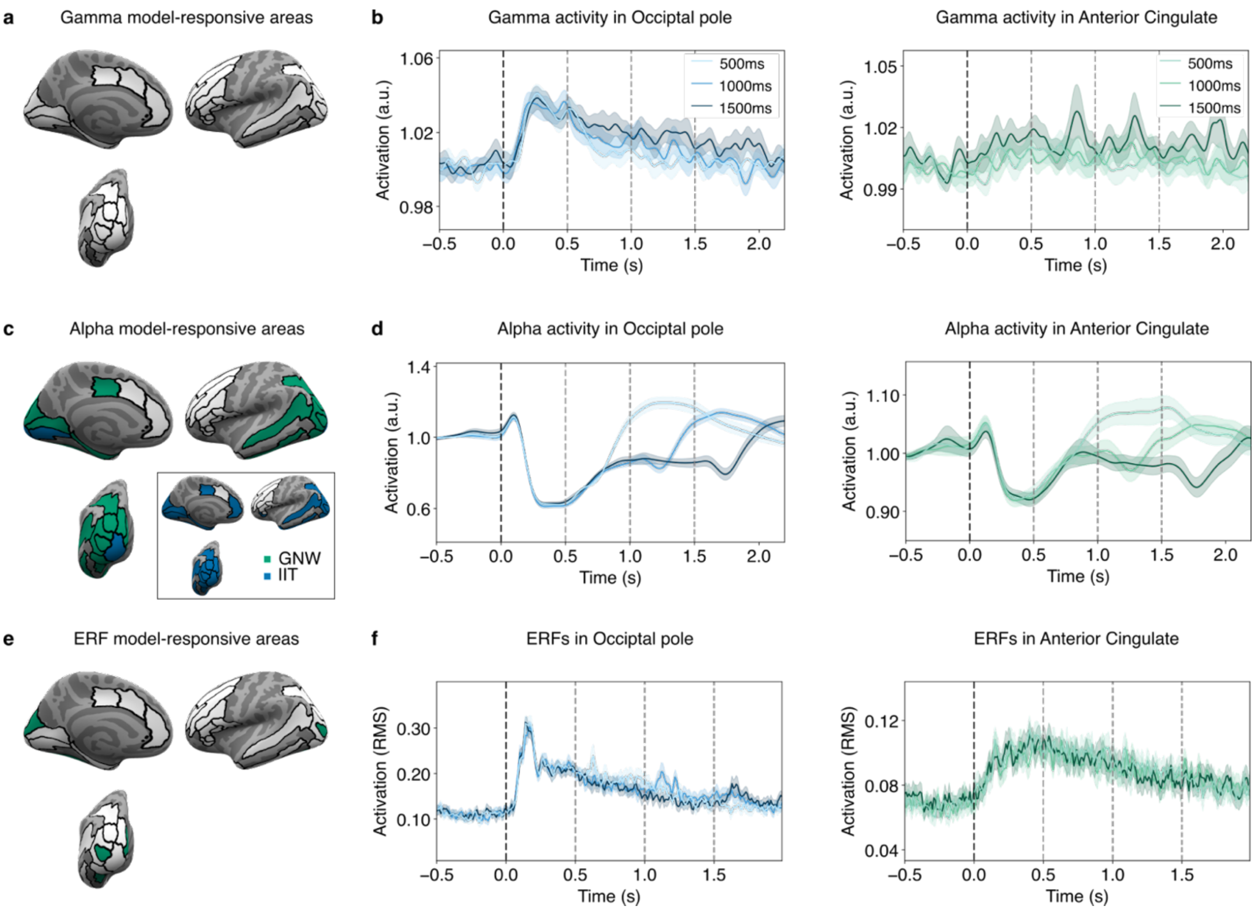

**Supplementary Figure 40**. Results of the task relevant analyses. **a.** Results of the LMM analysis on the gamma activity, where no parcel was captured by the theoretical models. **b**. Gamma activity time series in the Occipital pole and Middle-Posterior Cingulate Cortex for each duration. **c**. Results of the LMM analysis on the alpha activity. The inset represents the LMM results on the late time bins. **d**. Alpha activity time series in the Occipital Pole and Middle-Posterior Cingulate Cortex for each duration. **e**. Results of the LMM analysis on the ERFs. **f**. ERFs time series in the Occipital Pole and Middle-Posterior Cingulate Cortex for each duration. Vertical dotted lines represent stimulus onset (black) and offset (gray, for the three different offsets). Shaded areas denote the confidence intervals.

### Pre-registered analyses: Representational Similarity Analysis (RSA)

#### iEEG: RSA analysis

For this analysis, data from 29 patients with complete data sets were used. Among these patients, 28 had electrodes implanted in the posterior region of the brain, and 28 had electrodes implanted in the prefrontal cortex (PFC) region. In total, there were 583 electrodes in the posterior region and 576 electrodes in the PFC region. We performed a cross-temporal RSA on each theory-defined ROI, separately for each of the stimulus properties (category, identity and orientation). We correlated the temporal generalization matrices with the two theory-predicted matrices using Kendall's Tau correlation and assessed the significance of the correlation value for each theory-predicted matrix through a label shuffling permutation test. For every matrix that was found significant (for either of the theories), we then directly compared the correlation values between the theory-predicted matrices by subtracting the correlation values obtained for the GNWT-predicted matrix from the IIT-predicted matrix, and testing the significance of the difference. Using this approach, a difference greater than zero would indicate an advantage for IIT while a difference smaller than zero would indicate an advantage for GNWT. This allowed us to determine if a theory is empirically validated by performing better than the competing theory within its own ROI. This approach provides a stricter test than simply evaluating the significance of the correlation between a theory's predicted pattern and the observed data. Figure 3 and Extended Data Figure 5 illustrate the results for each of the stimulus properties and theory-defined ROI separately. In Supplementary Table 15, we report the full set of statistical results for each of these tests.

As we report in the main text, in the posterior ROI, all contrasts including category and identity were found to be significantly correlated with the IIT predicted temporal patterns, although for orientation none of the stimulus categories showed a sustained representation, and thus did not correlate with the IIT predicted temporal pattern. When directly contrasting the differences in correlation between the IIT and the GNWT predicted matrix in posterior cortex, we found that faces vs. objects in the task irrelevant condition and object identity were significantly better explained by the IIT predicted temporal pattern. The lack of significant advantage of the IIT vs. GNWT models for the other contrasts, despite the significant correlations with the IIT model itself, is mostly explained by the different type of test applied when directly comparing the two theory predicted matrices against each other. Because there is a partial overlap between the theories' predictions (e.g., at stimulus onset) observing a significant advantage for one theory over the other becomes more challenging. Reflecting this point, GNWT predicts content representation from 0.3-0.5 s following stimulus onset, which was observed in most cases. Yet while the correlation with the GNWT predicted matrix in posterior cortex was mostly found to be positive, it was almost never significant (with the exception of the contrast: task relevant letters vs. false-fonts).

In PFC, none of the contrasts for category, identity or orientation yielded patterns significantly correlated with the GNWT model, which can be explained by the lack of the predicted patterns after stimulus offset. In a few cases, the correlation with the IIT predicted temporal matrix was found to be significant: namely, faces vs. objects in the task relevant condition, and identity for letters and false fonts; with a significant difference in favor of the IIT predicted temporal pattern over the GNWT predicted temporal pattern for false font identity. These results indicate that at least certain stimulus features might be represented in a sustained fashion in prefrontal regions. Notably, they do not constitute evidence for IIT, as they were observed in PFC and not in posterior cortex as predicted by the theory.

| contrast | # electrodes | roi | $\tau_{iit}$ | $p_{iit}$ | $\tau_{gnwt}$ | $p_{gnwt}$ | $\tau_{iit} - \tau_{gnwt}$ | $p_{diff}$ |
|---|---|---|---|---|---|---|---|---|

| | | | | | | | | |
|---|---|---|---|---|---|---|---|---|
| Faces vs. objects (task irrelevant) | 583 | posterior | 0.62 | 0.000 * | -0.01 | 0.507 | 0.31 | 0.007 * |
| Faces vs. objects (task relevant) | 583 | posterior | 0.41 | 0.004 * | 0.09 | 0.210 | 0.16 | 0.127 |
| Letters vs false fonts (task irrelevant) | 583 | posterior | 0.25 | 0.019 * | -0.06 | 0.703 | 0.15 | 0.106 |
| Letters vs false fonts (task relevant) | 583 | posterior | 0.22 | 0.033 * | 0.20 | 0.017 * | 0.01 | 0.899 |
| Face identity | 583 | posterior | 0.42 | 0.003 * | -0.06 | 0.651 | 0.24 | 0.060 |
| Objects identity | 583 | posterior | 0.66 | 0.000 * | -0.06 | 0.657 | 0.36 | 0.003 * |
| Letters identity | 583 | posterior | 0.36 | 0.011 * | 0.01 | 0.473 | 0.18 | 0.125 |
| False-fonts identity | 583 | posterior | 0.45 | 0.002 * | 0.07 | 0.298 | 0.19 | 0.109 |
| Face orientation | 583 | posterior | 0.22 | 0.133 | 0.00 | 0.509 | | |
| Objects orientation | 583 | posterior | -0.07 | 0.561 | -0.08 | 0.729 | | |
| Letters orientation | 583 | posterior | -0.27 | 0.938 | 0.05 | 0.348 | | |
| False fonts orientation | 583 | posterior | -0.20 | 0.837 | 0.08 | 0.287 | | |
| Faces vs objects (task irrelevant) | 576 | PFC | 0.02 | 0.431 | 0.17 | 0.051 | | |
| Faces vs objects (task relevant) | 576 | PFC | 0.40 | 0.003 * | 0.08 | 0.246 | 0.16 | 0.767 |
| Letters vs false-fonts (task irrelevant) | 576 | PFC | -0.02 | 0.541 | 0.10 | 0.226 | | |
| Letters vs false-fonts (task relevant) | 576 | PFC | 0.13 | 0.181 | -0.01 | 0.486 | | |
| Faces identity | 576 | PFC | -0.40 | 0.992 | -0.04 | 0.607 | | |
| Objects identity | 576 | PFC | 0.16 | 0.187 | 0.17 | 0.130 | | |
| Letters identity | 576 | PFC | 0.45 | 0.011* | -0.10 | 0.746 | 0.28 | 0.311 |
| False-fonts identity | 576 | PFC | 0.44 | 0.003* | -0.24 | 0.971 | 0.34 | 0.004 * |
| Faces orientation | 576 | PFC | 0.07 | 0.338 | 0.13 | 0.215 | | |
| Objects orientation | 576 | PFC | 0.02 | 0.458 | -0.03 | 0.601 | | |
| Letters orientation | 576 | PFC | 0.16 | 0.193 | 0.20 | 0.060 | | |
| False-fonts orientation | 576 | PFC | -0.08 | 0.698 | -0.10 | 0.749 | | |

**Supplementary Table 15:** Results of the RSA for each of the investigated stimulus properties (category, identity and orientation) in each theory-defined ROI (N subjects=28 each). First column: stimulus property studied. Second column: number of electrodes within the theory-defined ROI. Third and fourth column: kendall's Tau correlation between the IIT ($\tau_{iit}$) predicted matrix and the temporal generalization matrices, and associated P-value from the label shuffle permutation test ($p_{iit}$), respectively. Fifth and sixth column: kendall's Tau correlation and P-value for GNWT ($\tau_{gnwt}$, $p_{gnwt}$). Seventh and eighth column: 0.5 centered correlation difference between the IIT and GNWT correlation and associated P-value ($\tau_{iit} - \tau_{gnwt}$ and $p_{diff}$ columns). The stars in the P-value column represent the significance of the test at alpha=0.05.

### Exploratory analyses: RSA

#### iEEG: Feature selection

As described above, we had a total of 583 and 576 electrodes in posterior and PFC ROIs, respectively. However, only a fraction of those were found to be responsive to our task (see Onset/offset analysis above). While multivariate analyses are thought to be robust to noise, adding irrelevant features might nonetheless obscure the findings. This in turn might explain the lack of significant representation for some of the contrasts in each ROIs. To rule out this possibility, we performed another RSA analysis, selecting the top 200 most informative features using select-k-best in a cross-validated fashion (5 folds for category and orientation, 3 folds for identity as only 3 trials were available for each label in this contrast level). Supplementary Figure 41 summarizes these results using this feature selection approach and Supplementary Table 16 provides a full description of the results.

In the posterior ROI, we observed strong evidence for IIT-predicted temporal patterns at the level of category: we found a significant correlation difference in favor of IIT for faces vs. objects in the task relevant and irrelevant condition and also for the letters vs. false-fonts contrast in the task irrelevant condition (Supplementary Figure 41, first row). However, for identity level analyses no significant representation was found for any of the stimulus categories, except objects (Supplementary Figure 41, second row). These results diverge from those reported without any feature selection, yet they are easily explained by the low number of trials available for each identity when using feature selection. As feature selection must be cross validated to avoid double dipping, only ⅔ of the trials were used to compute the within class corrected distances, while the remaining ⅓ were used for feature selection. The increased SNR offered by the feature selection is therefore counteracted by the lower number of trials available for computing the relevant metric. As such, while the RSA matrices both for faces and objects identity significantly correlated with IIT predicted matrices, the difference between the correlation for IIT and GNWT did not reach significance. Finally, for orientation, no significant representation was observed for any of the stimulus categories (Supplementary Figure 41, third row). Interestingly, the temporal generalization matrix for face orientation was observed to be significantly correlated both with IIT and GNWT predicted temporal matrices.

In PFC, feature selection yielded a larger improvement for the representation of category. Consistent with the results found without feature selection, we observed significant faces vs. objects representation in the task relevant and task relevant conditions, as well as significant representation of letters vs. false-fonts in the task relevant condition. For all of those contrasts, information was present only at stimulus onset but not at stimulus offset, contrary to GNWT's predictions (Supplementary Figure 41, fourth row). The RSA patterns themselves were similar to those observed without feature selection. Yet, the latter were not significant at stimulus onset, suggesting that feature selection might confer higher sensitivity to the RSA analysis. These results are consistent with the temporal generalization patterns observed in the decoding analysis.

Though these results suggest category representation exclusively at stimulus onset in PFC, no significant representation for identity or orientation for any of the stimulus categories was observed in PFC, either at stimulus onset or at stimulus offset. Also, the predicted offset patterns were not observed for any of the contrasts (Supplementary Figure 41, fifth and sixth row). Accordingly, none of the temporal generalization matrices were found to be significantly correlated with the GNWT predicted patterns.

Taken together, the RSA analysis with feature selection, while at times improving the statistical significance of certain results, does not change the main conclusions as we found no support for the GNWT prediction of both an onset and an offset reactivation of information.

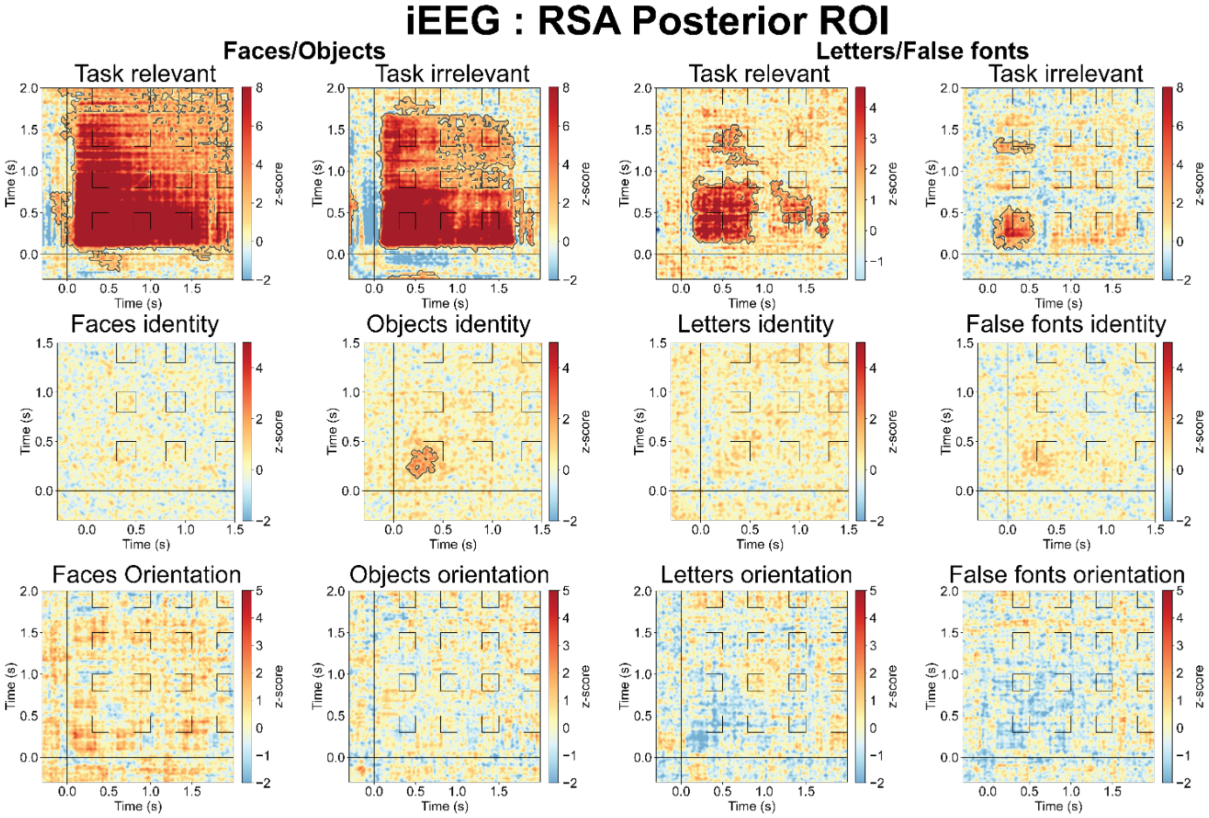
iEEG : RSA Posterior ROI
Faces/Objects
Letters/False fonts
Task relevant
Task irrelevant
Task relevant
Task irrelevant
Faces identity
Objects identity
Letters identity
False fonts identity
Faces Orientation
Objects orientation
Letters orientation
False fonts orientation
Time (s)
z-score

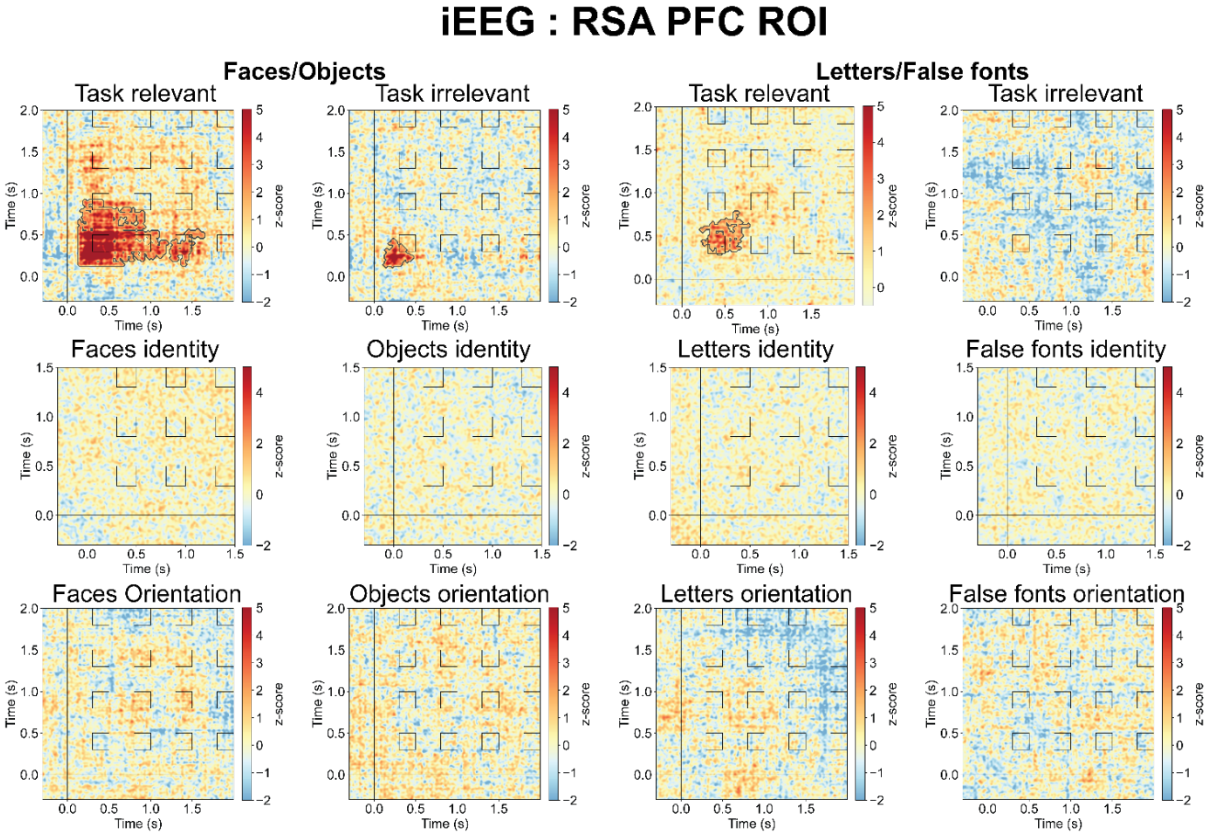
iEEG : RSA PFC ROI
Faces/Objects
Letters/False fonts
Task relevant
Task irrelevant
Task relevant
Task irrelevant
Faces identity
Objects identity
Letters identity
False fonts identity
Faces Orientation
Objects orientation
Letters orientation
False fonts orientation
Time (s)
z-score

**Supplementary Figure 41**. Results of temporal generalization RSA with 200 features selection. The upper three rows show the results from the posterior ROIs, the lower three rows show the results from the PFC ROIs. For each ROI, results are reported for category level contrasts (first row), identity level contrast for each category separately (second row) and orientation level contrasts for each category separately (last row). For the category level contrasts, task relevant and irrelevant trials were investigated separately. For identity and orientation contrasts, task relevant and irrelevant trials were combined. Furthermore, for category and orientation, only 1.5 s trials were analyzed. For identity, 1.0 s and 1.5 s trials were combined due to the low number of trials across subjects. As a result of this, the X-axis for the category and orientation conditions differs from the one for identity. The contour in the matrices depicts the statistically significant clusters determined using cluster-based permutation.

| contrast | # features | roi | $\tau_{iit}$ | $p_{iit}$ | $\tau_{gnwt}$ | $p_{gnwt}$ | $\tau_{iit} - \tau_{gnwt}$ | $p_{diff}$ |
|---|---|---|---|---|---|---|---|---|
| Faces vs. objects (task irrelevant) | 200 | posterior | 0.61 | 0.000 * | 0.00 | 0.452 | 0.30 | 0.002 ** |
| Faces vs. objects (task relevant) | 200 | posterior | 0.51 | 0.000 * | 0.03 | 0.342 | 0.24 | 0.007 ** |
| Letters vs false fonts (task irrelevant) | 200 | posterior | 0.29 | 0.002 * | -0.07 | 0.792 | 0.18 | 0.019 ** |
| Letters vs false fonts (task relevant) | 200 | posterior | 0.28 | 0.003 * | 0.15 | 0.041 * | 0.07 | 0.388 |
| Face identity | 200 | posterior | 0.24 | 0.012 * | -0.03 | 0.619 | 0.13 | 0.153 |
| Objects identity | 200 | posterior | 0.24 | 0.012 * | 0.17 | 0.046 * | 0.04 | 0.654 |
| Letters identity | 200 | posterior | -0.09 | 0.743 | 0.09 | 0.190 | | |
| False-fonts identity | 200 | posterior | 0.23 | 0.102 | 0.00 | 0.489 | | |
| Face orientation | 200 | posterior | 0.21 | 0.032 * | 0.20 | 0.029 * | 0.00 | 0.983 |
| Objects orientation | 200 | posterior | -0.09 | 0.636 | -0.04 | 0.644 | | |
| Letters orientation | 200 | posterior | -0.27 | 0.965 | 0.07 | 0.256 | | |
| False fonts orientation | 200 | posterior | -0.17 | 0.814 | 0.08 | 0.249 | | |
| Faces vs objects (task irrelevant) | 200 | PFC | 0.03 | 0.355 | 0.17 | 0.050 | -0.07 | 0.461 |
| Faces vs objects (task relevant) | 200 | PFC | 0.45 | 0.001 * | 0.08 | 0.214 | 0.19 | 0.458 |
| Letters vs false-fonts (task irrelevant) | 200 | PFC | -0.01 | 0.506 | 0.08 | 0.224 | | |
| Letters vs false-fonts (task relevant) | 200 | PFC | 0.14 | 0.140 | 0.03 | 0.365 | | |
| Faces identity | 200 | PFC | -0.26 | 0.990 | 0.02 | 0.409 | | |
| Objects identity | 200 | PFC | 0.09 | 0.200 | 0.04 | 0.336 | | |
| Letters identity | 200 | PFC | 0.24 | 0.049 * | 0.02 | 0.435 | 0.11 | 1.000 |

| | | | | | | | | |
|---|---|---|---|---|---|---|---|---|
| False-fonts identity | 200 | PFC | 0.26 | 0.010 * | -0.05 | 0.653 | 0.15 | 0.456 |
| Faces orientation | 200 | PFC | 0.40 | 0.001 * | -0.25 | 0.985 | 0.32 | 0.003 * |
| Objects orientation | 200 | PFC | -0.02 | 0.563 | 0.00 | 0.471 | | |
| Letters orientation | 200 | PFC | 0.14 | 0.176 | 0.20 | 0.044 * | -0.03 | 1.000 |
| False-fonts orientation | 200 | PFC | -0.06 | 0.685 | -0.10 | 0.786 | | |

**Supplementary Table 16.** Results of the RSA for each of the investigated stimulus properties (category, identity and orientation) in each theory-defined ROI with 200 features selection.

#### GNWT extended windows

The investigated time windows were determined based on theoretical considerations, and were aimed at maximizing the difference between the theories' predictions. In particular, for IIT it was critical that the tested time windows did not include the stimulus evoked response as much as possible, while for GNWT it was critical to capture the late responses marking the predicted ignition (i.e., >0.25 s). As a compromise between those two competing needs, a time window starting at 0.3 s was chosen. However, upon inspecting the data, we observed several cases where the responses in PFC had earlier latencies than the preregistered time window (0.3-0.5 s), yet still within the range of latencies predicted by GNWT (<0.25 s). To rule out the possibility that the lack of evidence for GNWT simply stemmed from the selection of the time window, we performed an exploratory analysis on the 1.5 s stimulus duration trials, using a more extended and earlier time window to capture a representation of conscious content in the workspace, had it occurred (from 0.25-0.5 s for GNWT onset ignitions and 1.75-2.0 s for GNWT offset ignitions, and from 0.25-1.5 s for IIT's sustained activity prediction). This also enabled us to investigate whether the offset responses predicted by GNWT may have also occurred earlier. Since this analysis only tested GNWT's prediction, it was only carried out in the PFC ROIs.

Results from this extended time window analysis are described in Supplementary Table 17. The observed findings did not change the main conclusions (as shown in Supplementary Table 15). Notably, for the category representation of faces/objects in task irrelevant trials, we found a significant correlation with the GNWT predicted matrix but not with the IIT predicted matrix, yet the direct contrast between the two theory predicted matrices was not statistically significant, thus failing the predefined test. Notably, this correlation seems to be driven by off-diagonal non-significant increases in within-class corrected distances, while GNWT predicts generalization between stimulus onset and offset, which should be observed also along the diagonal in the predicted time window (from 1.75-2.0 s). Thus, even if the statistical test would have yielded a significant result, the results would not be entirely compatible with the GNWT predicted offset patterns.

| contrast | # electrodes | roi | $\tau_{iit}$ | $p_{iit}$ | $\tau_{gnwt}$ | $p_{gnwt}$ | $\tau_{iit} - \tau_{gnwt}$ | $p_{diff}$ |
|---|---|---|---|---|---|---|---|---|
| Faces vs objects (task irrelevant) | 576 | PFC | 0.00 | 0.479 | 0.26 | 0.004 * | -0.13 | 0.617 |
| Faces vs objects (task relevant) | 576 | PFC | 0.45 | 0.000 * | 0.02 | 0.403 | 0.22 | 0.767 |
| Letters vs false-fonts (task irrelevant) | 576 | PFC | -0.06 | 0.657 | 0.09 | 0.224 | | |
| Letters vs false-fonts (task relevant) | 576 | PFC | 0.13 | 0.166 | 0.01 | 0.431 | | |
| Faces identity | 576 | PFC | -0.37 | 0.990 | -0.03 | 0.577 | | |
| Objects identity | 576 | PFC | 0.19 | 0.130 | 0.19 | 0.087 | | |
| Letters identity | 576 | PFC | 0.41 | 0.015 * | -0.17 | 0.870 | 0.29 | 0.311 |
| False-fonts identity | 576 | PFC | 0.18 | 0.137 | 0.14 | 0.190 | | |
| Faces orientation | 576 | PFC | 0.43 | 0.005 * | -0.22 | 0.966 | 0.33 | 0.004* |
| Objects orientation | 576 | PFC | 0.03 | 0.398 | -0.05 | 0.656 | | |
| Letters orientation | 576 | PFC | 0.19 | 0.139 | 0.17 | 0.072 | | |
| False-fonts orientation | 576 | PFC | -0.11 | 0.778 | -0.06 | 0.661 | | |

**Supplementary Table 17.** Results of the RSA for each of the investigated stimulus properties (category, identity and orientation; N subjects=28 for each ROI), when extending the predicted time-windows by 0.05 s, to explore the possibility of offset responses at earlier latencies than expected.

#### GNWT model including only onset

GNWT's preregistered prediction stated that the update of the workspace following stimulus offset should reinstate the information conveyed by the stimulus that has just disappeared. However, one might argue instead that an alternative mechanism for maintaining conscious perceptions over time only requires an update conveying information about the next conscious content. Under this interpretation of GNWT, information about the content of consciousness will be represented transiently following the onset of the stimulus only, without requiring an offset response.

We tested this prediction using an alternative GNWT derived matrix, where representation is predicted to occur only from 0.3-0.5 s (post stimulus onset). The results are described in Supplementary Table 18. Notably, while all category level contrasts were found to be significantly correlated with this GNWT 'onset only' model, and only one of the four category contrasts (faces vs. objects, task relevant) was correlated with the IIT model, we did not observe any significant differences between the IIT and GNWT models for any of the category contrasts when they were directly compared, indicating no stronger support for the GNWT prediction over the IIT prediction or vice versa.

With respect to identity, significant correlations were evident for both the GNWT and IIT models for letters identity, yet with no stronger support for GNWT when directly compared to IIT. No other identities (i.e., within the face, object, or false-font category) showed any significant correlations with either theory's model.

For orientation, we found no support for a GNWT onset-only model for any of the stimulus categories, and a significant correlation with the IIT model only for face orientation. As mentioned above in the preregistered RSA analysis section, these results do not lend support to IIT as they were observed in PFC and not in the posterior cortex.

| contrast | # electrodes | roi | $\tau_{iit}$ | $p_{iit}$ | $\tau_{gnwt}$ | $p_{gnwt}$ | $\tau_{iit} - \tau_{gnwt}$ | $p_{diff}$ |
|---|---|---|---|---|---|---|---|---|
| Faces vs objects (task irrelevant) | 576 | PFC | 0.02 | 0.431 | 0.26 | 0.002 * | -0.12 | 0.617 |
| Faces vs objects (task relevant) | 576 | PFC | 0.40 | 0.003 ** | 0.35 | 0.000 ** | 0.02 | 0.767 |
| Letters vs false-fonts (task irrelevant) | 576 | PFC | -0.02 | 0.541 | 0.21 | 0.042 * | -0.11 | 1.000 |
| Letters vs false-fonts (task relevant) | 576 | PFC | 0.13 | 0.181 | 0.34 | 0.000 * | -0.10 | 1.000 |
| Faces identity | 576 | PFC | -0.40 | 0.992 | -0.35 | 0.997 | | 1.000 |
| Objects identity | 576 | PFC | 0.16 | 0.187 | 0.14 | 0.177 | | 1.000 |
| Letters identity | 576 | PFC | 0.45 | 0.011 * | 0.28 | 0.043 * | 0.08 | 0.318 |
| False-fonts identity | 576 | PFC | 0.07 | 0.338 | 0.17 | 0.146 | | 1.000 |
| Faces orientation | 576 | PFC | 0.44 | 0.003 * | -0.11 | 0.767 | 0.27 | 0.004** |
| Objects orientation | 576 | PFC | 0.02 | 0.458 | 0.03 | 0.391 | | 1.000 |
| Letters orientation | 576 | PFC | 0.16 | 0.193 | 0.11 | 0.209 | | 1.000 |
| False-fonts orientation | 576 | PFC | -0.08 | 0.698 | -0.10 | 0.727 | | 1.000 |

**Supplementary Table 18.** Results of the RSA for each of the investigated stimulus properties (category, identity and orientation) in the PFC ROIs (N subjects=28), testing a GNWT model which includes only a representation of information only at stimulus onset.

#### Exploratory analysis: iEEG cross-task decoding at stimulus offset

To investigate the decodability of stimulus category at stimulus offset, which was relevant to one of the predictions made by GNWT, we trained separate classifiers on electrodes from posterior and prefrontal ROIs (GNWT ROIs $N_{electrodes}$=576, IIT ROIs $N_{electrodes}$=583). Data from all stimulus durations were combined and aligned to the stimulus offset (-0.5 to 0.5 s) for each duration (0.5, 1.0, 1.5 s). Classifiers were trained to discriminate stimulus category (faces vs. objects) in the task irrelevant condition at each time-point and tested in the task relevant condition across all time-points. Significant decoding of stimulus category (faces vs. objects) was observed in the posterior

ROI extending to approximately 0.3 s after stimulus offset (Supplementary Figure 42). In the prefrontal ROI, decoding of stimulus category after stimulus offset was not observed.

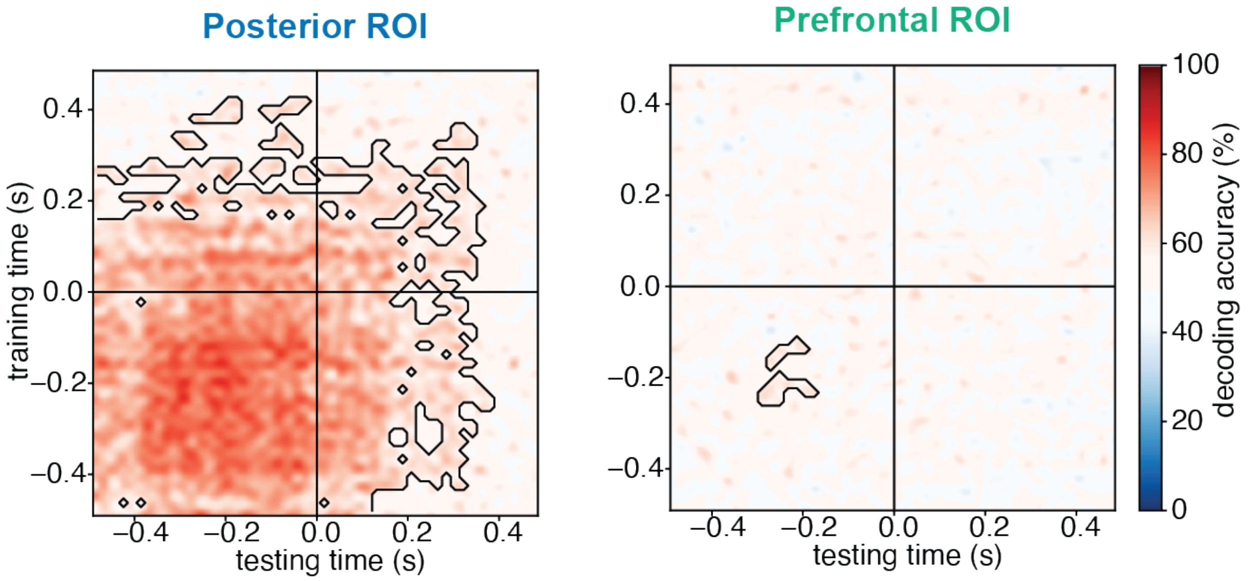

**Supplementary Figure 42**. Cross-task temporal generalization of decoding aligned to stimulus offset (-0.5 to 0.5 s) for iEEG. Pattern classifiers were trained to discriminate stimulus category (faces vs. objects) in the task irrelevant condition at each time-point and tested in the task relevant condition across all time-points (left: posterior ROIs; right: prefrontal ROIs). All trials from different durations were aligned to the stimulus offset (which is marked as time 0 in this figure).

#### MEG RSA analysis

We performed cross-temporal RSA on MEG cortical time series data, using the same methods as iEEG. For RSA of category and orientation, only 1.5s duration trials were entered into the analysis. We also used pseudotrial aggregation. For RSA of identity we combined 1.0 s and 1.5 s duration trials to compensate for the lower number of repetitions per identity. Pseudotrials were not applied for the identity analyses also due to too low number of trials (less than 20) for each identity.

Neither the posterior ROI nor prefrontal ROI exhibited a pattern consistent with the theory predictions. In the posterior ROI (Supplementary Figure 43), we observed information about category (faces vs. objects) only at stimulus onset, while category information for letters vs. false fonts was absent in the task irrelevant condition. Information about orientation was not observed for any of the four stimulus categories investigated. Identity information was observed for objects, letters and false fonts but not for faces. For the prefrontal ROI (Supplementary Figure 44), other than false font identity (only significant at the early time window of stimulus onset) we did not observe any information for any of the stimulus properties (category, orientation, identity).

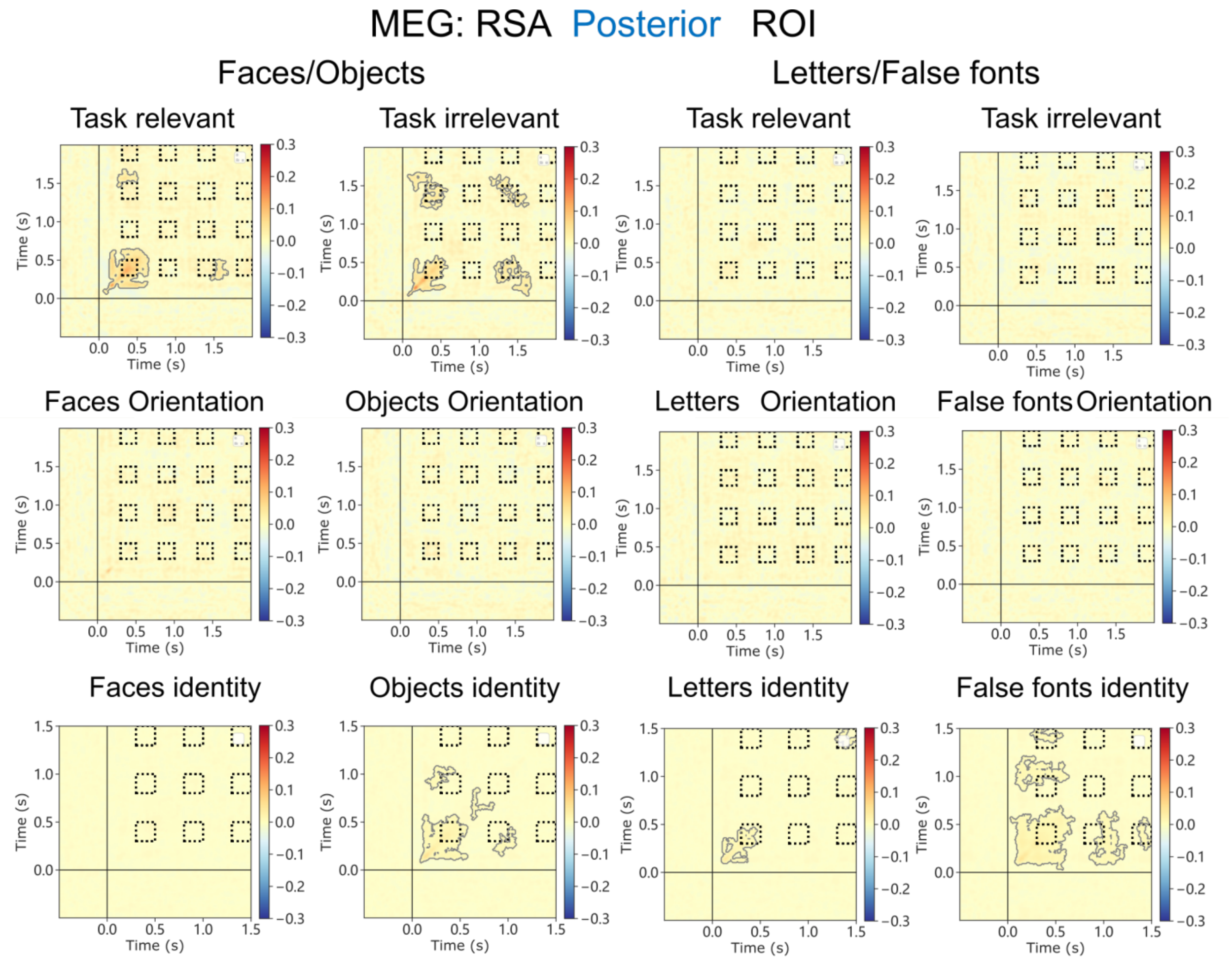

**Supplementary Figure 43**. Results of temporal generalization RSA for posterior ROIs. For each ROI, results are reported for category level contrasts (top row), orientation level contrasts for each category separately (middle row) and identity level contrast for each category separately (bottom row). For the category level contrasts, task relevant and irrelevant trials were investigated separately. For identity and orientation contrasts, task relevant and irrelevant trials were combined. Furthermore, for category and orientation, only 1.5 s trials were analyzed. For identity, 1.0 s and 1.5 s trials were combined due to the low number of trials for each identity. The contour in the matrices depicts the statistically significant clusters determined using cluster-based permutation.

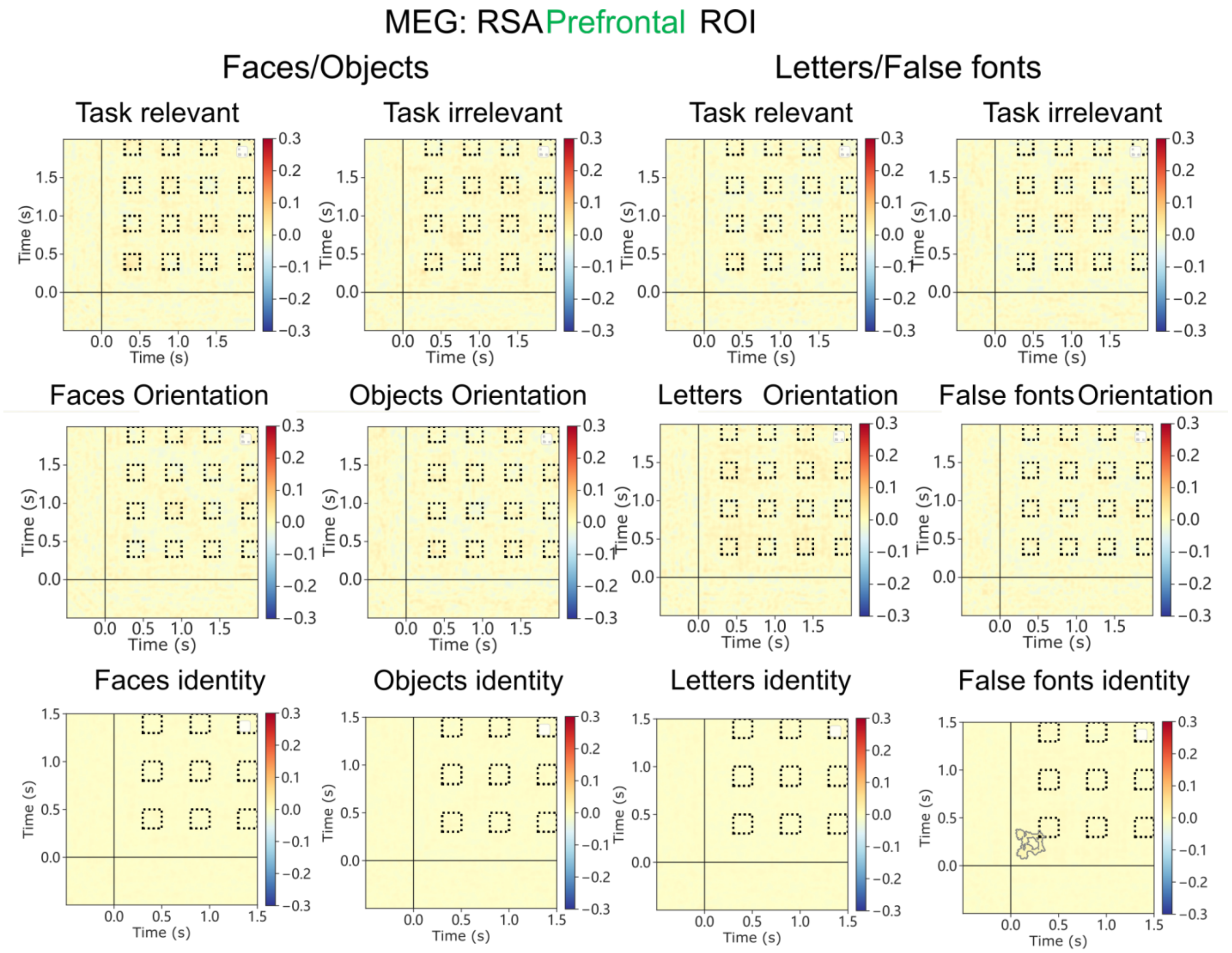

**Supplementary Figure 44**. Results of temporal generalization RSA for Prefrontal ROIs. Same conventions as in Supplementary Figure 43.

### Prediction #3: Interareal functional connectivity

In the main paper, we report the connectivity results for the task irrelevant trials, which constitute the most critical test for the theories. Here, we further describe analyses conducted either on the task relevant condition, where the signal is expected to be stronger due to task-based attentional amplification, or on the combined data from both task conditions, thus improving the signal-to-noise ratio by doubling the number of trials. These additional analyses allowed us to increase the chances of finding the predicted results, as well as to examine how one's choice of different tests influences the assessment of the predicted patterns. Since the fMRI analysis reported in the main text was already conducted across task conditions (i.e., combining all trials), we only report additional results from the iEEG and MEG analyses here (after first providing supplementary results for the fMRI gPPI analysis reported in the main paper).

#### Pre-registered analyses

##### fMRI Generalized Psycho-Physiological Interaction (gPPI) Table

In the main paper, we reported the results of the Generalized Psycho-Physiological Interaction (gPPI) analysis, combining task relevant and irrelevant trials. Supplementary Table 19 provides the full set of results. It shows that several regions such as Inferior Frontal Gyrus, Intra-Parietal Sulcus, Cuneus, and V1/V2 showed content-specific connectivity with the FFA seed. No significant clusters were observed when investigating connectivity with the FFA seed separately for task relevant and irrelevant conditions. Extended Data Figure 8 shows the clusters at an uncorrected $p<0.01$. Notably, no significant clusters were observed with the seed in Lateral Occipital Cortex, either when combining task relevant or irrelevant trials, or when performing the analysis separately per task.

| **Anatomical ROIs (Destrieux atlas)** | **Task relevant and task irrelevant trials combined** | |
|---|---|---|
| | n voxels | % voxels |
| **Posterior ROI** | | |
| G_and_S_occipital_inf | 0 | 0 |
| G_oc-temp_lat-fusifor | 0 | 0 |
| G_occipital_middle | 2 | 0.008 |
| S_oc_middle_and_Lunatus | 12 | 1.188 |
| G_cuneus | 294 | 11.732 |
| G_occipital_sup | 32 | 1.621 |
| G_oc-temp_med-Lingual | 115 | 3.836 |
| G_oc-temp_med-Parahip | 0 | 0 |
| G_temporal_inf | 0 | 0 |
| Pole_occipital | 78 | 3.161 |
| Pole_temporal | 0 | 0 |
| S_calcarine | 204 | 8.409 |

| | | |
|---|---|---|
| S_intrapariet_and_P_trans | 463 | 12.207 |
| S_oc_sup_and_transversal | 12 | 0.848 |
| S_temporal_sup | 0 | 0 |
| **PFC ROI** | | |
| G_and_S_cingul-Mid-Post | 0 | 0 |
| Lat_Fis-ant-Horizont | 0 | 0 |
| Lat_Fis-ant-Vertical | 0 | 0 |
| G_and_S_cingul-Ant | 0 | 0 |
| G_and_S_cingul-Mid-Ant | 0 | 0 |
| G_front_inf-Opercular | 0 | 0 |
| G_front_inf-Orbital | 0 | 0 |
| G_front_inf-Triangul | 122 | 7.345 |
| G_front_middle | 22 | 0.358 |
| S_front_middle | 0 | 0 |
| S_front_sup | 0 | 0 |
| S_front_inf | 68 | 3.274 |

**Supplementary Table 19**. Number and percentage of voxels in each ROI found significant in the gPPI analysis with combined task relevant and task irrelevant trials using FFA as a seed.

#### Task Relevant condition

##### Pre-registered analyses

The same Pairwise Phase Consistency (PPC) analysis was conducted solely on task relevant trials (See Supplementary Figure 45). In the iEEG data, cluster-based permutation tests revealed a significant difference in synchronization between face-selective and object selective electrodes and V1/V2 electrodes. This effect was found in an early time window and in a low-frequency band, in line with what was found in the task irrelevant condition. These effects were mostly explained by the synchronous activity elicited by the stimulus evoked response (Supplementary Figure 45 a-b, top row). In contrast, no content-selective PPC was found between face- and object-selective electrodes and PFC in the relevant time window (Supplementary Figure 45 a-b, bottom row).

In the MEG source data, cluster-based permutation tests revealed a significant difference in content specific synchronization between category selective nodes and V1/V2 in the task relevant condition: higher synchronization was found between face-selective nodes and V1/V2 for face stimuli, which remained significant even after removing the stimulus evoked response. We also found content-specific synchronization between face-selective nodes, object-selective nodes and PFC. Removing the evoked response reduced but did not completely abolish this synchronization. Notably, both of these synchronization effects (between category-selective areas and V1/V2 or PFC) were found in low-frequency bands in early time-windows.

Overall, compared to the main analyses on task irrelevant trials, the results of the PPC analysis on task relevant ones showed stronger modulations in content-specific synchronization. However, after removing the evoked response, the observed effects were too early in time, around the onset of the stimulus, to be considered meaningful, particularly in relation to V1/V2.

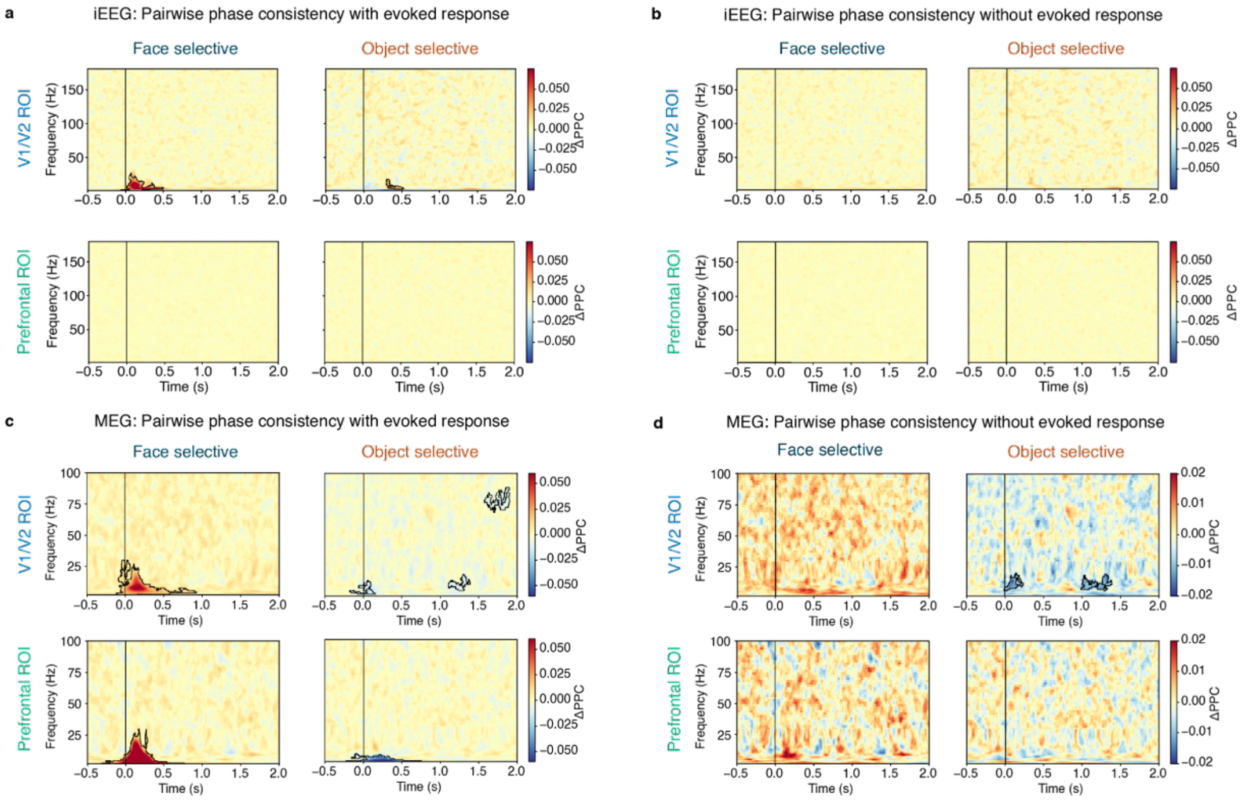

**Supplementary Figure 45**. Results of the PPC analysis on task relevant trials before and after removing the evoked response on iEEG and MEG source data. **a** iEEG PPC analysis of task relevant trials revealed significant content-selective synchrony (faces > objects for face-selective electrodes; objects > faces for object-selective electrodes) in V1/V2 ROIs (top row), but not in PFC ROIs (bottom row). **b.** After regressing out the evoked response, iEEG showed no significant content-selective connectivity in task relevant trials. **c.** MEG PPC analysis of task relevant trials revealed significant category-selective synchrony below 25 Hz for the face-selective GED filter (i.e., faces > objects for face-selective electrodes) in V1/V2 (top row) and PFC ROIs (bottom row) and for object-selective synchrony (objects > faces for object-selective electrodes) in PFC only. **d.** Removing the evoked response from MEG data significantly reduced but did not completely abolish the synchronization.

#### Exploratory analyses: DFC

The task relevant trials were also analyzed using the Dynamic Functional Connectivity (DFC)[7] method, with the same parameters as the PPC analysis, including restricting the analysis to the intermediate (1.0 s) and long (1.5 s) duration trials (see methods). The results are described in Supplementary Figure 46.

In iEEG, we observed significant connectivity between face selective electrodes and V1/V2. Connectivity was sustained up to 1 s after stimulus onset and present across several frequency bands, most predominantly in the gamma band between 50-100 Hz. Significant connectivity between object selective electrodes and V1/V2 was also observed predominantly in the gamma band, but it was briefer, lasting up to 0.5 s after stimulus onset. Significant, content-specific connectivity between face-selective electrodes and PFC was also observed, spanning a range of frequencies from the beta band up to the HG band, which was also extended in time up to 1 s after stimulus onset. In contrast, DFC between object selective electrodes and PFC was spottier and briefer, with an initial, peak in the HG range up until ~0.4 s, followed by a brief increase in the beta/low gamma range around 0.8 s after stimulus onset (but stronger for face stimuli). These effects were not entirely explained by the synchronous activity elicited by the stimulus evoked response as they remain after regressing out the evoked response (Supplementary Figure 46b).

For the MEG cortical time series, the results of the cluster-based permutation tests on the data (without regressing out the evoked response) revealed a significant difference between conditions for all nodes in the time window of 0 to 0.5 s from stimulus presentation (Supplementary Figure 46c). Specifically, we observed a significant increase in content-specific DFC in the low frequency range between the face-selective node and both PFC and V1/V2. This was accompanied by a simultaneous reduction in content-specific DFC in the high frequency range. We also found a smaller but significant modulation in synchrony between the object-selective node and both PFC and V1/V2 within the first 0.5 s after stimulus onset, mainly in the low frequency range.

To further investigate the stability of these results, we repeated the MEG DFC analysis after removing the evoked response, which largely removed the changes in connectivity observed at high frequencies while leaving virtually intact those observed at low frequencies (Supplementary Figure 46d). DFC between V1/V2 and object selective nodes was sustained up to 1.0 second in the alpha band, while briefer in time, lasting up to 0.5 seconds, between face-selective nodes and V1/V2. A comparable pattern of connectivity was observed between PFC and face-selective nodes. Overall, DFC was more pronounced between the face-selective node and both PFC and V1/V2 than in the object-selective node, in line with what we observed in the main analysis on task irrelevant trials.

These findings demonstrate that including the evoked responses can affect the DFC analysis by altering the high frequency synchronization, as well as potentially blocking the observation of smaller non-evoked synchronization patterns that are only noticeable if the evoked response is removed. This should accordingly be taken into account in future studies.

Overall, the results of our PPC analyses did not provide convincing evidence for either theory's prediction of either sustained (IIT) or phasic (GNWT) connectivity between relevant theory ROIs and content-selective nodes. Most of the results either showed no connectivity, or connectivity attributed to the evoked response and thus not related to content-specific synchrony. On the other hand, the power-based DFC analysis did show evidence of content-specific responses, but were either inconsistent between the iEEG and MEG modalities, or did not align with either theory's prediction. Therefore, no conclusive argument could be drawn for either theory's prediction.

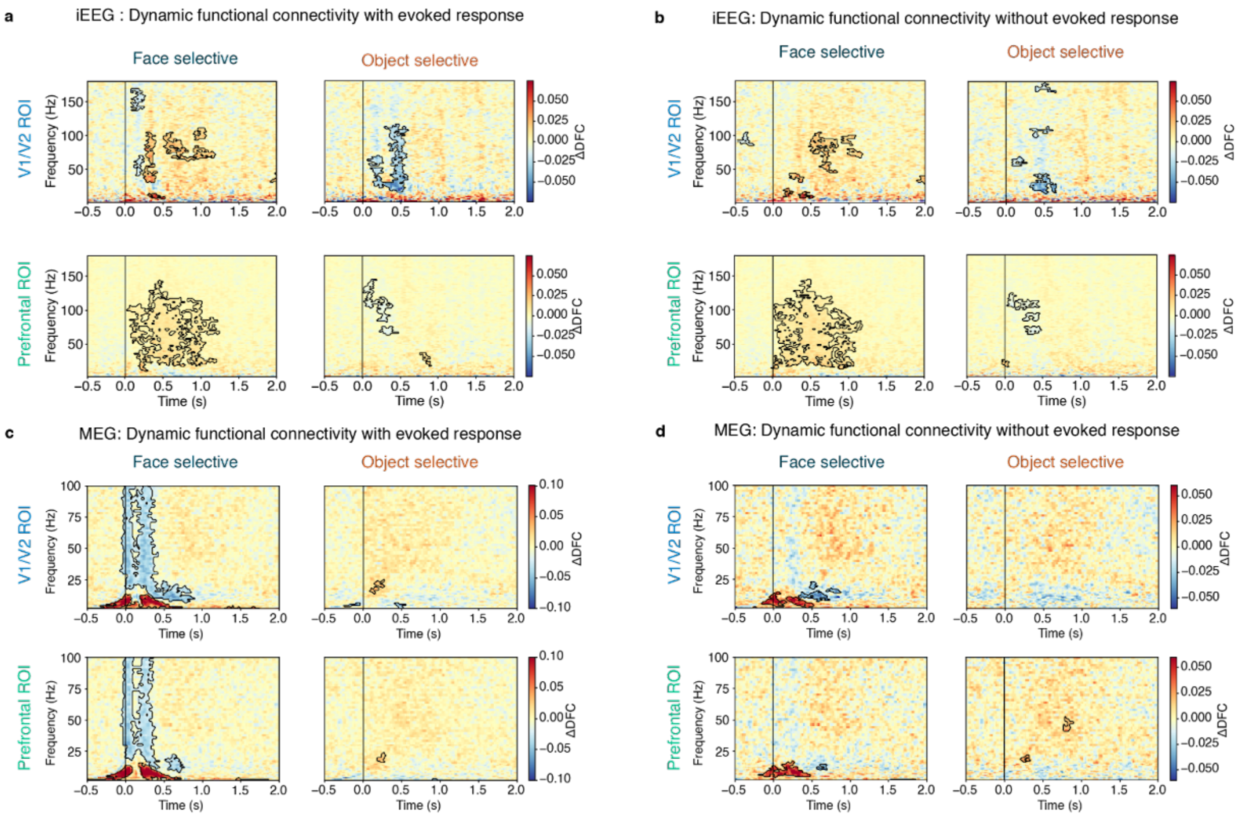

**Supplementary Figure 46**. Results of the DFC analysis on task relevant trials before and after removing the evoked response on iEEG and MEG source data. **a**. iEEG connectivity showed sustained (0-1 s) synchrony for face-selective electrodes to both V1/V2 regions and PFC. **b**. iEEG synchrony remained largely intact after removing the evoked response, but with reduced synchrony during the initial 0- 0.5s window for the V1/V2 ROI. **c**. MEG connectivity showed low-frequency DFC (< 25 Hz) between the face-selective node and both V1/V2 and PFC during the initial 0-0.5 s time window for faces (red) and high-frequency DFC (25-100 Hz) for objects (blue) in these same face-selective nodes. **d**. Removing the evoked response from MEG data abolished the high-frequency connectivity, while largely preserving the low-frequency effects, and revealing some object-selective (blue) connectivity in the alpha-band.

#### Task Relevant and Task Irrelevant combined

##### Pre-registered analyses

In the main paper, we reported the iEEG and MEG results on connectivity focusing on the task irrelevant trials, as those were the most diagnostic to testing the theories' predictions. In the supplement above, we repeated these analyses for the task relevant condition. However, as removing the evoked response from the single trials yielded no consistent connectivity either with V1/V2 or PFC, we conducted the same analyses combining task relevant and task irrelevant trials. The aim of this control analysis was to maximize statistical power by increasing the number of trials. We report the PPC results before and after subtracting the evoked responses from the single trials both for iEEG and MEG (Supplementary Figure 47). For the iEEG data, PPC results were comparable to those found in the analyses conducted separately on the task irrelevant and relevant trials: early content-specific synchronization was observed between face and object selective electrodes and V1/V2 predominantly in a low frequency band. However, this effect was abolished when removing the evoked response. No significant synchronization between PFC and face or object selective electrodes was found. This was true regardless of whether the evoked responses were removed from the data.

In MEG source data, cluster-based permutation tests showed a significant difference in low-frequency phase-synchronization between the face-selective node and PFC and V1/V2 in the period right after stimulus onset. A significant difference was also detected between the object-selective node and V1/V2. However, this effect was not sustained throughout presentation of the stimuli (1.5 s), and instead appeared 0.5 s after stimulus onset. Removing the evoked response again significantly reduced the synchronization and in some cases completely removed it (e.g., face-selective nodes and V1/V2 connectivity and the object-selective node and PFC connectivity).

Overall, the results of the phase-synchronization analyses on iEEG and MEG source data on the combined task relevant and task irrelevant conditions did not provide clear support for either GNWT or IIT, akin to the main conclusion of the analysis conducted on the task irrelevant condition. Specifically, the results provide weak evidence for a content-specific modulation of synchronization between the category-selective nodes and both the PFC and the V1/V2 ROIs. Connectivity with V1/V2 was however not sustained, and observed mostly in low-frequencies <25 Hz, in contrast with IIT's predictions. In contrast, the MEG data revealed synchronization between PFC and the face-selective node at the ignition time window, as predicted by GNWT. However, this effect appears to be driven mostly by the evoked responses, as after removal of the evoked response, the synchronization that remained was earlier than the predicted GNWT time-window of ignition, i.e., ~0.1-0.3 s.

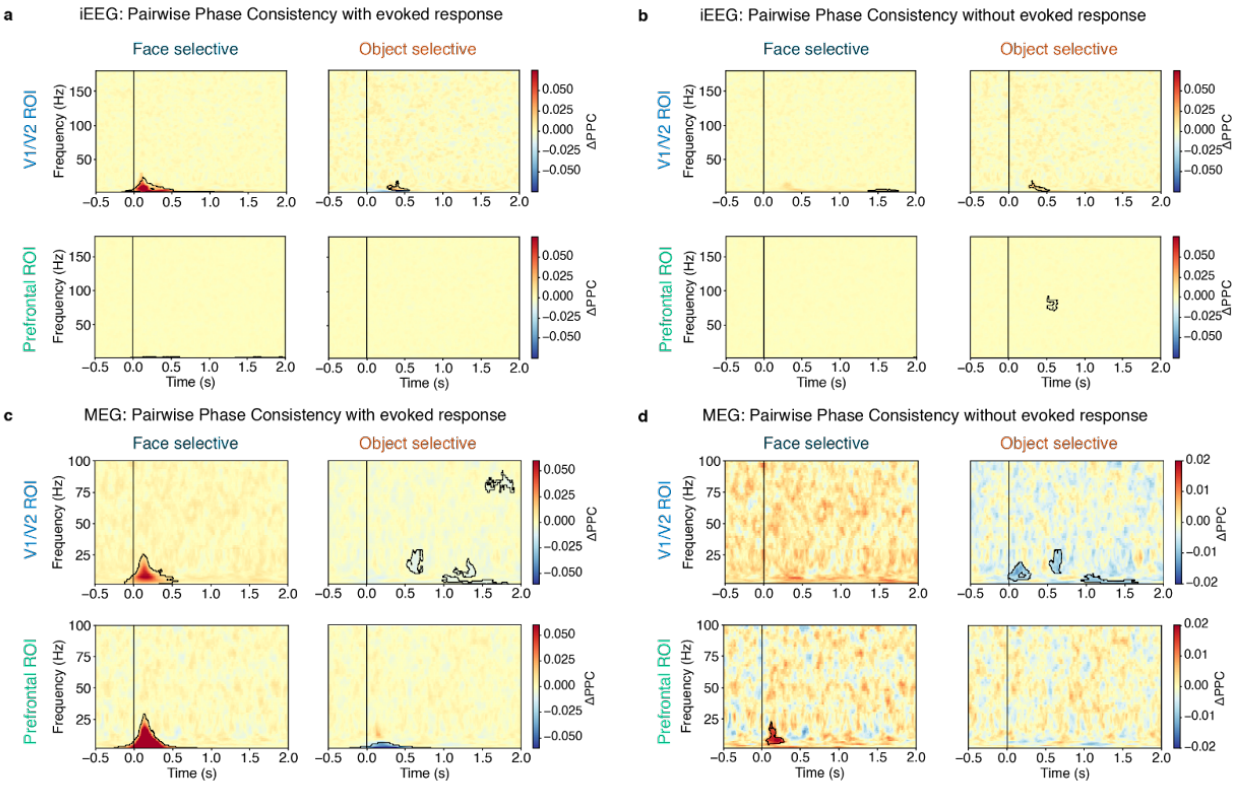

**Supplementary Figure 47**. Results of the PPC analysis on the combined task relevant and task irrelevant trials, before and after removing the evoked response on iEEG and MEG source data. **a-b**. iEEG results remained consistent with those found when analyzing the task irrelevant and task relevant conditions separately. **c-d**. MEG results were also consistent with those obtained from separate analyses on task irrelevant and task relevant data.

#### Exploratory analyses: DFC

Following the DFC analysis reported in the main text for task irrelevant trials and above for task relevant trials, we ran the same analysis on the combined task relevant and task irrelevant trials to increase statistical power (Supplementary Figure 48). In iEEG, we observed significant connectivity between face-selective electrodes and V1/V2. Connectivity was sustained up to 1 second after stimulus onset and present across several frequency bands, most predominantly in the high-gamma band between 70-100 Hz. Significant connectivity between object-selective electrodes and V1/V2 was also observed over a broad frequency range (e.g., > 30Hz), but only up to 0.5 s after stimulus onset. Significant, content-specific connectivity between face-selective electrodes and PFC was also observed, spanning a range of frequencies from the beta band up to the HG band, which was also extended in time up to 1 second after stimulus onset. Again, and in contrast to face-selective nodes, DFC between object-selective electrodes and PFC was spottier and briefer. After regressing out the evoked response, these effects remained largely consistent (Supplementary Figure 48b).

In MEG, the results of the DFC analysis indicated a significant difference between the face-selective node and both PFC and V1/V2 ROIs within the initial 0.5-s time window (Supplementary Figure 48c). Similar to our findings in the task relevant condition, we observed a significant increase in content-specific connectivity in the low-frequency range (i.e., DFC for faces in the face-selective node), accompanied by a decrease in content-specific connectivity in the high-frequency range (i.e., DFC for objects in the face-selective node). In addition, smaller yet significant changes in DFC were observed between the object-selective node and both PFC and V1/V2 ROIs during the same time period (0-0.5 s), but this result was difficult to interpret due to the generalized decrease in DFC observed when object stimuli were presented (i.e., indicating stronger DFC for faces in the object-node connectivity).

When we conducted the DFC analysis after removing the evoked response, the significant effects in the low frequency bands persisted, while most of the high-frequency modulations disappeared (Supplementary Figure 48d). A significant modulation in DFC was observed between the face-selective node and both PFC and V1/V2 ROIs in the alpha band up to 0.5 s. Additionally, the transient low-frequency object-selective DFC that was found before with V1/V2 in object selective nodes disappeared once the evoked response was removed.

Overall, the combined analysis of task relevant and task irrelevant conditions in the MEG data replicated our main findings from task irrelevant trials, with a pronounced modulation of DFC between face-selective node and both PFC and V1/V2 in the low-frequency range in the first 0.5 s from stimulus presentation.

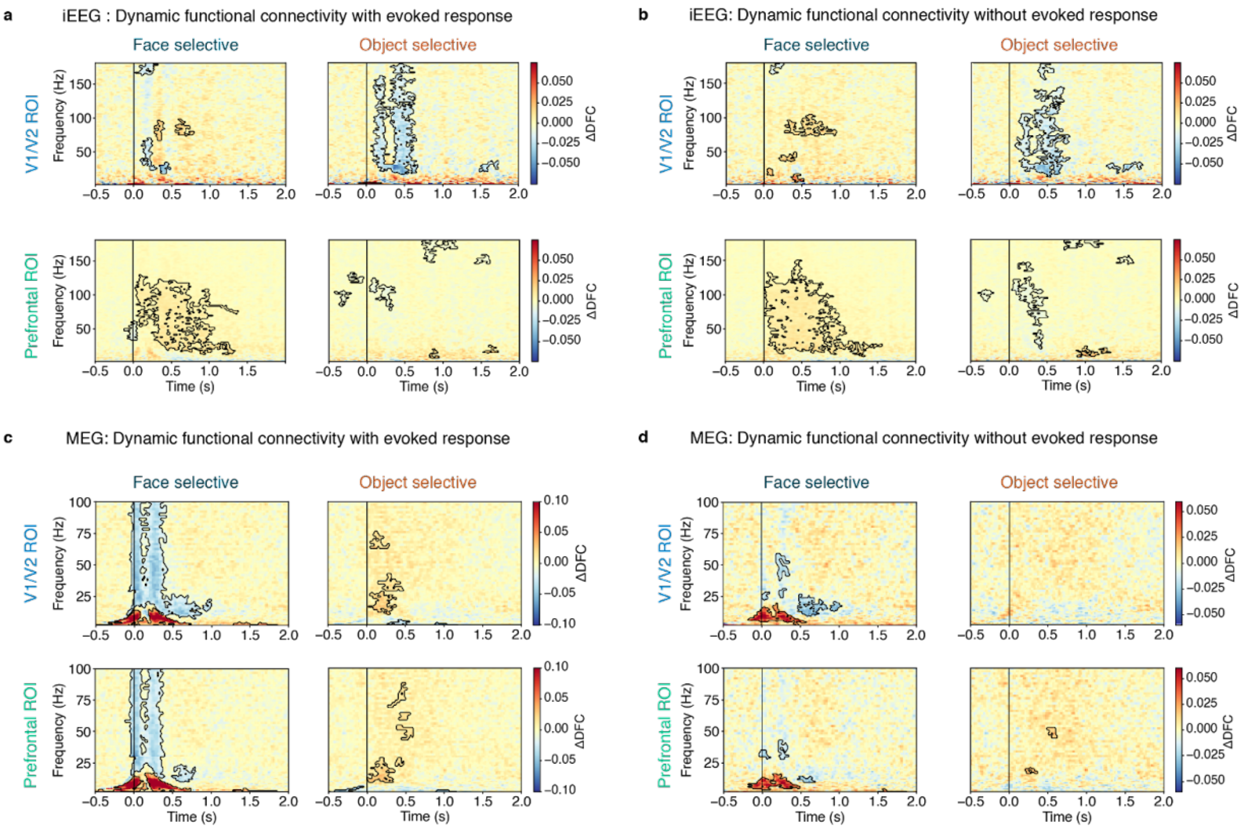

**Supplementary Figure 48**. Results of the DFC analysis on combined task relevant and task irrelevant trials before and after removing the evoked response on iEEG and MEG source data. **a-b**. iEEG results remained consistent with those performed on task irrelevant and task relevant, separately. **c-d**. MEG results were also consistent with those obtained from separate analyses on task irrelevant and task relevant data.

### Putative Neural Correlates of Consciousness (pNCC)

#### Pre-registered analyses

##### Univariate pNCC analysis

In the main text, we presented the results of the pNCC analysis (Figure 5). Here, we describe the full results for both the activation findings (Supplementary Table 20) and the deactivation ones (Supplementary Table 21). These tables detail the count of voxels detected by the analysis in each of the anatomical ROIs that were a priori defined by the proponents of the theories (see Extended Data Table 2).

| **Anatomical ROIs (Destrieux atlas)** | | **Face** | | **Object** | | **Letter** | | **False Font** | |
|---|---|---|---|---|---|---|---|---|---|
| **Theory** | **Short name** | **n voxels** | **% voxels** | **n voxels** | **% voxels** | **n voxels** | **% voxels** | **n voxels** | **% voxels** |
| **Posterior ROIs** | | | | | | | | | |
| IIT | G_and_S_occipital_inf | 1897 | 94.57 | 1934 | 96.41 | 1717 | 85.59 | 1798 | 89.63 |
| | G_oc-temp_lat-fusifor | 2144 | 82.91 | 2306 | 89.17 | 1491 | 57.66 | 1838 | 71.08 |
| | G_occipital_middle | 797 | 32.37 | 1379 | 56.01 | 1045 | 42.45 | 1246 | 50.61 |
| | S_oc_middle_and_Lunatus | 646 | 63.96 | 931 | 92.18 | 832 | 82.38 | 889 | 88.02 |
| | G_cuneus | 100 | 3.99 | 60 | 2.39 | 38 | 1.52 | 21 | 0.84 |
| | G_occipital_sup | 71 | 3.62 | 299 | 15.23 | 233 | 11.87 | 369 | 18.80 |
| | G_oc-temp_med-Lingual | 377 | 12.56 | 520 | 17.32 | 326 | 10.86 | 309 | 10.29 |
| | G_oc-temp_med-Parahip | 168 | 11.94 | 321 | 22.81 | 0 | 0.00 | 36 | 2.56 |
| | G_temporal_inf | 83 | 5.73 | 296 | 20.44 | 337 | 23.27 | 472 | 32.60 |
| | Pole_occipital | 2105 | 87.34 | 2039 | 84.61 | 1352 | 56.10 | 1278 | 53.03 |
| | S_calcarine | 58 | 2.39 | 67 | 2.76 | 25 | 1.03 | 30 | 1.24 |
| | S_intrapariet_and_P_trans | 0 | 0.00 | 243 | 6.41 | 769 | 20.27 | 1107 | 29.19 |
| | S_oc_sup_and_transversal | 22 | 1.55 | 1118 | 79.01 | 807 | 57.03 | 1025 | 72.44 |
| | S_temporal_sup | 13 | 0.26 | 3 | 0.06 | 1 | 0.02 | 4 | 0.08 |
| IIT extended | G_orbital | 74 | 4.87 | 39 | 2.57 | 0 | 0.00 | 0 | 0.00 |
| | G_pariet_inf-Angular | 0 | 0.00 | 15 | 0.43 | 71 | 2.04 | 56 | 1.61 |
| | G_pariet_inf-Supramar | 0 | 0.00 | 0 | 0.00 | 1 | 0.02 | 58 | 1.34 |
| | G_precentral | 87 | 2.07 | 30 | 0.72 | 77 | 1.84 | 59 | 1.41 |
| | G_temporal_middle | 2 | 0.06 | 5 | 0.14 | 7 | 0.20 | 19 | 0.53 |
| | S_occipital_ant | 270 | 36.83 | 433 | 59.07 | 468 | 63.85 | 516 | 70.40 |
| | S_oc-temp_lat | 682 | 58.79 | 820 | 70.69 | 781 | 67.33 | 876 | 75.52 |
| | S_precentral-inf-part | 0 | 0.00 | 0 | 0.00 | 235 | 10.63 | 136 | 6.15 |
| | S_temporal_inf | 23 | 1.24 | 97 | 5.21 | 101 | 5.43 | 146 | 7.85 |
| **PFC ROIs** | | | | | | | | | |
| GNWT, IIT extended | S_front_inf | 167 | 8.04 | 0 | 0.00 | 209 | 10.06 | 1 | 0.05 |
| GNWT | Lat_Fis-ant-Horizont | 0 | 0.00 | 5 | 0.91 | 0 | 0.00 | 0 | 0.00 |
| GNWT, IIT excluded | G_front_inf-Opercular | 43 | 1.92 | 0 | 0.00 | 66 | 2.95 | 0 | 0.00 |
| | G_front_inf-Orbital | 4 | 0.64 | 0 | 0.00 | 0 | 0.00 | 0 | 0.00 |
| | G_front_inf-Triangul | 8 | 0.49 | 0 | 0.00 | 20 | 1.21 | 0 | 0.00 |
| | G_front_middle | 19 | 0.31 | 0 | 0.00 | 71 | 1.16 | 0 | 0.00 |
| | S_front_sup | 0 | 0.00 | 0 | 0.00 | 8 | 0.21 | 0 | 0.00 |
| IIT excluded | G_front_sup | 21 | 0.20 | 11 | 0.11 | 123 | 1.18 | 1 | 0.01 |
| | G_subcallosal | 31 | 4.51 | 0 | 0.00 | 0 | 0.00 | 0 | 0.00 |
| | S_orbital_lateral | 7 | 1.32 | 0 | 0.00 | 0 | 0.00 | 0 | 0.00 |
| | S_orbital_med-olfact | 4 | 0.89 | 0 | 0.00 | 0 | 0.00 | 0 | 0.00 |
| | S_orbital-H_Shaped | 242 | 17.97 | 129 | 9.58 | 0 | 0.00 | 16 | 1.19 |

**Supplementary Table 20**. Results from the univariate fMRI contrast-conjunction pNCC analysis, aimed at "ruling in" putative NCCs, reporting activation in response to the presentation of stimuli, regardless of their relevance. For each of the predefined anatomical ROIs, we count the number of voxels showing activation, and calculate their proportion with respect to the total number of voxels of the ROI.

| Anatomical ROIs (Destrieux atlas) | | Face | | Object | | Letter | | False Font | |
|---|---|---|---|---|---|---|---|---|---|
| Theory | Short name | n voxels | % voxels | n voxels | % voxels | n voxels | % voxels | n voxels | % voxels |
| **Posterior ROIs** | | | | | | | | | |
| IIT | G_oc-temp_lat-fusifor | 1 | 0.04 | 0 | 0.00 | 0 | 0.00 | 0 | 0.00 |
| | G_occipital_middle | 258 | 10.48 | 81 | 3.29 | 203 | 8.25 | 180 | 7.31 |
| | S_oc_middle_and_Lunatus | 0 | 0.00 | 0 | 0.00 | 1 | 0.10 | 0 | 0.00 |
| | G_cuneus | 954 | 38.05 | 1609 | 64.18 | 1589 | 63.38 | 1474 | 58.80 |
| | G_occipital_sup | 676 | 34.44 | 601 | 30.62 | 523 | 26.64 | 586 | 29.85 |
| | G_oc-temp_med-Lingual | 727 | 24.22 | 1010 | 33.64 | 1024 | 34.11 | 1194 | 39.77 |
| | G_oc-temp_med-Parahip | 0 | 0.00 | 0 | 0.00 | 6 | 0.43 | 0 | 0.00 |
| | G_temporal_inf | 0 | 0.00 | 13 | 0.90 | 6 | 0.41 | 0 | 0.00 |
| | Pole_occipital | 1 | 0.04 | 1 | 0.04 | 0 | 0.00 | 2 | 0.08 |
| | S_calcarine | 406 | 16.74 | 671 | 27.66 | 739 | 30.46 | 570 | 23.50 |
| | S_intrapariet_and_P_trans | 65 | 1.71 | 235 | 6.20 | 154 | 4.06 | 6 | 0.16 |
| | S_oc_sup_and_transversal | 91 | 6.43 | 62 | 4.38 | 43 | 3.04 | 60 | 4.24 |
| | S_temporal_sup | 197 | 3.99 | 878 | 17.78 | 911 | 18.45 | 1526 | 30.90 |
| IIT extended | G_orbital | 28 | 1.84 | 92 | 6.06 | 80 | 5.27 | 101 | 6.65 |
| | G_pariet_inf-Angular | 293 | 8.40 | 740 | 21.22 | 509 | 14.60 | 806 | 23.11 |
| | G_pariet_inf-Supramar | 706 | 16.34 | 629 | 14.56 | 556 | 12.87 | 900 | 20.83 |
| | G_precentral | 6 | 0.14 | 111 | 2.65 | 89 | 2.12 | 41 | 0.98 |
| | G_temp_sup-Lateral | 42 | 1.11 | 31 | 0.82 | 198 | 5.21 | 355 | 9.35 |
| | G_temp_sup-Plan_tempo | 273 | 16.27 | 79 | 4.71 | 45 | 2.68 | 357 | 21.28 |
| | G_temporal_middle | 23 | 0.64 | 272 | 7.61 | 408 | 11.41 | 592 | 16.55 |
| | S_interm_prim-Jensen | 335 | 41.72 | 437 | 54.42 | 416 | 51.81 | 359 | 44.71 |
| | S_precentral-inf-part | 117 | 5.29 | 698 | 31.57 | 668 | 30.21 | 344 | 15.56 |
| | S_temporal_inf | 0 | 0.00 | 14 | 0.75 | 55 | 2.96 | 9 | 0.48 |
| **PFC ROIs** | | | | | | | | | |
| GNWT, IIT extended | S_front_inf | 62 | 2.99 | 314 | 15.12 | 428 | 20.61 | 193 | 9.29 |
| GNWT | G_and_S_cingul-Mid-Post | 47 | 2.44 | 74 | 3.85 | 73 | 3.80 | 182 | 9.46 |
| | Lat_Fis-ant-Horizont | 0 | 0.00 | 200 | 36.50 | 223 | 40.69 | 79 | 14.42 |
| | Lat_Fis-ant-Vertical | 79 | 17.14 | 179 | 38.83 | 206 | 44.69 | 192 | 41.65 |
| GNWT, IIT excluded | G_and_S_cingul-Ant | 313 | 9.41 | 339 | 10.19 | 359 | 10.79 | 376 | 11.30 |
| | G_and_S_cingul-Mid-Ant | 3 | 0.15 | 130 | 6.60 | 93 | 4.72 | 34 | 1.73 |
| | G_front_inf-Opercular | 183 | 8.19 | 562 | 25.16 | 573 | 25.65 | 427 | 19.11 |
| | G_front_inf-Orbital | 3 | 0.48 | 56 | 9.00 | 29 | 4.66 | 46 | 7.40 |
| | G_front_inf-Triangul | 37 | 2.24 | 280 | 16.98 | 398 | 24.14 | 292 | 17.71 |
| | G_front_middle | 414 | 6.74 | 1105 | 17.98 | 956 | 15.55 | 1409 | 22.93 |
| | S_front_middle | 9 | 0.41 | 64 | 2.95 | 29 | 1.34 | 91 | 4.19 |
| | S_front_sup | 503 | 13.46 | 581 | 15.55 | 463 | 12.39 | 693 | 18.54 |
| IIT excluded | G_and_S_frontomargin | 29 | 3.70 | 81 | 10.33 | 44 | 5.61 | 61 | 7.78 |
| | G_and_S_transv_frontopol | 79 | 7.66 | 118 | 11.45 | 134 | 13.00 | 198 | 19.20 |
| | G_front_sup | 281 | 2.70 | 746 | 7.16 | 666 | 6.39 | 929 | 8.92 |
| | G_subcallosal | 111 | 16.16 | 44 | 6.40 | 64 | 9.32 | 97 | 14.12 |
| | S_orbital_lateral | 0 | 0.00 | 88 | 16.57 | 88 | 16.57 | 130 | 24.48 |
| | S_orbital_med-olfact | 88 | 19.69 | 56 | 12.53 | 83 | 18.57 | 54 | 12.08 |
| | S_orbital-H_Shaped | 1 | 0.07 | 0 | 0.00 | 0 | 0.00 | 0 | 0.00 |
| | S_suborbital | 2 | 0.28 | 0 | 0.00 | 0 | 0.00 | 0 | 0.00 |

**Supplementary Table 21**. Results of the univariate fMRI contrast-conjunction pNCC analysis reported in Supplementary Table 20 (with the same conventions), but for voxels showing deactivation rather than activation.

The pNCC analysis identifies visually responsive areas, while ruling out areas that are responsive to aspects of the task. Namely, we focused on two types of areas: First, areas that are sensitive to task goal, showing greater activity for task relevant targets vs. baseline, and no differential activity for non-targets vs. baseline (blank ITIs) (defined by the following contrast: [targets > bsl & task relevant = bsl & task irrelevant = bsl]). Second, areas that are sensitive to task-relevance, expected to be responsive to all task relevant stimuli, but not to task irrelevant stimuli (defined by the following contrast: [targets > bsl & task relevant ≠ bsl & task irrelevant = bsl]. Areas that are sensitive to task goals and task relevance are listed in Supplementary Table 22 (see also Extended Data Figure 9a for a visual display of the results).

| **Anatomical ROIs (Destrieux atlas)** | | **Task goals** | | **Task relevance** | |
|---|---|---|---|---|---|
| **Theory** | **Short name** | **n voxels** | **% voxels** | **n voxels** | **% voxels** |
| **Posterior ROIs** | | | | | |
| IIT | G_occipital_middle | 20 | 0.81 | 4 | 0.16 |
| | S_oc_middle_and_Lunatus | 3 | 0.30 | 0 | 0.00 |
| | G_cuneus | 33 | 1.32 | 0 | 0.00 |
| | G_occipital_sup | 3 | 0.15 | 0 | 0.00 |
| | G_oc-temp_med-Lingual | 69 | 2.30 | 0 | 0.00 |
| | G_oc-temp_med-Parahip | 30 | 2.13 | 0 | 0.00 |
| | G_temporal_inf | 93 | 6.42 | 42 | 2.90 |
| | Pole_occipital | 16 | 0.66 | 0 | 0.00 |
| | S_calcarine | 55 | 2.27 | 0 | 0.00 |
| | S_intrapariet_and_P_trans | 90 | 2.37 | 630 | 16.61 |
| | S_oc_sup_and_transversal | 5 | 0.35 | 19 | 1.34 |
| | S_temporal_sup | 168 | 3.40 | 7 | 0.14 |
| IIT extended | G_orbital | 13 | 0.86 | 28 | 1.84 |
| | G_pariet_inf-Angular | 77 | 2.21 | 109 | 3.13 |
| | G_pariet_inf-Supramar | 219 | 5.07 | 237 | 5.49 |
| | G_precentral | 725 | 17.29 | 27 | 0.64 |
| | G_temp_sup-Lateral | 12 | 0.32 | 3 | 0.08 |
| | G_temp_sup-Plan_tempo | 0 | 0.00 | 9 | 0.54 |
| | G_temporal_middle | 100 | 2.80 | 4 | 0.11 |
| | S_interm_prim-Jensen | 93 | 11.58 | 0 | 0.00 |
| | S_occipital_ant | 18 | 2.46 | 0 | 0.00 |
| | S_oc-temp_lat | 7 | 0.60 | 10 | 0.86 |
| | S_precentral-inf-part | 82 | 3.71 | 114 | 5.16 |

| | | | | | |
|---|---|---|---|---|---|
| | S_temporal_inf | 73 | 3.92 | 12 | 0.64 |
| **PFC ROIs** | | | | | |
| GNWT, IIT extended | S_front_inf | 161 | 7.75 | 61 | 2.94 |
| GNWT | G_and_S_cingul-Mid-Post | 301 | 15.65 | 86 | 4.47 |
| | Lat_Fis-ant-Horizont | 24 | 4.38 | 0 | 0.00 |
| | Lat_Fis-ant-Vertical | 27 | 5.86 | 0 | 0.00 |
| GNWT, IIT excluded | G_and_S_cingul-Ant | 109 | 3.28 | 25 | 0.75 |
| | G_and_S_cingul-Mid-Ant | 374 | 18.99 | 141 | 7.16 |
| | G_front_inf-Opercular | 584 | 26.14 | 19 | 0.85 |
| | G_front_inf-Orbital | 16 | 2.57 | 0 | 0.00 |
| | G_front_inf-Triangul | 19 | 1.15 | 9 | 0.55 |
| | G_front_middle | 375 | 6.10 | 110 | 1.79 |
| | S_front_middle | 378 | 17.42 | 1 | 0.05 |
| | S_front_sup | 102 | 2.73 | 3 | 0.08 |
| IIT excluded | G_and_S_frontomargin | 119 | 15.18 | 6 | 0.77 |
| | G_front_sup | 1033 | 9.92 | 154 | 1.48 |
| | G_subcallosal | 22 | 3.20 | 1 | 0.15 |
| | S_orbital_lateral | 10 | 1.88 | 0 | 0.00 |
| | S_orbital_med-olfact | 11 | 2.46 | 0 | 0.00 |
| | S_orbital-H_Shaped | 108 | 8.02 | 0 | 0.00 |
| | S_suborbital | 4 | 0.15 | 0 | 0.00 |

**Supplementary Table 22**. Results from the univariate fMRI contrast-conjunction pNCC analysis, describing areas responsive to task goals and task relevance. The same conventions from Supplementary Table 20 are used here.

As expected, the two conjunction analyses designed to “rule out” areas responsive to task goals and task relevance identified several regions in the PFC ROIs (most prominently in inferior, middle and superior frontal gyrus, but also in cingulate cortex and others). Interestingly, these conjunctions also identified regions in posterior ROIs (e.g., inferior temporal gyrus, supramarginal gyrus and intraparietal sulcus). All of the voxels detected by these conjunctions were excluded from the reported putative NCCs.

#### Individual Z-maps

To compute the above conjunctions, we first created thresholded individual z maps by contrasting the presence of stimuli vs. baseline (corrected for multiple comparisons, using gaussian random-field cluster thresholding, with a cluster formation threshold of one-sided $p < 0.001$ ($z \geq 3.1$,) and a cluster significance threshold of $p < 0.05$) in the task relevant and irrelevant conditions separately. These maps are presented in Supplementary Figure 49. As can be seen, besides effects located in the visual cortex and other posterior regions, the presentation of stimuli elicits both activations and deactivations in several prefrontal areas for both task conditions.

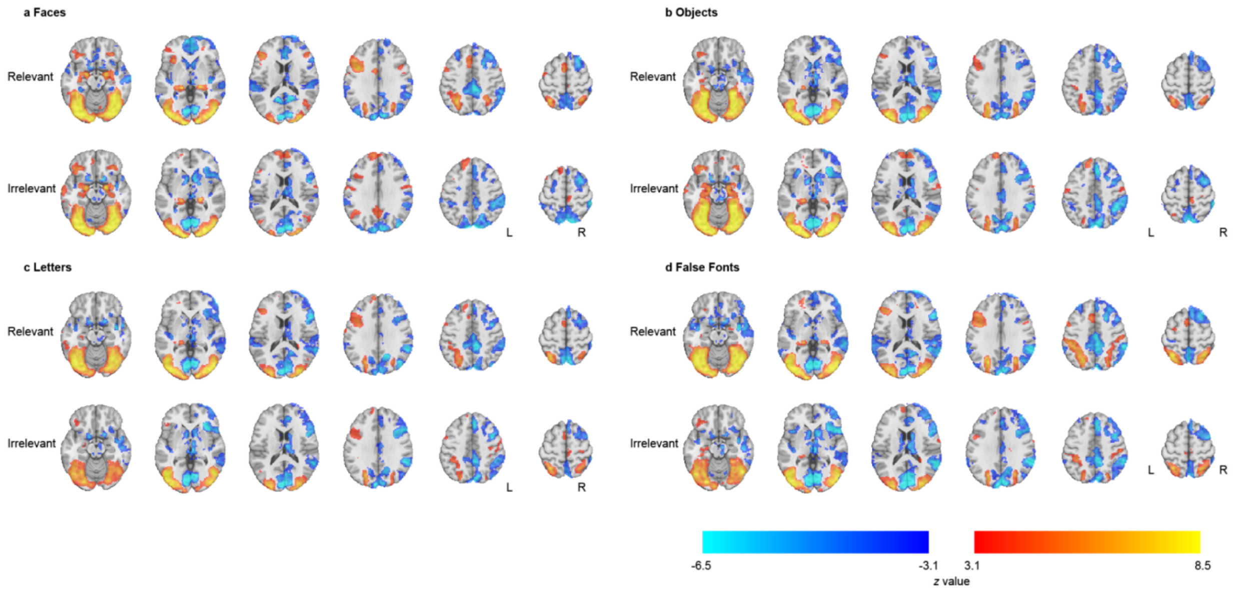

**Supplementary Figure 49**. Contrast of parameter estimates (stimulus vs. baseline) z maps used in the conjunction that identifies the putative NCCs (main text Figure 5). Here we show z maps for each stimulus category (**a**. faces; **b**. objects; **c**. letters; **d**. false fonts) and condition (top, Relevant; bottom, Irrelevant).

#### Multivariate pNCC analysis

The pNCC analysis was further conducted taking a multivariate approach (Extended Data Figure 9b). Below in Supplementary Table 23 we provide the details of this analysis, akin to the univariate analysis reported above.

| **Anatomical ROIs (Destrieux atlas)** | | **Face** | | **Object** | | **Letter** | | **False Font** | |
|---|---|---|---|---|---|---|---|---|---|
| **Theory** | **Short name** | **n voxels** | **% voxels** | **n voxels** | **% voxels** | **n voxels** | **% voxels** | **n voxels** | **% voxels** |
| **Posterior ROIs** | | | | | | | | | |
| IIT | G_and_S_occipital_inf | 123 | 6.13 | 71 | 3.54 | 18 | 0.90 | 118 | 5.88 |
| | G_oc-temp_lat-fusifor | 104 | 4.02 | 19 | 0.73 | 15 | 0.58 | 17 | 0.66 |
| | G_occipital_middle | 128 | 5.20 | 109 | 4.43 | 87 | 3.53 | 213 | 8.65 |
| | S_oc_middle_and_Lunatus | 139 | 13.76 | 108 | 10.69 | 44 | 4.36 | 107 | 10.59 |
| | G_cuneus | 429 | 17.11 | 346 | 13.80 | 433 | 17.27 | 355 | 14.16 |
| | G_occipital_sup | 598 | 30.46 | 563 | 28.68 | 397 | 20.22 | 512 | 26.08 |
| | G_oc-temp_med-Lingual | 878 | 29.25 | 560 | 18.65 | 524 | 17.46 | 580 | 19.32 |
| | G_temporal_inf | 1 | 0.07 | 2 | 0.14 | 6 | 0.41 | 51 | 3.52 |
| | Pole_occipital | 1085 | 45.02 | 656 | 27.22 | 374 | 15.52 | 391 | 16.22 |
| | S_calcarine | 310 | 12.78 | 298 | 12.28 | 287 | 11.83 | 241 | 9.93 |
| | S_intrapariet_and_P_trans | 0 | 0.00 | 12 | 0.32 | 64 | 1.69 | 32 | 0.84 |
| | S_oc_sup_and_transversal | 175 | 12.37 | 192 | 13.57 | 149 | 10.53 | 341 | 24.10 |
| | S_temporal_sup | 2 | 0.04 | 29 | 0.59 | 39 | 0.79 | 40 | 0.81 |

| | | | | | | | | | |
|---|---|---|---|---|---|---|---|---|---|
| IIT extended | G_pariet_inf-Angular | 0 | 0.00 | 36 | 1.03 | 35 | 1.00 | 68 | 1.95 |
| | G_pariet_inf-Supramar | 0 | 0.00 | 15 | 0.35 | 20 | 0.46 | 55 | 1.27 |
| | G_precentral | 0 | 0.00 | 11 | 0.26 | 37 | 0.88 | 50 | 1.19 |
| | G_temp_sup-Lateral | 0 | 0.00 | 0 | 0.00 | 0 | 0.00 | 24 | 0.63 |
| | G_temporal_middle | 0 | 0.00 | 2 | 0.06 | 52 | 1.45 | 15 | 0.42 |
| | S_interm_prim-Jensen | 0 | 0.00 | 3 | 0.37 | 35 | 4.36 | 2 | 0.25 |
| | S_occipital_ant | 3 | 0.41 | 0 | 0.00 | 1 | 0.14 | 9 | 1.23 |
| | S_oc-temp_lat | 0 | 0.00 | 0 | 0.00 | 0 | 0.00 | 29 | 2.50 |
| | S_precentral-inf-part | 0 | 0.00 | 49 | 2.22 | 235 | 10.63 | 174 | 7.87 |
| | S_temporal_inf | 0 | 0.00 | 0 | 0.00 | 11 | 0.59 | 23 | 1.24 |
| **PFC ROIs** | | | | | | | | | |
| GNWT, IIT extended | S_front_inf | 0 | 0.00 | 17 | 0.82 | 136 | 6.55 | 17 | 0.82 |
| GNWT | Lat_Fis-ant-Horizont | 0 | 0.00 | 15 | 2.74 | 0 | 0.00 | 0 | 0.00 |
| | Lat_Fis-ant-Vertical | 0 | 0.00 | 3 | 0.65 | 0 | 0.00 | 0 | 0.00 |
| GNWT, IIT excluded | G_and_S_cingul-Ant | 0 | 0.00 | 0 | 0.00 | 6 | 0.18 | 0 | 0.00 |
| | G_front_inf-Opercular | 1 | 0.04 | 66 | 2.95 | 114 | 5.10 | 96 | 4.30 |
| | G_front_inf-Triangul | 2 | 0.12 | 4 | 0.24 | 43 | 2.61 | 64 | 3.88 |
| | G_front_middle | 1 | 0.02 | 27 | 0.44 | 82 | 1.33 | 37 | 0.60 |
| | S_front_middle | 0 | 0.00 | 0 | 0.00 | 2 | 0.09 | 0 | 0.00 |
| | S_front_sup | 0 | 0.00 | 0 | 0.00 | 3 | 0.08 | 0 | 0.00 |
| IIT excluded | G_front_sup | 19 | 0.18 | 0 | 0.00 | 64 | 0.61 | 0 | 0.00 |

**Supplementary Table 23**. Results of the multivariate fMRI conjunction pNCC analysis. The same conventions from Supplementary Table 20 are used here.

#### Subject-level pNCC analysis

The preregistered univariate pNCC analysis was aimed at testing the theories' predictions about the areas potentially subserving conscious processing. A critical point of disagreement here pertains to prefrontal areas, where GNWT predicts they should be included in the resulting pNCCs, while IIT claims otherwise. However, the global workspace is held to be widely distributed in prefrontal and parietal areas[8], in a manner that might be idiosyncratic to a specific participant. Thus, a group level analysis might fail to detect prefrontal pNCCs even if they do exist. To account for this possibility, we complemented our analysis with an additional univariate pNCC analysis, performed at the subject level. Supplementary Table 24 shows the proportion of subjects that passed the conjunction analysis for each stimulus category in each a priori defined anatomical ROI.

As the results show, besides the majority of subjects showing activations in posterior ROIs, there is some agreement also around activations in prefrontal regions. For example, more than 10% of subjects showed activations in superior frontal gyrus across all stimulus categories. Similar results were also found in inferior and middle frontal gyri. Interestingly, these areas showed a considerable proportion of subjects also showing deactivations, e.g., in superior and middle frontal gyrus, more than 30% of subjects show deactivations in all stimulus categories, and 24% subjects showing the same in inferior frontal gyrus.

It should be noted that the conjunction analyses used here implement a logical AND operation between two maps that were corrected for multiple comparisons to have an alpha level of 0.05.

Therefore, the conjunction test is conservative, as assuming independence of the maps, the alpha level of the conjunction is 0.0025.

| Anatomical ROIs (Destrieux atlas) | | Activation (% subjects) | | | | Deactivation (% subjects) | | | |
|---|---|---|---|---|---|---|---|---|---|
| Theory | Short name | Face | Object | Letter | False Font | Face | Object | Letter | False Font |
| IIT | G_and_S_occipital_inf | 94.52 | 94.52 | 89.04 | 94.52 | 1.37 | 4.11 | 6.85 | 6.85 |
| | G_oc-temp_lat-fusifor | 94.52 | 91.78 | 68.49 | 89.04 | 8.22 | 5.48 | 9.59 | 6.85 |
| | G_occipital_middle | 94.52 | 94.52 | 91.78 | 93.15 | 41.10 | 17.81 | 23.29 | 27.40 |
| | S_oc_middle_and_Lunatus | 93.15 | 93.15 | 91.78 | 93.15 | 8.22 | 1.37 | 5.48 | 6.85 |
| | G_cuneus | 83.56 | 72.60 | 56.16 | 46.58 | 72.60 | 75.34 | 56.16 | 68.49 |
| | G_occipital_sup | 89.04 | 90.41 | 86.30 | 87.67 | 71.23 | 65.75 | 52.05 | 64.38 |
| | G_oc-temp_med-Lingual | 94.52 | 94.52 | 83.56 | 83.56 | 64.38 | 64.38 | 49.32 | 50.68 |
| | G_oc-temp_med-Parahip | 53.42 | 68.49 | 2.74 | 12.33 | 1.37 | 0.00 | 2.74 | 0.00 |
| | G_temporal_inf | 71.23 | 60.27 | 54.79 | 79.45 | 17.81 | 13.70 | 12.33 | 15.07 |
| | Pole_occipital | 94.52 | 94.52 | 91.78 | 94.52 | 47.95 | 49.32 | 32.88 | 46.58 |
| | Pole_temporal | 9.59 | 0.00 | 2.74 | 1.37 | 1.37 | 1.37 | 2.74 | 2.74 |
| | S_calcarine | 84.93 | 89.04 | 63.01 | 57.53 | 65.75 | 69.86 | 45.21 | 54.79 |
| | S_intrapariet_and_P_trans | 32.88 | 36.99 | 53.42 | 69.86 | 30.14 | 24.66 | 20.55 | 13.70 |
| | S_oc_sup_and_transversal | 73.97 | 91.78 | 82.19 | 91.78 | 46.58 | 30.14 | 21.92 | 35.62 |
| | S_temporal_sup | 52.05 | 24.66 | 20.55 | 26.03 | 32.88 | 32.88 | 32.88 | 41.10 |
| IIT extended | G_orbital | 21.92 | 8.22 | 4.11 | 1.37 | 17.81 | 19.18 | 15.07 | 17.81 |
| | G_pariet_inf-Angular | 39.73 | 30.14 | 35.62 | 50.68 | 39.73 | 35.62 | 36.99 | 43.84 |
| | G_pariet_inf-Supramar | 9.59 | 8.22 | 12.33 | 30.14 | 30.14 | 30.14 | 27.40 | 38.36 |
| | G_precentral | 26.03 | 15.07 | 13.70 | 21.92 | 15.07 | 21.92 | 24.66 | 23.29 |
| | G_temp_sup-Lateral | 17.81 | 2.74 | 5.48 | 2.74 | 23.29 | 28.77 | 28.77 | 35.62 |
| | G_temp_sup-Plan_tempo | 4.11 | 4.11 | 4.11 | 1.37 | 17.81 | 24.66 | 17.81 | 19.18 |
| | G_temporal_middle | 53.42 | 32.88 | 38.36 | 58.90 | 35.62 | 26.03 | 34.25 | 38.36 |
| | S_interm_prim-Jensen | 1.37 | 1.37 | 2.74 | 5.48 | 23.29 | 21.92 | 20.55 | 23.29 |
| | S_occipital_ant | 84.93 | 86.30 | 78.08 | 94.52 | 5.48 | 4.11 | 2.74 | 6.85 |
| | S_oc-temp_lat | 91.78 | 86.30 | 67.12 | 89.04 | 1.37 | 2.74 | 2.74 | 2.74 |
| | S_precentral-inf-part | 26.03 | 9.59 | 23.29 | 19.18 | 23.29 | 34.25 | 31.51 | 27.40 |
| | S_temporal_inf | 60.27 | 64.38 | 60.27 | 82.19 | 15.07 | 13.70 | 17.81 | 15.07 |
| GNWT, IIT extended | S_front_inf | 27.40 | 13.70 | 21.92 | 17.81 | 23.29 | 31.51 | 31.51 | 27.40 |
| GNWT | G_and_S_cingul-Mid-Post | 0.00 | 0.00 | 0.00 | 0.00 | 8.22 | 10.96 | 12.33 | 8.22 |
| | Lat_Fis-ant-Horizont | 4.11 | 5.48 | 5.48 | 2.74 | 10.96 | 16.44 | 19.18 | 12.33 |
| | Lat_Fis-ant-Vertical | 1.37 | 4.11 | 4.11 | 1.37 | 16.44 | 21.92 | 21.92 | 17.81 |
| GNWT, IIT excluded | G_and_S_cingul-Ant | 5.48 | 2.74 | 4.11 | 5.48 | 20.55 | 19.18 | 12.33 | 16.44 |
| | G_and_S_cingul-Mid-Ant | 1.37 | 1.37 | 4.11 | 2.74 | 8.22 | 15.07 | 9.59 | 6.85 |
| | G_front_inf-Opercular | 23.29 | 9.59 | 13.70 | 12.33 | 24.66 | 38.36 | 31.51 | 32.88 |
| | G_front_inf-Orbital | 12.33 | 6.85 | 4.11 | 2.74 | 13.70 | 17.81 | 15.07 | 16.44 |
| | G_front_inf-Triangul | 17.81 | 6.85 | 12.33 | 12.33 | 24.66 | 26.03 | 28.77 | 32.88 |
| | G_front_middle | 28.77 | 13.70 | 19.18 | 17.81 | 34.25 | 39.73 | 36.99 | 34.25 |
| | S_front_middle | 6.85 | 4.11 | 6.85 | 2.74 | 19.18 | 19.18 | 13.70 | 21.92 |
| | S_front_sup | 12.33 | 6.85 | 4.11 | 4.11 | 24.66 | 31.51 | 23.29 | 24.66 |
| IIT excluded | G_and_S_frontomargin | 8.22 | 2.74 | 4.11 | 2.74 | 10.96 | 16.44 | 9.59 | 17.81 |
| | G_and_S_transv_frontopol | 6.85 | 2.74 | 5.48 | 4.11 | 24.66 | 16.44 | 12.33 | 17.81 |
| | G_front_sup | 15.07 | 10.96 | 16.44 | 12.33 | 36.99 | 41.10 | 31.51 | 36.99 |
| | G_rectus | 8.22 | 1.37 | 2.74 | 0.00 | 0.00 | 0.00 | 1.37 | 2.74 |
| | G_subcallosal | 4.11 | 1.37 | 0.00 | 0.00 | 4.11 | 8.22 | 2.74 | 5.48 |
| | S_orbital_lateral | 6.85 | 5.48 | 4.11 | 4.11 | 13.70 | 12.33 | 12.33 | 16.44 |
| | S_orbital_med-olfact | 9.59 | 0.00 | 0.00 | 0.00 | 5.48 | 5.48 | 1.37 | 4.11 |
| | S_orbital-H_Shaped | 23.29 | 10.96 | 2.74 | 4.11 | 5.48 | 8.22 | 6.85 | 5.48 |
| | S_suborbital | 12.33 | 1.37 | 1.37 | 1.37 | 2.74 | 0.00 | 2.74 | 5.48 |

**Supplementary Table 24**. Results of the univariate fMRI contrast-conjunction pNCC analysis, performed at the subject level. The same conventions from Supplementary Table 20 are used here.

### Supplementary discussion: IIT proponents

The present results failed to confirm IIT's prediction about sustained, content-specific synchrony in the gamma range between category-specific cortical regions and V1/V2. By IIT, when we see, say, a face with its contours, features, and location in space, there should be 'causal relations' among all the units contributing those contents. Relations require an overlap between the intrinsic causes and effects of those units,[9,10] and those are likely to result in increased synchrony among active units, hence the prediction. Given the lack of evidence for sustained synchrony in the gamma band, the prediction may be wrong with respect to the frequency range. Synchrony may occur instead in lower frequency ranges that may be more sensitive to broader cortical interactions (see e.g.[11]). Indeed, main results using iEEG (Figure 4b) show increased content-specific synchrony (PPC) between category-specific cortical regions and V1/V2 (but not with PFC) in the 2-25 Hz frequency range between 0 and ~750 ms post-stimulus onset.

The failure to confirm sustained synchrony in the gamma range with iEEG in the present study may also stem from technical limitations. As pointed out in the main text, there were only 12 iEEG electrodes in V1/V2 (against 472 in PFC), with only a minority showing sustained activity for longer trials – which was required to include them in the synchrony analysis. Due to limited electrode coverage, the analysis of iEEG data did not strictly follow pre-registered analyses methods, which required restricting measures of synchrony with anatomical V1/V2 to 'category-selective' channels (showing a stronger response to either faces compared to objects or vice versa). Instead, the final analysis selected channels that were merely 'task-responsive' (showing increased activity for either faces or objects compared to baseline). Any overlap between neural responses to faces and objects within higher-order visual areas[12] would be expected to decrease signal-to-noise for the analysis. Results of an ongoing adversarial collaboration replicating the present paradigm using large-scale iEEG recordings along with single-units in monkeys (https://www.templetonworldcharity.org/accelerating-research-consciousness-our-structured-adversarial-collaboration-projects) may help resolve these issues and further test this prediction of IIT.

For technical reasons MEG ROIs for synchrony analyses had to be chosen using spatial filters defined on broad-band ERFs, rather than based on local gamma activity. Because broad-band ERFs likely comprise signals from both activated and deactivated brain areas (e.g. [13,14]), measuring changes in synchrony averaged across such ROIs did not formally test IIT's pre-registered prediction, which only concerns activated areas.

While several findings revealed sustained tracking of duration in iEEG HGP both within V1/V2 and inferior temporal cortex, results of other analyses (especially for orientation RSA and synchrony) failed to identify such a sustained pattern (instead decaying beyond 500 ms). One possible explanation for such discrepancy is that the two latter analyses strongly depend on content-specificity over time. This poses a challenge in the case of V1-V2 units, which contribute low-level stimulus features and were sparsely sampled. Note that micro-saccades appear to be more numerous beyond 500 ms after stimulus onset (Supplementary Figure 6), implying that the perceived location of faces or objects in the visual field may have shifted slightly. If so, IIT predicts that a different set of units in V1/V2 (sensitive to the different retinotopic location) would contribute to experienced low-level features, and thus to representational similarity and synchronization with category-specific units in higher areas. Again, animal experiments with denser iEEG sampling of V1/V2 may be necessary to resolve this issue.

The finding of a few fMRI-activated voxels in the PFC in the pNCC analysis may have various explanations. One possibility, mentioned in the main text, is that such areas may genuinely contribute some content of consciousness (such as some abstract/evaluative/actionable aspect of faces[15-17]). However, the pNCC contrast against baseline is admittedly not very specific (much less than duration-tracking using iEEG). Additionally, none of the activated PFC areas also showed significant fMRI decoding against baseline, further questioning their relevance for consciousness. The weak PFC fMRI activation patterns may also reflect inputs to PFC from posterior cortex or non-specific onset responses (found to be widespread within PFC areas using iEEG, in contrast with absent PFC duration-tracking). Further experiments employing slowly morphing stimuli may shed light on what determines such onset (or offset) responses.

IIT's prediction #1 of less consistent decoding of conscious contents in PFC compared to posterior cortex was verified not only through 1) a failure of decoding orientation in both iEEG and fMRI datasets, but also by 2) a failure of cross-task decoding for letter vs false fonts in PFC using fMRI (Extended Figure 1a); 3) a lack of duration tracking for decoding in PFC using iEEG (Extended Figure 1d-e); and 4) a lack of cross-task iEEG decoding for faces vs. objects using pseudotrials (Extended Figure 1g). In contrast, positive results were consistently found in posterior cortical areas. IIT's prediction of maximum decodability within posterior regions, with no significant additional information about content added by PFC regions, was confirmed using two different multivariate model comparison methods. These results all fit with IIT's prediction that posterior cortical areas are the primary constituents of a complex having maximal integrated information.

Of note, both the sparsity of iEEG electrodes showing duration-tracking (prediction #2) and the widespread deactivation patterns found in PFC (pNCC analysis) strongly suggest that changes in contents of consciousness may be supported by localized activity changes, rather than by a global 'broadcasting' and ignition across the brain. These findings are in line with IIT's prediction that local changes in the state of the substrate of consciousness are sufficient to determine changes in its contents.

### Participants

#### Ethics approvals

The iEEG arm of the study received approval from the New York University Langone Health Office of Science and Research Institutional Review Board (IRB), Harvard's IRBs (including Boston Children's Hospital and Brigham and Women's Hospital) and the IRB for human studies at the University of Wisconsin-Madison. The MEG arm of the study was approved by the Committee for Protecting Human and Animal Subjects in the School of Psychological and Cognitive Sciences at Peking University, and the Science, Technology, Engineering and Mathematics Ethical Review Committee in the University of Birmingham. The fMRI arm of the study was approved by Yale University's Human Research Protection Program IRB and the Commissie Mensgebonden Onderzoek Regio Arnhem-Nijmegen at DCCN..

#### iEEG demographics

Below we describe the characteristics of the iEEG patients (Supplementary Table 25). Also, we provide here further details about the three patients whose behavior fell short of the predefined behavioral criteria (i.e. hits < 70%, FA > 30%), but were nonetheless included in the analysis: one of them kept the response button pressed for most of the time during experiment, the other's low performance was driven by one of the categories only (which the patient reported having difficulty to detect), and the third's performance was very close to the threshold (65%) and had very low FA rate (2%).

| Subject ID | Sex | Age [years] | Handed-ness | Electrode Scheme | Number of Implanted Electrodes | Implant hemisphere | IQ [value, test] | WADA | Seizure Type | Age of Onset | Native Language |
|---|---|---|---|---|---|---|---|---|---|---|---|
| SE103 | F | 49 | L | stereo | 58 | B | 109, FSIQ | N/A | N/A | 27 | English |
| SE106 | F | 18 | R | stereo | 118 | L | >70, FSIQ | N/A | N/A | 12 | English |
| SE107 | M | 24 | R | stereo | 168 | L | >70, FSIQ | N/A | N/A | 13 | English |
| SE108 | F | 16 | R | stereo | 108 | L | >70, FSIQ | N/A | N/A | 12 | English |
| SE109 | F | 50 | R | stereo | 104 | B | >70, FSIQ | N/A | N/A | 45 | English |
| SE110 | F | 15 | R | stereo | 186 | R | >70, FSIQ | N/A | N/A | 10 | English |
| SE112 | F | 17 | R | stereo | 158 | R | >70, FSIQ | N/A | N/A | 7 | English |
| SE113 | F | 26 | A | stereo | 60 | B | >70, FSIQ | N/A | N/A | 19 | English |
| SE115 | M | 17 | R | stereo | 88 | R | >70, FSIQ | N/A | N/A | 6 | English |
| SE118 | M | 11 | R | stereo | 164 | L | >70, FSIQ | N/A | N/A | 8 | English |
| SE119 | M | 29 | R | stereo | 104 | B | >70, FSIQ | N/A | N/A | 28 | Polish |

| | | | | | | | | | | | |
|---|---|---|---|---|---|---|---|---|---|---|---|
| SE120 | M | 12 | L | stereo | 164 | L | >70, FSIQ | N/A | N/A | 1 | English |
| SF102 | F | 30 | R | subdural grid & strips, depths | 133 | L | 98, VCI; 90, POI; 83, WMI; 100, PSI | L | FBTC | 22 | English |
| SF103 | M | 24 | R | subdural grid & strips, depths | 189 | L | 145, VCI; 96, POI; 95, WMI; 86, PSI | predominantly L, mild R contribution | FA | 11 | English |
| SF104 | F | 23 | R | subdural grid & strips, depths | 116 | L | 79, VCI; 62, POI | L | FBTC | 13 | English |
| SF105 | M | 31 | R | subdural grid & strips, depths | 176 | L | 116, VCI; 111, POI; 102, WMI; 114, PSI | L | FBTC | 22 | English |
| SF106 | M | 17 | R | subdural grid & strips, depths | 156 | L | N/A | N/A | FM | 11 | English |
| SF107 | F | 31 | R | subdural grid & strips, depths | 242 | R | 104, VCI | N/A | FIA | 23 | English |
| SF109 | F | 30 | L | subdural grid & strips, depths | 102 | R | 107, VCI; 86, POI; 95, WMI; 92, PSI | predominantly L, mild R contribution | FIA | 3 | English |
| SF110 | F | 17 | R | subdural grid & strips, depths | 174 | L | 89, VCI; 100, POI; 83, WMI; 70, PSI | N/A | FIA, FBTC | 2 | English |
| SF112 | M | 23 | R | subdural strips, depths | 180 | B | 107, VCI; 123, POI; 131, WMI; 100, PSI | L | FIA | 19 | English |
| SF113 | F | 38 | R(converted from L) | subdural grid & strips, depths | 132 | R | 100, VCI; 102, POI; 114, WMI; 102, PSI | L | FA, FIA | 34 | English |
| SF116 | M | 43 | L | stereo | 166 | B | 144, VCI | N/A | FIA | 38 | English |
| SF117 | M | 28 | R | stereo | 174 | B | 105, VCI; 107, POI; 95, WMI; 108, PSI | N/A | FBTC, FIA | 26 | English |
| SF119 | M | 37 | R | subdural grid & | 104 | L | 102, VCI; 98, POI; 97, | L | FBTC | 36 | English |

| | | | | | | | | | | | |
|---|---|---|---|---|---|---|---|---|---|---|---|
| | | | | strips, depths | | | WMI; 108, PSI | | | | |
| SF120 | F | 61 | R | stereo | 79 | L | N/A | N/A | FA, FIA | 44 | English |
| SF121 | F | 50 | R | stereo | 75 | R | 83, VCI; 77, POI; 77, WMI; 76, PSI | L | FIA | 1 | English |
| SF122 | F | 27 | R | stereo | 99 | R | 114, VCI; 96, POI; 94, WMI; 94, PSI | N/A | FH | 14 | English |
| SG101 | M | 40 | R | stereo | 104 | B | 115, FSIQ | N/A | FIA | 24 | English |
| SG102 | M | 49 | R | stereo | 86 | B | 86, FSIQ | N/A | FA, FIA, FBTC | 12 | English |
| SG103 | F | 57 | R | stereo | 76 | B | 77, FSIQ | N/A | FA, FIA, FBTC | 1.5 | English |
| SG104 | F | 48 | R | stereo | 72 | B | 90, FSIQ | N/A | FIA | 30 | English |

N/A – not applicable, F – female, M – male, L – left, R – right, A – ambidextrous, B – bilateral; FSIQ – Full Scale Intelligence Quotient, VCI – Verbal Comprehension Index, POI – Perceptual Organization Index, WMI – Working Memory Index, PSI – Processing Speed Index; FBTC – focal to bilateral tonic-clonic, FIA – focal impaired awareness, FA – focal aware seizures, FM – focal motor, FH – focal hemiclonic seizure.

**Supplementary Table 25**. Characteristics of iEEG patients.

### Author Contributions

In an effort to provide greater transparency and assign the appropriate credit to the authors of this paper, Supplementary Figure 50 illustrates the rated CRediT Contribution Matrix. All listed authors provided a self-assessment of their respective contribution for each of the fourteen CRediT categories on a four-point scale. Here we clearly see each person's total contribution across the project (vertically) and the distribution of work for each role (horizontally). After we accumulated all author's self-rankings, all members were given the opportunity to view a draft of the Contribution Matrix and adjust their own ratings relative to their peers, as well as review and comment on other authors ratings, as a means of normalizing the ratings.

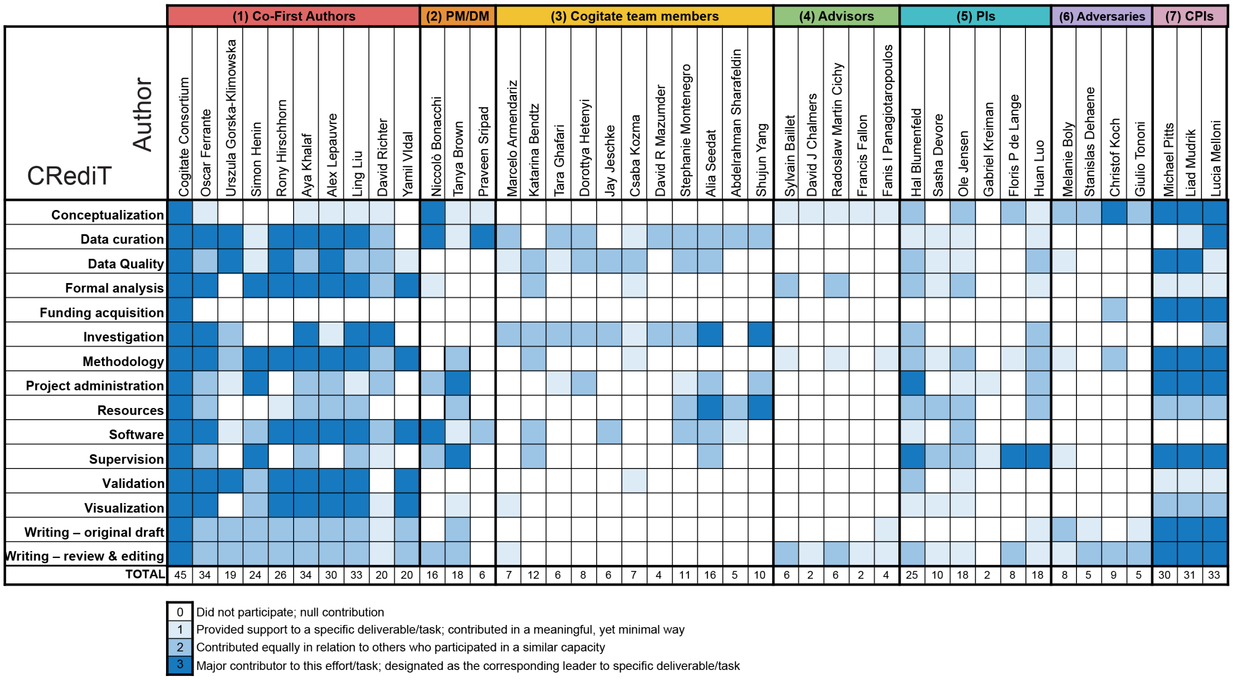

**Supplementary Figure 50.** Contribution matrix towards the work that went into the production of this paper. CRediT roles are listed vertically in the left column, while each author's name is listed horizontally along the top, in the order in which they appear in the author listing. Above the author names is the seven designated author categories, in accordance with the Cogitate Publication Policy v2; (1) co-first authors; (2) Project/Data Managers; (3) additional Cogitate members; (4) Scientific advisors; (5) Site Principle Investigators (PIs); (6) Adversaries; and (7) Centre PIs. Within each author category, authors are listed in alphabetical order. Each member ranked their own contributions for each of the fourteen CRediT roles according to a four-point scale: 0 – null contribution; 1 – support or minimal contributor; 2 – equal or moderate contributor; and 3 – lead or major contributor.
